# Genomes of Poaceae sisters reveal key metabolic innovations preceding the evolution of grasses

**DOI:** 10.1101/2024.11.06.622220

**Authors:** Yuri Takeda-Kimura, Bethany Moore, Samuel Holden, Sontosh K. Deb, Matt Barrett, David Lorence, Marcos V. V. de Oliveira, Jane Grimwood, Melissa Williams, Lori Beth Boston, Jerry Jenkins, Christopher Plott, Shengqiang Shu, Kerrie Barry, David M. Goodstein, Jeremy Schmutz, Matthew J. Moscou, Michael R. McKain, James H. Leebens-Mack, Hiroshi A. Maeda

## Abstract

The grass family (Poaceae, Poales) holds immense economic and ecological significance, exhibiting unique metabolic traits, including dual starch and lignin biosynthetic pathways. To investigate when and how the metabolic innovations known in grasses evolved, we sequenced the genomes of four Poales species, including *Joinvillea ascendens* and *Ecdeiocolea monostachya* representing the sister clade to Poaceae. The *rho* whole genome duplication (ρWGD) in the ancestral lineage for all grasses contributed to the gene family expansions underlying cytosolic starch biosynthesis, whereas an earlier tandem duplication of *phenylalanine ammonia lyase* (*PAL*) gave rise to *phenylalanine/tyrosine ammonia lyase* (*PTAL*) responsible for the dual lignin biosynthesis. Integrated functional genomic and biochemical analyses of grass relatives further revealed the molecular basis of key metabolic innovations predating the evolution of grasses.

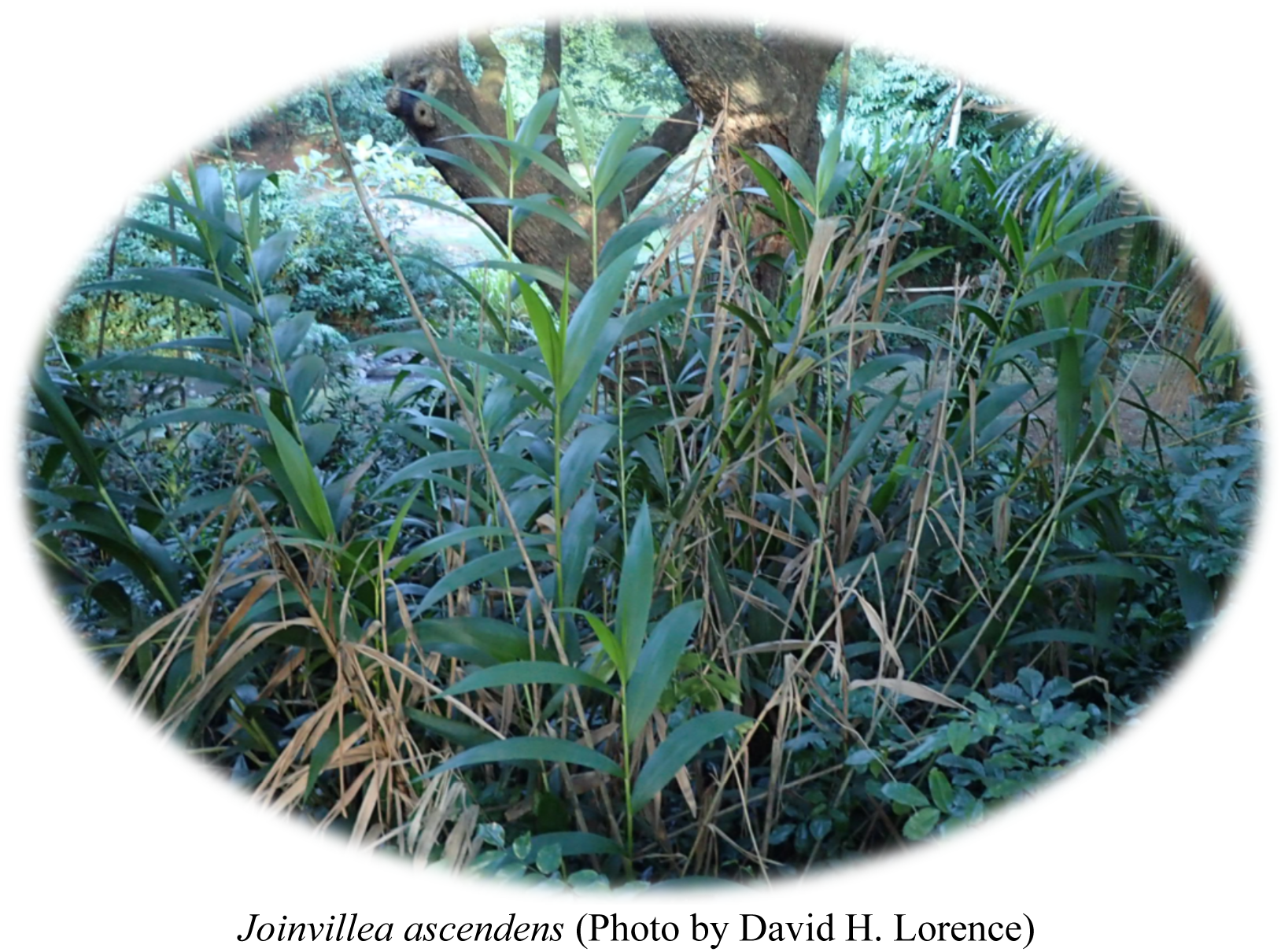

## Introduction

The grass family, Poaceae, emerged around 100 million years ago (Mya) within the monocot order Poales (*1*) and contains more than 11,000 species (**Fig. 1A**) (*1–3*). Grasses act as foundational primary producers for diverse ecosystems across the globe and contain many of the most economically important crops. Cereals, such as rice, wheat, and maize, provide more than 50% of calories for humans worldwide (*4*), while bioenergy crops, such as sugarcane and sorghum, produce abundant sugars and lignocellulosic biomass. Grasses exhibit a series of unique morphological and physiological traits, such as the highly specialized inflorescence, the ear, which has been a target for domestication to reduce shattering (*5–7*). Grasses also exhibit striking metabolic innovations, such as their ability to synthesize starch in both plastids and cytosol, leading to the starch-rich endosperm of grass seeds, e.g., grains and kernels (*8*). Grasses also have two entry pathways to synthesize lignin (**Fig. 1B**), the major cell wall phenolic polymer, which represents up to 30% of grass dry mass (*9*) and provides renewable aromatic feedstocks (*10–12*). Lignin is typically synthesized from the aromatic amino acid L-phenylalanine (Phe) via Phe ammonia-lyase (PAL) enzyme (**Fig. 1B**), but grasses can additionally utilize L-tyrosine (Tyr) to make lignin via the bifunctional Phe/Tyr ammonia-lyase (PTAL) (*13*, *14*). The PTAL pathway contributes to nearly half of grass lignin biosynthesis (*13*, *14*), potentially facilitating their rapid growth while depositing substantial lignin in their scattered vascular bundles (**Fig. 1B**). However, the evolutionary history and molecular basis of grass metabolic innovations are largely unknown. Recent advancement in genome sequencing has permitted tracing whole genome duplications (WGD), gene family evolution, and gene innovations (*15–20*). Several key features of monocot genome evolutions include numerous polyploidy events (*21–23*), including the *rho* WGD (ρWGD) which occurred within the stem lineage leading to Poaceae (**Fig. 1A**) (*3*, *19*, *23*). The ρWGD event facilitated the evolution of complex traits and an increased speciation rate (*24*, *25*) in association with proliferation of MADS-box genes and diversification of the grass floral spikelet (*6*). The majority of sequenced Poales genomes are found in the Poaceae, especially cereal crops due to their agricultural significance (*26–29*). However, genome assemblies are currently unavailable from the sister clade to Poaceae, which includes two families, Ecdeiocoleaceae (N=3) and Joinvilleaceae (N=4) (*3*, *30*).

**Fig. 1:**
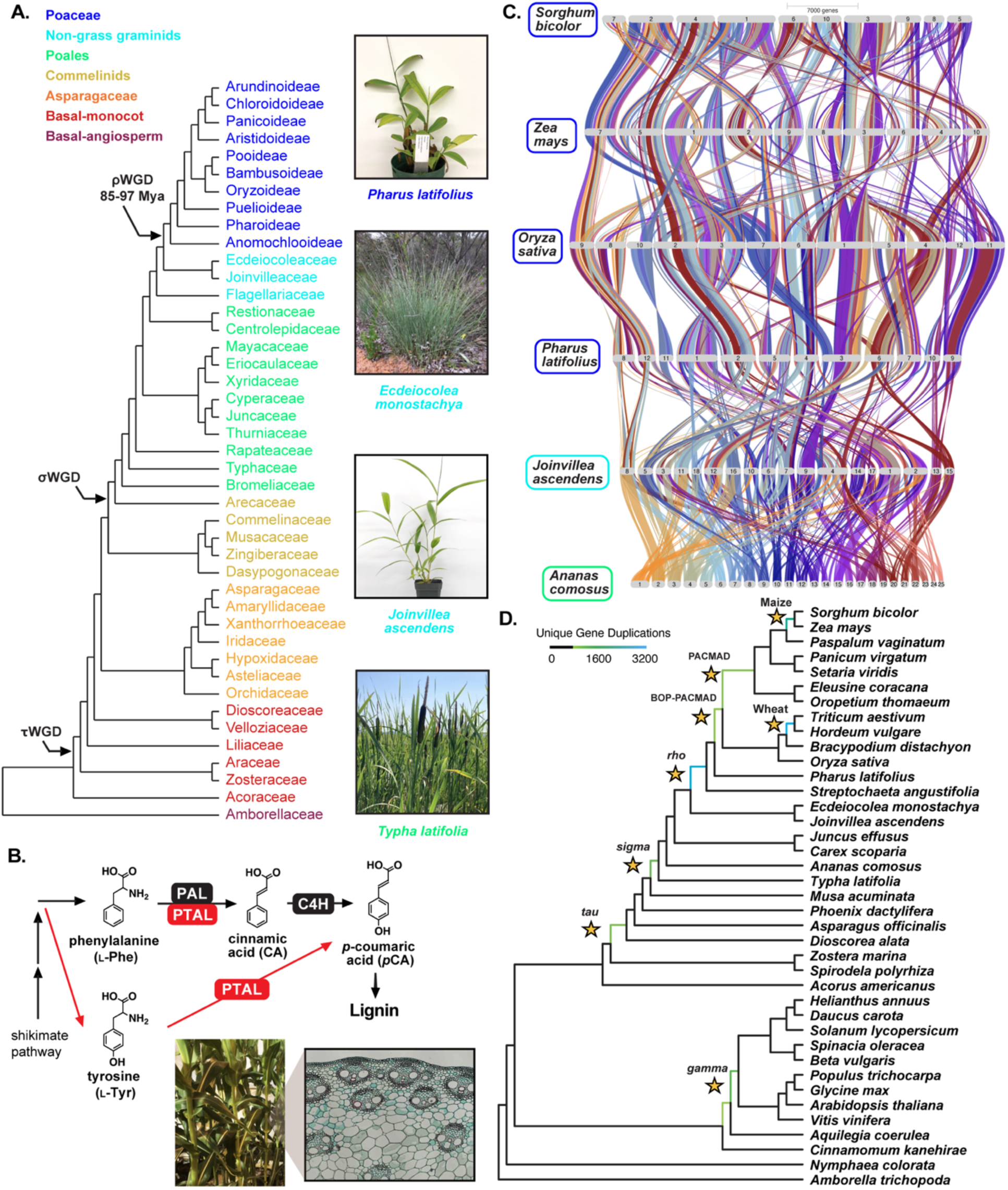
Sequencing genomes of Poaceae sisters to trace the evolutionary history of grass metabolic innovations. **(A)** The family tree of monocot plants showing three ancient whole genome duplication (WGD) events and the images of three plant species sister to core grasses. **(B)** Grasses have a unique metabolic trait, the dual entry pathways to synthesize lignin and phenylpropanoids from both L-phenylalanine (Phe) and L-tyrosine (Tyr) due to the presence of Phe/Tyr ammonia lyase (PTAL). A stem cross section of *Zea mays* showing scattered vascular bundles (Image by Michael Clayton). **(C)** The whole genome comparisons of gene order (synteny) across representative Poaceae and Poales. **(D)** Mapping of gene duplications counted in gene trees using the Phylogenetic Placement of Polyploidy Using Genomes (PUG) toolkit highlights ρWGD in the last common ancestor of all grasses (Poaceae), as well as sigma (σ) and tau (τ) WGDs.

Here we document the utility of these sister genomes for comparative genomics and facilitate identification of genome and molecular features associated with grass-specific functional traits (**Fig. 1A**). Reference-quality genomes were first generated for *Joinvillea ascendens* Gaudich. ex Brongn. & Gris and *Ecdeiocolea monostachya* F.Muell., representing the sister clade to the entire grasses, along with *Pharus latifolius* L., a non-core Poaceae grass species, and the cattail species *Typha latifolia* L. representing the sister lineage to all other families in the order Poales (*3*, *30*, *31*). Our integrated genomic and biochemical analyses of grass relatives in the order Poales unveiled the evolutionary history and molecular basis of key metabolic innovations that predated the emergence of grasses. The study highlights the critical roles of both WGDs and tandem gene duplications underlying evolutionary innovations and identifies a gene editing target that can introduce the dual lignin entry pathway in non-grass plants. The Poales genomes obtained here offer valuable resources for dissecting diverse and complex grass traits uniquely evolved in this critical plant family.

## Results

### Genomes of Joinvillea ascendens, Ecdeiocolea monostachya, Pharus latifolius, and Typha latifolia provide critical resources to study grass evolution

*Ecdeiocolea monostachya* is a wild perennial native to Western Australia and one of three species within the Ecdeiocoleaceae (**Fig. 1A**). The flow cytometry estimated between 38 to 42 chromosomes and haploid genome size of 980 Mb (4C of ∼4.0 pg), consistent with an outcrossing species with an estimated chromosome number (2N) of 38 (*32*). The *E. monostachya* genome was assembled by HiFiAsm using PacBio HiFi long-reads, and gene models were predicted using *ab initio*, homology, and evidence-based gene models (see Methods). The final haploid consensus assembly of a heterozygous, diploid *E. monostachya* isolate EM_001 encompasses 897 Mb over 1,114 scaffolds with N50 of 1.7 Mb and a total of 27,801 genes, with 97.9% of the Embryophyte BUSCO genes (**Table 1**).

**Table 1.**
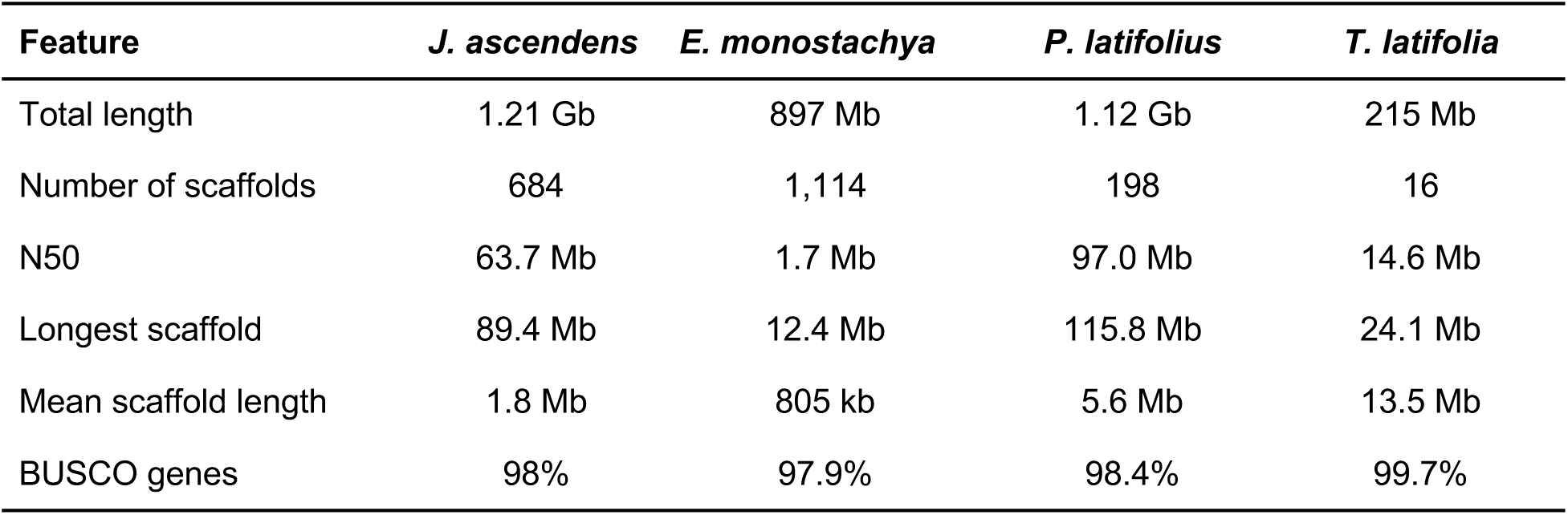
Summary statistics of the genome assembly of *J. ascendens*, *E. monostachya*, *P. latifolius,* and *T. latifolia*.

| Feature | *J. ascendens* | *E. monostachya* | *P. latifolius* | *T. latifolia* |
| --- | --- | --- | --- | --- |
| Total length | 1.21 Gb | 897 Mb | 1.12 Gb | 215 Mb |
| Number of scaffolds | 684 | 1,114 | 198 | 16 |
| N50 | 63.7 Mb | 1.7 Mb | 97.0 Mb | 14.6 Mb |
| Longest scaffold | 89.4 Mb | 12.4 Mb | 115.8 Mb | 24.1 Mb |
| Mean scaffold length | 1.8 Mb | 805 kb | 5.6 Mb | 13.5 Mb |
| BUSCO genes | 98% | 97.9% | 98.4% | 99.7% |

*Joinvillea ascendens* is native to the Hawaiian Islands and Oceania and one of four species within Joinvilleaceae (**Fig. 1A**). The genome of *J. ascendens* was sequenced to 125x coverage yielding a scaffold assembly with 18 chromosomes with a genome size of 1.21 Gb with N50 of 63.7 Mb and 29,122 protein coding genes, with 98% BUSCO score (**Table 1**). The *P. latifolius* genome was sequenced to a depth of 44x coverage yielding a chromosome level assembly spanning 1.12 Gb over 198 scaffolds. Gene models were inferred for 32,144 protein-coding genes, including 98.4% of the Embryophyte BUSCO genes (**Table 1**). As compared to the *P. latifolius* genome released by Ma et al. (*33*), the use of PacBio HiFi reads generated a reduced number of contigs (222 vs. 535) and slightly larger final genome size (1.12 Gb vs. 1.00 Gb). The *T. latifolia* genome was sequenced to depths of 74.70x coverage resulting in a chromosomal assembly including 215 Mb spanning the expected 15 chromosomes and one additional 54 Kb scaffold. A total of 22,107 protein-coding gene models were characterized in the *T. latifolia* genome including 99.7% of the Embryophyte BUSCO genes (**Table 1**). These three genomes were assembled using HiFiAsm, which integrates PacBio HiFi and Hi-C data, and polished using RACON (*34*). Scaffold orientation, ordering, and joining were based on Hi-C contact mapping, while evidence-based gene annotations were performed using Illumina RNAseq and PacBio IsoSeq reads (See Methods).

Variation in chromosomal structure across Poales genomes was elucidated using GENESPACE (*35*), which combines sequence similarity with gene order to robustly estimate orthology. Synteny patterns were consistent with the previously estimated timing of the ρWGD after divergence of Poaceae and its sister clade (**Fig. 1A**). A riverine plot reveals rearrangements of chromosomal blocks over the 120 My history of Poales (**Fig. 1C**). Gene family divergence and conservation were analyzed using 39 species with complete genomes, representing monocots with taxa focused in Poales along with representative eudicots, Magnoliids, and the basal angiosperms (ANA grade) (**table S1**). A Phylogenetic Phylogenetic Placement of Polyploidy Using Genomes (PUG) analysis (*3*, *30*, *31*) estimated a total of 2,493,258 putative paralog pairs from the 10,255 orthogroup trees across all taxa. After filtering for well supported nodes in the gene trees (see Methods), 21,105 putative paralog pairs provided support for unique duplication events (**Fig. 1D**). The comparisons between gene duplications on specific branches and previously characterized ancient polyploidy events showed that the highest number (3,200) of gene duplications was found on the branch leading to the last common ancestor of wheat (*Triticum aestivum*) and barley (*Hordeum vulgare*). The *rho* event—previously placed just prior to the diversification of core-grasses (*3*, *7*, *36*)—was supported with 3,004 gene duplications placed on the branch leading to the last common ancestor of all grasses (**Fig. 1D; table S2**). We also found 1122, 972, and 2196 gene duplications associated with the *sigma* WGD event (*37*), the *tau* WGD event (*3*, *38*), and the maize allopolyploid event (*37*, *39*), respectively (**table S2**). Additionally, 868 and 936 unique gene duplications were mapped to before the divergence of the BOP-PACMAD clades and the PACMAD clade, respectively (**table S2**), likely representing either homoeologous exchange between *rho* duplications (*40*) or possibly other types of duplication. Finally, we see evidence of the *gamma* event (1,212 duplications) prior to the divergence of *Vitis* from other Pentapetalae (*41*, *42*), as well as 803 unique duplications that map to branch leading to the common ancestor of all eudicots (**table S2**). In sum, the genomes of *E. monostachya*, *J. ascendens*, *P. latifolius,* and *T. latifolia* enable improved resolution of genome evolution with the grasses and prior to their origin.

### Duplication of genes involved in starch and fatty acid biosynthesis during grass evolution

Starch stores carbon and energy for maintaining metabolism and growth (*43*, *44*). Grasses have starch-rich seeds that support rapid seedling establishment (*45*) and provide the major source of calories consumed by humans (e.g., cereal grains and kernels). Besides plastids where starch is stored and typically made (*46*), grass endosperms can synthesize starch also in the cytosol (*8*, *47*) (**Fig. 2A**). To investigate the origin of the cytosolic starch biosynthesis, we constructed phylogenetic trees of three genes required for the cytosolic starch biosynthesis—ADP-glucose pyrophosphorylase (AGPase) large and small subunits (LSU and SSU, respectively) and ADP glucose transporter (**Fig. 2A**). Grasses have four types of AGPase LSUs with the duplication of Type 3 AGPase LSUs that likely generated Type 2 cytosolic AGPase LSUs (*8*). Consistent with this notion, our phylogeny showed two distinct, well-supported Type 3 and Type 2 AGPase LSU clades both having sequences from *P. latifolius* and another non-core grass, *Streptochaeta angustifolia* (*7*). In contrast, *J. ascendens* and *E. monostachya* sequences were found in their outgroup (**Fig. 2B; fig. S1A**), suggesting that the duplication event occurred within grasses. Similarly, AGPase SSUs duplicated within grasses, giving rise to Type 1 AGPase SSUs (**Fig. 2C; fig. S1B**), which is dual localized but reside in the cytosol in the endosperm, unlike Type 2 AGPase SSUs targeted to the plastids in leaves (*48*). The plastidial ADP glucose transporter, required to import ADP-glucose into the plastids (**Fig. 2A**) (*8*), also underwent gene duplication within the common ancestor of all grasses (**Fig. 2D**), giving rise to the plastidial ADP glucose transporter within the Type 2 Plastidial Adenine Nucleotide Transporters (PANTs) (*49*). The long branch leading to the grass-specific plastidial ADP glucose transporter suggests potential positive selection for neofunctionalization. Notably, however, *J. ascendens* and all grasses, including *S. angustifolia* and *P. latifolius*, had both Type 1 and Type 2 PANTs (**Fig. 2D**). This finding suggests that there was an earlier duplication event preceding the emergence of grasses, giving rise to the Type 2 PANTs, from which the plastidial ADP glucose transporter later evolved.

**Fig. 2.**
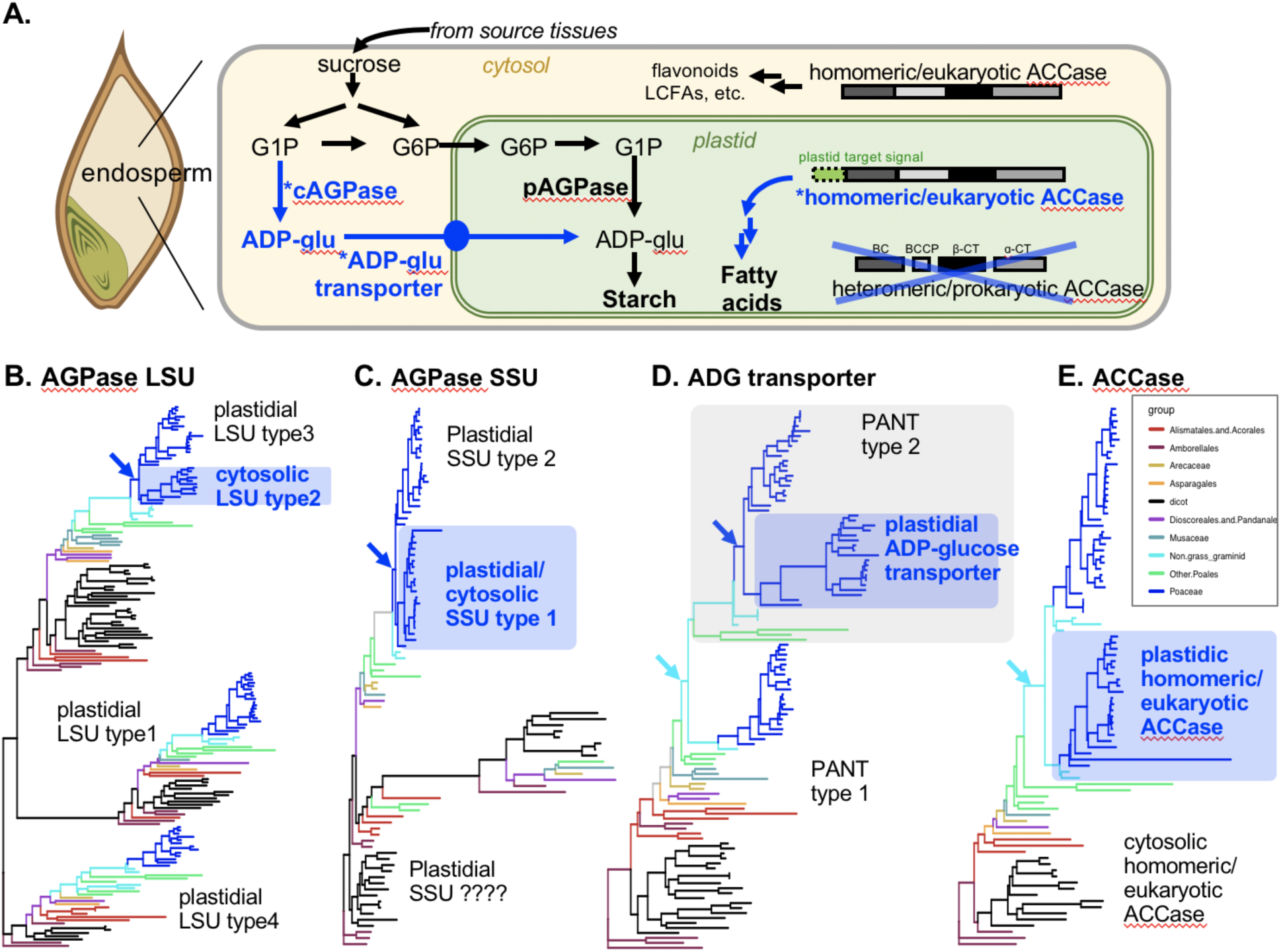
Alteration of starch and fatty acid biosynthesis during the grass evolution. **(A)** In the endosperm of grasses, starch can be synthesized in both cytosol and plastids due to the presence of cytosolic ADP-glucose pyrophosphorylase (AGPase) and ADP-glucose (ADG) transporter. Grasses also have unique fatty acid biosynthetic enzymes, homomeric eukaryotic-type acetyl-CoA carboxylase (ACCase) in the plastids, instead of heteromeric prokaryotic-type ACCase typically found in other plant plastids. **(B to E)** The phylogenetic trees of AGPase large subunit (LSU, **B**), small subunit (SSU, **C**), ADG transporter **(D)**, and ACCase **(E)** of grasses, Poales, as well as other monocots and angiosperms. Blue and gray boxes denote clades specific to Poaceae and non-grass graminids that likely contributed to the evolution of their unique starch and fatty acid biosynthetic pathways. Blue and sky-blue arrows indicate the underlying duplication events that occurred after and before the emergence of grasses, respectively. The original trees with branch supports and labels are shown in **fig. S1**.

Grasses also have a unique biochemical feature for fatty acid biosynthesis. Plants typically have heteromeric prokaryotic-type and homomeric eukaryotic-type acetyl-CoA carboxylase (ACCase) in the cytosol and plastids, respectively, whereas grasses have homomeric eukaryotic-type ACCase in both cytosol and plastids (*50*, *51*) (**Fig. 2A**). This makes grasses sensitive to certain herbicides that inhibit homomeric but not heteromeric ACCase (*50*, *52*). Notably, the homomeric eukaryotic-type *ACCase* gene duplicated in ancestral non-grass graminids, giving rise to the plastidic homomeric eukaryotic-type ACCase in grasses as well as *J. ascendens* and *E. monostachya* (**Fig. 2E**). These results together revealed the evolutionary history of metabolic genes underlying the unique metabolic traits of grasses, some of which have already appeared *before* the emergence of grasses and ρWGD.

### PTAL evolved within a common ancestor of grasses and non-grass graminid *Joinvillea*, before ρWGD and the appearance of grasses

Utilizing these novel genomic resources, we further investigated the evolutionary history of the dual lignin entry pathways, uniquely found in grasses (**Fig. 1B**) (*13*, *14*, *53*). Synteny analyses revealed that *J. ascendens* has two copies of *PAL/PTAL* homologs (Joasc.05G060400.1, Joasc.05G060500.1) located in tandem within the same syntenic block at the chromosome 5 (**Fig. 3A**), whereas the corresponding syntenic block in *Ananas comosus* (pineapple) has just one *PAL/PTAL* homolog. Within grasses, the *PAL/PTAL* syntenic block was further duplicated in association with the ρWGD, and *PAL* genes in one of the two duplicated blocks were expanded in core grasses (**Fig. 3A**). One *PAL/PTAL* copy within these syntenic blocks in grasses encodes previously characterized or predicted PTAL enzymes (*14*). Analyses of *Ks* values (synonymous substitutions) for genes within and between *PAL/PTAL* syntenic blocks further supports the timing of the ancestral *PAL/PTAL* tandem duplication before ρWGD **(fig. S2)** and the origin of Poaceae.

**Fig. 3:**
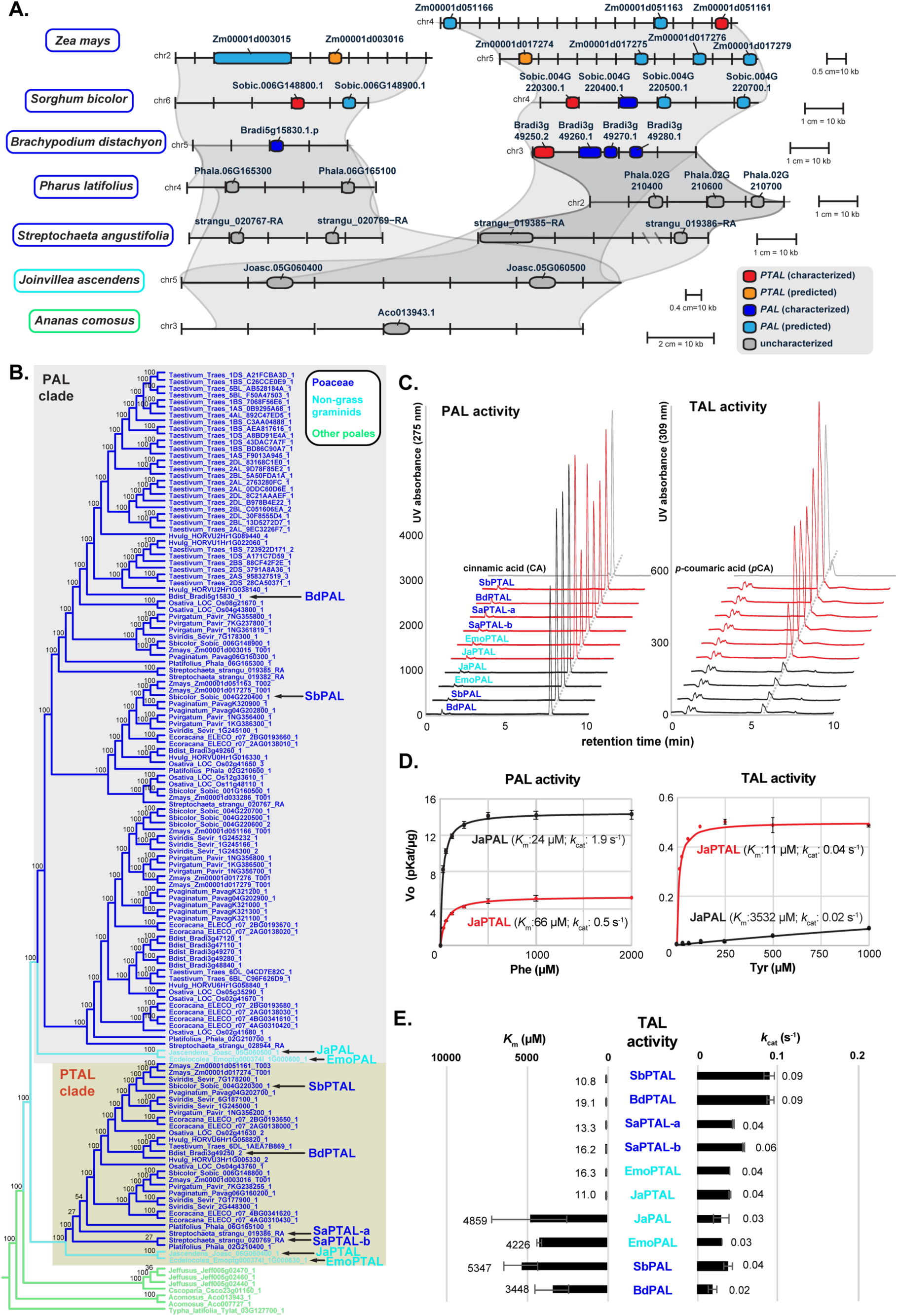
The PTAL enzyme evolved before the emergence of grasses through tandem duplication predating ρWGD. **(A)** *PAL/PTAL* gene synteny is shown for *A. comosus, J. ascendens, S. angustifolia, P. latifolius, B. distachyon, S. bicolor,* and *Z. mays*, illustrating the gains and losses of PAL/PTAL synteny across Poales. The *PAL* gene is duplicated and neofunctionalized to PTAL in the ancestor of *J. ascendens*. The *PAL*/*PTAL* pair is then duplicated within grasses, likely due to the ρWGD, and again in *Z. mays*, likely due to the *Z. mays* WGD. **(B)** Phylogenetic tree of *PAL*/*PTAL* genes, with focus on Poales species, generated with RAxML-NG and reconciled with TreeSolve. *PAL*/*PTAL* homologs characterized in this study are highlighted with black arrows. The full-scaled phylogenetic tree is shown in **fig. S5**. **(C)** HPLC trace of PAL and TAL activity assay for PTALs/PALs from *S. bicolor* (SbPTAL and SbPAL), *B. distachion* (BdPTAL and BdPAL), *S. angustifolia* (SaPTAL-a and SaPTAL-b), *J. ascendens* (JaPTAL and JaPAL) and *Ecdeiocolea monostachya* (EmoPTAL and EmoPAL). See **fig. S6** for additional traces of their control reactions. **(D)** Michaelis-Menten curves of TAL and PAL assay for JaPTAL and JaPAL. **(E)** Kinetic parameters of TAL activity for PTALs/PALs. Data are means ± SD (*n* = 3).

In contrast to the findings of the synteny analysis, the Maximum Likelihood (ML) gene tree for the *PAL/PTAL* gene family resolved *PAL* homologs from *J. ascendens*, and *E. monostachya* on a grade leading to a clade with all *PTAL* homologs from *J. ascendens*, *E. monostachya* and grasses (**figs. S3 and S4**). Using the TreeSolve dynamic programming algorithm for species-tree-aware estimation of the *PAL/PTAL* tree while accounting for the gene duplication-and-loss process (*54*), monophyly of the *PAL* and *PTAL* orthologs was recovered with a support values of 100% for each clade **(Fig. 3B, fig. S5)**. Consistent with the inference of an ancestral tandem duplication based on the synteny analysis (**Fig. 3A**), *J. ascendens* Joasc.05G060500.1 and *E. monostachya* Emoptg000374l_1G000600.1 genes formed a clade sister to a clade with all grass *PAL* orthologs, and Joasc.05G060400.1 and Emoptg000374l_1G000630.1 were in a clade sister to all grass *PTAL* genes (**Fig. 3B, fig. S5**). In sum, these analyses implicate divergence of *PAL* and *PTAL* and neofunctionalization of *PTAL* following a tandem duplication in a graminid ancestor of the grasses. The His140 residue, located at the substrate binding pocket, was previously shown to be critical for the Tyr substrate recognition in the bacterial TAL enzyme (*55*). We noted that the corresponding His140 residues were present in PTAL orthologs of all grasses, including both of the *S. angustifolia* enzymes (strangu_020769−RA and strangu_019386−RA), as well as one of each enzyme from *J. ascenden*s (Joasc.05G060400.1) and *E. monostachya* (Emoptg000374l_1G000630.1), suggesting that these proteins may be functional PTAL enzymes. To test this hypothesis experimentally, we cloned, expressed, and purified the recombinant enzymes of PAL/PTAL orthologs from *S. angustifolia*, *J. ascendens*, and *E. monostachya*, along with PALs and PTALs from *Sorghum bicolor* (SbPAL and SbPTAL) and *Brachypodium distachyon* (BdPAL and BdPTAL) as positive controls (*14*, *56*). All of these purified enzymes showed robust PAL activity, efficiently converting Phe to cinnamic acid (CA) as confirmed by HPLC analysis (**Fig. 3C**), unlike negative controls (i.e., boiled enzyme or no substrate controls, **fig. S6**). Efficient TAL activity, the production of *p*CA from Tyr, was observed from SbPTAL, BdPTAL, strangu_020769−RA, strangu_019386−RA, Emoptg000374l_1G000630.1, and Joasc.05G060400.1, whereas much lower TAL activity was detected from the remaining enzymes (**Fig. 3C**). These results showed that the PAL/PTAL orthologs with His140 are bifunctional PTAL enzymes having both PAL and TAL activity. Therefore, we named both of *S. angustifolia* enzymes and one of *E. monostachya*, and *J. ascendens* enzymes having His140 as SaPTAL-a, SaPTAL-b, EmoPTAL, and JaPTAL, whereas another one of *E. monostachya*, and *J. ascendens* enzymes with Phe140 as EmoPAL and JaPAL, respectively (**Fig. 3B**).

Enzyme kinetic analyses further revealed that the apparent *K*m of these PTAL enzymes for Tyr were 11 to 19 μM, whereas those of PAL enzymes were much higher (3449-6211 μM) and hence inefficient in utilizing Tyr. The *k*cat values of the TAL activity were approximately twice higher for PTALs than for PALs (**Fig. 3, D and E**; **table S2**). Consequently, the catalytic efficiency (*k*cat/*K*m) of TAL activity for PTAL enzymes were much higher (485-fold in average) than those for PAL enzymes (**Fig. 3, D and E; table S3**). These quantitative data further support that *S. angustifolia*, *E. monostachya*, and *J. ascendens* have at least one PTAL enzyme having strong TAL activity. We also noted that grass PTAL enzymes showed higher *K*m toward Phe (150-227 μM) than graminid PTALs (64-66 μM) (**table S3**), leading to ∼3.1-fold higher TAL/PAL activity ratio for PTALs from grasses than non-grass graminids (**fig. S7A**). These experimental data demonstrate that the bifunctional PTAL enzymes emerged within the common ancestor of grasses and non-grass graminids (e.g., *J. ascendens*), and later gained higher TAL/PAL ratio in grasses.

### Ile112 and His140 are critical for acquisition of the TAL activity in graminid PTALs

To experimentally test the role of His140 for the acquisition of TAL activity, site-directed mutagenesis was conducted on the PAL and PTAL enzymes of grasses and non-grass graminids characterized above. The conversion of Phe140 to His in PAL enzymes (e.g., JaPAL^F140H^) increased overall TAL activity (9.7-fold in *k*cat/*K*m average) with significant reduction of *K*m values towards Tyr (**table S3**). The reciprocal mutants of PTAL enzymes (e.g., JaPTAL^H125F^) also exhibited decreased TAL activity (0.01-fold *k*cat/*K*m in average) with significant increase in *K*m towards Tyr (**fig. S7B; table S3**). These results support the importance of His140 for TAL activity, consistent with prior studies (*55*, *57*); however, the introduction of His140 itself was not *sufficient* to convert PALs into PTALs. The catalytic efficiency of TAL activity (*k*cat/*K*m) of PAL^F140H^ mutants was still much lower (∼10% in average) than that of wild-type PTALs, because of the much higher *K*m values (222-450 μM) than wild-type PTALs (11-19 μM) (**fig, S7B; table S3**). Furthermore, PTAL^H140F^ mutants still showed higher TAL activity than that of wild-type PALs (e.g., *K*m value of 531-765 vs. 3448-6211 μM). Thus, unlike the bacterial TAL enzyme (*57*), other residues besides the His140 are required for the acquisition of strong TAL activity in PTALs of grasses and closely-related non-grass graminids.

To search for the additional residue(s) required for the efficient TAL activity, we first conducted a positive selection analysis of *dN/dS* (nonsynonymous to synonymous substitution rate) using PAML (*58*). A total of 30 positively selected sites were identified between the PTAL vs. PAL clade of the graminids with a probability of >0.7 (**table S4**). A phylogeny-guided amino acid sequence comparison (*59*) utilizing the phylogenetic distribution of the functional PAL and PTALs (**Fig. 3**) identified 16 residues, besides His140, that are highly conserved in PTAL enzymes (**Fig. 4A; fig. S8**), all of which were under positive selection (**table S4**). Eight residues (magenta in **Fig. 4A**) were highly conserved within PAL and PTAL groups but distinct between these two groups; the additional 8 residues (purple in **Fig. 4A**) were highly conserved only among PTALs and variable in different PALs (**Fig 4A; table S4**). Based on a homology model of JaPAL generated using the well-characterized parsley PAL structure as a template (PDB:6F6T) (*60*), most of these 16 residues were found to be located near the active site, except for a few peripheral residues highlighted in purple (**Fig. 4B**).

**Fig. 4:**
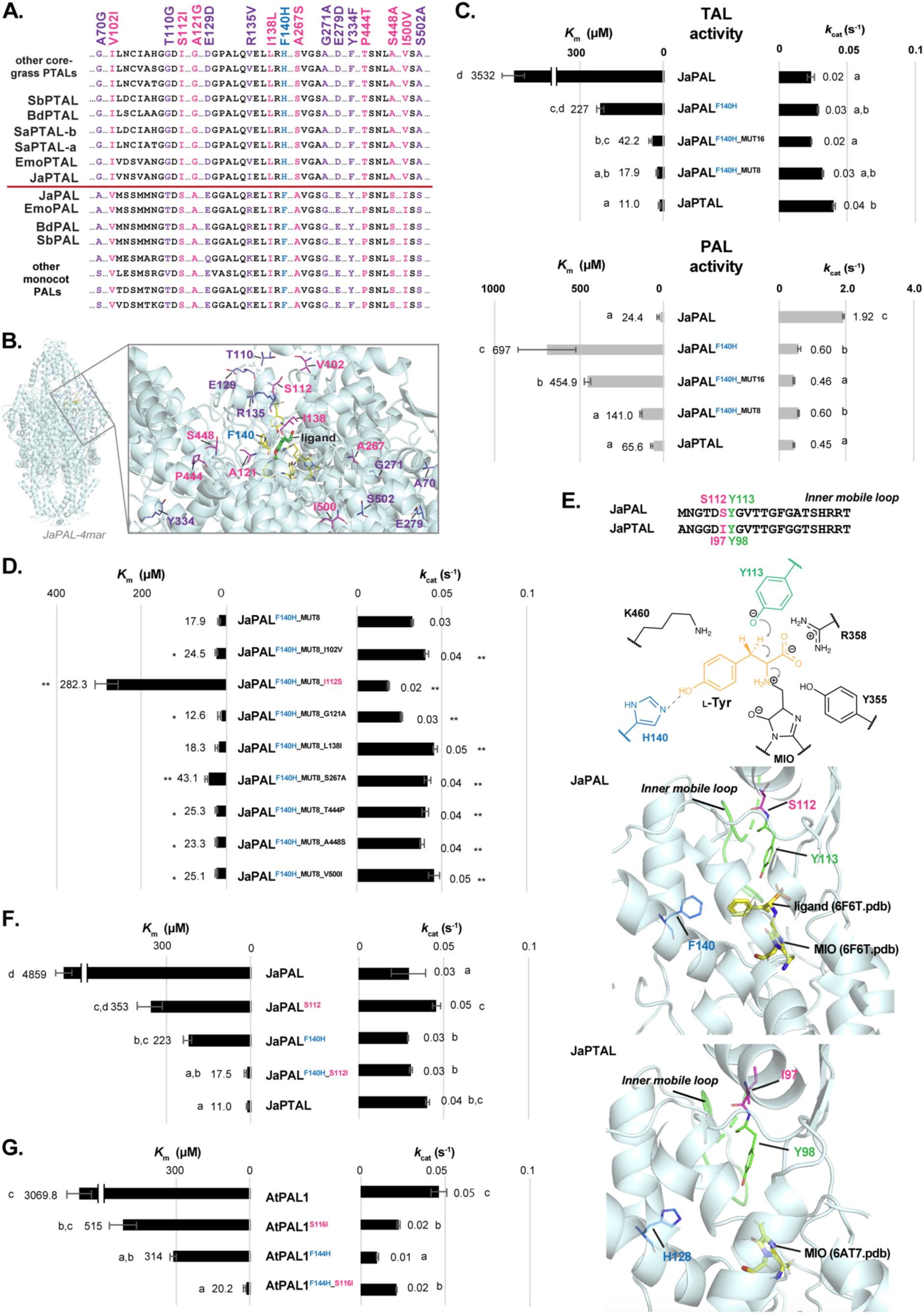
Two mutations, F140H and S112I, convert PAL into PTAL enzymes. **(A)** Partial amino acid alignment highlighting residue His/Phe140 (cyan), which is essential for recognizing substrate phenylalanine and tyrosine, and residues highly conserved uniquely in PTALs (magenta: conserved in both PALs and PTALs, but in distinct ways; purple: conserved among PTALs but not PALs). Full-scaled alignment is shown in **fig. S8**. **(B)** Protein structure model of JaPAL highlighting selected residues. The structures of the substrate ligand and MIO cofactor are derived from the template 6F6T.pdb. Residues important for enzyme activity are displayed in yellow. **(C)** Kinetic parameters of TAL and PAL activity for JaPAL, JaPAL with mutation at His140 site (JaPAL^F140H^) and additional mutations (JaPAL^F140H_MUT8^ and JaPAL^F140H_MUT16^), and JaPTAL. **(D)** Kinetic parameters of TAL activity for JaPAL^F140H_MUT8^ with mutation on 8 of each residue back to PAL type. **(E)** Potential mechanism of TAL reaction (*77*) and roles of His140 and Ile112 for the catalysis of the PTAL enzyme. **(F)** Kinetic parameters of TAL activity for JaPAL with site-directed mutation at His140 site (JaPAL^F140H^) and Ile112 site (JaPAL^S112I^). **(G)** Kinetic parameters of TAL activity for Arabidopsis AtPAL1 with mutation at His140 (AtPAL1^F144H^) and Ile112 (AtPAL1^S116I^). Data are means ± SD (*n* = 3). Different letters indicate significant differences (*p* < 0.05).

To test functions of these residues, JaPAL^F140H^ was further mutated at these 8 magenta residues or both magenta and purple (total 16) residues from PAL- to PTAL-type (**table S4**), generating JaPAL^F140H_MUT8^ and JaPAL^F140H_MUT16^, respectively. The apparent *K*m value of JaPAL^F140H_MUT8^ (17.9 μM) became significantly improved than that of the JaPAL^F140H^ single mutant (222.7 μM) and approached closely to that of wild-type JaPTAL (10.9 μM) with the similar *k*cat (**Fig. 4C; table S3**). The JaPAL^F140H_MUT16^ mutant also showed significantly improved *K*m (42.2 μM) for TAL activity than JaPAL^F140H^ (and JaPAL wild-type) but to a lesser extent than JaPAL^F140H_MUT8^ (17.9 μM, **Fig. 4C**). Thus, these results demonstrate that some of these 8 magenta residues, besides His140, are involved in the efficient TAL activity of non-grass graminid PTALs. To determine which of the eight magenta residues are essential in the conversion of PAL to PTAL enzyme, each one of these residues on JaPAL^F140H_MUT8^ was mutated back to the PAL type, one by one. The substitution of seven out of eight residues had no to minor impacts on the overall TAL activity (**Fig. 4D**). In contrast, when the I112S substitution was introduced to JaPAL^F140H_MUT8^ to make JaPAL^F140H_MUT8_I112S^, TAL activity was significantly decreased because of the drastic increase of *K*m and slight reduction of *k*cat (**Fig. 4D; table S3**). Based on the structure models of JaPAL and JaPTAL proteins generated with parsley PAL and sorghum PTAL structures (*56*, *61*), respectively, as a template (**Fig. 4E**), the Ser/Ile112 is not in direct contact with the substrate but is located next to the Tyr113/98 (for PAL/PTAL), a critical proton acceptor for the catalysis. Therefore, the Ile112 residue of the PTAL enzyme near the active site appears to be crucial for the efficient TAL activity.

### F140H and S112I substitutions are sufficient to change plant PALs into bifunctional PTALs

To further test the role of the Ile112 residue, the reciprocal S112I mutation was introduced to the JaPAL wild-type and JaPAL^F140H^ single mutant to generate the JaPAL^S112I^ single and JaPAL^F140H_S112I^ double mutants, respectively. While *k*cat was not drastically affected by these mutations, *K*m of JaPAL^F140H_S112I^ for Tyr (17.5 μM) became significantly lower than those of the wild-type JaPAL (4859 μM) and the single mutants JaPAL^F140H^ and JaPAL^S112I^ (223 μM and 354 μM, respectively) and reached to the level of wild-type JaPTAL (11.0 μM) (**Fig. 4F**). Therefore, Ile112 is essential for gaining the efficient TAL activity. Strikingly, introduction of these two amino acid substitutions equivalent to F140H and S112I in a distantly-related PAL from *Arabidopsis thaliana* (*55*, *62*) (AtPAL1^F144H_S116I^) also conferred efficient TAL activity. The AtPAL1^F144H_S116I^ mutant showed dramatic reduction in its *K*m towards Tyr (20.2 μM) as compared to wild-type AtPAL1 (3070 μM) and its single mutants, AtPAL1^F144H^ and AtPAL1^S116I^ (314 μM and 515 μM, respectively). Overall, the kinetic behavior of the AtPAL1^F144H_S116I^ and JaPAL^H140F_I112S^ double mutants were similar (**Fig. 4, F and G; table S3**). Thus, the introduction of two key residues Ile112 and His140, identified from the PTAL evolutionary analysis in Poales, can alter monofunctional PALs into bifunctional PTALs even in a distantly-related plant PAL enzyme.

## Discussion

Genomes of closely related species in sister lineages provide critical resources to dissect the genetic and molecular basis of unique traits that evolved in a target organism(s). For example, the chimpanzee and bonobo genomes have accelerated our understanding of the evolution of human traits (*63*, *64*). In the current study, integrated genomic and biochemical analyses of Poales that are close-relative to core-grasses unveiled the evolutionary origin and molecular basis of key metabolic innovations that predated the emergence of grasses. While some genes involved in the cytosolic starch biosynthetic pathway emerged through the ρWGD, a preceding duplication into PANT Type 1 and 2 potentiated the evolution of ADP-glucose transport across the plastid membrane (**Fig. 2E**). Strikingly, the dual lignin entry pathway catalyzed by bifunctional PTAL enzyme, which was believed to be specific to grasses (*14*), actually emerged *before* the evolution of grasses at the common ancestor of Joinvilleaceae, Ecdeiocoleaceae, and Poaceae. Although WGD-associate diversification has been attributed to sub/neofunctionalization of duplicates leading to morphological and functional innovations (*65*, *66*), our results uncovered that tandemly duplicated gene families also contribute to evolutionary innovations, through subsequent neofunctionalization and further expansion by later WGD (**Fig. 3**).

Precise determination of the timing of the PTAL evolution also revealed its underlying molecular basis. A novel residue Ile112 responsible for the transition from PAL to PTAL function is located at the ‘inner mobile loop’ (**Fig. 4E**). Based on prior structure analyses (*61, 67*), such modification likely alters the substrate binding pocket and improves TAL activity. We further demonstrated that simultaneously introducing two substitutions, Ser112 to Ile and Phe140 to His, converted both monocot and dicot PAL into PTAL enzymes (**Fig. 4, F and G**). The dual entry pathway via PTAL likely enabled efficient synthesis of phenylpropanoids in fast-growing grasses that still accumulate lignin at up to 30% of dry weight (*9, 12, 68*). Therefore, leveraging insights from grass evolution, the introduction of a second entry pathway of lignin biosynthesis, via the Ile112 and His140 residues, presents a promising gene editing strategy to enhance the production of diverse phenylpropanoid compounds in various plants.

Grasses have many derived functional traits that are hypothesized to have contributed to their evolutionary success of this prominent plant family with immense economic and ecological significance (*45, 69*). While many grass genomes have been sequenced (*26–29*), the ancestral characteristics of grasses remain poorly understood due to the lack of reference-quality genomes in the sister lineage to the grass family. The novel Poales genomes obtained in this study now offer valuable resources to dissect complex traits uniquely evolved in grasses and their relatives. Understanding the evolutionary basis of grass-specific traits will inform efforts to conserve grass-dominated ecosystems and accelerate grass breeding and engineering for sustainable production of food, feed, bioenergy, and biomaterials.

## Supporting information

Supplemental informations

## Acknowledgements

We greatly appreciate valuable discussions with Drs. Thomas J. Givnish for the selection of monocot species, Drs. Barbara Briggs and Elizabeth Kellogg for the *Ecdeiocolea monostachya* genome sequencing and analysis, and Drs. Jaime Barros-Rios and Richard Dixon for grass PTAL functions. We thank Dr. Jorge El-Azaz for his assistance with the biochemical analysis of PALs/PTALs, Sarah Friedrich from the UW Botany Media Studio for illustration, and Cara Steekstra from the UW-Madison Botany Greenhouse for the growth of *Joinvillea ascendens*.

## Authors contributions

H.A.M., J.H.L-M, M.R.M., M.J.M., and J.S. conceptualized the research. Y.T-K., B.M., S.H., M.B., M.O., D.L., J.G., M.W., L.B.B., and K.B. performed the experiments. Y.T-K., B.M., M.R.M., S.K.D., M.J.M., S.H., J.J., C.P., S.S., K.B., D.M.G., J.S., and H.A.M. analyzed data. The original draft was written by Y.T-K., H.A.M., J.H.L-M., S.H., M.J.M. and J.J. The manuscript was reviewed and edited by Y.T-K., B.M., H.A.M., J.H.L-M., S.H., M.J.M., M.R.M., J.J. and J.S.

## Data and materials availability

The raw PacBio and Illumina gDNA sequencing and RNAseq data for *Ecdeiocolea monostachya* is deposited in the NCBI database under BioProject code PRJNA894727. This *Ecdeiocolea monostachya* Whole Genome Shotgun project has been deposited at DDBJ/ENA/GenBank under BioProject codes PRJNA894727 (Illumina/Oxford Nanopore Technology hybrid assembly) and PRJNA1179411 (PacBio HiFi assembly; this version used in this paper). Code not previously published for the high-throughput production of cleaned alignments for phylogenetic reconstruction is available here: https://github.com/mrmckain/Polyploid_Genomics_Utilities. The rest of genomes available in Phytozome as listed in the Resource Table (table S16). All other data reported in this paper will be shared by the lead contacts upon request.

## Supplementary Materials

### Materials and Methods

Figs. S1 to S11

Table S1 to S16

References (*70-109*)

## References

1. W. Huang, L. Zhang, J. T. Columbus, Y. Hu, Y. Zhao, L. Tang, Z. Guo, W. Chen, M. McKain, M. Bartlett, C.-H. Huang, D.-Z. Li, S. Ge, H. Ma, A well-supported nuclear phylogeny of Poaceae and implications for the evolution of C4 photosynthesis. Mol. Plant 15, 755–777 (2022).

2. T. J. Givnish, M. Ames, J. R. McNeal, M. R. McKain, P. R. Steele, C. W. Depamphilis, S. W. Graham, J. C. Pires, D. W. Stevenson, W. B. Zomlefer, Others, Assembling the tree of the monocotyledons: plastome sequence phylogeny and evolution of Poales1. Ann. Mo. Bot. Gard. 97, 584–616 (2010).

3. M. R. McKain, H. Tang, J. R. McNeal, S. Ayyampalayam, J. I. Davis, C. W. dePamphilis, T. J. Givnish, J. C. Pires, D. W. Stevenson, J. H. Leebens-Mack, A Phylogenomic Assessment of Ancient Polyploidy and Genome Evolution across the Poales. Genome Biol. Evol. 8, 1150–1164 (2016).

4. ESS: 9.3 World Calories - Total. https://www.fao.org/economic/the-statistics-division-ess/chartroom-and-factoids/chartroom/93-world-calories-total/en/.

5. Z. Lin, X. Li, L. M. Shannon, C.-T. Yeh, M. L. Wang, G. Bai, Z. Peng, J. Li, H. N. Trick, T. E. Clemente, J. Doebley, P. S. Schnable, M. R. Tuinstra, T. T. Tesso, F. White, J. Yu, Parallel domestication of the Shattering1 genes in cereals. Nat. Genet. 44, 720–724 (2012).

6. Y. Wang, X. Bi, J. Zhong, Revisiting the origin and identity specification of the spikelet: A structural innovation in grasses (Poaceae). Plant Physiol. 190, 60–71 (2022).

7. A. S. Seetharam, Y. Yu, S. Bélanger, L. G. Clark, B. C. Meyers, E. A. Kellogg, M. B. Hufford, The Genome and the Evolution of the Grasses. Front. Plant Sci. 12, 710383 (2021).

8. S. Comparot-Moss, K. Denyer, The evolution of the starch biosynthetic pathway in cereals and other grasses. J. Exp. Bot. 60, 2481–2492 (2009).

9. R. Bhatia, J. A. Gallagher, L. D. Gomez, M. Bosch, Genetic engineering of grass cell wall polysaccharides for biorefining. Plant Biotechnol. J. 15, 1071–1092 (2017).

10. A. J. Ragauskas, G. T. Beckham, M. J. Biddy, R. Chandra, F. Chen, M. F. Davis, B. H. Davison, R. A. Dixon, P. Gilna, M. Keller, P. Langan, A. K. Naskar, J. N. Saddler, T. J. Tschaplinski, G. A. Tuskan, C. E. Wyman, Lignin Valorization: Improving Lignin Processing in the Biorefinery. Science 344, 1246843–1246843 (2014).

11. R. Rinaldi, R. Jastrzebski, M. T. Clough, J. Ralph, M. Kennema, P. C. A. Bruijnincx, B. M. Weckhuysen, Paving the Way for Lignin Valorisation: Recent Advances in Bioengineering, Biorefining and Catalysis. Angew. Chem. Int. Ed. 55, 8164–8215 (2016).

12. T. Umezawa, Lignin modification *in planta* for valorization. Phytochem. Rev., 1–23 (2018).

13. J. Rosler, F. Krekel, N. Amrhein, J. Schmid, Maize phenylalanine ammonia-lyase has tyrosine ammonia-lyase activity. Plant Physiol. 113, 175–179 (1997).

14. J. Barros, J. C. Serrani-Yarce, F. Chen, D. Baxter, B. J. Venables, R. A. Dixon, Role of bifunctional ammonia-lyase in grass cell wall biosynthesis. Nat. Plants 2, 16050 (2016).

15. Y. Van de Peer, T.-L. Ashman, P. S. Soltis, D. E. Soltis, Polyploidy: an evolutionary and ecological force in stressful times. Plant Cell 33, 11–26 (2021).

16. F. Almeida-Silva, Y. Van de Peer, Whole-genome Duplications and the Long-term Evolution of Gene Regulatory Networks in Angiosperms. Mol. Biol. Evol. 40 (2023).

17. N. Walden, D. A. German, E. M. Wolf, M. Kiefer, P. Rigault, X.-C. Huang, C. Kiefer, R. Schmickl, A. Franzke, B. Neuffer, K. Mummenhoff, M. A. Koch, Nested whole-genome duplications coincide with diversification and high morphological disparity in Brassicaceae. Nat. Commun. 11, 3795 (2020).

18. K. C. Moharana, T. M. Venancio, Polyploidization events shaped the transcription factor repertoires in legumes (Fabaceae). Plant J. 103, 726–741 (2020).

19. T. Zhang, W. Huang, L. Zhang, D.-Z. Li, J. Qi, H. Ma, Phylogenomic profiles of whole-genome duplications in Poaceae and landscape of differential duplicate retention and losses among major Poaceae lineages. Nat. Commun. 15, 3305 (2024).

20. J. W. Clark, P. C. J. Donoghue, Whole-Genome Duplication and Plant Macroevolution. Trends Plant Sci. 23, 933–945 (2018).

21. A. H. Paterson, J. E. Bowers, B. A. Chapman, Ancient polyploidization predating divergence of the cereals, and its consequences for comparative genomics. Proc. Nat. Acad. Sci. U. S. A. 101, 9903– 9908 (2004).

22. J. Yu, J. Wang, W. Lin, S. Li, H. Li, J. Zhou, P. Ni, W. Dong, S. Hu, C. Zeng, J. Zhang, Y. Zhang, R. Li, Z. Xu, S. Li, X. Li, H. Zheng, L. Cong, L. Lin, J. Yin, J. Geng, G. Li, J. Shi, J. Liu, H. Lv, J. Li, J. Wang, Y. Deng, L. Ran, X. Shi, X. Wang, Q. Wu, C. Li, X. Ren, J. Wang, X. Wang, D. Li, D. Liu, X. Zhang, Z. Ji, W. Zhao, Y. Sun, Z. Zhang, J. Bao, Y. Han, L. Dong, J. Ji, P. Chen, S. Wu, J. Liu, Y. Xiao, D. Bu, J. Tan, L. Yang, C. Ye, J. Zhang, J. Xu, Y. Zhou, Y. Yu, B. Zhang, S. Zhuang, H. Wei, B. Liu, M. Lei, H. Yu, Y. Li, H. Xu, S. Wei, X. He, L. Fang, Z. Zhang, Y. Zhang, X. Huang, Z. Su, W. Tong, J. Li, Z. Tong, S. Li, J. Ye, L. Wang, L. Fang, T. Lei, C. Chen, H. Chen, Z. Xu, H. Li, H. Huang, F. Zhang, H. Xu, N. Li, C. Zhao, S. Li, L. Dong, Y. Huang, L. Li, Y. Xi, Q. Qi, W. Li, B. Zhang, W. Hu, Y. Zhang, X. Tian, Y. Jiao, X. Liang, J. Jin, L. Gao, W. Zheng, B. Hao, S. Liu, W. Wang, L. Yuan, M. Cao, J. McDermott, R. Samudrala, J. Wang, G. K.-S. Wong, H. Yang, The Genomes of Oryza sativa: A History of Duplications. PLOS Biology 3, e38 (2005).

23. H. Tang, J. E. Bowers, X. Wang, A. H. Paterson, Angiosperm genome comparisons reveal early polyploidy in the monocot lineage. Proc. Natl. Acad. Sci. U. S. A. 107, 472–477 (2010).

24. S. Ohno, Evolution by Gene Duplication. 1970, London: G. Allen Google Scholar.

25. K. Vanneste, G. Baele, S. Maere, Y. Van de Peer, Analysis of 41 plant genomes supports a wave of successful genome duplications in association with the Cretaceous-Paleogene boundary. Genome Res. 24, 1334–1347 (2014).

26. S. A. Goff, D. Ricke, T.-H. Lan, G. Presting, R. Wang, M. Dunn, J. Glazebrook, A. Sessions, P. Oeller, H. Varma, D. Hadley, D. Hutchison, C. Martin, F. Katagiri, B. M. Lange, T. Moughamer, Y. Xia, P. Budworth, J. Zhong, T. Miguel, U. Paszkowski, S. Zhang, M. Colbert, W.-L. Sun, L. Chen, B. Cooper, S. Park, T. C. Wood, L. Mao, P. Quail, R. Wing, R. Dean, Y. Yu, A. Zharkikh, R. Shen, S. Sahasrabudhe, A. Thomas, R. Cannings, A. Gutin, D. Pruss, J. Reid, S. Tavtigian, J. Mitchell, G. Eldredge, T. Scholl, R. M. Miller, S. Bhatnagar, N. Adey, T. Rubano, N. Tusneem, R. Robinson, J. Feldhaus, T. Macalma, A. Oliphant, S. Briggs, A draft sequence of the rice genome (Oryza sativa L. ssp. japonica). Science 296, 92–100 (2002).

27. J. Yu, S. Hu, J. Wang, G. K.-S. Wong, S. Li, B. Liu, Y. Deng, L. Dai, Y. Zhou, X. Zhang, M. Cao, J. Liu, J. Sun, J. Tang, Y. Chen, X. Huang, W. Lin, C. Ye, W. Tong, L. Cong, J. Geng, Y. Han, L. Li, W. Li, G. Hu, X. Huang, W. Li, J. Li, Z. Liu, L. Li, J. Liu, Q. Qi, J. Liu, L. Li, T. Li, X. Wang, H. Lu, T. Wu, M. Zhu, P. Ni, H. Han, W. Dong, X. Ren, X. Feng, P. Cui, X. Li, H. Wang, X. Xu, W. Zhai, Z. Xu, J. Zhang, S. He, J. Zhang, J. Xu, K. Zhang, X. Zheng, J. Dong, W. Zeng, L. Tao, J. Ye, J. Tan, X. Ren, X. Chen, J. He, D. Liu, W. Tian, C. Tian, H. Xia, Q. Bao, G. Li, H. Gao, T. Cao, J. Wang, W. Zhao, P. Li, W. Chen, X. Wang, Y. Zhang, J. Hu, J. Wang, S. Liu, J. Yang, G. Zhang, Y. Xiong, Z. Li, L. Mao, C. Zhou, Z. Zhu, R. Chen, B. Hao, W. Zheng, S. Chen, W. Guo, G. Li, S. Liu, M. Tao, J. Wang, L. Zhu, L. Yuan, H. Yang, A draft sequence of the rice genome (Oryza sativa L. ssp. indica). Science 296, 79–92 (2002).

28. P. S. Schnable, D. Ware, R. S. Fulton, J. C. Stein, F. Wei, S. Pasternak, C. Liang, J. Zhang, L. Fulton, T. A. Graves, P. Minx, A. D. Reily, L. Courtney, S. S. Kruchowski, C. Tomlinson, C. Strong, K. Delehaunty, C. Fronick, B. Courtney, S. M. Rock, E. Belter, F. Du, K. Kim, R. M. Abbott, M. Cotton, A. Levy, P. Marchetto, K. Ochoa, S. M. Jackson, B. Gillam, W. Chen, L. Yan, J. Higginbotham, M. Cardenas, J. Waligorski, E. Applebaum, L. Phelps, J. Falcone, K. Kanchi, T. Thane, A. Scimone, N. Thane, J. Henke, T. Wang, J. Ruppert, N. Shah, K. Rotter, J. Hodges, E. Ingenthron, M. Cordes, S. Kohlberg, J. Sgro, B. Delgado, K. Mead, A. Chinwalla, S. Leonard, K. Crouse, K. Collura, D. Kudrna, J. Currie, R. He, A. Angelova, S. Rajasekar, T. Mueller, R. Lomeli, G. Scara, A. Ko, K. Delaney, M. Wissotski, G. Lopez, D. Campos, M. Braidotti, E. Ashley, W. Golser, H. Kim, S. Lee, J. Lin, Z. Dujmic, W. Kim, J. Talag, A. Zuccolo, C. Fan, A. Sebastian, M. Kramer, L. Spiegel, L. Nascimento, T. Zutavern, B. Miller, C. Ambroise, S. Muller, W. Spooner, A. Narechania, L. Ren, S. Wei, S. Kumari, B. Faga, M. J. Levy, L. McMahan, P. Van Buren, M. W. Vaughn, K. Ying, C.-T. Yeh, S. J. Emrich, Y. Jia, A. Kalyanaraman, A.-P. Hsia, W. B. Barbazuk, R. S. Baucom, T. P. Brutnell, N. C. Carpita, C. Chaparro, J.-M. Chia, J.-M. Deragon, J. C. Estill, Y. Fu, J. A. Jeddeloh, Y. Han, H. Lee, P. Li, D. R. Lisch, S. Liu, Z. Liu, D. H. Nagel, M. C. McCann, P. SanMiguel, A. M. Myers, D. Nettleton, J. Nguyen, B. W. Penning, L. Ponnala, K. L. Schneider, D. C. Schwartz, A. Sharma, C. Soderlund, N. M. Springer, Q. Sun, H. Wang, M. Waterman, R. Westerman, T. K. Wolfgruber, L. Yang, Y. Yu, L. Zhang, S. Zhou, Q. Zhu, J. L. Bennetzen, R. K. Dawe, J. Jiang, N. Jiang, G. G. Presting, S. R. Wessler, S. Aluru, R. A. Martienssen, S. W. Clifton, W. R. McCombie, R. A. Wing, R. K. Wilson, The B73 maize genome: complexity, diversity, and dynamics. Science 326, 1112–1115 (2009).

29. International Wheat Genome Sequencing Consortium (IWGSC), A chromosome-based draft sequence of the hexaploid bread wheat (Triticum aestivum) genome. Science 345, 1251788 (2014).

30. P. R. Timilsena, E. K. Wafula, C. F. Barrett, S. Ayyampalayam, J. R. McNeal, J. D. Rentsch, M. R. McKain, K. Heyduk, A. Harkess, M. Villegente, J. G. Conran, N. Illing, B. Fogliani, C. Ané, J. C. Pires, J. I. Davis, W. B. Zomlefer, D. W. Stevenson, S. W. Graham, T. J. Givnish, J. Leebens-Mack, C. W. dePamphilis, Phylogenomic resolution of order- and family-level monocot relationships using 602 single-copy nuclear genes and 1375 BUSCO genes. Front. Plant Sci. 13, 876779 (2022).

31. T. L. Elliott, D. Spalink, I. Larridon, A. R. Zuntini, M. Escudero, J. Hackel, R. L. Barrett, S. Martín-Bravo, J. I. Márquez-Corro, C. Granados Mendoza, A. C. Mashau, K. J. Romero-Soler, D. A. Zhigila, B. Gehrke, C. O. Andrino, D. M. Crayn, M. S. Vorontsova, F. Forest, W. J. Baker, K. L. Wilson, D. A. Simpson, A. M. Muasya, Global analysis of Poales diversification - parallel evolution in space and time into open and closed habitats. New Phytol. 242, 727–743 (2024).

32. L. Hanson, A. Boyd, M. A. T. Johnson, M. D. Bennett, First Nuclear DNA C-values for 18 Eudicot Families. Ann. Bot. 96, 1315–1320 (2005).

33. P.-F. Ma, Y.-L. Liu, G.-H. Jin, J.-X. Liu, H. Wu, J. He, Z.-H. Guo, D.-Z. Li, The Pharus latifolius genome bridges the gap of early grass evolution. Plant Cell 33, 846–864 (2021).

34. R. Vaser, I. Sović, N. Nagarajan, M. Šikić, Fast and accurate de novo genome assembly from long uncorrected reads. Genome Res. 27, 737–746 (2017).

35. J. T. Lovell, A. Sreedasyam, M. E. Schranz, M. Wilson, J. W. Carlson, A. Harkess, D. Emms, D. M. Goodstein, J. Schmutz, GENESPACE tracks regions of interest and gene copy number variation across multiple genomes. Elife 11 (2022).

36. H. Tang, J. E. Bowers, X. Wang, R. Ming, M. Alam, A. H. Paterson, Synteny and Collinearity in Plant Genomes. Science 320, 486–488 (2008).

37. A. H. Paterson, J. E. Bowers, R. Bruggmann, I. Dubchak, J. Grimwood, H. Gundlach, G. Haberer, U. Hellsten, T. Mitros, A. Poliakov, J. Schmutz, M. Spannagl, H. Tang, X. Wang, T. Wicker, A. K. Bharti, J. Chapman, F. A. Feltus, U. Gowik, I. V. Grigoriev, E. Lyons, C. A. Maher, M. Martis, A. Narechania, R. P. Otillar, B. W. Penning, A. A. Salamov, Y. Wang, L. Zhang, N. C. Carpita, M. Freeling, A. R. Gingle, C. T. Hash, B. Keller, P. Klein, S. Kresovich, M. C. McCann, R. Ming, D. G. Peterson, Mehboob-ur-Rahman, D. Ware, P. Westhoff, K. F. X. Mayer, J. Messing, D. S. Rokhsar, The Sorghum bicolor genome and the diversification of grasses. Nature 457, 551–556 (2009).

38. Y. Jiao, J. Li, H. Tang, A. H. Paterson, Integrated Syntenic and Phylogenomic Analyses Reveal an Ancient Genome Duplication in Monocots. Plant Cell 26, 2792–2802 (2014).

39. M. R. McKain, M. C. Estep, R. Pasquet, D. J. Layton, D. M. V. Díaz, J. Zhong, J. G. Hodge, S. T. Malcomber, G. Chipabika, B. Pallangyo, E. A. Kellogg, Ancestry of the two subgenomes of maize. bioRxiv [Preprint] (2018). 10.1101/352351.

40. S. K. Deb, P. P. Edger, J. C. Pires, M. R. McKain, Patterns, mechanisms, and consequences of homoeologous exchange in allopolyploid angiosperms: a genomic and epigenomic perspective. New Phytol. 238, 2284–2304 (2023).

41. O. Jaillon, J.-M. Aury, B. Noel, A. Policriti, C. Clepet, A. Casagrande, N. Choisne, S. Aubourg, N. Vitulo, C. Jubin, A. Vezzi, F. Legeai, P. Hugueney, C. Dasilva, D. Horner, E. Mica, D. Jublot, J. Poulain, C. Bruyère, A. Billault, B. Segurens, M. Gouyvenoux, E. Ugarte, F. Cattonaro, V. Anthouard, V. Vico, C. Del Fabbro, M. Alaux, G. Di Gaspero, V. Dumas, N. Felice, S. Paillard, I. Juman, M. Moroldo, S. Scalabrin, A. Canaguier, I. Le Clainche, G. Malacrida, E. Durand, G. Pesole, V. Laucou, P. Chatelet, D. Merdinoglu, M. Delledonne, M. Pezzotti, A. Lecharny, C. Scarpelli, F. Artiguenave, M. E. Pè, G. Valle, M. Morgante, M. Caboche, A.-F. Adam-Blondon, J. Weissenbach, F. Quétier, P. Wincker, The grapevine genome sequence suggests ancestral hexaploidization in major angiosperm phyla. Nature 449, 463–467 (2007).

42. J. H. Leebens-Mack, M. S. Barker, E. J. Carpenter, M. K. Deyholos, M. A. Gitzendanner, S. W. Graham, I. Grosse, Z. Li, M. Melkonian, S. Mirarab, M. Porsch, M. Quint, S. A. Rensing, D. E. Soltis, P. S. Soltis, D. W. Stevenson, K. K. Ullrich, N. J. Wickett, L. DeGironimo, P. P. Edger, I. E. Jordon-Thaden, S. Joya, T. Liu, B. Melkonian, N. W. Miles, L. Pokorny, C. Quigley, P. Thomas, J. C. Villarreal, M. M. Augustin, M. D. Barrett, R. S. Baucom, D. J. Beerling, R. M. Benstein, E. Biffin, S. F. Brockington, D. O. Burge, J. N. Burris, K. P. Burris, V. Burtet-Sarramegna, A. L. Caicedo, S. B. Cannon, Z. Çebi, Y. Chang, C. Chater, J. M. Cheeseman, T. Chen, N. D. Clarke, H. Clayton, S. Covshoff, B. J. Crandall-Stotler, H. Cross, C. W. dePamphilis, J. P. Der, R. Determann, R. C. Dickson, V. S. Di Stilio, S. Ellis, E. Fast, N. Feja, K. J. Field, D. A. Filatov, P. M. Finnegan, S. K. Floyd, B. Fogliani, N. García, G. Gâteblé, G. T. Godden, F. (Qi Y. Goh, S. Greiner, A. Harkess, J. M. Heaney, K. E. Helliwell, K. Heyduk, J. M. Hibberd, R. G. J. Hodel, P. M. Hollingsworth, M. T. J. Johnson, R. Jost, B. Joyce, M. V. Kapralov, E. Kazamia, E. A. Kellogg, M. A. Koch, M. Von Konrat, K. Könyves, T. M. Kutchan, V. Lam, A. Larsson, A. R. Leitch, R. Lentz, F.-W. Li, A. J. Lowe, M. Ludwig, P. S. Manos, E. Mavrodiev, M. K. McCormick, M. McKain, T. McLellan, J. R. McNeal, R. E. Miller, M. N. Nelson, Y. Peng, P. Ralph, D. Real, C. W. Riggins, M. Ruhsam, R. F. Sage, A. K. Sakai, M. Scascitella, E. E. Schilling, E.-M. Schlösser, H. Sederoff, S. Servick, E. B. Sessa, A. J. Shaw, S. W. Shaw, E. M. Sigel, C. Skema, A. G. Smith, A. Smithson, C. N. Stewart, J. R. Stinchcombe, P. Szövényi, J. A. Tate, H. Tiebel, D. Trapnell, M. Villegente, C.-N. Wang, S. G. Weller, M. Wenzel, S. Weststrand, J. H. Westwood, D. F. Whigham, S. Wu, A. S. Wulff, Y. Yang, D. Zhu, C. Zhuang, J. Zuidof, M. W. Chase, J. C. Pires, C. J. Rothfels, J. Yu, C. Chen, L. Chen, S. Cheng, J. Li, R. Li, X. Li, H. Lu, Y. Ou, X. Sun, X. Tan, J. Tang, Z. Tian, F. Wang, J. Wang, X. Wei, X. Xu, Z. Yan, F. Yang, X. Zhong, F. Zhou, Y. Zhu, Y. Zhang, S. Ayyampalayam, T. J. Barkman, N. Nguyen, N. Matasci, D. R. Nelson, E. Sayyari, E. K. Wafula, R. L. Walls, T. Warnow, H. An, N. Arrigo, A. E. Baniaga, S. Galuska, S. A. Jorgensen, T. I. Kidder, H. Kong, P. Lu-Irving, H. E. Marx, X. Qi, C. R. Reardon, B. L. Sutherland, G. P. Tiley, S. R. Welles, R. Yu, S. Zhan, L. Gramzow, G. Theißen, G. K.-S. Wong, One Thousand Plant Transcriptomes Initiative, One thousand plant transcriptomes and the phylogenomics of green plants. Nature 574, 679–685 (2019).

43. S. C. Zeeman, J. Kossmann, A. M. Smith, Starch: Its Metabolism, Evolution, and Biotechnological Modification in Plants. Ann. Rev. Plant Biol. 61, 209–234 (2010).

44. G. J. MacNeill, S. Mehrpouyan, M. A. A. Minow, J. A. Patterson, I. J. Tetlow, M. J. Emes, Starch as a source, starch as a sink: the bifunctional role of starch in carbon allocation. J. Exp. Bot. 68, 4433– 4453 (2017).

45. H. P. Linder, C. E. R. Lehmann, S. Archibald, C. P. Osborne, D. M. Richardson, Global grass (Poaceae) success underpinned by traits facilitating colonization, persistence and habitat transformation. Biol. Rev. Camb. Philos. Soc. 93, 1125–1144 (2018).

46. B. Pfister, S. C. Zeeman, Formation of starch in plant cells. Cell. Mol. Life Sci. 73, 2781–2807 (2016).

47. D. M. Beckles, A. M. Smith, T. ap Rees, A Cytosolic ADP-Glucose Pyrophosphorylase Is a Feature of Graminaceous Endosperms, But Not of Other Starch-Storing Organs1. Plant Physiol. 125, 818–827 (2001).

48. B. Huang, T. A. Hennen-Bierwagen, A. M. Myers, Functions of Multiple Genes Encoding ADP-Glucose Pyrophosphorylase Subunits in Maize Endosperm, Embryo, and Leaf. Plant Physiol. 164, 596–611 (2014).

49. N. J. Patron, B. Greber, B. F. Fahy, D. A. Laurie, M. L. Parker, K. Denyer, The lys5 Mutations of Barley Reveal the Nature and Importance of Plastidial ADP-Glc Transporters for Starch Synthesis in Cereal Endosperm. Plant Physiol. 135, 2088–2097 (2004).

50. T. Konishi, Y. Sasaki, Compartmentalization of two forms of acetyl-CoA carboxylase in plants and the origin of their tolerance toward herbicides. Proc. Natl. Acad. Sci. U. S. A. 91, 3598–3601 (1994).

51. Y. Sasaki, Y. Nagano, Plant Acetyl-CoA Carboxylase: Structure, Biosynthesis, Regulation, and Gene Manipulation for Plant Breeding. Biosci. Biotechnol. Biochem. 68, 1175–1184 (2004).

52. C. Délye, X.-Q. Zhang, S. Michel, A. Matéjicek, S. B. Powles, Molecular Bases for Sensitivity to Acetyl-Coenzyme A Carboxylase Inhibitors in Black-Grass. Plant Physiol. 137, 794–806 (2005).

53. A. C. Neish, Formation of *m*- and *p*-coumaric acids by enzymatic deamination of the corresponding isomers of tyrosine. Phytochemistry 1, 1–24 (1961).

54. M. Kordi, M. S. Bansal, “TreeSolve: Rapid Error-Correction of Microbial Gene Trees” in Algorithms for Computational Biology, C. Martín-Vide, M. A. Vega-Rodríguez, T. Wheeler, Eds. (Springer International Publishing, Cham, 2020), pp. 125–139.

55. K. T. Watts, P. C. Lee, C. Schmidt-Dannert, Biosynthesis of plant-specific stilbene polyketides in metabolically engineered Escherichia coli. BMC biotechnol. 6, 22 (2006).

56. S.-Y. Jun, S. A. Sattler, G. S. Cortez, W. Vermerris, S. E. Sattler, C. Kang, Biochemical and Structural Analysis of Substrate Specificity of a Phenylalanine Ammonia-Lyase. Plant Physiology 176, 1452– 1468 (2018).

57. G. V. Louie, M. E. Bowman, M. C. Moffitt, T. J. Baiga, B. S. Moore, J. P. Noel, Structural Determinants and Modulation of Substrate Specificity in Phenylalanine-Tyrosine Ammonia-Lyases. Chem. Biol. 13, 1327–1338 (2006).

58. Z. Yang, PAML 4: phylogenetic analysis by maximum likelihood. Mol. Biol. Evol. 24, 1586–1591 (2007).

59. H. A. Maeda, Harnessing evolutionary diversification of primary metabolism for plant synthetic biology. J. Biol. Chem. 294, 16549–16566 (2019).

60. Z. Bata, Z. Molnár, E. Madaras, B. Molnár, E. Sánta-Bell, A. Varga, I. Leveles, R. Qian, F. Hammerschmidt, C. Paizs, B. G. Vértessy, L. Poppe, Substrate Tunnel Engineering Aided by X-ray Crystallography and Functional Dynamics Swaps the Function of MIO-Enzymes. ACS Catal. 11, 4538–4549 (2021).

61. D. Röther, L. Poppe, G. Morlock, S. Viergutz, J. Rétey, An active site homology model of phenylalanine ammonia-lyase from P. crispum. Eur. J. Biochem. 269, 3065–3075 (2002).

62. F. C. Cochrane, L. B. Davin, N. G. Lewis, The Arabidopsis phenylalanine ammonia lyase gene family: kinetic characterization of the four PAL isoforms. Phytochemistry 65, 1557–1564 (2004).

63. Chimpanzee Sequencing and Analysis Consortium, Initial sequence of the chimpanzee genome and comparison with the human genome. Nature 437, 69–87 (2005).

64. K. Prüfer, K. Munch, I. Hellmann, K. Akagi, J. R. Miller, B. Walenz, S. Koren, G. Sutton, C. Kodira, R. Winer, J. R. Knight, J. C. Mullikin, S. J. Meader, C. P. Ponting, G. Lunter, S. Higashino, A. Hobolth, J. Dutheil, E. Karakoç, C. Alkan, S. Sajjadian, C. R. Catacchio, M. Ventura, T. Marques-Bonet, E. E. Eichler, C. André, R. Atencia, L. Mugisha, J. Junhold, N. Patterson, M. Siebauer, J. M. Good, A. Fischer, S. E. Ptak, M. Lachmann, D. E. Symer, T. Mailund, M. H. Schierup, A. M. Andrés, J. Kelso, S. Pääbo, The bonobo genome compared with the chimpanzee and human genomes. Nature 486, 527– 531 (2012).

65. M. E. Schranz, S. Mohammadin, P. P. Edger, Ancient whole genome duplications, novelty and diversification: the WGD Radiation Lag-Time Model. Curr. Opin. Plant Biol. 15, 147–153 (2012).

66. J. C. Preston, A. Christensen, S. T. Malcomber, E. A. Kellogg, MADS-box gene expression and implications for developmental origins of the grass spikelet. Am. J. Bot. 96, 1419–1429 (2009).

67. J. Barros, R. A. Dixon, Plant Phenylalanine/Tyrosine Ammonia-lyases. Trend Plant Sci. 25, 66–79 (2020).

68. Y. Y. Tye, K. T. Lee, W. N. Wan Abdullah, C. P. Leh, The world availability of non-wood lignocellulosic biomass for the production of cellulosic ethanol and potential pretreatments for the enhancement of enzymatic saccharification. Renew. Sust. Energ. Rev. 60, 155–172 (2016).

69. T. R. Hodkinson, Evolution and taxonomy of the grasses (Poaceae): A model family for the study of species-rich groups, Ann. Plant Rev. (2018)pp. 1–39.

