## Supplemental informations for "Genomes of Poaceae sisters reveal key metabolic innovations preceding the evolution of grasses"

**The file includes:**

Materials and Methods
Figs. S1 to S11
Tables S1 to S16
References

### Materials and Methods:

***Genome sequencing of Joinvillea ascendens***

To obtain genome sequence of *J. ascendens*, in May 2017, we first harvested drupes, fruits containing seeds (the seed batch of NTBG #800379), from the *J. ascendens* subsp. glabra plant grown at the National Tropical Garden (Kalaheo, HI, Accession No. 800379, herbarium voucher Lorence 7657, PTBG) from seed originally collected in New Caledonia. These drupes were sent to University of Wisconsin-Madison, physically removed for fruits using a laser blade, and the seeds were soaked with 5% liquid smoke (Wright's Liquid Smoke, Hickory) for 24 hours (*70*), rinsed in distilled water, and then planted on three parts perlite to one part potting mix. After two months in July 2017, one out of >100 planted seeds germinated. After 6 months of the germination in January 2018, newly developed young leaves were sprayed with clean distilled water, and ~100 mg fresh tissues were harvested and ground in liquid nitrogen and used for genomic DNA isolation using the DNeasy Plant Mini Kit (Qiagen).

The isolated DNA was sent to and sequenced at JGI using PACBIO platform at 125x coverage (average read length of 10,007). Main assembly was performed and polished using MECAT (*71*). Misjoins in the assembly were identified using Hi-C data. A total of 27 misjoins were identified in the polished assembly. Scaffolds were then oriented, ordered, and joined together using Hi-C scaffolding. Significant telomeric sequence was properly oriented in the assembly. A total of 186 joins were applied to the broken assembly to form the final assembly consisting of 18 chromosomes (2N), with a total of 96.5% of the assembled sequence contained in the chromosomes. Hi-C reads were then aligned to the joined release. The alignments were converted to contact map to quality control check on the order/orientation of contigs in the chromosomes. Seven discrepancies were corrected. Finally, care was taken to ensure that telomere sequence was properly oriented in the chromosomes, and the resulting sequence was screened for retained vector and/or contaminants. Adjacent alternative haplotypes were identified on the joined contig set. Alternative haplotype regions were collapsed using the longest common substring between the two haplotypes. A total of 67 adjacent alternative haplotypes were collapsed. Chromosomes were numbered from largest to smallest, and the p-arm of each chromosome was oriented to the 5' end. Additionally, homozygous SNPs and INDELs were corrected in the release sequence using 40x of Illumina reads (2x150, 400bp insert).

To aid in annotation by MAKER (*72*), ~2B pairs of 2X150 stranded paired-end Illumina RNA sequencing was performed on various tissues: the leaf tip, leaf base, upper internodes, base internodes, and root tissues harvested into liquid nitrogen at University of Wisconsin-Madison, as well as the inflorescence tissues harvested and immediately cooled by dry ice at the National Tropical Garden. transcript assemblies were generated from 2 × 150 paired-end Illumina RNA-seq reads using PERTRAN (*73*), which conducts genome-guided transcriptome short read assembly via GSNAP (*74*) and builds splice alignment graphs after alignment validation, realignment and correction. The highest-scoring predictions for each locus were selected using multiple positive factors including EST and protein support, and one negative factor: overlap with repeats. Improvement includes adding UTRs, splicing correction, and adding alternative transcripts. The transcripts were selected if the Cscore was larger than or equal to 0.5 and protein coverage larger than or equal to 0.5, or if it had EST coverage, but the CDS overlapping with repeats was less than 20%. For gene models whose CDS overlaps with repeats for more that 20%, its Cscore had to be at least 0.9 and homology coverage at least 70% to be selected. The selected gene models were subject to Pfam analysis and gene models whose protein was more than 30% in Pfam Transposable elements (TE) domains were removed. Repetitive DNA elements were identified de novo with RepeatModeler.

***Genome sequencing and analysis of Ecdeiocolea monostachya***

The accession of *E. monostachya* E001 from Western Australia (PERTH 09450289) was sampled and isolated at flowering stage on 2018 Sep 12 (**fig. S9**). A genome size estimate of *E. monostachya* accession E001 was 2.0 pg (2C) with 38 or 42 chromosomes (2N). Analysis of three additional accessions (E002, E010, and E014) had similar estimates of genome size and chromosome number. Hanson and colleagues (2005) previous estimated chromosome number at 38 and genome size of 1.98 pg (2C). Therefore, *E. monostachya* is estimated to have a haploid genome size of 1.94 Gb.

The genome assembly was performed using HiFiAsm using PacBio HiFi long-reads. Gene models were predicted using *ab initio*, homology, and evidence-based gene models using RNAseq from sheath, root and flower tissue. For genome assembly, genomic DNA was collected from sheath tissue of *E. monostachya* accession E001 with a CTAB-based extraction protocol (*75*) and sequenced using paired-end Illumina short-read and PacBio HiFi long-read sequencing. The estimated haploid genome size using GenomeScope (*76*) ranged from 734.0 Mb (k=17) to 839.2 (k=24) and heterozygosity estimate of 2.33% (k=31) to 2.65% (k=17). findGSE (*77*) and KmerGenie (*78*) predicted diploid genome sizes of 1.47 Gb (k=24) and 1.31 Gb (k=107).

A total of 16.84 Gb of PacBio HiFi data was generated with an approximate depth of 21x and average read length of 14.4 kb. Genome assembly was performed using HiFiAsm (v0.16) (*79*, *80*) using parameters '-I = 2' and '-n=4'. Contigs derived from heterozygous regions were removed using purge_dups (v.1.2.5)(*81*) using default parameters. The final draft haploid genome assembly was 897 Mbp over 1,114 scaffolds (**Table 1**). The majority of the genome (>50%) spans contigs greater than 1.7 Mb. Evaluation of *k*-mer composition of the resulting assembly relative to raw Illumina reads found that the genome was partially phased (**fig. S10**).

An integrated gene annotation approach using de novo prediction, homology search, and evidence-based (RNAseq) assembly was used to annotate protein-coding genes in the genome. Prediction of *de novo* gene models was performed with Augustus (v2.4) (*82*) and SNAP (v2006-07-28) (*83*). The homolog-based approach involved GeMoMa (v1.7)(*84*) using reference gene model from Poales species (*Brachypodium distachyon*, *Setaria italica*, *Oryza sativa*, and *Joinvillea ascendens*). Evidence-based gene prediction used paired-end Illumina RNAseq derived from sheath, root, and flower (immature embryos) samples of *E. monostachya* accession E001. Total RNA was extracted using a Trizol-based extraction protocol. For flower samples, developing seeds were extracted from whole flowers in liquid nitrogen. Spliced alignment using hisat2 (*85*) found that the majority of RNAseq reads (89.5-91.0%) aligned to the genome with non-aligning RNAseq reads (9.1-9.6%) associated with bacterial, human, Achaea, or viral contamination based on kraken2 (*86*) classification. Reads were mapped using hisat (v2.0.4)(*85*) and assembled by Stringtie (v1.2.3) (*87*). Genes were predicted from assembled transcripts using GeneMarkS-T (v5.1)(*88*). Gene prediction from Trinity (v2.11)(*89*) was performed with PASA (v2.0.2)(*90*). EVM (v1.1.1)(*91*) was used to merge gene models from these multiple approaches and was updated by PASA. Analysis of the benchmark universal single copy orthologs (BUSCO)(*92*) using the predicted transcripts found 94.6% complete, 2.9% fragmented, and 2.5% missing genes. The majority of BUSCO genes were not duplicated (7.0%) reflecting the purging of redundant contigs. The final set of protein encoding genes included 27,801 gene models.

Previous work has shown that the GC content of exons in Poaceae genomes has a distinct bimodal distribution (*3*, *93*). Evaluation of the GC content of *E. monostachya* found monomodal distribution that is skewed towards high GC content (**fig. S11**). This suggests that a subset of genes have a higher GC content than expected through a neutral process.

For short read and long-read sequencing, Illumina and Oxford Nanopore Technology (ONT) library construction and sequencing using *E. monostachya* accession E001 genomic DNA was performed by Novogene (Beijing, China). Illumina paired end libraries were generated using inserts of 250 bp and 350 bp and sequenced using Illumina HiSeq. In total, 123.2 Gb of short-read Illumina sequencing data was generated. In total, 12.8 million ONT reads were generated that ranged in length from 500 bp to 849 kb with a median length of 4.6 kb and total size of 69.7 Gb.

Hybrid assembly of the *E. monostachya* accession E001 genome was carried out using MaSuRCA (v3.3.0) using default parameters on an Amazon AWS x1.32xlarge instance (128 CPUs, 1,952 GB RAM) running SUSE Linux with raw Illumina and ONT reads. BUSCO genes were identified using BUSCO (v3.1.0) (*92*) with default parameters and Embryophyta (ODB9) database.

For k-mer analysis, Illumina reads were trimmed using Trimmomatic (v0.36)(*94*) using the parameters 2:30:10 for clipping reads based on TruSeq3 paired end adapters, leading and trailing of 5 bp, sliding window of 4:10, and a minimum length of 36 bp. Initial *k*-mer analysis of Illumina short-read data to determine coverage was performed using jellyfish (v1.1.12) with k of 17, 24, 27, and 31 bp. Genome size was estimated using KmerGenie (v1.7048)(*95*), findGSE (*77*), and GenomeScope (*76*) (http://genomescope.org/). The *kat comp* command in the *k*-mer analysis tookit (v2.4.1) using default parameters (k=27) was used to generate the stacked histogram to determine the proportion and frequency of *k*-mers in short-read sequencing data relative to the hybrid genome assembly.

To determine potential contamination in *E. monostachya* RNAseq samples, kraken2 (v2.0.8-beta) (*86*) was used to classify the kingdom origin of individual paired reads using default parameters and libraries: archaea, bacteria, plasmid, viral, human, fungi, plant, protozoa, UniVec_core, and nt. The kraken2 database was generated on Nov 27th, 2019.

For gene annotation, RNAseq reads were trimmed using Trimmomatic (v0.36)(*94*) using the same parameters for genomic DNA reads. Spliced alignments to the hybrid genome were performed using hisat2 (v2.2.1)(*85*) with the following parameters: maximum intron length of 20 kb, k of 1, no soft clipping, and alignments tailored for Cufflinks. SAM files were converted into sorted BAM files using samtools (v1.11). Cufflinks/Cuffmerge (v2.2.1) pipeline using default parameters was used to identify evidence-based gene models. TransDecoder (v5.5.0) was used with default parameters to identify open reading frames (LongORFs) and predicted proteins (Predict) using Pfam. Completeness of conserved orthologs was performed using BUSCO (v3)(*92*) with the embryophyta_odb9 lineage and default parameters.

***Genome sequencing of Pharus latifolius***

We sequenced *Pharus latifolius* (accession 1993-0885-2 from the Missouri Botanical Garden; herbarium voucher McKain 320) using a whole genome shotgun sequencing strategy and standard sequencing protocols. This specimen was originally collected from near Napo, Ecuador on February 7, 1992 by MacDougal & Lalumondier (collection number 4792). Since that time, it has been maintained and propagated in the climatron at the Missouri Botanical Garden. Sequencing reads were collected using Illumina and PACBIO platforms. Illumina and PACBIO reads were sequenced at the HudsonAlpha Institute in Huntsville, Alabama using the Illumina NovoSeq 6000 platform and the SEQUEL II platform, respectively. One 400bp insert 1x250 Illumina fragment library (170.65x) was sequenced along with one 2x150 HiC library (76.53x) (**table S5**). Prior to assembly, Illumina fragment reads were screened for phix contamination. Reads composed of >95% simple sequences were removed. Illumina reads <50 bp after trimming for adapter and quality (q<20) were removed. The final Illumina read set consists of 1,901,151,696 reads for a total high-quality base pair yield of 247.18x. For the PACBIO sequencing, a total raw sequence yield of 39.2 Gb, with a total coverage of 35.08x (**tables S5 and S6**).

For genome assembly and construction of pseudomolecule chromosomes, the version 1.0 assembly was generated by assembling the 2,058,145 PACBIO CCS reads (35.08x) using the HiFiAsm assembler (*79*, *80*) and subsequently polished using RACON (*34*). This produced an initial assembly consisting of 3,267 scaffolds (3,267 contigs), with a contig N50 of 31.6 Mb, and a total genome size of 1,234.5 Mb (**table S7**).

Hi-C Illumina reads from *Pharus latifolius* (var. MRM_UAlabama_McKain320), were separately aligned to the contigs with Juicer (*96*), and chromosome scale scaffolding was performed with 3D-DNA (*97*). No misjoins were identified in the assembly, and the contigs were then oriented, ordered, and joined together into 12 chromosomes using the HiC data. A total of 47 joins were applied to the assembly. Each chromosome join is padded with 10,000 Ns. Contigs terminating in significant telomeric sequence were identified using the $(TTTAGGG)_n$ repeat, and care was taken to make sure that they were properly oriented in the production assembly. The remaining scaffolds were screened against bacterial proteins, organelle sequences, GenBank nr and removed if found to be a contaminant. After forming the chromosomes, it was observed that some small (<20Kb) redundant sequences were present on adjacent contig ends within chromosomes. To resolve this issue, adjacent contig ends were aligned to one another using BLAT (*98*), and duplicate sequences were collapsed to close the gap between them. A total of 23 adjacent contig pairs were collapsed in the assembly.

Finally, homozygous SNPs and INDELs were corrected in the V1 release using ~40x of Illumina reads (2x150, 400bp insert) by aligning the reads using bwa-mem (*99*) and identifying homozygous SNPs and INDELs with the GATK's UnifiedGenotyper tool (*100*). A total of 291 homozygous SNPs and 12,285 homozygous INDELs were corrected in the V1 release. The final version 1.0 release contained 1,117.9 Mb of sequence, consisting of 222 contigs with a contig N50 of 57.1 Mb and a total of 99% of assembled bases in chromosomes (**table S8**).

Completeness of the euchromatic portion of the version 1.0 assembly was assessed by aligning the existing IsoSeq reads to the V1 release. The aim of this analysis is to obtain a measure of completeness of the assembly, rather than a comprehensive examination of gene space. The IsoSeq alignments indicate that 99.68% of the reads are aligned to the version 1 release.

For screening and final assembly releases, scaffolds that were not anchored in a chromosome were classified into bins depending on sequence content. Contamination was identified using blastn against the NCBI non-redundant nucleotide collection (NR/NT) and blastx using a set of known microbial proteins. Additional scaffolds were classified in the version 1 release as repetitive (>95% masked with 24mers that occur more than 4 times in the chromosomes) (815 scaffolds, 30.7 Mb), redundant (>95% masked with 24mers that occur 2 or more times in all scaffolds) (2,220 scaffolds, 85.5 Mb), prokaryote (9 scaffolds, 414.5 Kb), and mitochondria (1 scaffolds, 41.3 Kb).

***Genome sequencing of Typha latifolia***

The accession of *Typha latifolia*, CWD-2019.6, was collected from the Penn State Arboretum and vouchered by Claude dePamphilis in 2019. We sequenced *Typha latifolia* (var. 2019.6) using a whole genome shotgun sequencing strategy and standard sequencing protocols. Sequencing reads were collected using Illumina and PACBIO platforms. Illumina and PACBIO reads were sequenced at the HudsonAlpha Institute in Huntsville, Alabama. Illumina reads were sequenced using the Illumina NovoSeq6000 platform, and the PACBIO reads were sequenced using the REVIO platform. One 400 bp insert 2x150 Illumina fragment library (261.16x per haplotype) was sequenced along with one 2x150 OmniC library (230.01x per haplotype, **table S9**). Prior to assembly, Illumina fragment reads were screened for phix contamination. Reads composed of >95% simple sequences were removed. Illumina reads <50 bp after trimming for adapter and quality (q<20) were removed. The final read set consists of 755,162,004 reads for a total of 491.17x of high-quality Illumina bases. For the PACBIO sequencing, a total raw sequence yield of 22.4 Gb, with a total coverage of 104.23x per haplotype (**table S10**).

For genome assembly and construction of pseudomolecule chromosomes, the version 1.0 assemblies were generated by assembling the 1,066,852 PACBIO CCS reads (104.23x per haplotype) using the HiFiAsm assembler (*79*, *80*) and subsequently polished using RACON (*34*). This produced initial assemblies of both haplotypes. The HAP1 assembly consisted of 410 scaffolds (410 contigs), with a contig N50 of 11.7 Mb, and a total genome size of 235.5 Mb (**table S11**). The HAP2 assembly consisted of 421 scaffolds (421 contigs), with a contig N50 of 9.5 Mb, and a total genome size of 233.9 Mb (**table S12**).

Hi-C Illumina reads from *Typha latifolia* (var. 2019.6), were separately aligned to the HAP1 and HAP2 contig sets with Juicer (*96*), and chromosome scale scaffolding was performed with 3D-DNA (*97*). No misjoins were identified in either the HAP1 or HAP2 assemblies. The contigs were then oriented, ordered, and joined together into 15 chromosomes per haplotype using the HiC data. A total of 9 joins were applied to the HAP1 assembly, and 14 joins for the HAP2 assembly. Chromosomes were numbered by size from largest to smallest with the p-arm to the left. Each chromosome join is padded with 10,000 Ns. Contigs terminating in significant telomeric sequence were identified using the $(TTTAGGG)_n$ repeat, and care was taken to make sure that they were properly oriented in the production assembly. The remaining scaffolds were screened against bacterial proteins, organelle sequences, GenBank nr and removed if found to be a contaminant. After forming the chromosomes, it was observed that some small (<20 Kb) redundant sequences were present on adjacent contig ends within chromosomes. To resolve this issue, adjacent contig ends were aligned to one another using BLAT (*98*), and duplicate sequences were collapsed to close the gap between them. A total of 2 adjacent contig pairs were collapsed in the HAP1 assembly and 1 in the HAP2 assembly.

Finally, homozygous SNPs and INDELs were corrected in the HAP1 and HAP2 releases using ~60x of Illumina reads (2x150, 400 bp insert) by aligning the reads using bwa-mem (*99*) and identifying homozygous SNPs and INDELs with the GATK's UnifiedGenotyper tool (*100*). A total of 107 homozygous SNPs and 6,519 homozygous INDELs were corrected in the HAP1 release, while a total of 121 homozygous SNPs and 7,009 homozygous INDELs were corrected in the HAP2 release. The final version 1.0 HAP1 release contained 215.3 Mb of sequence, consisting of 30 contigs with a contig N50 of 9.9 Mb and a total of 99.975% of assembled bases integrated into chromosomes (**table S13**). The final version 1.0 HAP2 release contained 214.7 Mb of sequence, consisting of 20 contigs with a contig N50 of 13.5 Mb and a total of 100% of assembled bases in chromosomes (**table S14**).

Completeness of the euchromatic portion of the version 1.0 assemblies was assessed using rnaSEQ reads. The aim of this analysis is to obtain a measure of completeness of the assembly, rather than a comprehensive examination of gene space. The reads were aligned to the assembly using bwa-mem2 (*99*). The screened alignments indicate that 99.82% of the ranaSEQ reads are aligned to the V1 HAP1 release, and 99.83% aligned to the V1 HAP2 release.

***Gene tree reconstruction***

We identified putative orthogroups for gene models of 39 species (**table S1**) using OrthoFinder (v.2.5.4) (*101*) under default settings. Following McKain et al. (*3*), we filtered the orthogroups to have a minimum of 20 taxa. Amino acids for each orthogroup were aligned using MAFFT (v.7.490) (*102*) with the "auto" option for algorithm selection. The PAL/PTAL tree and other gene trees were reconstructed on these filtered alignments using IQ-TREE 2 (v.2.2.6) (*103*) allowing for the optimal model and 1000 bootstrap replicates to be estimated for each alignment. For building a coding sequence tree for the PAL/PTAL orthogroup, we used the aligned amino acid sequences to create a codon alignment of nucleotide sequences using default parameters of PAL2NAL (v.14) (*104*). Codon alignments were filtered so that the length of a gene sequence had to be at least 50% of the final alignment length and sequences were not permitted to have more than 30% gaps in the alignment (*3*). Then, the tree was generated using RAxML-NG (*105*) with the evolutionary model TVM+G4 determined by ModelTest-NG (*106*). To reconcile the PAL/PTAL gene tree with the species tree, we used TreeSolve (*54*) to generate optimal trees. For input, we used the species tree generated from OrthoFinder and generated an amino acid tree for PAL/PTAL genes from these species using RAxML-NG with the evolutionary model JTT+I+G4+F. We also used 100 bootstrap trees generated from RAxML-NG as input. Duplication-Transfer-Loss (DTL) reconciliation from TreeSolve showed that the optimally resolved gene tree has a lower reconciliation cost than the original gene tree (389 vs. 448) and hence is a more probable tree given the species tree.

***Identification and placement of polyploidy events in the species phylogeny***
A total of 10,055 multicopy gene trees were analyzed to identify potential polyploidy events and place them on the species tree obtained from Timilsena et al. (*30*) using the PUG software and algorithm (*3*). The "estimate_paralogs" option was selected in PUG allowing identification of all possible paralogs within a single gene tree by identifying all gene model pairs from the same taxon. Each multicopy gene tree was re-rooted to a preferred list of outgroups (*Amborella*, *Nymphaea*, *Cinnamon*, *Aquilegia*, *Vitis*, *Arabidopsis*, *Glycine*, *Populus*, *Beta*, *Spinacia*, *Solanum*, *Helianthus*, and *Acorus* in that preferred order. PUG attempts to use the farthest outgroup, as designated by the user, and works down the list until a sequence from that taxon is found. That sequence is then set as the outgroup for the gene tree. PUG identifies the most recent common ancestor node in the gene tree for each potential paralog pair, and the taxa composition of the subtree with the MRCA as the root is used to locate the equivalent node in the species tree. A duplication event placed on the species phylogeny was considered valid if two criteria were met: 1) the taxa above the node matched those in the gene tree and 2) at least one sister taxon to the queried node was found in both the species and gene trees. PUG does not consider potential duplications on tip branches of the species phylogeny, so species-specific duplications are ignored. PUG was run for all gene trees and putative paralogs across all species to identify all potential WGD events. Assessment for putative WGD events was limited to support from orthogroup tree nodes that have a bootstrap support value of 80 or more though we also report values for BSV 50 in the supplemental materials. Unique duplications were mapped on the species tree using the PUG_Figure_Maker.R script (https://github.com/mrmckain/PUG) on R v.4.3.3 and ape v.5.7-1 (*107*). All branches with duplication counts less than 25% of the largest count on a single branch (3,200) were ignored when mapping.

***Synteny analyses***
The GENESPACE analysis was conducted using v.1.2.3 (*35*), default parameters, and genome assemblies and annotations for *Ananas cosmosus*, *Joinvillea ascendens*, *Pharus latifolius*, *Oryza sativa*, *Sorghum bicolor*, and *Zea mays* (**Table S1**) as representatives of the Poales with well-scaffolded genomes.

Syntenic analyses of *PAL*/*PTAL* genes were performed using COGE (*108*) across selected Poales species with the following options: DAGchainer: relative gene order, maximum distance between 2 matches: 20 genes, minimum number of aligned pairs: 5 genes, merged syntenic blocks with quota align, window size 100 genes, use all genes in the target genome, calculate synonymous substitution rates, and tandem duplication distance is 10. Synteny was calculated between the following species: *B. distachyon* (version 556, 3.0) and *J. ascendens* (version 1.1); *B. distachyon* (version 556, 3.0) and *Z. mays* (version PH207 UMN 1.0); *B. distachyon* (version 556, 3.0) and *P. latifolius* (version 1.0); *B. distachyon* (version 556, 3.0) and *S. bicolor* (version 454, 3.0.1); *P. latifolius* (version 1.0) and *S. angustifolia* (version 1.1); *S. angustifolia* (version 1.1) and *J. ascendens* (version 1.1); *J. ascendens* (version 1.1) and *A. comosus* (version 3); *J. ascendens* (version 1.1) and *P. latifolius* (version 1.0).

***Identification of the residues involved in the transition from PAL to PTAL***
To find positively selected residues, we selected the node in the coding sequence tree of PAL/PTAL containing all PTAL genes including JaPTAL, but excluding JaPAL (red arrow in **fig. S3**). PAML v4.9 software (*58*) was then used to determine the positively selected residues of this clade.

***Cloning of PAL and PTAL candidates with and without mutations***
The coding sequences of PAL and PTAL candidate enzymes from *S. bicolor*, *B. distachyon*, *S. angustifolia* and *J. ascendens* were first amplified by nested PCR using the corresponding cDNA and gene specific primer with PrimeSTAR MAX DNA polymerase (Takara Bio). The PCR fragments were gel-purified and then used as a PCR template for the second PCR with additional primers with in-fusion tag. The obtained PCR products were inserted into the pET28a vector at EcoRI and NdeI sites using the in-fusion HD Cloning Kit (Takara Bio) according to the manufacturer's instruction. The obtained plasmid vectors were submitted for the sequence analysis and confirmed that the coding sequences were matched with the ones in the database. BdPTAL, EmoPTAL, EmoPAL, $JaPAL^{F140H\text{-}MUT8}$, and $JaPAL^{F140H\text{-}MUT16}$ were gene synthesized and cloned into the pET28a vector (SynbioTech). For site-directed mutagenesis, the 1:100 diluted plasmid and mutagenesis primers were used for the PCR amplification with PrimeSTAR MAX DNA polymerase (Takara Bio). The primers used in this study are listed in **table S15**.

***Recombinant protein expression and purification***
For the recombinant protein expression, the cloned pET28a vectors were transformed into Rosetta-2 (DE3) *E. coli* and cultured in 3 ml of TB medium containing kanamycin (50 µg/ml), chloramphenicol (34 µg/ml), and 0.1 % glucose at 37 ℃ and 200 rpm. After the overnight cultivation, 500 µl of pre-culture solution were added to 50 ml TB medium containing the same antibiotics and further cultured at 27 °C and 200 rpm until the OD600 reached 0.5-0.7. After the bacterial cultures were cooled down on ice, isopropyl β-D-1-thiogalactopyranoside (IPTG, 0.5 mM of final concentration) incubated at 22 ℃ under the constant shaking at 200 rpm. After 24 h, the cultures were harvested by centrifugation (5000 g, 5 min, 4 °C) and the pellets were frozen at -30 °C. The pellets were thawed and resuspended with the lysis buffer containing 50 mM sodium phosphate buffer (pH 8.0), 300 mM NaCl, 10% glycerol, and 0.25 mg of lysozyme. After 30 min incubation on ice, the suspension was sonicated three times for 20 sec and the supernatant was recovered after the centrifugation (12500 g, 20 min, 4 °C). The supernatants were added to the new tube containing 100 µl of Ni-NTA beads (Millipore) and incubate the mixture at 25 °C for 30 min under constant inversion. After the unbound proteins were washed away for three times with washing buffer containing 50 mM sodium phosphate buffer (pH 8.0), 300 mM NaCl, 10% glycerol, and 10 mM imidazole, target proteins were eluted with elution buffer containing 50 mM sodium phosphate buffer (pH 8.0), 300 mM NaCl, 10% glycerol, and 300 mM imidazole. The collected enzyme solutions were desalted by using sephadex G-50 column (GE Healthcare). The protein concentration was determined using the BioRad protein assay dye (BioRad). The purity was confirmed to be >90% by SDS-PAGE and calculated by using ImageJ software.

***PAL and TAL enzyme assays***
All the substrate solutions were prepared with 0.01N NaOH to increase the solubility of L-Tyr. The mixture containing 100 mM Tris-HCl (pH 8.5), 1% glycerol and the purified enzyme in a total volume of 50 µl was preincubate for 3 min at 30 °C. PAL and TAL reactions were started by addition of the 50 µl of 1 mM substrate (L-Phe or L-Tyr, respectively) and incubated at 30 °C for 20 min unless otherwise

noted. The reaction was terminated by addition of 6N AcOH (10 µl).

The reaction products were analyzed using HPLC (1200 Infinitely Series - Infinitely better, Agilent Technologies) to directly detect the final products of PAL and TAL assay, cinnamic acid and *p*-coumaric acid, respectively. Analytical conditions were as follows: column, Neptune T3 C18 column (3 µm, 2.1 × 150 mm, ES industries); solvent system, solvent A (water including 0.1 % [v/v] formic acid) and solvent B (acetonitrile including 0.1 % [v/v] formic acid); gradient program: 99% A/1% B at 0 min, 99% A/1% B at 4.5 min, 95% A/5% B at 7.5 min, 85% A/15% B at 12 min, 75% A/25% B at 16.5 min, 70% A/30% B at 21 min, 5% A/95% B at 23 min, 5% A/95% B at 26 min, 99% A/5% B at 26.5 min, and 99% A/5% B at 30 min; flow rate: 0.3 mL/min; DAD: 275 nm for cinnamic acid, 309 nm for *p*-coumaric acid.

The kinetics parameters of the recombinant enzymes were determined using HPLC. The mixture contains 100 mM Tris-HCl (pH 8.5), 1% glycerol and purified enzymes (0.15 µg for PAL assay and 1 µg for TAL assay) in a 50 µl total volume were preincubate for 3 min at 30 ℃. PAL and TAL reactions were started by addition of 50 µl substrate solution prepared with the concentration range of 0-4 mM for L-Phe and 0-2 mM for L-Tyr. After 10 min and 20 min incubation for PAL and TAL assay, respectively, at 30 ℃, the reaction was terminated by addition of 6N AcOH (10 µl). Analytical conditions were as follows: column, Atlantis T3 C18 column (3 µm, 2.1 × 150 mm, Waters); solvent system, solvent A (water including 0.1 % [v/v] formic acid) and solvent B (acetonitrile including 0.1 % [v/v] formic acid); gradient program: 85% A/15% B at 0 min, 85% A/15% B at 1 min, 70% A/30% B at 3 min, 15% A/95% B at 6.5 min, 15% A/95% B at 7.5 min, 85% A/15% B at 8.5 min, and 85% A/15% B at 10 min; flow rate: 0.4 mL/min; DAD: 275 nm for cinnamic acid, 309 nm for *p*-coumaric acid. The products were quantified on the basis of the calibration curves generated with the authentic standards. Non-linear hyperbolic regression analyses were conducted by using the Excel Solver tool to calculate $K_m$ and $k_{cat}$ values.

***Protein modeling analysis***

The structures of JaPAL and JaPTAL were generated by SWISS-MODEL (*109*) using homo-tetrameric PAL structure from parsley 6F6T.pdb (*60*) and homo-dimeric PTAL structure from sorghum 6AT7.pdb (*56*), respectively, as the templates. The sequence identity against each template were 77.3 % and 80.5 % for JaPAL and JaPTAL, respectively.

***Data analysis***

For the PAL/PTAL enzyme assay, statistical significance was determined by the statistical tests indicated at the figure legend. Student's *t*-test and ANOVA with post hoc Tukey–Kramer test or Kruskal-Wallis nonparametric test were conducted using Microsoft Excel and R, respectively. Key resources, including software and algorisms, used in this study are summarized in **table S16**.

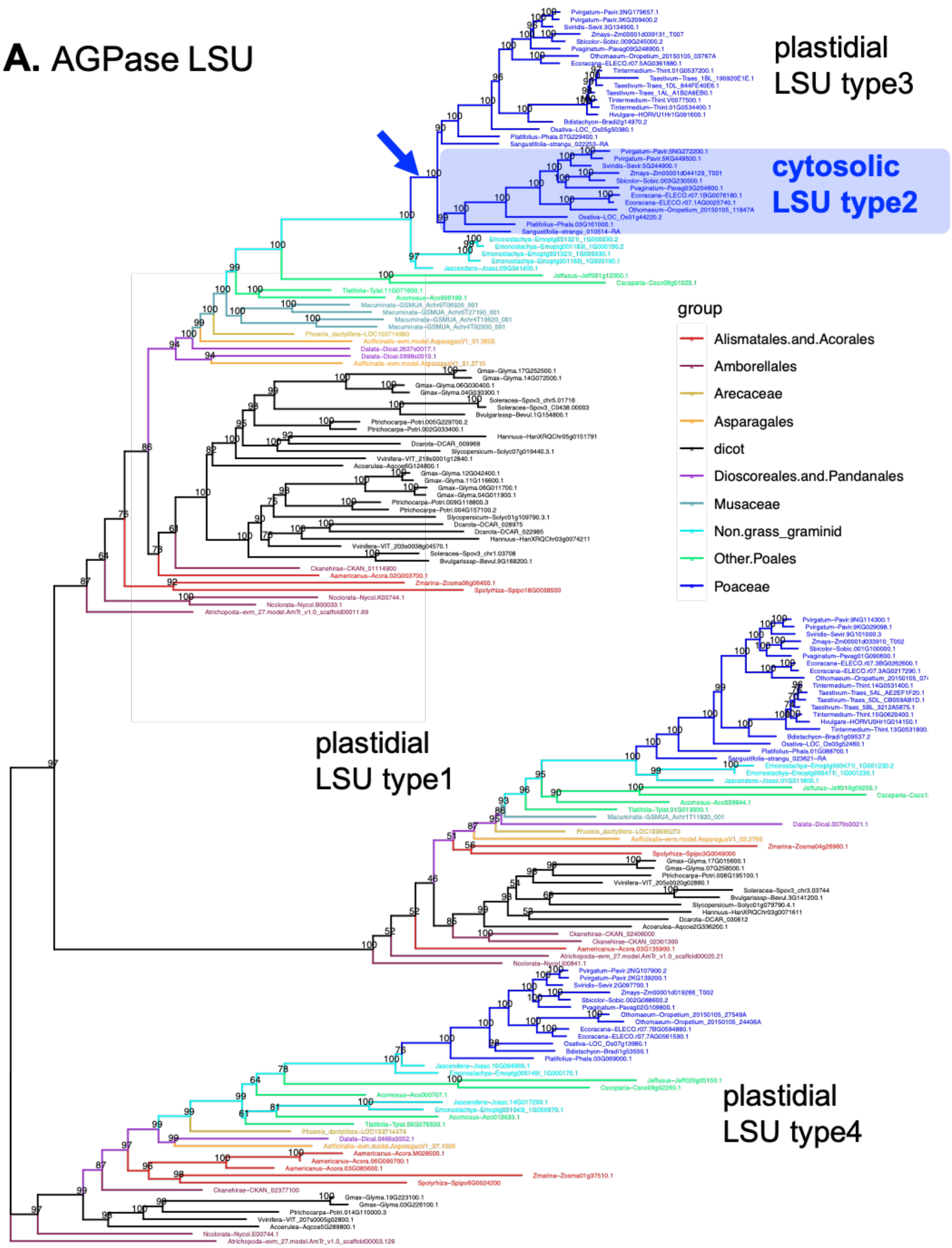
A. AGPase LSU
plastidial LSU type3
cytosolic LSU type2
plastidial LSU type1
plastidial LSU type4
group
Alismatales.and.Acorales
Amborellales
Arecaceae
Asparagales
dicot
Dioscoreales.and.Pandanales
Musaceae
Non.grass_graminid
Other.Poales
Poaceae

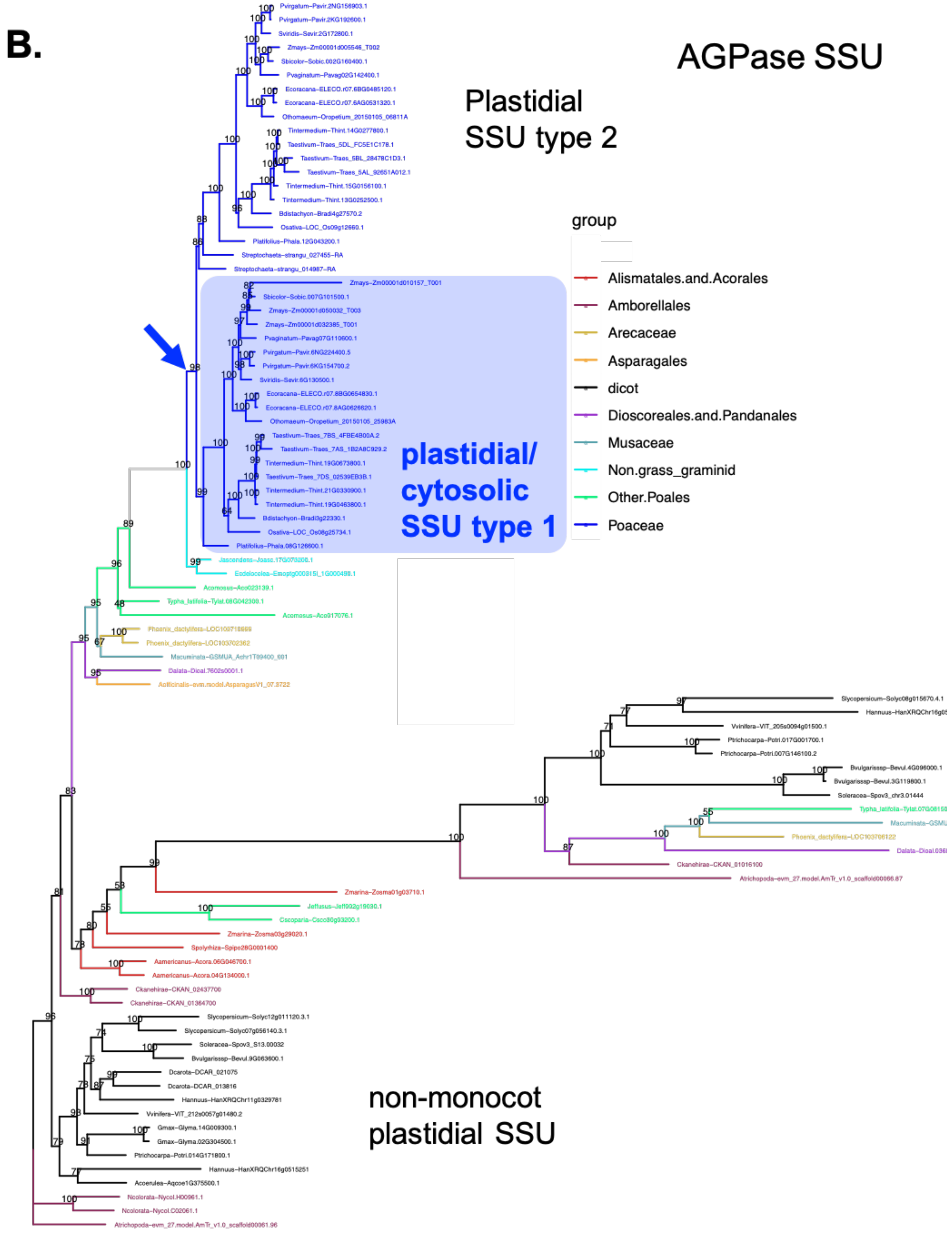
B.
AGPase SSU
Plastidial
SSU type 2
group
Alismatales.and.Acorales
Amborellales
Arecaceae
Asparagales
dicot
Dioscoreales.and.Pandanales
Musaceae
Non.grass_graminid
Other.Poales
Poaceae
plastidial/
cytosolic
SSU type 1
non-monocot
plastidial SSU

c. ADG transporter

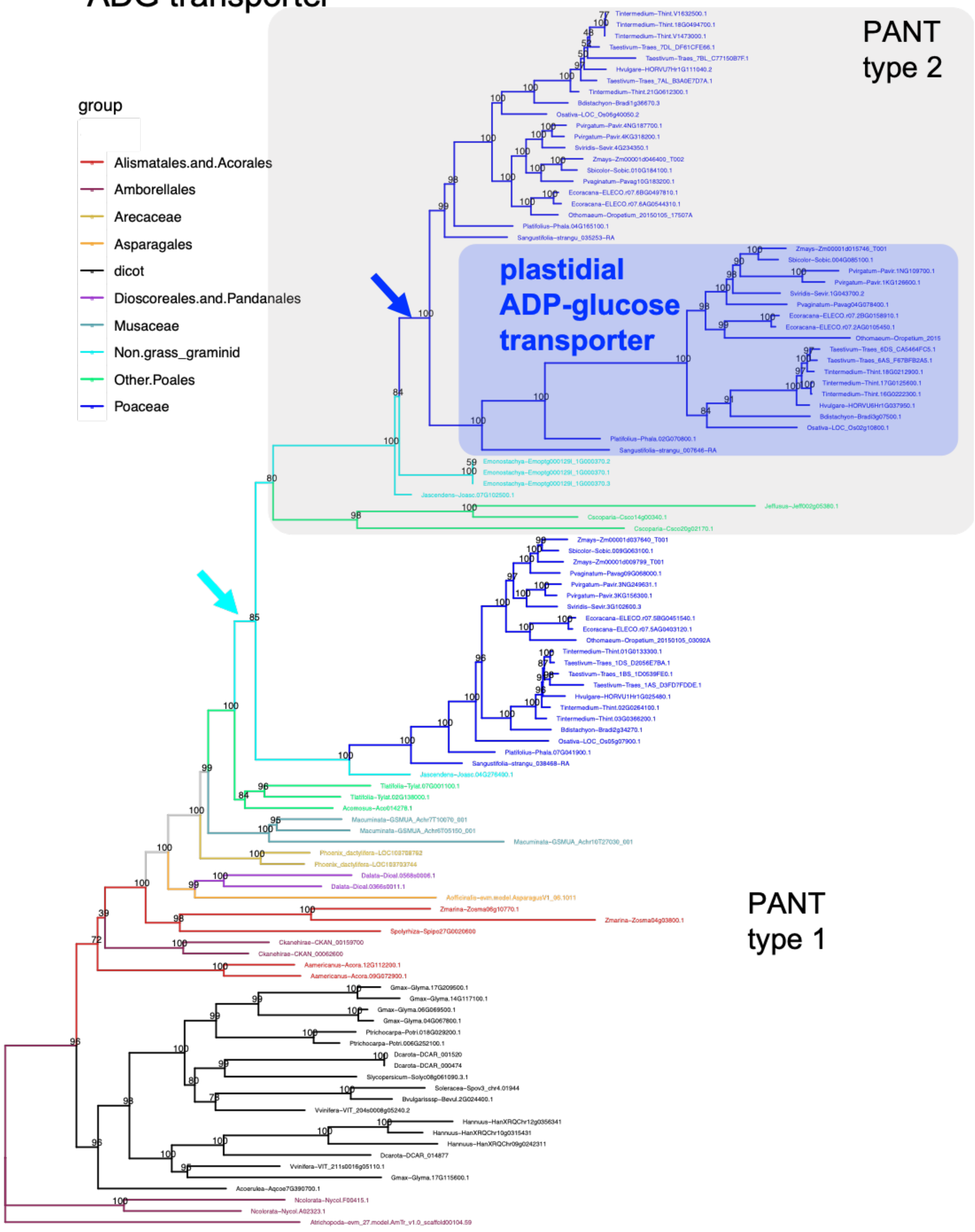

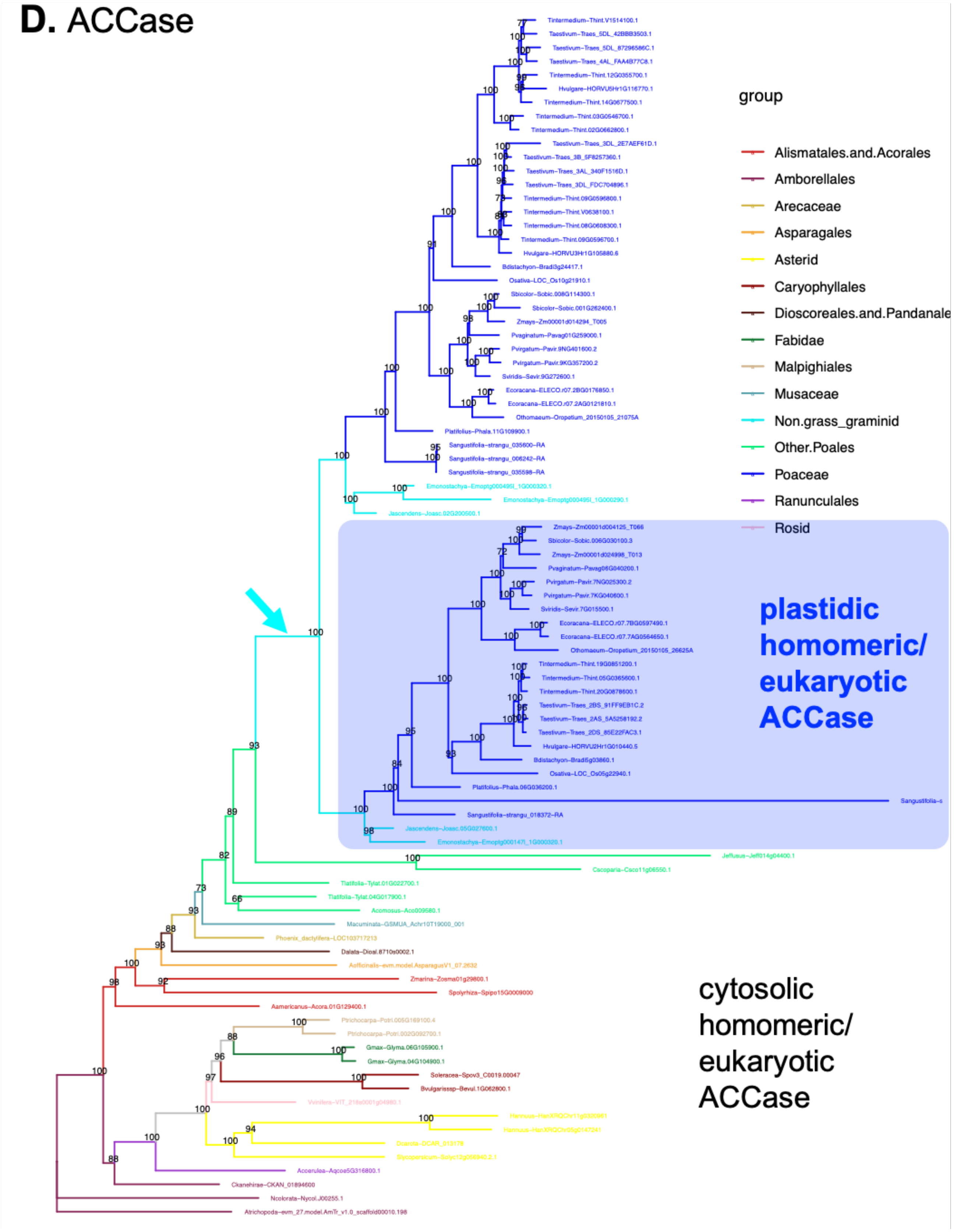
D. ACCase
group
Alismatales.and.Acorales
Amborellales
Arecaceae
Asparagales
Asterid
Caryophyllales
Dioscoreales.and.Pandanale
Fabidae
Malpighiales
Musaceae
Non.grass_graminid
Other.Poales
Poaceae
Ranunculales
Rosid
plastidic homomeric/ eukaryotic ACCase
cytosolic homomeric/ eukaryotic ACCase

**Fig. S1. Full-scaled gene trees for genes involved in starch and fatty acid biosynthesis**

Orthogroups were determined using OrthoFinder, then Maximum Likelihood (ML) gene trees were built from genes in the orthogroup using IQ-tree. Gene trees included are (**A**) the large subunit of ADP-glucose pyrophosphatase (AGPase LSU), (**B**) the small subunit of AGPase (AGPase SSU), and (**C**) ADP-glucose transporters and plastidial Adenine nucleotide transporters (PANT), involved in starch biosynthesis, as well as (**D**) acetyl-CoA carboxylase (ACCase) involved in fatty acid biosynthesis. Representative sequences from grasses, Poales, as well as other monocots and angiosperms were included in the analysis. Blue and gray boxes denote clades specific to Poaceae and non-grass graminids that likely contributed to the evolution of their unique starch and fatty acid biosynthetic pathways. Blue and sky blue arrows indicate the underlying duplication events that occurred after and before the emergence of grasses, respectively.

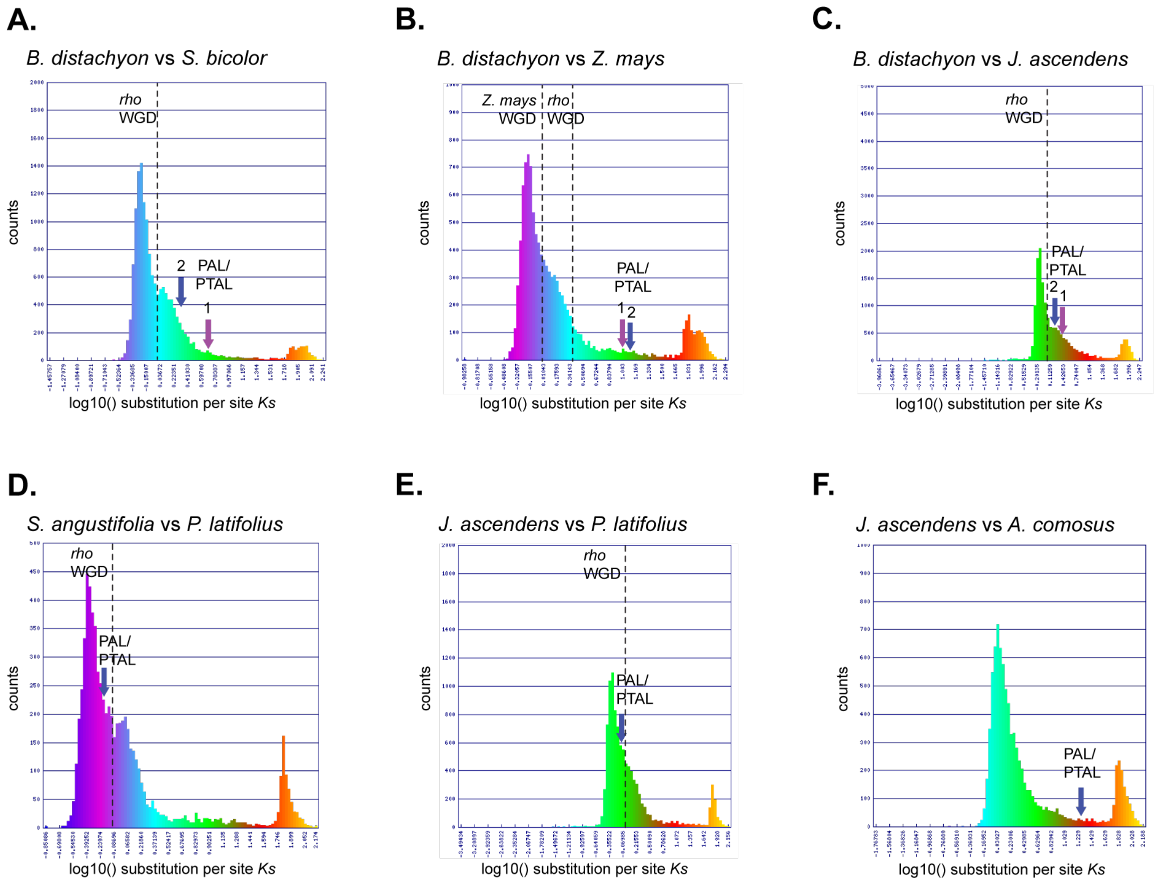


**Fig. S2. Distributions of synonymous substitution rates (*Ks*) across syntenic genes between Poales species.**

Distributions visualized using CoGe interface (https://genomevolution.org/coge/) (**A**) *Ks* between *Brachypodium distachyon* and *Sorghum bicolor*. (**B**) *Ks* between *B. distachyon* and *Zea mays*. (**C**) *Ks* between *B. distachyon* and *Joinvillea ascendens*. (**D**) *Ks* between *Streptochaeta angustifolia* and *Pharus latifolius*. (**E**) *Ks* between *J. ascendens* and *P. latifolius*. (**F**) *Ks* between *J. ascendens* and *Ananas comosus*. *PAL/PTAL Ks* rate indicated by arrows.

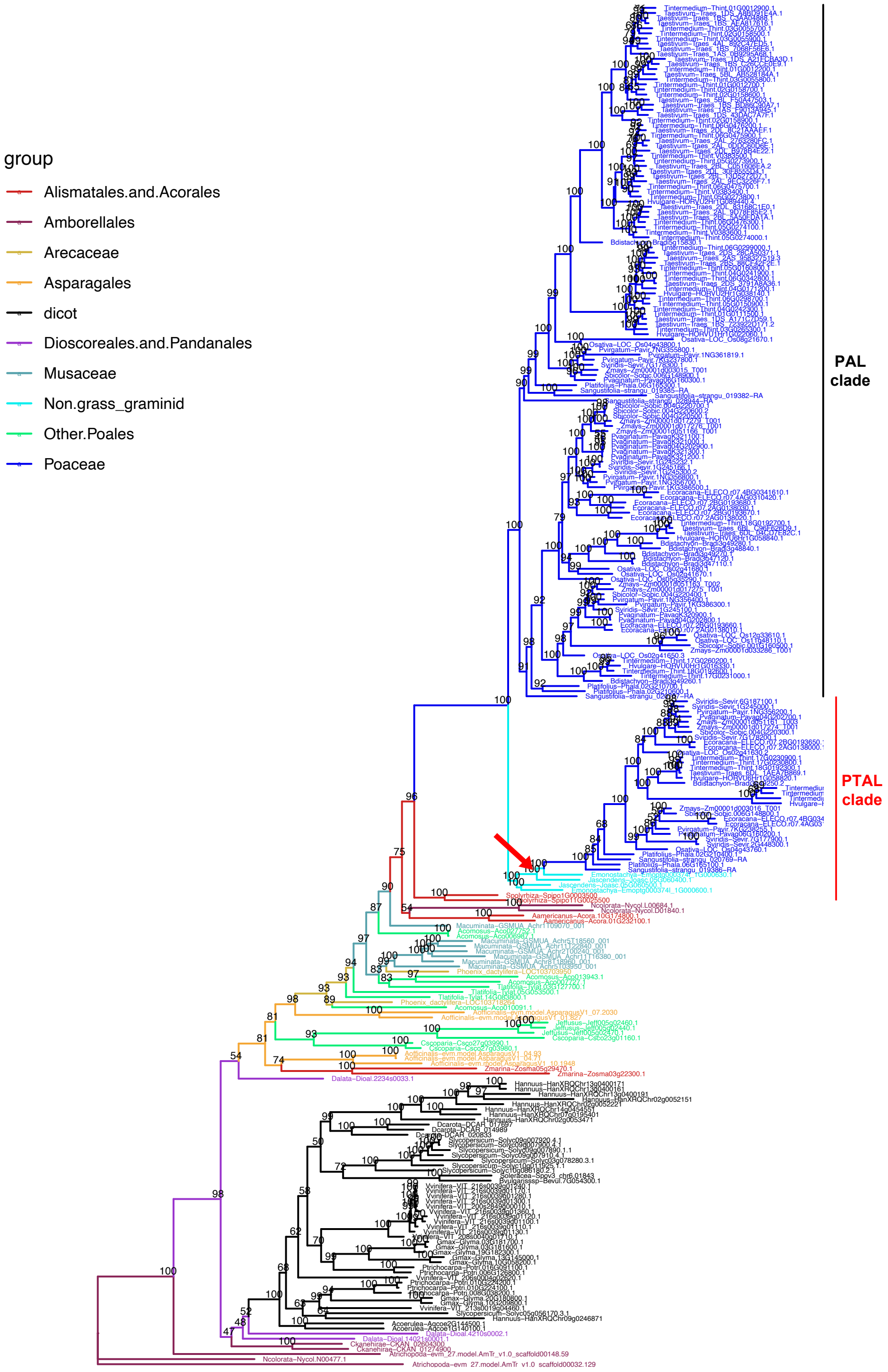


**Fig. S3. Amino acid based phylogenetic tree of *PAL*/*PTAL* genes.**

*PAL*/*PTAL* orthogroup was determined using OrthoFinder, then Maximum Likelihood (ML) gene trees were built from genes in the orthogroup using IQ-tree. PTAL clade includes genes where PTAL function is known in grasses, whereas PAL clade includes genes where only PAL function is known in grasses. The red arrow indicates the node used for the positive selection analysis.

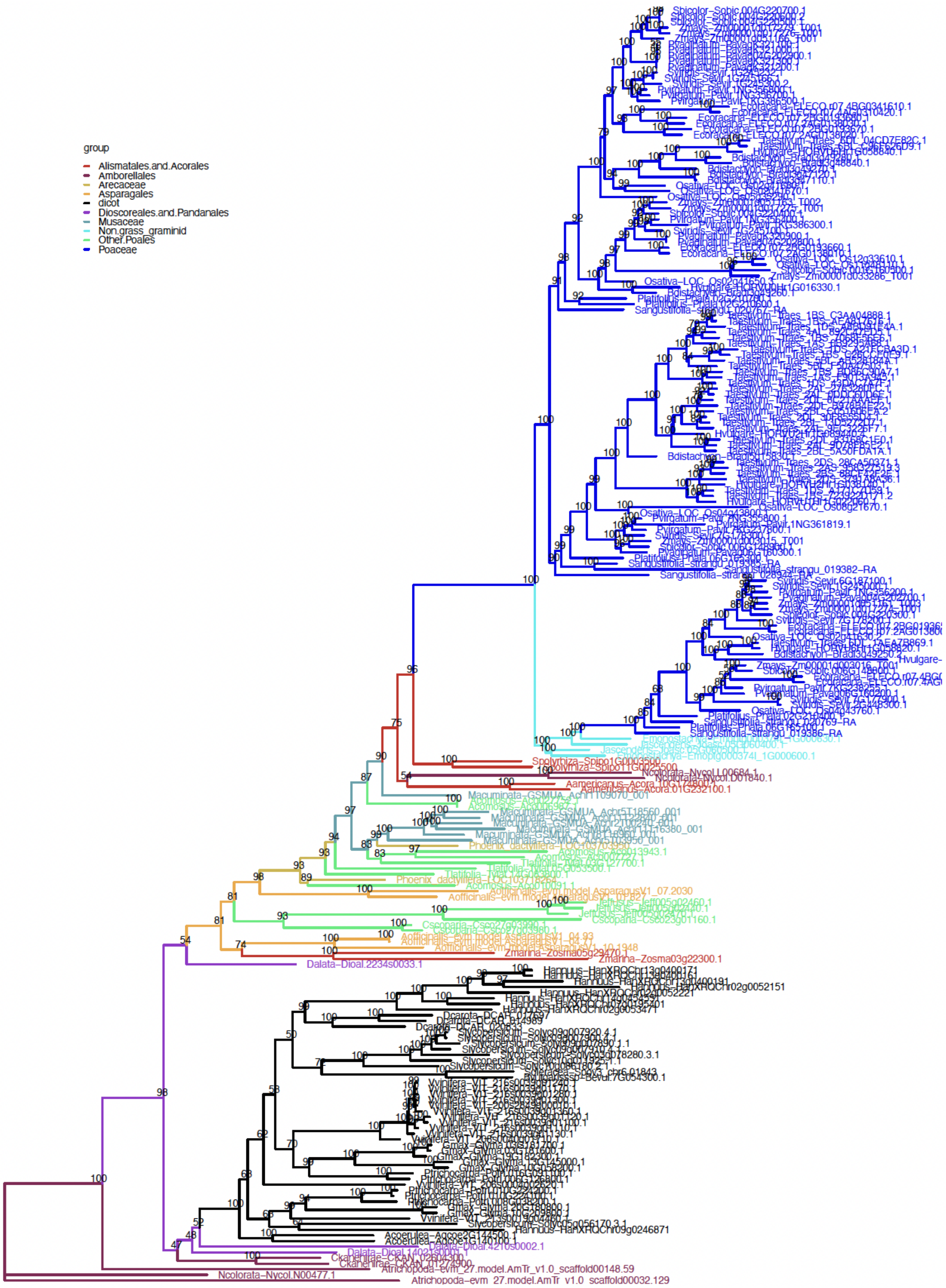


**Fig. S4. Coding sequence based phylogenetic tree of *PAL*/*PTAL* genes.**

Coding sequences from the *PAL*/*PTAL* orthogroup were aligned to the protein sequences using PAL2NAL. After alignment, IQ-tree was used to build the tree.

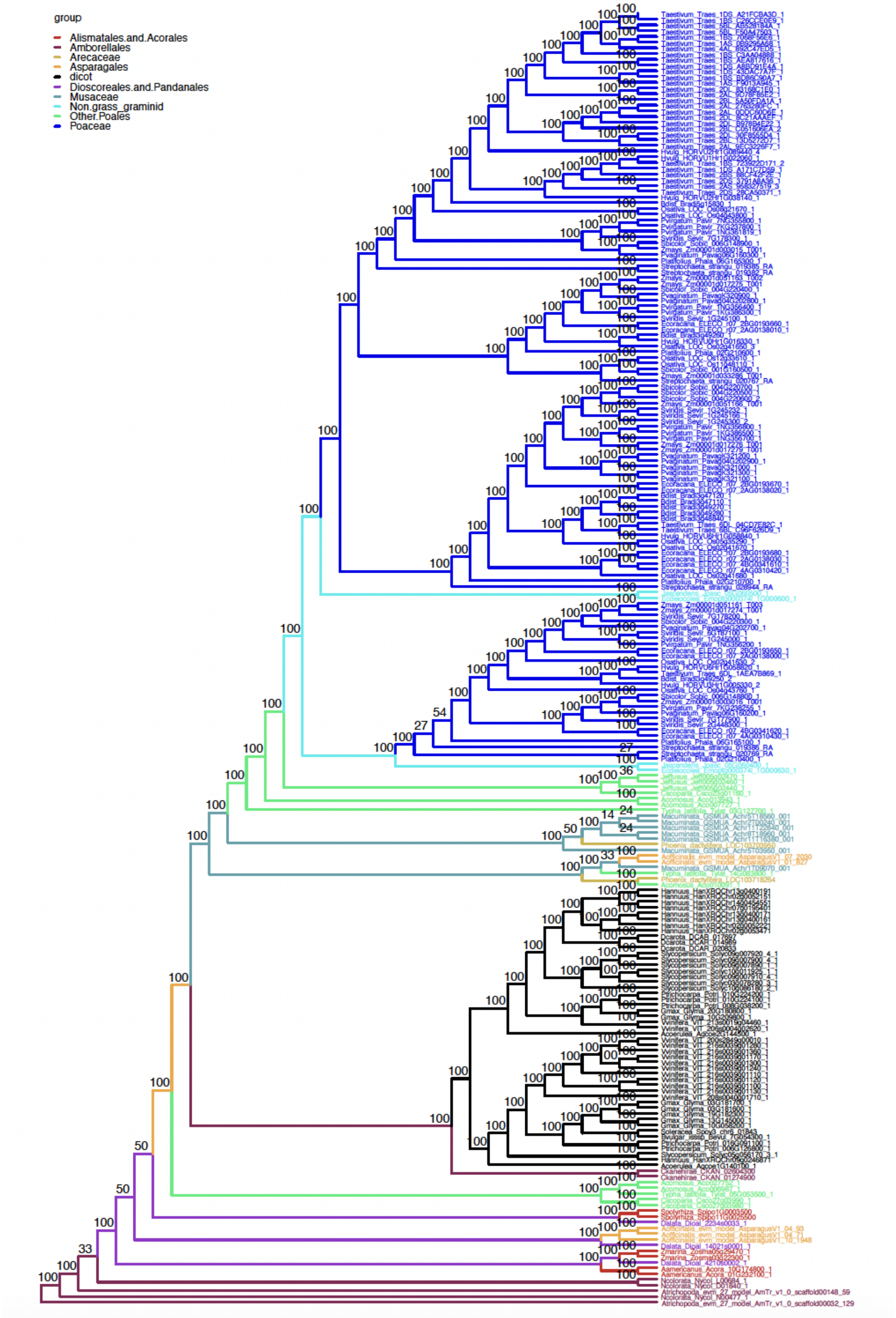


**Fig. S5. Amino acid based *PAL*/*PTAL* genes tree** reconciled with **TreeSolve**.

Final protein sequence tree based on TreeSolve. TreeSolve attempts to reconcile the gene tree with the species tree, and the optimal consensus tree places *PAL* genes from *J. ascendens* and *E. monostachya* in the PAL clade.

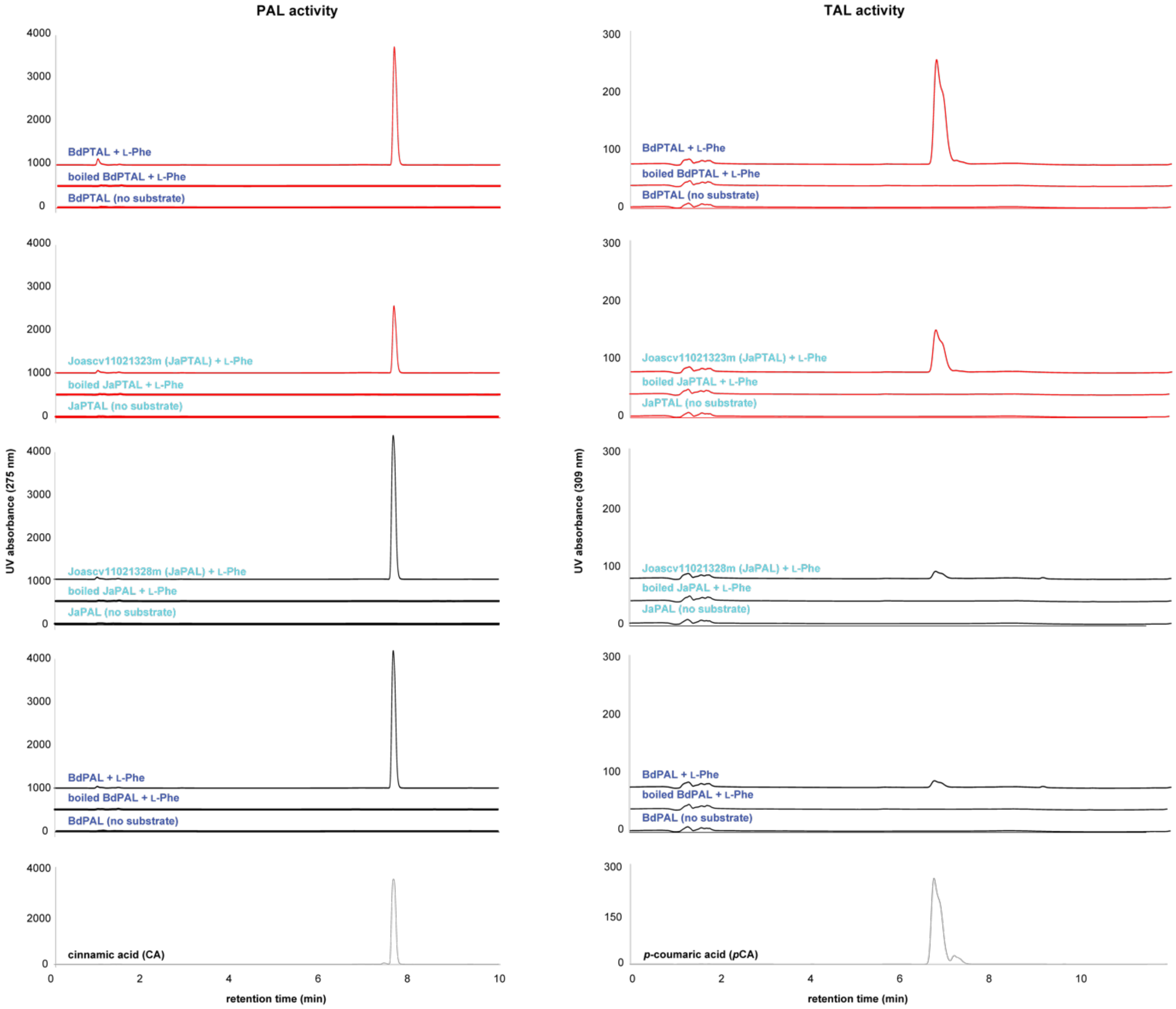


**Fig. S6. HPLC trace of PAL and TAL activity assay for PTALs/PALs from *B. distachyon* and *J. ascendens*.**

Purified PAL/PTAL recombinant enzymes were incubated with 1 mM substrate (L-Phe or L-Tyr). For negative controls, two reactions were performed with thermally inactivated recombinant enzymes and without substrate.

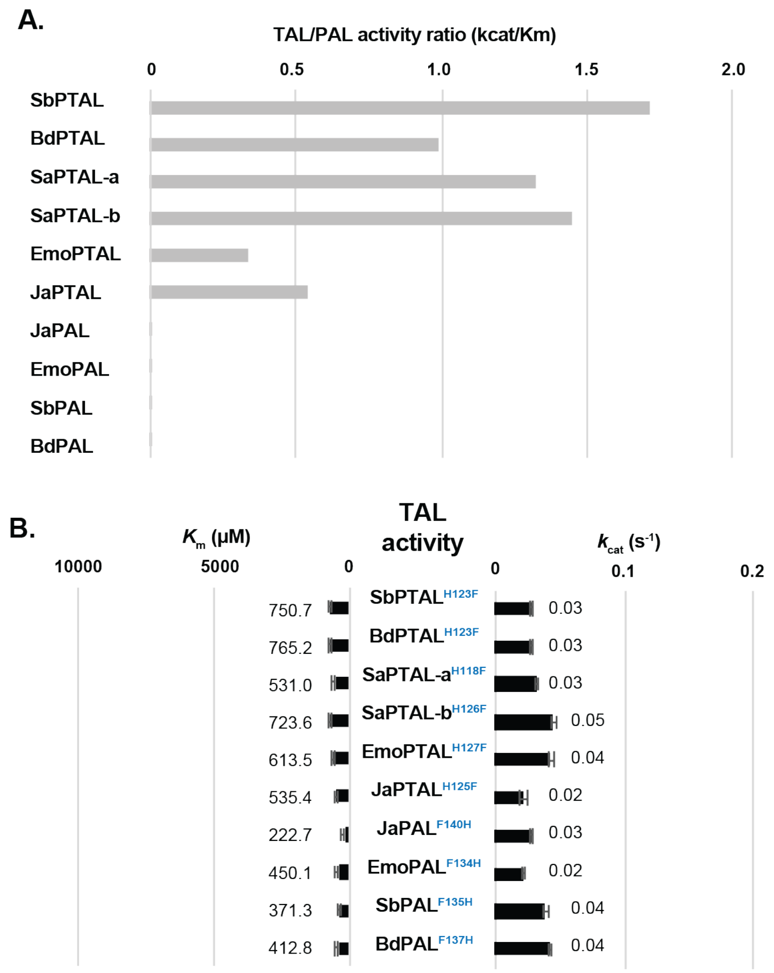


**Fig. S7. Kinetics assay for PALs/PTALs from grasses and non-grass graminids.**

(**A**) Ratio of TAL and PAL activity ($k_{cat}/K_m$) tends to be increased with the evolution of PTALs in graminids.

(**B**) $K_m$ and $k_{cat}$ of TAL activity for site-directed mutants of PTALs/PALs at His140 site, suggesting the involvement of additional residue(s) other than His140 in the efficient TAL activity.

```
Aamericanus-Acora.01G232100.1 ------------------------------------------------------------
Aamericanus-Acora.10G174800.1 ------------------------------------------------------------
Acomosus-Aco013943.1 ------------------------------------------------------------
Aofficinalis-evm.model.AsparagusV1_01.827 ------------------------------------------------------------
Typha_latifolia-Tylat.14G083800.1 ------------------------------------------------------------
Aofficinalis-evm.model.AsparagusV1_04.71 ------------------------------------------------------------
Aofficinalis-evm.model.AsparagusV1_04.93 ------------------------------------------------------------
Aofficinalis-evm.model.AsparagusV1_10.1948 ------------------------------------------------------------
Typha_latifolia-Tylat.03G127700.1 ------------------------------------------------------------
Spolyrhiza-Spipo11G0025500 ------------------------------------------------------------
Spolyrhiza-Spipo1G0003500 ------------------------------------------------------------
Phoenix_dactylifera-LOC103718264 ------------------------------------------------------------
Macuminata-GSMUA_Achr1T09070_001 ------------------------------------------------------------
Acomosus-Aco006987.1 ------------------------------------------------------------
Acomosus-Aco027752.1 ------------------------------------------------------------
Cscoparia-Csco27g03980.1 ------------------------------------------------------------
Cscoparia-Csco27g03990.1 ------------------------------------------------------------
Macuminata-GSMUA_Achr11T16380_001 ------------------------------------------------------------
Macuminata-GSMUA_Achr11T22840_001 ------------------------------------------------------------
Macuminata-GSMUA_Achr2T00240_001 ------------------------------------------------------------
Macuminata-GSMUA_Achr5T18560_001 ------------------------------------------------------------
Macuminata-GSMUA_Achr8T18960_001 ------------------------------------------------------------
Macuminata-GSMUA_Achr5T03950_001 ------------------------------------------------------------
Phoenix_dactylifera-LOC103703950 ------------------------------------------------------------
Acoerulea-Aqcoe1G140100.1 ------------------------------------------------------------
Acoerulea-Aqcoe2G144500.1 ------------------------------------------------------------
Typha_latifolia-Tylat.05G053500.1 ------------------------------------------------------------
Zmarina-Zosma05g29470.1 ------------------------------------------------------------
Acomosus-Aco007727.1 ------------------------------------------------------------
Bvulgarisssp-Bevul.7G054300.1 ------------------------------------------------------------
Bdistachyon-Bradi3g47110.1 ------------------------------------------------------------
Bdistachyon-Bradi3g47120.1 ------------------------------------------------------------
Bdistachyon-Bradi3g48840.1 ------------------------------------------------------------
Bdistachyon-Bradi3g49280.1 ------------------------------------------------------------
Hvulgare-HORVU6Hr1G058840.1 ------------------------------------------------------------
Taestivum-Traes_6BL_C96F626D9.1 ------------------------------------------------------------
Tintermedium-Thint.18G0192700.1 ------------------------------------------------------------
Taestivum-Traes_6DL_04CD7E82C.1 ------------------------------------------------------------
Bdistachyon-Bradi3g49270.1 ------------------------------------------------------------
Ecoracana-ELECO.r07.2AG0138020.1 ------------------------------------------------------------
Ecoracana-ELECO.r07.2BG0193670.1 ------------------------------------------------------------
Osativa-LOC_Os02g41670.1 ------------------------------------------------------------
Osativa-LOC_Os02g41680.1 ------------------------------------------------------------
Pvaginatum-Pavag04G202900.1 ------------------------------------------------------------
Pvaginatum-PavagK321000.1 ------------------------------------------------------------
Pvaginatum-PavagK321300.1 ------------------------------------------------------------
Pvaginatum-PavagK321100.1 MLYAPTGSHVLVVVGLVSTLHTAYHTTPSQAQLFNPRHSHQLSAGRAA------------
Pvaginatum-PavagK321200.1 ------------------------------------------------------------
Pvirgatum-Pavir.1KG386500.1 ------------------------------------------------------------
Pvirgatum-Pavir.1NG356700.1 ------------------------------------------------------------
Sviridis-Sevir.1G245166.1 ------------------------------------------------------------
Sviridis-Sevir.1G245232.1 ------------------------------------------------------------
Pvirgatum-Pavir.1NG356800.1 ------------------------------------------------------------
Sbicolor-Sobic.004G220500.1 ------------------------------------------------------------
Sbicolor-Sobic.004G220600.2 ------------------------------------------------------------
Sbicolor-Sobic.004G220700.1 ------------------------------------------------------------
Sviridis-Sevir.1G245300.2 ------------------------------------------------------------
Zmays-Zm00001d017276_T001 ------------------------------------------------------------
Zmays-Zm00001d017279_T001 ------------------------------------------------------------
Zmays-Zm00001d051166_T001 ------------------------------------------------------------
Ecoracana-ELECO.r07.2AG0138030.1 ------------------------------------------------------------
Ecoracana-ELECO.r07.2BG0193680.1 ------------------------------------------------------------
Osativa-LOC_Os05g35290.1 ------------------------------------------------------------
Platifolius-Phala.02G210700.1 ------------------------------------------------------------
Ecoracana-ELECO.r07.4AG0310420.1 ------------------------------------------------------------
Ecoracana-ELECO.r07.4BG0341610.1 ------------------------------------------------------------
Bdistachyon-Bradi3g49260.1 ------------------------------------------------------------
Hvulgare-HORVU0Hr1G016330.1 ------------------------------------------------------------
Tintermedium-Thint.17G0260200.1 ------------------------------------------------------------
Tintermedium-Thint.18G0192600.1 ------------------------------------------------------------
Tintermedium-Thint.17G0231000.1 ------------------------------------------------------------
Ecoracana-ELECO.r07.2AG0138010.1 ------------------------------------------------------------
Ecoracana-ELECO.r07.2BG0193660.1 ------------------------------------------------------------
Osativa-LOC_Os02g41650.3 ------------------------------------------------------------
Pvaginatum-Pavag04G202800.1 ------------------------------------------------------------
Pvaginatum-PavagK320900.1 ------------------------------------------------------------
Pvirgatum-Pavir.1NG356400.1 ------------------------------------------------------------
Sviridis-Sevir.1G245100.1 ------------------------------------------------------------
Sbicolor-Sobic.004G220400.1 ------------------------------------------------------------
Zmays-Zm00001d017275_T001 ------------------------------------------------------------
Zmays-Zm00001d051163_T002 ------------------------------------------------------------
Pvirgatum-Pavir.1KG386300.1 ------------------------------------------------------------
Platifolius-Phala.02G210600.1 ------------------------------------------------------------
Streptochaeta-strangu_020767-RA ------------------------------------------------------------
Bdistachyon-Bradi5g15830.1 ------------------------------------------------------------
Hvulgare-HORVU2Hr1G089440.4 ------------------------------------------------------------
Taestivum-Traes_2AL_0DDC60D6E.1 ------------------------------------------------------------
Tintermedium-Thint.06G0475900.1 ------------------------------------------------------------
Tintermedium-Thint.06G0476200.1 ------------------------------------------------------------
Taestivum-Traes_2AL_2763280FC.1 ------------------------------------------------------------
Taestivum-Traes_2DL_8C21AAAEF.1 ------------------------------------------------------------
Tintermedium-Thint.V0383500.1 ------------------------------------------------------------
Taestivum-Traes_2AL_9EC3226F7.1 ------------------------------------------------------------
Tintermedium-Thint.05G0273800.1 ------------------------------------------------------------
Tintermedium-Thint.V0383400.1 ------------------------------------------------------------
Taestivum-Traes_2BL_13D5272D7.1 ------------------------------------------------------------
Tintermedium-Thint.06G0475700.1 ------------------------------------------------------------
Taestivum-Traes_2BL_C051606EA.2 ------------------------------------------------------------
Tintermedium-Thint.V0383600.1 ------------------------------------------------------------
Taestivum-Traes_2AL_9D78F85E2.1 ------------------------------------------------------------
Tintermedium-Thint.05G0274100.1 ------------------------------------------------------------
Tintermedium-Thint.06G0476300.1 ------------------------------------------------------------
Taestivum-Traes_2BL_5A50FDA1A.1 ------------------------------------------------------------
Tintermedium-Thint.05G0273900.1 ------------------------------------------------------------
Taestivum-Traes_2DL_30F8555D4.1 ------------------------------------------------------------
Taestivum-Traes_2DL_83168C1E0.1 ------------------------------------------------------------
Taestivum-Traes_2DL_B978B4E22.1 ------------------------------------------------------------
Taestivum-Traes_1AS_0B9295A68.1 ------------------------------------------------------------
Taestivum-Traes_1BS_7068F56E6.1 ------------------------------------------------------------
Taestivum-Traes_1BS_AEA817616.1 ------------------------------------------------------------
Taestivum-Traes_4AL_892C47ED5.1 ------------------------------------------------------------
Tintermedium-Thint.03G0055900.1 ------------------------------------------------------------
Tintermedium-Thint.01G0012900.1 ------------------------------------------------------------
Tintermedium-Thint.02G0158500.1 ------------------------------------------------------------
Tintermedium-Thint.03G0055700.1 ------------------------------------------------------------
Taestivum-Traes_1BS_C3AA04888.1 ------------------------------------------------------------
Taestivum-Traes_1DS_A8BD91E4A.1 ------------------------------------------------------------
Taestivum-Traes_1BS_C26CCE0E9.1 ------------------------------------------------------------
Taestivum-Traes_1DS_A21FCBA3D.1 ------------------------------------------------------------
Tintermedium-Thint.01G0012200.1 ------------------------------------------------------------
Tintermedium-Thint.02G0158600.1 ------------------------------------------------------------
Tintermedium-Thint.01G0012700.1 ------------------------------------------------------------
Taestivum-Traes_5BL_F50A47503.1 ------------------------------------------------------------
Tintermedium-Thint.02G0158700.1 ------------------------------------------------------------
Taestivum-Traes_5BL_AB528184A.1 ------------------------------------------------------------
Tintermedium-Thint.03G0055800.1 ------------------------------------------------------------
Taestivum-Traes_1AS_F9013A945.1 ------------------------------------------------------------
Taestivum-Traes_1DS_43DAC7A7F.1 ------------------------------------------------------------
Taestivum-Traes_1BS_BD86C90A7.1 ------------------------------------------------------------
```

```
Tintermedium-Thint.02G0158900.1                   ------------------------------------------------------------
Tintermedium-Thint.05G0274000.1                   ------------------------------------------------------------
Osativa-LOC_Os04g43800.1                          ------------------------------------------------------------
Pvaginatum-Pavag06G160300.1                       ------------------------------------------------------------
Sbicolor-Sobic.006G148900.1                       ------------------------------------------------------------
Pvirgatum-Pavir.7KG237800.1                       ------------------------------------------------------------
Pvirgatum-Pavir.7NG355800.1                       ------------------------------------------------------------
Sviridis-Sevir.7G178300.1                         ------------------------------------------------------------
Zmays-Zm00001d003015_T001                         ------------------------------------------------------------
Hvulgare-HORVU1Hr1G022060.1                       ------------------------------------------------------------
Taestivum-Traes_1BS_723922D171.2                  ------------------------------------------------------------
Taestivum-Traes_1DS_A171C7D59.1                   ------------------------------------------------------------
Tintermedium-Thint.01G0111500.1                   ------------------------------------------------------------
Tintermedium-Thint.03G0265300.1                   ------------------------------------------------------------
Hvulgare-HORVU2Hr1G038140.1                       K------------------------TTPH-------------------------------
Tintermedium-Thint.04G0171200.1                   ------------------------------------------------------------
Taestivum-Traes_2AS_958327519.3                   ------------------------------------------------------------
Taestivum-Traes_2DS_28CA50371.1                   ------------------------------------------------------------
Tintermedium-Thint.05G0160800.1                   ------------------------------------------------------------
Tintermedium-Thint.06G0299000.1                   ------------------------------------------------------------
Taestivum-Traes_2DS_3791A8A36.1                   ------------------------------------------------------------
Tintermedium-Thint.06G0342800.1                   ------------------------------------------------------------
Tintermedium-Thint.05G0150900.1                   ------------------------------------------------------------
Tintermedium-Thint.04G0242300.1                   ------------------------------------------------------------
Tintermedium-Thint.06G0298700.1                   ------------------------------------------------------------
Taestivum-Traes_2BS_88CF42F2E.1                   ------------------------------------------------------------
Tintermedium-Thint.04G0241900.1                   ------------------------------------------------------------
Platifolius-Phala.06G165300.1                     ------------------------------------------------------------
Streptochaeta-strangu_019385-RA                   ------------------------------------------------------------
Streptochaeta-strangu_028944-RA                   ------------------------------------------------------------
Acomosus-Aco010091.1                              ------------------------------------------------------------
Zmarina-Zosma03g22300.1                           ------------------------------------------------------------
Ecdeiocolea-Emoptg0003741_1G000600.1              ------------------------------------------------------------
Jascendens-Joasc.05G060500.1                      ------------------------------------------------------------
Ecdeiocolea-Emoptg0003741_1G000630.1              ------------------------------------------------------------
Jascendens-Joasc.05G060400.1                      ------------------------------------------------------------
Bdistachyon-Bradi3g49250.2                        ------------------------------------------------------------
Hvulgare-HORVU6Hr1G058820.1                       ------------------------------------------------------------
Taestivum-Traes_6DL_1AEA7B869.1                   ------------------------------------------------------------
Tintermedium-Thint.18G0192300.1                   ------------------------------------------------------------
Tintermedium-Thint.17G0230800.1                   ------------------------------------------------------------
Tintermedium-Thint.17G0230900.1                   ------------------------------------------------------------
Ecoracana-ELECO.r07.2AG0138000.1                  ------------------------------------------------------------
Ecoracana-ELECO.r07.2BG0193650.1                  ------------------------------------------------------------
Pvaginatum-Pavag04G202700.1                       ------------------------------------------------------------
Sviridis-Sevir.1G245000.1                         ------------------------------------------------------------
Sviridis-Sevir.6G187100.1                         ------------------------------------------------------------
Pvirgatum-Pavir.1NG356200.1                       ------------------------------------------------------------
Sbicolor-Sobic.004G220300.1                       ------------------------------------------------------------
Zmays-Zm00001d017274_T001                         ------------------------------------------------------------
Zmays-Zm00001d051161_T003                         ------------------------------------------------------------
Sviridis-Sevir.7G178200.1                         ------------------------------------------------------------
Osativa-LOC_Os02g41630.2                          ------------------------------------------------------------
Ecoracana-ELECO.r07.4AG0310430.1                  ------------------------------------------------------------
Ecoracana-ELECO.r07.4BG0341620.1                  ------------------------------------------------------------
Pvaginatum-Pavag06G160200.1                       ------------------------------------------------------------
Pvirgatum-Pavir.7KG238255.1                       ------------------------------------------------------------
Sbicolor-Sobic.006G148800.1                       ------------------------------------------------------------
Zmays-Zm00001d003016_T001                         M----------------------RSTTPLNSPTRSP--TNPPTPGLSASPKPPLLLLVLV
Osativa-LOC_Os04g43760.1                          ------------------------------------------------------------
Platifolius-Phala.02G210400.1                     ------------------------------------------------------------
Platifolius-Phala.06G165100.1                     ------------------------------------------------------------
Streptochaeta-strangu_019386-RA                   ------------------------------------------------------------
Streptochaeta-strangu_020769-RA                   ------------------------------------------------------------
Sviridis-Sevir.2G448300.1                         ------------------------------------------------------------
Sviridis-Sevir.7G177900.1                         ------------------------------------------------------------


Aamericanus-Acora.01G232100.1                     --------------------------------------------------------ME--
Aamericanus-Acora.10G174800.1                     --------------------------------------------------------MEHS
Acomosus-Aco013943.1                              --------------------------------------------------------M---
Aofficinalis-evm.model.AsparagusV1_01.827         ------------------------------------------------------------
Typha_latifolia-Tylat.14G083800.1                 --------------------------------------------------------MECV
Aofficinalis-evm.model.AsparagusV1_04.71          ------------------------------------------------------------
Aofficinalis-evm.model.AsparagusV1_04.93          ------------------------------------------------------------
Aofficinalis-evm.model.AsparagusV1_10.1948        ------------------------------------------------------------
Typha_latifolia-Tylat.03G127700.1                 --------------------------------------------------------MECE
Spolyrhiza-Spipo11G0025500                        --------------------------------------------------------MEAA
Spolyrhiza-Spipo1G0003500                         --------------------------------------------------------MEAA
Phoenix_dactylifera-LOC103718264                  --------------------------------------------------------MENS
Macuminata-GSMUA_Achr1T09070_001                  --------------------------------------------------------MEFA
Acomosus-Aco006987.1                              --------------------------------------------------------MEF-
Acomosus-Aco027752.1                              --------------------------------------------------------MEF-
Cscoparia-Csco27g03980.1                          --------------------------------------------------------MEF-
Cscoparia-Csco27g03990.1                          --------------------------------------------------------MEF-
Macuminata-GSMUA_Achr11T16380_001                 --------------------------------------------------------MEV-
Macuminata-GSMUA_Achr11T22840_001                 --------------------------------------------------------MEV-
Macuminata-GSMUA_Achr2T00240_001                  --------------------------------------------------------MEV-
Macuminata-GSMUA_Achr5T18560_001                  --------------------------------------------------------MEN-
Macuminata-GSMUA_Achr8T18960_001                  --------------------------------------------------------MEV-
Macuminata-GSMUA_Achr5T03950_001                  --------------------------------------------------------MEF-
Phoenix_dactylifera-LOC103703950                  --------------------------------------------------------MEF-
Acoerulea-Aqcoe1G140100.1                         --------------------------------------------------------MAAT
Acoerulea-Aqcoe2G144500.1                         --------------------------------------------------------MGS-
Typha_latifolia-Tylat.05G053500.1                 --------------------------------------------------------MEH-
Zmarina-Zosma05g29470.1                           --------------------------------------------------------MDAM
Acomosus-Aco007727.1                              --------------------------------------------------------MEC-
Bvulgarisssp-Bevul.7G054300.1                     --------------------------------------------------------MEA-
Bdistachyon-Bradi3g47110.1                        --------------------------------------------------------MAC-
Bdistachyon-Bradi3g47120.1                        ------------------------------------------------------MTMAS-
Bdistachyon-Bradi3g48840.1                        --------------------------------------------------------MEC-
Bdistachyon-Bradi3g49280.1                        --------------------------------------------------------MEC-
Hvulgare-HORVU6Hr1G058840.1                       --------------------------------------------------------MEC-
Taestivum-Traes_6BL_C96F626D9.1                   ------------------------------------------------------------
Tintermedium-Thint.18G0192700.1                   --------------------------------------------------------MEC-
Taestivum-Traes_6DL_04CD7E82C.1                   --------------------------------------------------------MEY-
Bdistachyon-Bradi3g49270.1                        --------------------------------------------------------MAR-
Ecoracana-ELECO.r07.2AG0138020.1                  --------------------------------------------------------MEC-
Ecoracana-ELECO.r07.2BG0193670.1                  --------------------------------------------------------MEC-
Osativa-LOC_Os02g41670.1                          --------------------------------------------------------MAC-
Osativa-LOC_Os02g41680.1                          --------------------------------------------------------MEC-
Pvaginatum-Pavag04G202900.1                       --------------------------------------------------------MAC-
Pvaginatum-PavagK321000.1                         --------------------------------------------------------MAC-
Pvaginatum-PavagK321300.1                         --------------------------------------------------------MAC-
Pvaginatum-PavagK321100.1                         ---LPSLSATRALLCSSNPSKLA--TIHLHTFPYLLPSSASSVIHQFS-----ALAMAC-
Pvaginatum-PavagK321200.1                         --------------------------------------------------------MEC-
Pvirgatum-Pavir.1KG386500.1                       --------------------------------------------------------MEC-
Pvirgatum-Pavir.1NG356700.1                       --------------------------------------------------------MAC-
Sviridis-Sevir.1G245166.1                         --------------------------------------------------------MAC-
Sviridis-Sevir.1G245232.1                         --------------------------------------------------------MAC-
Pvirgatum-Pavir.1NG356800.1                       --------------------------------------------------------MEC-
Sbicolor-Sobic.004G220500.1                       --------------------------------------------------------MAC-
Sbicolor-Sobic.004G220600.2                       --------------------------------------------------------MAC-
Sbicolor-Sobic.004G220700.1                       --------------------------------------------------------MAC-
Sviridis-Sevir.1G245300.2                         --------------------------------------------------------MEC-
Zmays-Zm00001d017276_T001                         --------------------------------------------------------MAC-
Zmays-Zm00001d017279_T001                         --------------------------------------------------------MEC-
Zmays-Zm00001d051166_T001                         --------------------------------------------------------MEC-
```

```
Ecoracana-ELECO.r07.2AG0138030.1 --------------------------------------------------------MEC-
Ecoracana-ELECO.r07.2BG0193680.1 --------------------------------------------------------MEC-
Osativa-LOC_Os05g35290.1 --------------------------------------------------------MEC-
Platifolius-Phala.02G210700.1 --------------------------------------------------------MEC-
Ecoracana-ELECO.r07.4AG0310420.1 --------------------------------------------------------MEC-
Ecoracana-ELECO.r07.4BG0341610.1 --------------------------------------------------------MEC-
Bdistachyon-Bradi3g49260.1 --------------------------------------------------------MEC-
Hvulgare-HORVU0Hr1G016330.1 --------------------------------------------------------MEC-
Tintermedium-Thint.17G0260200.1 --------------------------------------------------------MEC-
Tintermedium-Thint.18G0192600.1 --------------------------------------------------------MEC-
Tintermedium-Thint.17G0231000.1 --------------------------------------------------------MEC-
Ecoracana-ELECO.r07.2AG0138010.1 --------------------------------------------------------MEC-
Ecoracana-ELECO.r07.2BG0193660.1 --------------------------------------------------------MEC-
Osativa-LOC_Os02g41650.3 --------------------------------------------------------MEC-
Pvaginatum-Pavag04G202800.1 --------------------------------------------------------MEC-
Pvaginatum-PavagK320900.1 --------------------------------------------------------MEC-
Pvirgatum-Pavir.1NG356400.1 --------------------------------------------------------MEC-
Sviridis-Sevir.1G245100.1 --------------------------------------------------------MEC-
Sbicolor-Sobic.004G220400.1 --------------------------------------------------------MEC-
Zmays-Zm00001d017275_T001 --------------------------------------------------------MEC-
Zmays-Zm00001d051163_T002 --------------------------------------------------------MES-
Pvirgatum-Pavir.1KG386300.1 --------------------------------------------------------MEC-
Platifolius-Phala.02G210600.1 --------------------------------------------------------MEC-
Streptochaeta-strangu_020767-RA --------------------------------------------------------MEC-
Bdistachyon-Bradi5g15830.1 --------------------------------------------------------MEY-
Hvulgare-HORVU2Hr1G089440.4 --------------------------------------------------------MEC-
Taestivum-Traes_2AL_0DDC60D6E.1 ------------------------------------------------------------
Tintermedium-Thint.06G0475900.1 --------------------------------------------------------MEY-
Tintermedium-Thint.06G0476200.1 --------------------------------------------------------MEC-
Taestivum-Traes_2AL_2763280FC.1 ------------------------------------------------------------
Taestivum-Traes_2DL_8C21AAAEF.1 ------------------------------------------------------------
Tintermedium-Thint.V0383500.1 --------------------------------------------------------MEC-
Taestivum-Traes_2AL_9EC3226F7.1 --------------------------------------------------------MEC-
Tintermedium-Thint.05G0273800.1 --------------------------------------------------------MEC-
Tintermedium-Thint.V0383400.1 --------------------------------------------------------MEC-
Taestivum-Traes_2BL_13D5272D7.1 --------------------------------------------------------MEC-
Tintermedium-Thint.06G0475700.1 --------------------------------------------------------MEC-
Taestivum-Traes_2BL_C051606EA.2 --------------------------------------------------------MEC-
Tintermedium-Thint.V0383600.1 --------------------------------------------------------MEC-
Taestivum-Traes_2AL_9D78F85E2.1 --------------------------------------------------------MEC-
Tintermedium-Thint.05G0274100.1 --------------------------------------------------------MEC-
Tintermedium-Thint.06G0476300.1 --------------------------------------------------------MEC-
Taestivum-Traes_2BL_5A50FDA1A.1 ------------------------------------------------------------
Tintermedium-Thint.05G0273900.1 ------------------------------------------------------------
Taestivum-Traes_2DL_30F8555D4.1 ------------------------------------------------------------
Taestivum-Traes_2DL_83168C1E0.1 ------------------------------------------------------------
Taestivum-Traes_2DL_B978B4E22.1 ------------------------------------------------------------
Taestivum-Traes_1AS_0B9295A68.1 --------------------------------------------------------MEC-
Taestivum-Traes_1BS_7068F56E6.1 --------------------------------------------------------MEC-
Taestivum-Traes_1BS_AEA817616.1 ------------------------------------------------------------
Taestivum-Traes_4AL_892C47ED5.1 --------------------------------------------------------MEC-
Tintermedium-Thint.03G0055900.1 --------------------------------------------------------MEC-
Tintermedium-Thint.01G0012900.1 --------------------------------------------------------MEC-
Tintermedium-Thint.02G0158500.1 --------------------------------------------------------MEC-
Tintermedium-Thint.03G0055700.1 --------------------------------------------------------MEC-
Taestivum-Traes_1BS_C3AA04888.1 ------------------------------------------------------------
Taestivum-Traes_1DS_A8BD91E4A.1 --------------------------------------------------------MEY-
Taestivum-Traes_1BS_C26CCE0E9.1 --------------------------------------------------------MEF-
Taestivum-Traes_1DS_A21FCBA3D.1 --------------------------------------------------------MEF-
Tintermedium-Thint.01G0012200.1 --------------------------------------------------------MEF-
Tintermedium-Thint.02G0158600.1 --------------------------------------------------------MEF-
Tintermedium-Thint.01G0012700.1 --------------------------------------------------------MEF-
Taestivum-Traes_5BL_F50A47503.1 --------------------------------------------------------MEC-
Tintermedium-Thint.02G0158700.1 --------------------------------------------------------MDC-
Taestivum-Traes_5BL_AB528184A.1 --------------------------------------------------------MEF-
Tintermedium-Thint.03G0055800.1 --------------------------------------------------------MEF-
Taestivum-Traes_1AS_F9013A945.1 --------------------------------------------------------MEC-
Taestivum-Traes_1DS_43DAC7A7F.1 --------------------------------------------------------MEC-
Taestivum-Traes_1BS_BD86C90A7.1 --------------------------------------------------------MEC-
Tintermedium-Thint.02G0158900.1 --------------------------------------------------------MEC-
Tintermedium-Thint.05G0274000.1 ------------------------------------------------------------
Osativa-LOC_Os04g43800.1 --------------------------------------------------------MEC-
Pvaginatum-Pavag06G160300.1 --------------------------------------------------------MEC-
Sbicolor-Sobic.006G148900.1 --------------------------------------------------------MEC-
Pvirgatum-Pavir.7KG237800.1 --------------------------------------------------------MEC-
Pvirgatum-Pavir.7NG355800.1 --------------------------------------------------------MEC-
Sviridis-Sevir.7G178300.1 --------------------------------------------------------MEC-
Zmays-Zm00001d003015_T001 --------------------------------------------------------MEC-
Hvulgare-HORVU1Hr1G022060.1 ------------------------------------------------------------
Taestivum-Traes_1BS_723922D171.2 ------------------------------------------------------------
Taestivum-Traes_1DS_A171C7D59.1 ------------------------------------------------------------
Tintermedium-Thint.01G0111500.1 ------------------------------------------------------------
Tintermedium-Thint.03G0265300.1 ------------------------------------------------------------
Hvulgare-HORVU2Hr1G038140.1 --------------------------------------------------------LPCC
Tintermedium-Thint.04G0171200.1 ------------------------------------------------------------
Taestivum-Traes_2AS_958327519.3 ------------------------------------------------------------
Taestivum-Traes_2DS_28CA50371.1 ------------------------------------------------------------
Tintermedium-Thint.05G0160800.1 ------------------------------------------------------------
Tintermedium-Thint.06G0299000.1 ------------------------------------------------------------
Taestivum-Traes_2DS_3791A8A36.1 ------------------------------------------------------------
Tintermedium-Thint.06G0342800.1 ------------------------------------------------------------
Tintermedium-Thint.05G0150900.1 ------------------------------------------------------------
Tintermedium-Thint.04G0242300.1 ------------------------------------------------------------
Tintermedium-Thint.06G0298700.1 ------------------------------------------------------------
Taestivum-Traes_2BS_88CF42F2E.1 ------------------------------------------------------------
Tintermedium-Thint.04G0241900.1 ------------------------------------------------------------
Platifolius-Phala.06G165300.1 --------------------------------------------------------MDS-
Streptochaeta-strangu_019385-RA --------------------------------------------------------MEF-
Streptochaeta-strangu_028944-RA --------------------------------------------------------MQR-
Acomosus-Aco010091.1 --------------------------------------------------------MER-
Zmarina-Zosma03g22300.1 --------------------------------------------------------M---
Ecdeiocolea-Emoptg0003741_1G000600.1 --------------------------------------------------------MEC-
Jascendens-Joasc.05G060500.1 --------------------------------------------------------MEC-
Ecdeiocolea-Emoptg0003741_1G000630.1 --------------------------------------------------------MAC-
Jascendens-Joasc.05G060400.1 --------------------------------------------------------MA--
Bdistachyon-Bradi3g49250.2 --------------------------------------------------------MAG-
Hvulgare-HORVU6Hr1G058820.1 --------------------------------------------------------MAG-
Taestivum-Traes_6DL_1AEA7B869.1 --------------------------------------------------------MAG-
Tintermedium-Thint.18G0192300.1 --------------------------------------------------------MAG-
Tintermedium-Thint.17G0230800.1 --------------------------------------------------------MAG-
Tintermedium-Thint.17G0230900.1 --------------------------------------------------------MAG-
Ecoracana-ELECO.r07.2AG0138000.1 --------------------------------------------------------MAG-
Ecoracana-ELECO.r07.2BG0193650.1 --------------------------------------------------------MAG-
Pvaginatum-Pavag04G202700.1 --------------------------------------------------------MAG-
Sviridis-Sevir.1G245000.1 --------------------------------------------------------MAG-
Sviridis-Sevir.6G187100.1 --------------------------------------------------------MAG-
Pvirgatum-Pavir.1NG356200.1 --------------------------------------------------------MAG-
Sbicolor-Sobic.004G220300.1 --------------------------------------------------------MAG-
Zmays-Zm00001d017274_T001 --------------------------------------------------------MAG-
Zmays-Zm00001d051161_T003 --------------------------------------------------------MAG-
Sviridis-Sevir.7G178200.1 --------------------------------------------------------MAS-
Osativa-LOC_Os02g41630.2 --------------------------------------------------------MAG-
Ecoracana-ELECO.r07.4AG0310430.1 --------------------------------------------------------MAS-
Ecoracana-ELECO.r07.4BG0341620.1 --------------------------------------------------------MAS-
Pvaginatum-Pavag06G160200.1 --------------------------------------------------------MAS-
Pvirgatum-Pavir.7KG238255.1 --------------------------------------------------------MAS-
Sbicolor-Sobic.006G148800.1 --------------------------------------------------------MAS-
Zmays-Zm00001d003016_T001 LVLLPYITASPPL---HFPSRRAPKSIHHHS---CLPSSSSATPHQHHHTHPLPAQMAS-
Osativa-LOC_Os04g43760.1 --------------------------------------------------------MAS-
Platifolius-Phala.02G210400.1 --------------------------------------------------------MAS-
```

```
Platifolius-Phala.06G165100.1              --------------------------------------------------------MAA-
Streptochaeta-strangu_019386-RA            --------------------------------------------------------MVA-
Streptochaeta-strangu_020769-RA            --------------------------------------------------------MAS-
Sviridis-Sevir.2G448300.1                  --------------------------------------------------------MAC-
Sviridis-Sevir.7G177900.1                  --------------------------------------------------------MAC-
```

```
Aamericanus-Acora.01G232100.1              ----------------------------------------------------------NG
Aamericanus-Acora.10G174800.1              ----------LN---------------GF--------------T--------NGSVNQNG
Acomosus-Aco013943.1                       ---------------------------GG--------------G----------------
Aofficinalis-evm.model.AsparagusV1_01.827  ------------------------------------------------------------
Typha_latifolia-Tylat.14G083800.1          PQIH------EN---------------GK--------------S--------------DG
Aofficinalis-evm.model.AsparagusV1_04.71   ------------------------------------------------------------
Aofficinalis-evm.model.AsparagusV1_04.93   ------------------------------------------------------------
Aofficinalis-evm.model.AsparagusV1_10.1948 ------------------------------------------------------------
Typha_latifolia-Tylat.03G127700.1          PTAH------VN---------------GS--------------A--------NGSV--AD
Spolyrhiza-Spipo11G0025500                 TQVH------CN---------------GN------------------------GNVKLEA
Spolyrhiza-Spipo1G0003500                  GQIN------SN---------------GH--------------S---------GVL--EG
Phoenix_dactylifera-LOC103718264           ---------------------------GA--------------K---------------S
Macuminata-GSMUA_Achr1T09070_001           PKAQV----VEN---------------GE--------------A----------------
Acomosus-Aco006987.1                       ---------------------------------------------------VEGY-----
Acomosus-Aco027752.1                       ---------------------------------------------------VEGY-----
Cscoparia-Csco27g03980.1                   ---------HLN---------------GN--------------T--------NGSM--TD
Cscoparia-Csco27g03990.1                   ---------HLN---------------GN--------------T--------NGSM--TD
Macuminata-GSMUA_Achr11T16380_001          ----------EN---------------GA--------------A--------------GG
Macuminata-GSMUA_Achr11T22840_001          ----------EN---------------GV--------------H-------GNGFA--DG
Macuminata-GSMUA_Achr2T00240_001           ----------EN---------------GA--------------Y-------TNSTT--DG
Macuminata-GSMUA_Achr5T18560_001           --------------------------GGV--------------H-------ANG-----S
Macuminata-GSMUA_Achr8T18960_001           ----------EN---------------GV--------------G--------------NG
Macuminata-GSMUA_Achr5T03950_001           ----------ERN--------------GH--------------H-------ANG-----G
Phoenix_dactylifera-LOC103703950           --------AHEN---------------GN--------------A--------NGAV--NG
Acoerulea-Aqcoe1G140100.1                  KI-------NEN---------------GN--------------S--------------NS
Acoerulea-Aqcoe2G144500.1                  ----------QG---------------QE--------------D--------------NA
Typha_latifolia-Tylat.05G053500.1          ------------------------------------------------------------
Zmarina-Zosma05g29470.1                    DLCQV---KLSN---------------GA--------------T--------NGI-----
Acomosus-Aco007727.1                       ---------DAN---------------GV--------------V----------------
Bvulgarisssp-Bevul.7G054300.1              ------NQQYQN---------------GS--------------SE-------------MD
Bdistachyon-Bradi3g47110.1                 ----------EN---------------GQ--------------V-----AANG-------
Bdistachyon-Bradi3g47120.1                 ----------KN---------------VH--------------V-----SADGY----LI
Bdistachyon-Bradi3g48840.1                 ----------EN---------------GQ--------------------FAANG----TG
Bdistachyon-Bradi3g49280.1                 ----------EN---------------GQ--------------------FAANG----TG
Hvulgare-HORVU6Hr1G058840.1                ----------EN---------------GE--------------------AAANG----NI
Taestivum-Traes_6BL_C96F626D9.1            ------------------------------------------------------------
Tintermedium-Thint.18G0192700.1            ----------EN---------------GE--------------------VAANG----NS
Taestivum-Traes_6DL_04CD7E82C.1            ----------EN---------------GE--------------------VAANG----NS
Bdistachyon-Bradi3g49270.1                 ----------EN---------------AR--------------V-----AAANG------
Ecoracana-ELECO.r07.2AG0138020.1           ----------ER---------------AN--------------V-----AAPSG----DA
Ecoracana-ELECO.r07.2BG0193670.1           ----------ER---------------AN--------------V-----AAPSG----DA
Osativa-LOC_Os02g41670.1                   ----------EN---------------GQ--------------V-----AADGI----NG
Osativa-LOC_Os02g41680.1                   ----------EN---------------GR--------------V-----SANGM----SG
Pvaginatum-Pavag04G202900.1                ----------------------------------------------------------DS
Pvaginatum-PavagK321000.1                  ----------------------------------------------------------DS
Pvaginatum-PavagK321300.1                  ----------------------------------------------------------DS
Pvaginatum-PavagK321100.1                  ----------------------------------------------------------DS
Pvaginatum-PavagK321200.1                  ----------EN---------------GH--------------A-----ASANA----GS
Pvirgatum-Pavir.1KG386500.1                ----------KN---------------GR--------------V-------ANG----DS
Pvirgatum-Pavir.1NG356700.1                ----------EK---------------SN--------------V------ATNG----DG
Sviridis-Sevir.1G245166.1                  ----------EK---------------SN--------------V------AANG----DG
Sviridis-Sevir.1G245232.1                  ----------EK---------------SN--------------V------AANG----DG
Pvirgatum-Pavir.1NG356800.1                ----------EN---------------GR---------------------GANG----DT
Sbicolor-Sobic.004G220500.1                ----------EN---------------GR--------------V-----AATNG----DG
Sbicolor-Sobic.004G220600.2                ----------EN---------------GR--------------V-----AATNG----DG
Sbicolor-Sobic.004G220700.1                ----------DN---------------GR--------------V-----AATNG----DG
Sviridis-Sevir.1G245300.2                  ----------EN---------------GR--------------V-----AAANG----DG
Zmays-Zm00001d017276_T001                  ----------------------------------------------------------DS
Zmays-Zm00001d017279_T001                  ----------EN---------------GRG-------------V-----AATNS----DS
Zmays-Zm00001d051166_T001                  ----------DN---------------GR--------------V-----AATNG----DS
Ecoracana-ELECO.r07.2AG0138030.1           ----------EN---------------GH--------------V-----AATNNG---VD
Ecoracana-ELECO.r07.2BG0193680.1           ----------EN---------------GH--------------V-----AATNNG---VD
Osativa-LOC_Os05g35290.1                   ----------ET---------------GY--------------V-----AAAAD----GG
Platifolius-Phala.02G210700.1              ----------EK---------------SQ--------------F-----VAANG----ND
Ecoracana-ELECO.r07.4AG0310420.1           ----------ET---------------GH--------------V-----AAANG----NG
Ecoracana-ELECO.r07.4BG0341610.1           ----------EN---------------GH--------------A-----ATANG----NG
Bdistachyon-Bradi3g49260.1                 ----------EN---------------GL--------------V-----GSLNG----EG
Hvulgare-HORVU0Hr1G016330.1                ----------ET---------------GL--------------V-----GSLNG----DG
Tintermedium-Thint.17G0260200.1            ----------ET---------------GL--------------V-----GSLNG----DG
Tintermedium-Thint.18G0192600.1            ----------ET---------------GL--------------V-----GSLNG----EG
Tintermedium-Thint.17G0231000.1            ----------EN---------------GD--------------V------AANG----NS
Ecoracana-ELECO.r07.2AG0138010.1           ----------ET---------------GV--------------V-----RSLHG----EG
Ecoracana-ELECO.r07.2BG0193660.1           ----------ET---------------GL--------------V-----RSLHG----EG
Osativa-LOC_Os02g41650.3                   ----------ET---------------GL--------------V-----RSLNG----DG
Pvaginatum-Pavag04G202800.1                ----------ET---------------GL--------------V-----RSLHG----DG
Pvaginatum-PavagK320900.1                  ----------ET---------------GL--------------V-----RSLHG----DG
Pvirgatum-Pavir.1NG356400.1                ----------ET---------------GL--------------V-----RSLHG----DG
Sviridis-Sevir.1G245100.1                  ----------ET---------------GL--------------V-----RSLHG----DG
Sbicolor-Sobic.004G220400.1                ----------ET---------------GL--------------V-----RSLNG----DG
Zmays-Zm00001d017275_T001                  ----------ET---------------GL--------------V-----RSLNG----EG
Zmays-Zm00001d051163_T002                  ----------EA---------------GLL-------------VR----SSLNG----EG
Pvirgatum-Pavir.1KG386300.1                ----------ET---------------GL--------------V-----RSLHG----DG
Platifolius-Phala.02G210600.1              ----------EN---------------GQ--------------F-----VGTGG----NG
Streptochaeta-strangu_020767-RA            ----------EN---------------SQ--------------V------AANG----NC
Bdistachyon-Bradi5g15830.1                 ----------EN---------------GH--------------A------ATYG----DG
Hvulgare-HORVU2Hr1G089440.4                ----------DN---------------GH--------------V------AANG----DG
Taestivum-Traes_2AL_0DDC60D6E.1            ------------------------------------------------------------
Tintermedium-Thint.U6G0475900.1            ----------EN---------------AH--------------V------AANG----DG
Tintermedium-Thint.06G0476200.1            ----------EN---------------AH--------------V------AANG----DG
Taestivum-Traes_2AL_2763280FC.1            ------------------------------------------------------------
Taestivum-Traes_2DL_8C21AAAEF.1            ------------------------------------------------------------
Tintermedium-Thint.V0383500.1              ----------EN---------------AH--------------V------AANG----DG
Taestivum-Traes_2AL_9EC3226F7.1            ----------EN---------------AH--------------V------AANG----DG
Tintermedium-Thint.U5G0273800.1            ----------EN---------------GQ--------------I------AANG----DG
Tintermedium-Thint.V0383400.1              ----------EN---------------GQ--------------V------AANG----DG
Taestivum-Traes_2BL_13D5272D7.1            ----------EN---------------AH--------------I------AANG----DG
Tintermedium-Thint.U6G0475700.1            ----------EN---------------AH--------------I------GANG----DG
Taestivum-Traes_2BL_C051606EA.2            ----------EN---------------AH--------------V------AANG----DG
Tintermedium-Thint.V0383600.1              ----------EN---------------GH--------------V------AANG----DG
Taestivum-Traes_2AL_9D78F85E2.1            ----------EN---------------GH--------------V------AANG----DG
Tintermedium-Thint.U5G0274100.1            ----------EN---------------GH--------------V------AANG----DG
Tintermedium-Thint.06G0476300.1            ----------EN---------------GR--------------V------AANG----DG
Taestivum-Traes_2BL_5A50FDA1A.1            ------------------------------------------------------------
Tintermedium-Thint.U5G0273900.1            ------------------------------------------------------------
Taestivum-Traes_2DL_30F8555D4.1            ------------------------------------------------------------
Taestivum-Traes_2DL_83168C1E0.1            ------------------------------------------------------------
Taestivum-Traes_2DL_B978B4E22.1            ------------------------------------------------------------
Taestivum-Traes_1AS_0B9295A68.1            ----------EN---------------GQ--------------V------AGNG----NS
Taestivum-Traes_1BS_7068F56E6.1            ----------EN---------------GK--------------V------AGNG----NS
Taestivum-Traes_1BS_AEA817616.1            ------------------------------------------------------------
Taestivum-Traes_4AL_892C47ED5.1            ----------EN---------------GQ--------------V------AGNG----NS
Tintermedium-Thint.U3G0055900.1            ----------EN---------------GQ--------------V------AGNG----NS
Tintermedium-Thint.01G0012900.1            ----------ENR--------------AH--------------V------AANG----DG
Tintermedium-Thint.02G0158500.1            ----------ENR--------------GH--------------V------AAND----DG
Tintermedium-Thint.03G0055700.1            ----------ENR--------------GH--------------V------AANG----DG
Taestivum-Traes_1BS_C3AA04888.1            ------------------------------------------------------------
Taestivum-Traes_1DS_A8BD91E4A.1            ----------ENR--------------GH--------------V------AAND----DG
Taestivum-Traes_1BS_C26CCE0E9.1            ----------E-----------------------------------------NA----NG
Taestivum-Traes_1DS_A21FCBA3D.1            ----------E-----------------------------------------NA----NG
Tintermedium-Thint.U1G0012200.1            ----------E-----------------------------------------NS----NS
Tintermedium-Thint.02G0158600.1            ----------E-----------------------------------------NT----NS
```

```
Tintermedium-Thint.01G0012700.1              ----------E-----------------------------------------NA----NG
Taestivum-Traes_5BL_F50A47503.1              ----------EN---------------GQ--------------V------AGNG----NG
Tintermedium-Thint.02G0158700.1              ----------DN---------------GQ--------------V------AGNG----NS
Taestivum-Traes_5BL_AB528184A.1              ----------Q-----------------------------------------NA----NG
Tintermedium-Thint.03G0055800.1              ----------E-----------------------------------------NA----NG
Taestivum-Traes_1AS_F9013A945.1              ----------DNR--------------AH--------------V------AANG----DG
Taestivum-Traes_1DS_43DAC7A7F.1              ----------ENR--------------AH--------------V------AAND----DG
Taestivum-Traes_1BS_BD86C90A7.1              ----------ENR--------------AH--------------V------AANG----DG
Tintermedium-Thint.02G0158900.1              ----------ENR--------------GH--------------V------AANG----DG
Tintermedium-Thint.05G0274000.1              ------------------------------------------------------------
Osativa-LOC_Os04g43800.1                     ----------EN---------------GH--------------VA----AAANG----SS
Pvaginatum-Pavag06G160300.1                  ----------EN---------------GH--------------V------AASG----NG
Sbicolor-Sobic.006G148900.1                  ----------EN---------------GH--------------A-----AGASG----NG
Pvirgatum-Pavir.7KG237800.1                  ----------EN---------------GR--------------V-----AAATG----NG
Pvirgatum-Pavir.7NG355800.1                  ----------EN---------------GH--------------V------AANG----NG
Sviridis-Sevir.7G178300.1                    ----------EN---------------GH--------------V-----VAANG----NG
Zmays-Zm00001d003015_T001                    ----------EN---------------GH--------------------LAASGN---GG
Hvulgare-HORVU1Hr1G022060.1                  -------------------------------------------------MAANGN---DG
Taestivum-Traes_1BS_723922D171.2             -------------------------------------------------MAANGN---DG
Taestivum-Traes_1DS_A171C7D59.1              -------------------------------------------------MAANGN---DG
Tintermedium-Thint.01G0111500.1              -------------------------------------------------MAANGN---DD
Tintermedium-Thint.03G0265300.1              -------------------------------------------------MAANGN---DG
Hvulgare-HORVU2Hr1G038140.1                  IQLFSLTAAPQNRHIDNQPLVSTRALLSHKARTAPSRLQDPLLIK---PMAANGN---DG
Tintermedium-Thint.04G0171200.1              ------------------------------------------------------------
Taestivum-Traes_2AS_958327519.3              ------------------------------------------------------------
Taestivum-Traes_2DS_28CA50371.1              ------------------------------------------------------------
Tintermedium-Thint.05G0160800.1              -------------------------------------------------MAANGN---DG
Tintermedium-Thint.06G0299000.1              -------------------------------------------------MAANGN---DG
Taestivum-Traes_2DS_3791A8A36.1              ------------------------------------------------------------
Tintermedium-Thint.06G0342800.1              ------------------------------------------------------------
Tintermedium-Thint.05G0150900.1              ------------------------------------------------------------
Tintermedium-Thint.04G0242300.1              -------------------------------------------------MATNGN---DG
Tintermedium-Thint.06G0298700.1              -------------------------------------------------MATNGS---DG
Taestivum-Traes_2BS_88CF42F2E.1              ------------------------------------------------------------
Tintermedium-Thint.04G0241900.1              -------------------------------------------------MAANGN---DG
Platifolius-Phala.06G165300.1                ----------EK---------------GH--------------V------AANG----SD
Streptochaeta-strangu_019385-RA              -------------------------------------------LS--C-VAANG----NG
Streptochaeta-strangu_028944-RA              ----------ES---------------GR--------------VEAYC----------NG
Acomosus-Aco010091.1                         --------------------------------------------------APSSTK--KD
Zmarina-Zosma03g22300.1                      ---------------------------G--------------------------------
Ecdeiocolea-Emoptg0003741_1G000600.1         ----------IN---------------GN-------------------GAAANG----SG
Jascendens-Joasc.05G060500.1                 ----------EN---------------GN--------------VA--AVAAVNGG---NG
Ecdeiocolea-Emoptg0003741_1G000630.1         --------------------------------------------------LSSD----NV
Jascendens-Joasc.05G060400.1                 --------------------------------------------------FQND----NV
Bdistachyon-Bradi3g49250.2                   ----------NG-------------------------------A----------------
Hvulgare-HORVU6Hr1G058820.1                  ----------NG-------------------------------A----------------
Taestivum-Traes_6DL_1AEA7B869.1              ----------NG-------------------------------A----------------
Tintermedium-Thint.18G0192300.1              ----------NG-------------------------------A----------------
Tintermedium-Thint.17G0230800.1              ----------NG-------------------------------A----------------
Tintermedium-Thint.17G0230900.1              ----------NG-------------------------------A----------------
Ecoracana-ELECO.r07.2AG0138000.1             ----------NG-------------------------------M----------------
Ecoracana-ELECO.r07.2BG0193650.1             ----------NG-------------------------------M----------------
Pvaginatum-Pavag04G202700.1                  ----------NG-------------------------------L----------------
Sviridis-Sevir.1G245000.1                    ----------NG-------------------------------L----------------
Sviridis-Sevir.6G187100.1                    ----------NG-------------------------------L----------------
Pvirgatum-Pavir.1NG356200.1                  ----------NG-------------------------------L----------------
Sbicolor-Sobic.004G220300.1                  ----------NG-------------------------------A----------------
Zmays-Zm00001d017274_T001                    ----------NG-------------------------------A----------------
Zmays-Zm00001d051161_T003                    ----------NG-------------------------------A----------------
Sviridis-Sevir.7G178200.1                    ----------NT-------------------------------A----------------
Osativa-LOC_Os02g41630.2                     ----------NG-------------------------------P----------------
Ecoracana-ELECO.r07.4AG0310430.1             ----------NT-------------------------------A----------------
Ecoracana-ELECO.r07.4BG0341620.1             ----------NT-------------------------------A----------------
Pvaginatum-Pavag06G160200.1                  ----------NT-------------------------------A----------------
Pvirgatum-Pavir.7KG238255.1                  ----------NT-------------------------------A----------------
Sbicolor-Sobic.006G148800.1                  ----------NT-------------------------------A----------------
Zmays-Zm00001d003016_T001                    ----------NT-------------------------------A----------------
Osativa-LOC_Os04g43760.1                     ----------QT-------------------------------A--------------DA
Platifolius-Phala.02G210400.1                ----------QA-------------------------------D--------------TV
Platifolius-Phala.06G165100.1                ----------QN-------------------------------T--------------N-
Streptochaeta-strangu_019386-RA              ----------QS-------------------------------D--------------SA
Streptochaeta-strangu_020769-RA              ----------Q-------------------------------------------------
Sviridis-Sevir.2G448300.1                    ----------NT-------------------------------A----------------
Sviridis-Sevir.7G177900.1                    ----------NT-------------------------------A----------------
```

```
Aamericanus-Acora.01G232100.1                FCMKKEEEV---------VDPLNWAKSAEPL--MGSHLEEVKRMVEEFRNPLVCLEGADL
Aamericanus-Acora.10G174800.1                LCLKKEAAS---------TDPLNWAKSTEPL--MGSHLEEVKKMVEEFRNPLVSLEGADL
Acomosus-Aco013943.1                         ------------------SRP----RAA-----RGRQADGIA------------------
Aofficinalis-evm.model.AsparagusV1_01.827    ------------------------------------------------------------
Typha_latifolia-Tylat.14G083800.1            LCLKN--------------DPLNWVMAAESM--AGSHLDEVKAMVEEFRKPSVKLEGATL
Aofficinalis-evm.model.AsparagusV1_04.71     ------------------------------------------------------------
Aofficinalis-evm.model.AsparagusV1_04.93     ------------------------------------------------------------
Aofficinalis-evm.model.AsparagusV1_10.1948   ------------------------------------------------------------
Typha_latifolia-Tylat.03G127700.1            FCVK---------------DPLNWGKAAEAV--SGSHLDEVKRMVEDYRSPVVRLEGATL
Spolyrhiza-Spipo11G0025500                   FCLKPAAAAAAPAVAPVSGDPLNWGAAADSL--RGSHLDEVKRMVEDFRRPLVRLEGADL
Spolyrhiza-Spipo1G0003500                    FCLKPTIAA---------GDPLNWGAAAESL--SGSHLEEVKRMVEEFRQPLVQLEGADL
Phoenix_dactylifera-LOC103718264             LCL---------------SDPLHWAKAAETL--TGSHLDEVKHMVEEYRKPLVRLEGATL
Macuminata-GSMUA_Achr1T09070_001             FCLK--------------ADPLNWIKAAESL--TGSHLDEVKRMVEEFRKPQVRLEGATL
Acomosus-Aco006987.1                         LCADEKKQR----------DPLNWGRAAEAM--SGSHLEEVKRMVEEFRRPLVRLEGADL
Acomosus-Aco027752.1                         LCADEKKQR----------DPLNWGRAAEAM--SGSHLEEVKRMVEEFRRPLVRLEGADL
Cscoparia-Csco27g03980.1                     LCMSK-------------KDALNWGKAAEAL--SGSHLDEVKRMVEEFRAPLVKLEGADL
Cscoparia-Csco27g03990.1                     LCISK-------------KDALNWGKAAEAL--SGSHLDEVKRMVEEFRAPLVKLEGADL
Macuminata-GSMUA_Achr11T16380_001            LRVE--------------TDPLNWAAAAEAL--AGSHLDEVRRMTWEFRQPLVKLEGSTL
Macuminata-GSMUA_Achr11T22840_001            LCVK--------------ADPLNWGAAAEAL--TGSHLDEVKRMVEEFRRPLVRLEGATL
Macuminata-GSMUA_Achr2T00240_001             LCIK--------------ADALNWATAAEAL--SGNHLDEVRRMVEESRRPLVRLEGATL
Macuminata-GSMUA_Achr5T18560_001             LCIR--------------TDPLNWGAAAEAV--TGSHLEEVKRMVGEFRQPLVRLEGATL
Macuminata-GSMUA_Achr8T18960_001             PCIK--------------ADPLNWGAAAEAM--TRSHLDDVKRMVEEFRQPLVRLEGATL
Macuminata-GSMUA_Achr5T03950_001             LCIK---------------DPLNWAAAAEAL--TGSHLDEVKKMVAESRQPLVSLEGTTL
Phoenix_dactylifera-LOC103703950             LCIK---------------DPLNWAAAAEAL--TGSHLDEVKRMVEEFRQPAVRLEGATL
Acoerulea-Aqcoe1G140100.1                    LCINK-------------IDPLNWGLAAESL--KGSHLDEVKRMVEEFRKPGLKLGGATL
Acoerulea-Aqcoe2G144500.1                    FCVK---------------DPLNWGVAAESL--KGSHLDEVKRMVEEFRKPVVKLGGESL
Typha_latifolia-Tylat.05G053500.1            -CAPQ----------VHDKDPLNWAKAAESM--AGSHLDEVRLMVEEFKRPLVMLEGASL
Zmarina-Zosma05g29470.1                      -----------------TDADPLNWGAAAEEL--TGSHLQEVKRMVEDFNNPLVKIEGATL
Acomosus-Aco007727.1                         ------------------TDPLGWAAAAAAAAESGSHLEEVKRMVEEFRAPAVTLKGVEL
Bvulgarisssp-Bevul.7G054300.1                LCIGGK---PI------QADPLNWGKAAEAI--TGSHLDEVKKMVAEYRNPAIRLGGETL
Bdistachyon-Bradi3g47110.1                   ICTAI----QH-------ADPLNWGKAAEAL--TGSHLEEVKRMVAEYRQPVVTIEGASL
Bdistachyon-Bradi3g47120.1                   LCPATS---QH-------ADPLNWGKAAEAL--TGSHLEEVKRMVAEYRQPVVTIEGASL
Bdistachyon-Bradi3g48840.1                   LCMAT----PS-------ADPLNWGKAAEEL--TGSHLDEVKRMVAEYRKPVVTIEGATL
Bdistachyon-Bradi3g49280.1                   LCMAT----PR-------ADPLNWGKAAEEL--TGSHLDEVKRMVAEYRKPVVTIEGATL
Hvulgare-HORVU6Hr1G058840.1                  LCMAI----QH-------ADPLNWGKAAEEL--TGSHLDEVKRMVAEYRKPVVTIEGASL
Taestivum-Traes_6BL_C96F626D9.1              ------------------------------------------------------------
Tintermedium-Thint.18G0192700.1              VCMAI----QH-------ADPLNWGKAAEEL--TGSHLDEVKRMVADYRKPVVTIEGASL
Taestivum-Traes_6DL_04CD7E82C.1              VCMAI----QH-------ADPLNWGKAAEEL--TGSHLDEVKRMVADYRKPVVTIEGASL
Bdistachyon-Bradi3g49270.1                   ICTAI----QH-------ADPLNWGKAAEEL--TGSHLDEVKRMVVEYREPVVTIEGASL
Ecoracana-ELECO.r07.2AG0138020.1             LCMAM----PR-------ADPLNWGKAAE------------------YRQPLVKIEGASL
Ecoracana-ELECO.r07.2BG0193670.1             LCMAV----PR-------ADPLNWGKAAEEL--MGSHLDEVKRMVAEYRQPLVKIEGASL
Osativa-LOC_Os02g41670.1                     LCMAA----PR-------ADPLNWGKAAEEM--SGSHLDEVKRMVAEYRQPLVKIEGASL
Osativa-LOC_Os02g41680.1                     LCVAA----PR-------ADPLNWGKATEEM--TGSHLDEVKRMVAEYRQPLVKIEGASL
Pvaginatum-Pavag04G202900.1                  LCMAT----PR-------ADPLNWGKAAEEL--MGSHLDEVKRMVAEYREPLVKIEGASL
Pvaginatum-PavagK321000.1                    LCMAT----PR-------ADPLNWGKAAEEL--MGSHLDEVKRMVAEYREPLVKIEGASL
Pvaginatum-PavagK321300.1                    LCMAT----PR-------ADPLNWGKAAEEL--MGSHLDEVKRMVAEYREPLVKIEGASL
Pvaginatum-PavagK321100.1                    LCMAT----PR-------ADPLNWGKAAEEL--MGSHLDEVKRMVAEYREPLVKIEGASL
Pvaginatum-PavagK321200.1                    LCMAT----PR-------ADPLNWGKAAEEL--MGSHLDEVKRMVAEYREPLVKIEGASL
Pvirgatum-Pavir.1KG386500.1                  LCVAT----PR-------ADPLNWGKAAEEL--MGSHLDEVKRMVAEYRQPLVKIEGASL
Pvirgatum-Pavir.1NG356700.1                  LCMAT----PR-------ADPLNWGKAAEEL--MGSHLDEVKRMVAEYRQPLVKIEGASL
Sviridis-Sevir.1G245166.1                    LCMAT----PR-------ADPLNWGKAAEEL--MGSHLDEVKRMVADYRQPLVKIEGASL
Sviridis-Sevir.1G245232.1                    LCMAT----PR-------ADPLNWGKAAEEL--MGSHLDEVKRMVADYRQPLVKIEGASL
```

```
Pvirgatum-Pavir.1NG356800.1                LCMAT----PR-------ADPLNWGKAAEEL--MGSHLDEVKRMVAEYRQPLVKIEGASL
Sbicolor-Sobic.004G220500.1                LCMAT----PR-------ADPLNWGKAAEEL--MGSHLDEVKRMVAEYRQPLVKIEGASL
Sbicolor-Sobic.004G220600.2                LCMAT----PR-------ADPLNWGKAAEEL--MGSHLDEVKRMVAEYRQPLVKIEGASL
Sbicolor-Sobic.004G220700.1                LCMAT----PR-------ADPLNWGKAAEEL--MGSHLDEVKRMVAEYRQPLVKIEGASL
Sviridis-Sevir.1G245300.2                  LCMAT----PR-------ADPLNWGKAAEEL--MGSHLDEVKRMVADYRQPLVKIEGASL
Zmays-Zm00001d017276_T001                  PCMAT----PR-------ADPLNWGKAAEEL--MGSHLDEVKRMVAEYRQPLVKIEGASL
Zmays-Zm00001d017279_T001                  LCMAT----PR-------ADPLNWGKAAEEL--MGSHLDEVKRMVAEYRQPLVKIEGASL
Zmays-Zm00001d051166_T001                  LCMAL----PR------AADPLNWGKAAEEM--MGSHLDEVKRMVAEYRQPLVKIEGASL
Ecoracana-ELECO.r07.2AG0138030.1           LCMAT----PR-------ADPLNWGKAAEEL--MGSHLDEVKQMVAEYRKPLVTIEGASL
Ecoracana-ELECO.r07.2BG0193680.1           LCMAT----PR-------ADPLNWGKAAEEL--MGSHLDEVKQMVTEYRKPLVTIEGASL
Osativa-LOC_Os05g35290.1                   LCMEV----PR-------ADPLNWGKAAEEM--AGSHLDEVKRMVAEYRQPVVRIEGASL
Platifolius-Phala.02G210700.1              LCLAT----PR-------ADRLNWGKAAEEL--KGSHLDEVKRMVEEYRQPVVRIEGASL
Ecoracana-ELECO.r07.4AG0310420.1           LCMPPP---PR------AADPLNWGKAAEDL--TRSHLDAVKRMVEEYRRPVVRIEGASL
Ecoracana-ELECO.r07.4BG0341610.1           LCMTR------------AADPLNWGKAAEDL--SGSHLDAVKRMVEEYRRPVVRIEGASL
Bdistachyon-Bradi3g49260.1                 LCMSA----PP----RAAADPLNWAKTAEEL--AGSHLEEVKKMVAQFRMPLVKIEGATL
Hvulgare-HORVU0Hr1G016330.1                LCVSA----PR------AADPLNWVKTAEEL--TGSHLDEVKKMVAQFREPLVTIEGASL
Tintermedium-Thint.17G0260200.1            LCVSA----PR-------ADPLNWVKTAEEL--TGSHLDEVKKMVAQFREPLVTIEGASL
Tintermedium-Thint.18G0192600.1            LCMSA----PR-------ADPLNWVKTAEEL--TGSHLEEVKKMVAQFREPLVTIEGASL
Tintermedium-Thint.17G0231000.1            LCMAI----QH-------AAPLNWGKAAQEL--TGSHLDEVKRMVADYRKPVVTIEGASL
Ecoracana-ELECO.r07.2AG0138010.1           LCMSS----AA------RADPLNWGKAAEDL--SGSHLDEVKRMVAEFRDPVVTIQGASL
Ecoracana-ELECO.r07.2BG0193660.1           LCMSS----AAS--APRRADPLNWGKAAEDL--SGSHLDEVKRMVAEFRDPVVTIQGASL
Osativa-LOC_Os02g41650.3                   LCMSS----VS---APPRADPLNWGKAADEL--AGSHLDEVKRMVEDFRQPLVKIEGASL
Pvaginatum-Pavag04G202800.1                LCVVS----PA----PRAADPLNWGKAAEDL--SGSHVGEVKRMVAEFREPVVKIQGASL
Pvaginatum-PavagK320900.1                  LCVVS----PA----PRAADPLNWGKAAEDL--SGSHVGEVKRMVAEFREPVVKIQGASL
Pvirgatum-Pavir.1NG356400.1                LCVPA----PA----AARADPLNWGKAAEDL--SGSHLGEVQRMVAEFREPLVRIQGASL
Sviridis-Sevir.1G245100.1                  LCMSA----QA---AAPRADPLNWGKAAEDL--SGSHLDEVKRMVAEFREPLVKIQGASL
Sbicolor-Sobic.004G220400.1                LCMSA----QA---APRGADPLNWGKAAEDL--SGSHLEEVKRMVAEFRDPVVKIQGASL
Zmays-Zm00001d017275_T001                  LCMSA----AA---AAPRADALNWGKAAEDL--SGSHLDEVKRMVVEFREPLVRIQGASL
Zmays-Zm00001d051163_T002                  LCMPA----PR-------ADPLNWGKAAEGL--SGSHLDEVKRMVAEFRDPLVKIQGASL
Pvirgatum-Pavir.1KG386300.1                LCAPA----PA----PRAADPLNWGKAAEDL--SGSHLGEVQRMVAEFREPLVRIQGASL
Platifolius-Phala.02G210600.1              LCLAT----PP------RVDPLNWGKAAEEL--TGSHLDEVKSMVEEYRRPVVRIEGASL
Streptochaeta-strangu_020767-RA            LCLAK----PQ-------ADPLNWGKAAEEL--TGSHLDEVKRMVAEYREPVVRIEGAGL
Bdistachyon-Bradi5g15830.1                 LCVAA----PL----APRADPLNWGKAAEEL--SGSHLDAVKRMVEEYRRPVVKMEGASL
Hvulgare-HORVU2Hr1G089440.4                LCVAQ----P------ARADPLNWGKAAEEL--SGSHLDAVKRMVEEYRKPVVTMEGASL
Taestivum-Traes_2AL_0DDC60D6E.1            ------------------------------------------------------------
Tintermedium-Thint.06G0475900.1            LCVAQ----P------ARADPLNWGKAAEEL--SGSHLDAVKRMVEEYRRPVVTMEGASL
Tintermedium-Thint.06G0476200.1            LCVAQ----P------ARADPLNWGKAAEEL--SGSHLDAVKRMVEEYRRPVVTMEGASL
Taestivum-Traes_2AL_2763280FC.1            ------------------------------------------------------------
Taestivum-Traes_2DL_8C21AAAEF.1            ------------------------------------------------------------
Tintermedium-Thint.V0383500.1              LCVAQ----P------ARADPLNWGKAAEEL--SGSHLEAVKRMVEEYRKPVVTMEGASL
Taestivum-Traes_2AL_9EC3226F7.1            LCVAQ----P------ARADPLNWGKAAEEL--SGSHLEAVKRMVEEYRKPVVTMEGASL
Tintermedium-Thint.05G0273800.1            LCVAQ----P------ARADPLNWGKAAEEL--SGSHLEAVKRMVEEYRKPVVTMEGASL
Tintermedium-Thint.V0383400.1              LCVAQ----P------ARADPLNWGKAAEEL--SGSHLDAVKRMVEEYRKPVVTMEGASL
Taestivum-Traes_2BL_13D5272D7.1            LCVAQ----P------ARADPLNWGKAAEEL--SGSHLDAVKRMVKEYRKPVVTMEGASL
Tintermedium-Thint.06G0475700.1            LCVAQ----T------ARADPLNWGKAAEEL--SGSHLDAVKRMVEEYRKPVVTMEGASL
Taestivum-Traes_2BL_C051606EA.2            LCVAQ----P------ARADPLNWGKAADEL--SGSHLDAVKRMVEEYRKPVVTMEGASL
Tintermedium-Thint.V0383600.1              LCVAQ----P------ARPDPLNWGKAAEEL--SGSHLDAVKRMVEEYRKPVVTMEGASL
Taestivum-Traes_2AL_9D78F85E2.1            LCVAQ----P------ARADPLNWGKAAEEL--SGSHLDAVKRMVEEYRKPVVTMEGASL
Tintermedium-Thint.05G0274100.1            LCVAQ----P------ARADPLNWGKAAEEL--SGSHLDAVKRMVEEYRKPVVTMEGASL
Tintermedium-Thint.06G0476300.1            LCVAQ----P------ARADPLNWGKAAEEL--SGSHLDAVKRMVEEYRKPVVTMEGASL
Taestivum-Traes_2BL_5A50FDA1A.1            ------------------------------------------------------------
Tintermedium-Thint.05G0273900.1            ------------------------------------------------------------
Taestivum-Traes_2DL_30F8555D4.1            ------------------------------------------------------------
Taestivum-Traes_2DL_83168C1E0.1            ------------------------------------------------------------
Taestivum-Traes_2DL_B978B4E22.1            ------------------------------------------------------------
Taestivum-Traes_1AS_0B9295A68.1            LCVAQ----P------ARADPLNWGKAAEGL--SGSHLDAVKQMVEEYRRPVVTMEGASL
Taestivum-Traes_1BS_7068F56E6.1            LCVAQ----P------ARADPLNWGKAAEEL--SVSHLDAVKRMVEEYRRPVVTMEGASL
Taestivum-Traes_1BS_AEA817616.1            ------------------------------------------------------------
Taestivum-Traes_4AL_892C47ED5.1            LCLAQ----P------ARADPLNWGKAAEEL--SGSHLDAVKRMVEEYRRPVVTMEGASL
Tintermedium-Thint.03G0055900.1            LCVAQ----P------ARADPLNWGKAAEEL--SGSHLDAVKRMVEEYRRPVVTMEGASL
Tintermedium-Thint.01G0012900.1            FCVAQ----P------ARADPLNWGKAAEDL--SGSHLDAVKRMVEEYRRPVVTMEGASL
Tintermedium-Thint.02G0158500.1            LCVAQ----P------ARADPLNWGKAAEEL--SGSHLDAVKRMVEEYRRPVVTMEGASL
Tintermedium-Thint.03G0055700.1            LCVAQ----P------ARADPLNWGKAAEEL--SGSHLDAVKRMVEEYRRPVVTMEGASL
Taestivum-Traes_1BS_C3AA04888.1            ------------------------------------------------------------
Taestivum-Traes_1DS_A8BD91E4A.1            LCVVQ----P------ARADPLNWGKAAEEL--SGSHLDAVKRMVEEYRRPVVTMEGASL
Taestivum-Traes_1BS_C26CCE0E9.1            LCVAQ----P------ARADPLNWGKAAEEL--SGSHLDGVKRMVEEYRRPVVTMEGASL
Taestivum-Traes_1DS_A21FCBA3D.1            LCVAQ----P------ARADPLNWGKAAEEL--SGSHLDAVKRMVEEYRRPVVTMEGASL
Tintermedium-Thint.01G0012200.1            LCVAQ----P------ARADPLNWGKAAEEL--SGSHLDAVKRMVEEYHRPVVTMEGASL
Tintermedium-Thint.02G0158600.1            LCVVQ----P------ARADPLNWGKAAEEL--SGSHLDAVKRMVEEYRRPVVTMEGASL
Tintermedium-Thint.01G0012700.1            LCVAQ----P------ARADPLNWGKAAEEL--SGSHLEAVKRMVEEYRRPVVTMEGASL
Taestivum-Traes_5BL_F50A47503.1            LCVAQ----P------GRADPLNWGKAAEEL--SGSHLDAVKRMVEEYRRPVVTMEGASL
Tintermedium-Thint.02G0158700.1            LCVAQ----P------ARADPLNWGKAAEEL--SGSHLDAVKRMVEEYRRPMVTMEGASL
Taestivum-Traes_5BL_AB528184A.1            LCVAQ----P------ARADPLNWGKAAEEL--SGSHLEAVKRMVEEYRRPVVTMEGASL
Tintermedium-Thint.03G0055800.1            LCVAQ----P------ARADPLNWGKAAEEL--SGSHLDAVKRMVEEYRRPVVTMEGASL
Taestivum-Traes_1AS_F9013A945.1            LCVAQ----L------ARADPLNWGKAAEEL--SGSHLDAVKRMVEEYRRPVVTMEGASL
Taestivum-Traes_1DS_43DAC7A7F.1            LCVAQ----P------ARADPLNWGKAAEGL--SGSHLDAVKRMVEEYRRPVVTMEGASL
Taestivum-Traes_1BS_BD86C90A7.1            LCVAQ----P------ARADPLNWGKAAEEL--SGSHLDAVKRMVEEYRRPVVTMEGASL
Tintermedium-Thint.02G0158900.1            LCVAQ----P------ARADPLNWGKAAEEL--SGSHLDAVKRMVEEYRRPVVTMEGASL
Tintermedium-Thint.05G0274000.1            ------------------------------------------------------------
Osativa-LOC_Os04g43800.1                   LCVAK----P-------RADPLNWGKAAEEL--SGSHLDAVKRMVEEYRRPVVTIEGASL
Pvaginatum-Pavag06G160300.1                VCLAA----P-------RADPLNWGKAAVDL--TGSHLEAVKRMVEEYRRPLVKIEGASL
Sbicolor-Sobic.006G148900.1                VCLAT----P------RAADPLNWGKAAEDL--TGSHLEAVKRMVEEYRRPLVKIEGASL
Pvirgatum-Pavir.7KG237800.1                VCLPV----P------PRADPLNWGKAAENL--AGSHLEAVKRMVEEYRRPLVRIEGASL
Pvirgatum-Pavir.7NG355800.1                VCLPV----P------PRADPLNWGKAAEDL--AGSHLEAVRRMVEEYRRPLVKIEGASL
Sviridis-Sevir.7G178300.1                  VCLPV----P------PRADPLNWGKAAEDL--SGSHLEAVKRMVEEYRRPLVRIEGSSL
Zmays-Zm00001d003015_T001                  VCLAT----PA-----PRADPLNWAKAAEDL--AGSHLDAVKRMVEEYRRPLVKIEGASL
Hvulgare-HORVU1Hr1G022060.1                LCVAK----PR-----S-ADPLNWGKAAEEL--SGSHLDAVKRMVEEYRRPVVVMEGASL
Taestivum-Traes_1BS_723922D171.2           LCVAK----PR-----SAADPLNWGKAAAEL--SGSHVDAVKRMVEEYRRPVVVMEGASL
Taestivum-Traes_1DS_A171C7D59.1            LCVAE----PR-----SAADPLNWGKAAEEL--SGSHLDAVKRMVEEYRRPVVVMEGASL
Tintermedium-Thint.01G0111500.1            LCVAE----PR-----SAADPLNWGKAAEEL--SGSHLDAVKRMVEEYRRPVVVMEGASL
Tintermedium-Thint.03G0265300.1            LCVAK----PG-----NAADPLNWGKAAAEL--SGSHLDAVKRMVEEYRRPVVVMEGASL
Hvulgare-HORVU2Hr1G038140.1                LCVAE----PR-----S-ADPLNWGKAAAEL--SGSHLDAVKRMVEEYRRPVVVMEGASL
Tintermedium-Thint.04G0171200.1            ------------------------------------------------------------
Taestivum-Traes_2AS_958327519.3            ------------------------------------------------------------
Taestivum-Traes_2DS_28CA50371.1            ------------------------------------------------------------
Tintermedium-Thint.05G0160800.1            LCVAE----PR-----S-ADPLNWGKAAQEL--SGSHLDAVKRMVEEYRRPVVVMEGASL
Tintermedium-Thint.06G0299000.1            LCVAE----PR-----S-ADPLNWGKAAEAL--SGSHLDAVKRMVEEYRRPVVVMEGASL
Taestivum-Traes_2DS_3791A8A36.1            ------------------------------------------------------------
Tintermedium-Thint.06G0342800.1            -------------------------------------------MVEEYRRPVVVMEGASL
Tintermedium-Thint.05G0150900.1            ------------------------------------------------------------
Tintermedium-Thint.04G0242300.1            LCVAK----PR-----S-ADPLNWGKAAEEL--SGSHLDAVKQMVEEYRRPVVVMEGASL
Tintermedium-Thint.06G0298700.1            LCVAK----PP-----S-ADPLNWGKAAEEL--SGSHLDAVKRMVEEYRRPVVVMEGASL
Taestivum-Traes_2BS_88CF42F2E.1            -------------------------------------------MVEEYRRPVVVMEGASL
Tintermedium-Thint.04G0241900.1            LCVAE----PR-----S-ADPLNWGKAAQEL--SGSHLDAVKRMVEEYRRPVVVMEGASL
Platifolius-Phala.06G165300.1              LCLER----PQ-------ADPLNWGKAAEEL--AGSHLDAVKRMVEEYRKPAVRIEGASL
Streptochaeta-strangu_019385-RA            LCLAT----PQ-------ADPLNWGKAAAEL--TGSHLDEVKRMVAEYRKPVVRIEGANL
Streptochaeta-strangu_028944-RA            LRLAR----PY-------ADPLNWGTAAVEL--TGSHLEEVKRMVAESRLPVVTIEGAGL
Acomosus-Aco010091.1                       LSM---------------EDPLNWAGAAEPT--KVSHFDEVKRMVEEFRKPLVRLAGANL
Zmarina-Zosma03g22300.1                    ------------------ADPLKWEEAAEGM--KGSHVAEVKQMIEDFRNPLVVLEGKNL
Ecdeiocolea-Emoptg0003741_1G000600.1       LCLRAG------------GDPLNWGKAAEEM--AGSHLEEVKRMVAEFRAPAVRIEGASL
Jascendens-Joasc.05G060500.1               LCLQK----PQH------ADPLNWGKAAGEL--MGSHLEEVKRMVAEFRAPVVKIEGASL
Ecdeiocolea-Emoptg0003741_1G000630.1       VCIPK--------------DALNWAKAAEEL--AGSHLEEVKRMVAEFRSPAVRIEGASL
Jascendens-Joasc.05G060400.1               LCIKK--------------DPLNWGKAAEEL--TGSHLDEVKRMVAESRTPVVKIQGASL
Bdistachyon-Bradi3g49250.2                 --ISE-------------KDPLNWGAAAAEL--TGSHLDEVKRMVAQFREPVVKIEGASL
Hvulgare-HORVU6Hr1G058820.1                --IVE-------------KDPLNWGAAAAEL--TGSHLDEVKRMVAQFREPVVKIEGASL
Taestivum-Traes_6DL_1AEA7B869.1            --IVE-------------KDPLNWGAAAAEL--TGSHLDEVKRMVAQFREPVVKIEGASL
Tintermedium-Thint.18G0192300.1            --IVE-------------KDPLNWGAAAAEL--TGSHLDEVKRMVAQFREPVVKIEGASL
Tintermedium-Thint.17G0230800.1            --IVE-------------KDPLNWGAAAAEL--TGSHLDEVKRMVAQFREPVVKIEGASL
Tintermedium-Thint.17G0230900.1            --IVE-------------KDPLNWGAAAAEL--TGSHLDEVKRMVAQFREPVVKIEGASL
Ecoracana-ELECO.r07.2AG0138000.1           --ILE-------------SDPLNWGAAAAEM--AGSHLDEVKRMVEQARQPVVKIEGSTL
Ecoracana-ELECO.r07.2BG0193650.1           --ILE-------------SDPLNWGAAAAEM--AGSHLDEVKRMVEQARQPVVKIEGSTL
Pvaginatum-Pavag04G202700.1                --IVE-------------NDPLNWGAAAAEL--AGSHLDEVKRMVAQARQPVVKIEGSTL
Sviridis-Sevir.1G245000.1                  --IVE-------------NDPLNWGAAAAEL--AGSHLDEVKRMVAQARQPVVKIEGSTL
Sviridis-Sevir.6G187100.1                  --IVE-------------NDPLNWGAAAAEL--AGSHLDEVKRMVAQARQPVVKIEGSTL
Pvirgatum-Pavir.1NG356200.1                --IVE-------------NDPLNWGAAAAEL--AGSHLDEVKRMVAQARQPVVKIEGSTL
Sbicolor-Sobic.004G220300.1                --IVE-------------SDPLNWGAAAAEL--SGSHLDEVKRMVAQARQPVVKIEGSTL
Zmays-Zm00001d017274_T001                  --IVE-------------SDPLNWGAAAAEL--AGSHLDEVKRMVAQARQPVVKIEGSTL
Zmays-Zm00001d051161_T003                  --IVE-------------SDPLNWGAAAAEL--AGSHLDEVKRMVAQARQPVVKIEGSTL
Sviridis-Sevir.7G178200.1                  --ILE-------------SDPLNWGKAAAEM--AGSHLDEVKRMVAQFREPLVKIEGSSL
Osativa-LOC_Os02g41630.2                   --INK-------------EDPLNWGAAAAEM--AGSHLDEVKRMVAQFREPLVKIQGATL
```

```
Ecoracana-ELECO.r07.4AG0310430.1        --IVV-------------SDPLNWGKAASEM--TGSHLEEVKRMVAQSREPVVKIEGSSL
Ecoracana-ELECO.r07.4BG0341620.1        --IVE-------------SDPLNWGKAASEM--TGSHLEEVKRMVAQSREPVVKIEGSSL
Pvaginatum-Pavag06G160200.1             --ILA-------------SDPLNWGKAAAEL--AGSHLDEVKRMVAQYRDPVVKIEGSSL
Pvirgatum-Pavir.7KG238255.1             --ICE-------------SDPLNWGKAAAEM--AGSHLDEVKRMVAQFREPLVKIEGSSL
Sbicolor-Sobic.006G148800.1             --ILE-------------SDPLSWGKAAAEL--TGSHLDEVKRMVAQFRDPVVKIEGSTL
Zmays-Zm00001d003016_T001               --ILE-------------SDPLNWGKAAAEL--TGSHLDEVKRMVAQFRDPVVKIEGSSL
Osativa-LOC_Os04g43760.1                TGFVA-------------SDPLSWGKAALEM--TGSHLDEVKRMVAQSREAVVKIEGSSL
Platifolius-Phala.02G210400.1           L-SLP-------------KDPLNWGKAAQEM--MGSHFDEVKRMVAQSREPVVKIEGASL
Platifolius-Phala.06G165100.1           --ALP-------------SDPLNWGKAAEEL--TGSHLDEVKRMVAQFREPAVKIEGASL
Streptochaeta-strangu_019386-RA         VSIL--------------RDPLNWGKAAEEL--TGSHFDEVKRMVAQFREPAVTIEGASL
Streptochaeta-strangu_020769-RA         ------------------RDPLNWAAAAAEL--TGSHLDEVKRMVAQFREPQVKIEGASL
Sviridis-Sevir.2G448300.1               --IVT-------------SDPLNWGKAAAEL--TGSHLDEVRRMVAQSREPIVRVDGSRL
Sviridis-Sevir.7G177900.1               --IVT-------------SDPLNWGKAAAEL--TGSHLDEVRRMVAQSREPVVRVDGSRL


Aamericanus-Acora.01G232100.1           KISQVAAVA-----A---------------------------------------------
Aamericanus-Acora.10G174800.1           KISQVAAVA-----A---------------------------------------------
Acomosus-Aco013943.1                    ----VAAVA---G-A-----P---------------------------------------
Aofficinalis-evm.model.AsparagusV1_01.827 ------------------------------------------------------------
Typha_latifolia-Tylat.14G083800.1       KISQVAAVA---A-A---------------------------------------------
Aofficinalis-evm.model.AsparagusV1_04.71 ------------------------------------------------------------
Aofficinalis-evm.model.AsparagusV1_04.93 ------------------------------------------------------------
Aofficinalis-evm.model.AsparagusV1_10.1948 ------------------------------------------------------------
Typha_latifolia-Tylat.03G127700.1       NISQVAAVA---A-A---------------------------------------------
Spolyrhiza-Spipo11G0025500              KISQVAAVA-----S---------------------------------------------
Spolyrhiza-Spipo1G0003500               KISQVAAVA---A-A---------------------------------------------
Phoenix_dactylifera-LOC103718264        KISQVAAVA---T-S---------------------------------------------
Macuminata-GSMUA_Achr1T09070_001        TISQVAAVA---A-A---------------------------------------------
Acomosus-Aco006987.1                    KISQVAAVA---M-G-DF----------------AQAQAQ----------------AQPV
Acomosus-Aco027752.1                    KISQVAAVA---M-G-DF----------------AQAQAQ----------------AQAQ
Cscoparia-Csco27g03980.1                KISQVAAVA---M---KE------------------------------------------
Cscoparia-Csco27g03990.1                KISQVAAVA---M---KE------------------------------------------
Macuminata-GSMUA_Achr11T16380_001       RISQVAAIA---AGA---------------------------------------------
Macuminata-GSMUA_Achr11T22840_001       KISQVAAVA---M-A-PH------------------------------------------
Macuminata-GSMUA_Achr2T00240_001        TISQVAAVA---T-A---------------------------------------------
Macuminata-GSMUA_Achr5T18560_001        KISQVAAVA---T-E-PH------------------------------------------
Macuminata-GSMUA_Achr8T18960_001        NISQVAAVA---M-A-AH------------------------------------------
Macuminata-GSMUA_Achr5T03950_001        KIAHVAAVA---G-Q---------------------------------------------
Phoenix_dactylifera-LOC103703950        KISQVAAVA---M-A---------------------------------------------
Acoerulea-Aqcoe1G140100.1               TVSQVAGIA---A-H-G-------------------------------------------
Acoerulea-Aqcoe2G144500.1               TISQVAAIG---S-H-D-------------------------------------------
Typha_latifolia-Tylat.05G053500.1       KISQVAAIA---M-S---------------------------------------------
Zmarina-Zosma05g29470.1                 KISQVAAVA---T-A---------------------------------------------
Acomosus-Aco007727.1                    KISQVAAVA---M-A-KAVAPGVRVELAEAARGGVEASSEWVMESMARGTDSYGVAAVAM
Bvulgarisssp-Bevul.7G054300.1           TVAQVAAVA---A-A---------------------------------------------
Bdistachyon-Bradi3g47110.1              SIAKVAAVA---A-A-GE------------------------------------------
Bdistachyon-Bradi3g47120.1              SIAKVAAVA---A-A-GE------------------------------------------
Bdistachyon-Bradi3g48840.1              SIAKVAAVA---A-A-GE------------------------------------------
Bdistachyon-Bradi3g49280.1              SIAKVAAVA---A-A-GE------------------------------------------
Hvulgare-HORVU6Hr1G058840.1             SIAKVAAVA---A-A-GE------------------------------------------
Taestivum-Traes_6BL_C96F626D9.1         ------------------------------------------------------------
Tintermedium-Thint.18G0192700.1         SIAKVAAVA---A-A-GE------------------------------------------
Taestivum-Traes_6DL_04CD7E82C.1         SIAKVAAVA---A-A-GE------------------------------------------
Bdistachyon-Bradi3g49270.1              SIAKVAAVA---A-A-GE------------------------------------------
Ecoracana-ELECO.r07.2AG0138020.1        TIAQVAAVA---AGD-GE------------------------------------------
Ecoracana-ELECO.r07.2BG0193670.1        TIAQVAAIA---AGD-GE------------------------------------------
Osativa-LOC_Os02g41670.1                RIAQVAAVA---A-A-GE------------------------------------------
Osativa-LOC_Os02g41680.1                RIAQVAAVA---A-A-GE------------------------------------------
Pvaginatum-Pavag04G202900.1             RIAQVAAVA---AGA-GE------------------------------------------
Pvaginatum-PavagK321000.1               RIAQVAAVA---AGA-GE------------------------------------------
Pvaginatum-PavagK321300.1               RIAQVAAVA---AGA-GE------------------------------------------
Pvaginatum-PavagK321100.1               RIAQVAAVA---AGA-GE------------------------------------------
Pvaginatum-PavagK321200.1               RIAQVAAVA---AGA-GE------------------------------------------
Pvirgatum-Pavir.1KG386500.1             RIAQVAAVA---NGA-GE------------------------------------------
Pvirgatum-Pavir.1NG356700.1             RIAQVAAVA---NGA-GE------------------------------------------
Sviridis-Sevir.1G245166.1               NIAQVAAVA---NGA-GE------------------------------------------
Sviridis-Sevir.1G245232.1               NIAQVAAVA---NGA-GE------------------------------------------
Pvirgatum-Pavir.1NG356800.1             RIAQVAAVA---NGA-GE------------------------------------------
Sbicolor-Sobic.004G220500.1             RIAQVAAVA---AGA-GE------------------------------------------
Sbicolor-Sobic.004G220600.2             RIAQVAAVA---AGA-GE------------------------------------------
Sbicolor-Sobic.004G220700.1             RIAQVAAVA---AGA-GE------------------------------------------
Sviridis-Sevir.1G245300.2               NIAQVAAVA---NGA-GE------------------------------------------
Zmays-Zm00001d017276_T001               SIAQVAAVA---TGV-GE------------------------------------------
Zmays-Zm00001d017279_T001               SIAQVAAVA---TGA-GE------------------------------------------
Zmays-Zm00001d051166_T001               RIAQVAAVA---AGA-GE------------------------------------------
Ecoracana-ELECO.r07.2AG0138030.1        SIAQVAAVA---TGA-GE------------------------------------------
Ecoracana-ELECO.r07.2BG0193680.1        SIAQVAAVA---AGA-GE------------------------------------------
Osativa-LOC_Os05g35290.1                RIAQVAAVA---G-A-GD------------------------------------------
Platifolius-Phala.02G210700.1           RIAQVAAVA---TGA-GE------------------------------------------
Ecoracana-ELECO.r07.4AG0310420.1        TVAQVAAVAN--A-A-GE------------------------------------------
Ecoracana-ELECO.r07.4BG0341610.1        TVAQVAAVAN--A-A-GE------------------------------------------
Bdistachyon-Bradi3g49260.1              GIAQVAAVA---AGA-GE------------------------------------------
Hvulgare-HORVU0Hr1G016330.1             GIAQVAAVA---AGA-GV------------------------------------------
Tintermedium-Thint.17G0260200.1         GIAQVAAVA---AGA-GV------------------------------------------
Tintermedium-Thint.18G0192600.1         GIAQVAAVA---AGA-GV------------------------------------------
Tintermedium-Thint.17G0231000.1         SIAKVAAVA---A-A-GE------------------------------------------
Ecoracana-ELECO.r07.2AG0138010.1        SIAQVAAVA---TGA-GE------------------------------------------
Ecoracana-ELECO.r07.2BG0193660.1        SIAQVAAVA---TGA-GE------------------------------------------
Osativa-LOC_Os02g41650.3                TIAQVAAVA---AGA-GD------------------------------------------
Pvaginatum-Pavag04G202800.1             SIAQVAAVA---AGA-GE------------------------------------------
Pvaginatum-PavagK320900.1               SIAQVAAVA---AGA-GE------------------------------------------
Pvirgatum-Pavir.1NG356400.1             SIAQVAAVA---AGA-GE------------------------------------------
Sviridis-Sevir.1G245100.1               SIAQVAAVA---AGA-GE------------------------------------------
Sbicolor-Sobic.004G220400.1             SIAQVAAVA---AGA-GE------------------------------------------
Zmays-Zm00001d017275_T001               SIGQVAAVA---AGAGGE------------------------------------------
Zmays-Zm00001d051163_T002               SVAQVAAVAVG-AGG-GE------------------------------------------
Pvirgatum-Pavir.1KG386300.1             SIAQVAAVA---AGT-GE------------------------------------------
Platifolius-Phala.02G210600.1           RIAQVAAVA---AGA-GE------------------------------------------
Streptochaeta-strangu_020767-RA         RIAQVAAVA-AAADG-GE------------------------------------------
Bdistachyon-Bradi5g15830.1              TIAQVAAVA---A-G-AE------------------------------------------
Hvulgare-HORVU2Hr1G089440.4             TIAMVAAVA---A-G-SD------------------------------------------
Taestivum-Traes_2AL_0DDC60D6E.1         ------------------------------------------------------------
Tintermedium-Thint.06G0475900.1         TIAMVAAVA---A-G-SD------------------------------------------
Tintermedium-Thint.06G0476200.1         TIAMVAAVA---A-G-SD------------------------------------------
Taestivum-Traes_2AL_2763280FC.1         ------------------------------------------------------------
Taestivum-Traes_2DL_8C21AAAEF.1         ------------------------------------------------------------
Tintermedium-Thint.V0383500.1           TIAMVAAVA---A-G-SD------------------------------------------
Taestivum-Traes_2AL_9EC3226F7.1         TIAMVAAVA---A-G-SD------------------------------------------
Tintermedium-Thint.05G0273800.1         TIAMVAAVA---A-G-SD------------------------------------------
Tintermedium-Thint.V0383400.1           TIAMVAAVA---A-G-SD------------------------------------------
Taestivum-Traes_2BL_13D5272D7.1         TIAMVAAVA---A-G-SD------------------------------------------
Tintermedium-Thint.06G0475700.1         TIAMVAAVA---A-G-SD------------------------------------------
Taestivum-Traes_2BL_C051606EA.2         TIAMVAAVA---A-G-SD------------------------------------------
Tintermedium-Thint.V0383600.1           TIAMVAAVA---A-G-SD------------------------------------------
Taestivum-Traes_2AL_9D78F85E2.1         TIAMVAAVA---A-G-SD------------------------------------------
Tintermedium-Thint.05G0274100.1         TIAMVAAVA---A-G-SD------------------------------------------
Tintermedium-Thint.06G0476300.1         TIAMVAAVA---A-G-SD------------------------------------------
Taestivum-Traes_2BL_5A50FDA1A.1         ---MVAAVA---A-G-SD------------------------------------------
Tintermedium-Thint.05G0273900.1         ------------------------------------------------------------
Taestivum-Traes_2DL_30F8555D4.1         ------------------------------------------------------------
Taestivum-Traes_2DL_83168C1E0.1         ------------------------------------------------------------
Taestivum-Traes_2DL_B978B4E22.1         ------------------------------------------------------------
Taestivum-Traes_1AS_0B9295A68.1         TIAMVAAVA---A-G-SD------------------------------------------
Taestivum-Traes_1BS_7068F56E6.1         TIAMVAAVA---T-G-SD------------------------------------------
Taestivum-Traes_1BS_AEA817616.1         ------------------------------------------------------------
Taestivum-Traes_4AL_892C47ED5.1         TIAMVAAVA---A-G-AE------------------------------------------
Tintermedium-Thint.03G0055900.1         TIAMVAAVA---A-G-AD------------------------------------------
Tintermedium-Thint.01G0012900.1         TIAMVAAVA---A-G-SD------------------------------------------
```

```
Tintermedium-Thint.02G0158500.1              TIAMVAAVA---A-G-SD------------------------------------------
Tintermedium-Thint.03G0055700.1              TIAMVAAVA---A-G-SD------------------------------------------
Taestivum-Traes_1BS_C3AA04888.1              ------------------------------------------------------------
Taestivum-Traes_1DS_A8BD91E4A.1              TIAMVAAVA---A-G-SD------------------------------------------
Taestivum-Traes_1BS_C26CCE0E9.1              TIAMVAAVA---A-G-SD------------------------------------------
Taestivum-Traes_1DS_A21FCBA3D.1              TIAMVAAVA---A-G-SD------------------------------------------
Tintermedium-Thint.01G0012200.1              TIAMVAAVA---A-G-SD------------------------------------------
Tintermedium-Thint.02G0158600.1              TIAMVAAVA---A-G-SE------------------------------------------
Tintermedium-Thint.01G0012700.1              TIAMVAAVA---A-G-SD------------------------------------------
Taestivum-Traes_5BL_F50A47503.1              TIAMVAAVA---A-G-GE------------------------------------------
Tintermedium-Thint.02G0158700.1              TIAMVAAVA---A-G-AE------------------------------------------
Taestivum-Traes_5BL_AB528184A.1              TIAMVAAVA---A-G-SD------------------------------------------
Tintermedium-Thint.03G0055800.1              TIAMVAAVA---A-G-AE------------------------------------------
Taestivum-Traes_1AS_F9013A945.1              TIAMVAAVA---A-G-SD------------------------------------------
Taestivum-Traes_1DS_43DAC7A7F.1              TIAMVAAVA---A-G-SD------------------------------------------
Taestivum-Traes_1BS_BD86C90A7.1              TIAMVAAVA---A-G-CD------------------------------------------
Tintermedium-Thint.02G0158900.1              TIAMVAAVA---A-G-SD------------------------------------------
Tintermedium-Thint.05G0274000.1              ------------------------------------------------------------
Osativa-LOC_Os04g43800.1                     TIAQVAAVA---S-A-GA------------------------------------------
Pvaginatum-Pavag06G160300.1                  TVAQVAAVA---A-A-GE------------------------------------------
Sbicolor-Sobic.006G148900.1                  TVAQVAAVA---A-A-GE------------------------------------------
Pvirgatum-Pavir.7KG237800.1                  TVAQVAAVA---A-A-GE------------------------------------------
Pvirgatum-Pavir.7NG355800.1                  TVAQVAAVA---A-A-AE------------------------------------------
Sviridis-Sevir.7G178300.1                    TVAQVAAVAN--G-A-GE------------------------------------------
Zmays-Zm00001d003015_T001                    TVAQVAAVA---A-A-GE------------------------------------------
Hvulgare-HORVU1Hr1G022060.1                  TIAQVAAVA---A-A-GG------------------------------------------
Taestivum-Traes_1BS_723922D171.2             TIAQVAAVA---A-A-GG------------------------------------------
Taestivum-Traes_1DS_A171C7D59.1              TIAQVAAVA---A-A-GG------------------------------------------
Tintermedium-Thint.01G0111500.1              TIAQVAAVA---A-A-GG------------------------------------------
Tintermedium-Thint.03G0265300.1              TIAQVAAVA---A-A-GG------------------------------------------
Hvulgare-HORVU2Hr1G038140.1                  TIAQVAAVA---A-A-DG------------------------------------------
Tintermedium-Thint.04G0171200.1              ------------------------------------------------------------
Taestivum-Traes_2AS_958327519.3              ------------------------------------------------------------
Taestivum-Traes_2DS_28CA50371.1              ------------------------------------------------------------
Tintermedium-Thint.05G0160800.1              TIAQVAAVA---A-A-DG------------------------------------------
Tintermedium-Thint.06G0299000.1              TIAQVAAVA---A-A-DG------------------------------------------
Taestivum-Traes_2DS_3791A8A36.1              ------------------------------------------------------------
Tintermedium-Thint.06G0342800.1              TIAQVAAVA---A-A-DG------------------------------------------
Tintermedium-Thint.05G0150900.1              ------------------------------------------------------------
Tintermedium-Thint.04G0242300.1              TIAQVAAVA---A-A-GG------------------------------------------
Tintermedium-Thint.06G0298700.1              TIAQVAAVA---A-A-DG------------------------------------------
Taestivum-Traes_2BS_88CF42F2E.1              TIAQVAAVA---T-A-DG------------------------------------------
Tintermedium-Thint.04G0241900.1              TIAQVAAVA---A-A-NG------------------------------------------
Platifolius-Phala.06G165300.1                TIAQVAAVA---A-G-GD------------------------------------------
Streptochaeta-strangu_019385-RA              SIAQVVAVAL--A-G-GA------------------------------------------
Streptochaeta-strangu_028944-RA              RIAHVAAVA---T-ACGG------------------------------------------
Acomosus-Aco010091.1                         TISQVAAVA-----A-SE------------------------------------------
Zmarina-Zosma03g22300.1                      KIAHVAAVA-------GD------------------------------------------
Ecdeiocolea-Emoptg0003741_1G000600.1         RIAQVAAVA-----A-GG------------------------------------------
Jascendens-Joasc.05G060500.1                 RIAQVAAVA-----A-GE------------------------------------------
Ecdeiocolea-Emoptg0003741_1G000630.1         RVGQVAAVA-----G-AE------------------------------------------
Jascendens-Joasc.05G060400.1                 RIGQVAAVA-----V-RE------------------------------------------
Bdistachyon-Bradi3g49250.2                   RVGQVAAVA---Q-A-KD------------------------------------------
Hvulgare-HORVU6Hr1G058820.1                  RVGQVAAVA---Q-A-KD------------------------------------------
Taestivum-Traes_6DL_1AEA7B869.1              RVGQVAAVA---Q-A-KD------------------------------------------
Tintermedium-Thint.18G0192300.1              RVGQVAAVA---Q-A-KD------------------------------------------
Tintermedium-Thint.17G0230800.1              RVGQVAAVA---Q-A-KD------------------------------------------
Tintermedium-Thint.17G0230900.1              RVGQVAAVA---Q-A-KD------------------------------------------
Ecoracana-ELECO.r07.2AG0138000.1             RVGQVAAVA---A-A-KD------------------------------------------
Ecoracana-ELECO.r07.2BG0193650.1             RVGQVAAVA---A-A-KD------------------------------------------
Pvaginatum-Pavag04G202700.1                  RVGQVAAVA---A-A-KD------------------------------------------
Sviridis-Sevir.1G245000.1                    RVGQVAAVA---S-A-RD------------------------------------------
Sviridis-Sevir.6G187100.1                    RVGQVAAVA---A-A-KD------------------------------------------
Pvirgatum-Pavir.1NG356200.1                  RVGQVAAVA---A-A-RD------------------------------------------
Sbicolor-Sobic.004G220300.1                  RVGQVAAVA---S-A-KD------------------------------------------
Zmays-Zm00001d017274_T001                    RVGQVAAVA---S-A-KD------------------------------------------
Zmays-Zm00001d051161_T003                    RVGQVAAVA---S-A-RD------------------------------------------
Sviridis-Sevir.7G178200.1                    RVGQVAAVA---A-A-KD------------------------------------------
Osativa-LOC_Os02g41630.2                     RVGQVAAVA---Q-A-KD------------------------------------------
Ecoracana-ELECO.r07.4AG0310430.1             RVGQVAAVA---A-A-KD------------------------------------------
Ecoracana-ELECO.r07.4BG0341620.1             RVGQVAAVA---A-A-KD------------------------------------------
Pvaginatum-Pavag06G160200.1                  RVGQVAAVA---A-A-KD------------------------------------------
Pvirgatum-Pavir.7KG238255.1                  RVGQVAAVA---V-A-KD------------------------------------------
Sbicolor-Sobic.006G148800.1                  RVGQVAAVA---A-A-KD------------------------------------------
Zmays-Zm00001d003016_T001                    RVGQVAAVA---A-A-KD------------------------------------------
Osativa-LOC_Os04g43760.1                     RVGQVAAVS---A-A-KD------------------------------------------
Platifolius-Phala.02G210400.1                RVGQVASVA---A-A-RD------------------------------------------
Platifolius-Phala.06G165100.1                RVGQVASVA---A-A-RD------------------------------------------
Streptochaeta-strangu_019386-RA              RVGQVASVA---A-A-RD------------------------------------------
Streptochaeta-strangu_020769-RA              RVGQVAAVA---A-A-RD------------------------------------------
Sviridis-Sevir.2G448300.1                    HVGKVAAVA---A-A-KD------------------------------------------
Sviridis-Sevir.7G177900.1                    HVGQVAAVA---A-A-KD------------------------------------------
```

```
Aamericanus-Acora.01G232100.1                --EGGGHVRVELSEAARPRVKASSDWVMESMNKGTDSYGVTTGFGATSHRRTKQGGALQK
Aamericanus-Acora.10G174800.1                --GGGEGVRVQLSETARPRVKASSDWVMDSMSKGTDSYGVTTGFGATSHRRTKQGGALQK
Acomosus-Aco013943.1                         --P-----------------AAGAAW-----PRAPTAYRRHHGFGATSHRRTKEGAASRG
Aofficinalis-evm.model.AsparagusV1_01.827    ------------------------------------------------------------
Typha_latifolia-Tylat.14G083800.1            ---SDAAPRVELSEAARAGVKASSDWVMESMNKGTDSYGVTTGFGATSHRRTKQGGALQK
Aofficinalis-evm.model.AsparagusV1_04.71     ------------------------------------------------------------
Aofficinalis-evm.model.AsparagusV1_04.93     ------------------------------------------------------------
Aofficinalis-evm.model.AsparagusV1_10.1948   ------------------------------------------------------------
Typha_latifolia-Tylat.03G127700.1            ----ESAVRVELSESAREGVKASSDWVMDSMMKGTDSYGVTTGFGATSHRRTKQGGALQK
Spolyrhiza-Spipo11G0025500                   ---GRGTLRVELAESARERVKASSDWVMDSMNKGTDSYGVTTGFGATSHRRTKQGGALQR
Spolyrhiza-Spipo1G0003500                    --GGGASLRVELADSARPRVKASSDWVMDSMNRGTDSYGVTTGFGATSHRRTKQGAALQK
Phoenix_dactylifera-LOC103718264             ----ESPVTVELSEAARDGVKASSDWVMESMNKGTDSYGVTTGFGATSHRRTKQGGALQK
Macuminata-GSMUA_Achr1T09070_001             ----RSPVRVELSEEARDGVRASSEWVMESMNKGTDSYGVTTGFGATSHRRTKQGGALQK
Acomosus-Aco006987.1                         PVPVPVPVRVELSEGARDAVRASSDWVLASMDKGTDSYGVTTGFGATSHRRTKQGGALQR
Acomosus-Aco027752.1                         PVPVPVPVRVELSEGARDAVRASSDWVLASMDKGTDSYGVTTGFGATSHRRTKQGGALQR
Cscoparia-Csco27g03980.1                     ----VEVVKVELNESARERVKASSDWVMDSMNRGTDSYGVTTGFGATSHRRTKEGGALQK
Cscoparia-Csco27g03990.1                     ----VEAVKVELNESARERVKASSDWVMDSMNKGTDSYGVTTGFGATSHRRTKEGGALQR
Macuminata-GSMUA_Achr11T16380_001            ----HSAVRVELSESARDEVCASSRWVVDSTAKGTNSNGMTAGFGAASHRRTKEGGALQK
Macuminata-GSMUA_Achr11T22840_001            -----SAVRVELSESAREGVRASSQWVTDSMTNGTDSYGVTTGFGATSHRRTKEGGALQK
Macuminata-GSMUA_Achr2T00240_001             -----AAVRVELSESARDGVRASSQWVVDSMTKGTDSYGVTTGFGATSHRRTKEGGALQK
Macuminata-GSMUA_Achr5T18560_001             -----SAVRVELSESAREGVRASSQWVMDSMTKGTDSYGVTTGFGATSHRRTKQGGALQQ
Macuminata-GSMUA_Achr8T18960_001             ----SSAVRVELSESARDRVSASSEWVMDSMTKGTDSYGVTTGFGATCHRRTREGGALQR
Macuminata-GSMUA_Achr5T03950_001             -----SGARVELSESARGRVRASSDWVMDSMTKGTDSYGVTTGFGATSHRRTKQGGALQK
Phoenix_dactylifera-LOC103703950             ----ESAVRVELSEASRDGVQASSQWVMNSMSNGTDSYGVTTGFGATSHRRTKQGGALQK
Acoerulea-Aqcoe1G140100.1                    -----ANVKVELCESARAGVKASSDWVMESMNKGTDSYGVTTGFGATSHRRTKQGGALQK
Acoerulea-Aqcoe2G144500.1                    -----SGVKVELSESARAGVKASSDWVMESMNKGTDSYGVTTGFGATSHRRTKQGGALQK
Typha_latifolia-Tylat.05G053500.1            ----GEGVRVELSEAARPRVRASSEWVLESMAKGVDSYGVTTGFGSTSHRRTIQGGALQK
Zmarina-Zosma05g29470.1                      ------SAEVQLSETARERVTASSHWVMDSMNKGTDSYGVTTGFGATSHRRTKQGGALQK
Acomosus-Aco007727.1                         AKAVAPGVRVELAEAARGGVEASSEWVMESMARGTDSYGVTTGFGATSHRRTKQGGALQK
Bvulgarisssp-Bevul.7G054300.1                --TGPHEAAVELAESARAGVKASSDWVMDSMNKGTDSYGVTTGFGATSHRRTKQGGALQK
Bdistachyon-Bradi3g47110.1                   -------AQVQLDESARERVKASSDWVMDSMANGVDSYGVTTGFGATSHRRTKEGGALQR
Bdistachyon-Bradi3g47120.1                   -------AQVQLDESARERVKASSDWVMDSMANGVDSYGVTTGFGATSHRRTKEGGALQR
Bdistachyon-Bradi3g48840.1                   -------AKVELDESARERIKASSDWVMNSMMNGTDSYGVTTGFGATSHRRTKEGGALQR
Bdistachyon-Bradi3g49280.1                   -------AKVELDESARERVKASSDWVMNSMANGVDSYGVTTGFGATSHRRTKEGGALQR
Hvulgare-HORVU6Hr1G058840.1                  -------VVVELDESARERVKASSDWVMNSMMNGTDSYGVTTGFGATSHRRTKEGGALQR
Taestivum-Traes_6BL_C96F626D9.1              ---------------------------MNSMMNGTDSYGVTTGFGATSHRRTKEGGALQR
Tintermedium-Thint.18G0192700.1              -------VVVELDESARARVKASSDWVMNSMMNGTDSYGVTTRFGATSHRRTKEGGALQR
Taestivum-Traes_6DL_04CD7E82C.1              -------VVVELDESARARVKASSDWVMNSMMNGTDSYGVTTGFGATSHRRTKQGGALQR
Bdistachyon-Bradi3g49270.1                   -------AKVELDESARERVKASSDWVMNSMANGVDSYGVTTGFGATSHRRTKEGGALQR
Ecoracana-ELECO.r07.2AG0138020.1             -------ARVELDESARARVKASSDWVMDSMAKGTDSYGVTTGFGATSHRRTKEGGALQR
Ecoracana-ELECO.r07.2BG0193670.1             -------ARVELDESARERVKASSDWVMNSMMNGTDSYGVTTGFGATSHRRTKEGGALQR
Osativa-LOC_Os02g41670.1                     -------ARVELDESARERVKASSDWVMNSMMNGTDSYGVTTGFGATSHRRTKEGGALQR
Osativa-LOC_Os02g41680.1                     -------ARVELDESARERVKASSDWVMNSMMNGTDSYGVTTGFGATSHRRTKEGGALQR
Pvaginatum-Pavag04G202900.1                  -------ARVELDESARERVKASSDWVMNSMMNGTDSYGVTTGFGATSHRRTKEGGALQR
```

```
Pvaginatum-PavagK321000.1                  -------ARVELDESARERVKASSDWVMNSMMNGTDSYGVTTGFGATSHRRTKEGGALQR
Pvaginatum-PavagK321300.1                  -------ARVELDESARERVKASSDWVMNSMMNGTDSYGVTTGFGATSHRRTKEGGALQR
Pvaginatum-PavagK321100.1                  -------ARVELDESARERVKASSDWVMNSMMNGTDSYGVTTGFGATSHRRTKEGGALQR
Pvaginatum-PavagK321200.1                  -------ARVELDESARERVKASSDWVMNSMMNGTDSYGVTTGFGATSHRRTKEGGALQR
Pvirgatum-Pavir.1KG386500.1                -------ARVELDESARDRVKASSDWVMNSMMNGTDSYGVTTGFGATSHRRTKEGGALQR
Pvirgatum-Pavir.1NG356700.1                -------ARVELDESARDRVKASSDWVMNSMMNGTDSYGVTTGFGATSHRRTKEGGALQR
Sviridis-Sevir.1G245166.1                  -------ARVELDESARERVKASSDWVMNSMMNGTDSYGVTTGFGATSHRRTKEGGALQR
Sviridis-Sevir.1G245232.1                  -------ARVELDESARERVKASSDWVMNSMMNGTDSYGVTTGFGATSHRRTKEGGALQR
Pvirgatum-Pavir.1NG356800.1                -------ARVELDESARDRVKASSDWVMNSMMNGTDSYGVTTGFGATSHRRTKEGGALQR
Sbicolor-Sobic.004G220500.1                -------ARVELDESARGRVKASSDWVMNSMMNGTDSYGVTTGFGATSHRRTKEGGALQR
Sbicolor-Sobic.004G220600.2                -------ARVELDESARGRVKASSDWVMNSMMNGTDSYGVTTGFGATSHRRTKEGGALQR
Sbicolor-Sobic.004G220700.1                -------ARVELDESARGRVKASSDWVMNSMMNGTDSYGVTTGFGATSHRRTKEGGALQR
Sviridis-Sevir.1G245300.2                  -------ARVELDESARERVKASSDWVMNSMMNGTDSYGVTTGFGATSHRRTKEGGALQR
Zmays-Zm00001d017276_T001                  -------ARVELDESARSRVKASSDWVMSSMMNGTDSYGVTTGFGATSHRRTKEGGALQR
Zmays-Zm00001d017279_T001                  -------ARVELDESARSRVKASSDWVMTSMMNGTDSYGVTTGFGATSHRRTKEGGALQR
Zmays-Zm00001d051166_T001                  -------ARVELDESARGRVKASSDWVRDSMMNGTDSYGVTTGFGATSHRRTKEGGALQR
Ecoracana-ELECO.r07.2AG0138030.1           -------ARVELDESARARVKASSDWVMDSMAKGTDSYGVTTGFGATSHRRTKEGGALQR
Ecoracana-ELECO.r07.2BG0193680.1           -------ARVELDESARARVKASSDWVMDSMAKGTDSYGVTTGFGATSHRRTKEGGALQR
Osativa-LOC_Os05g35290.1                   ----GEAAMVELDESARERVKASSDWVMNSMANGTDSYGVTTGFGATSHRRTKEGGALQR
Platifolius-Phala.02G210700.1              -------AKVELDESARGRVKASSDWVMNSMMNGTDSYGVTTGFGATSHRRTKEGAALQR
Ecoracana-ELECO.r07.4AG0310420.1           ------ACRVELDESARERVKASSDWVMSSMMNGTDSYGVTTGFGATSHRRTKEGGALQR
Ecoracana-ELECO.r07.4BG0341610.1           ------ACRVELDESARERVKASSDWVMNSMMNGTDSYGVTTGFGATSHRRTKEGGALQR
Bdistachyon-Bradi3g49260.1                 -------ARVELDESARGRVKESSDWVMNSMMNGTDSYGVTTGFGATSHRRTKEGGALQR
Hvulgare-HORVU0Hr1G016330.1                -------ARVALDESARGRVKESSDWVMTSMMNGTDSYGVTTGFGATSHRRTKEGGALQR
Tintermedium-Thint.17G0260200.1            -------ARVALDESARGRVKESSDWVMTSMMNGTDSYGVTTGFGATSHRRTKEGGALQR
Tintermedium-Thint.18G0192600.1            -------ARVALDESARGRVKESSDWVMTSMMNGTDSYGVTTGFGATSHRRTKEGGALQR
Tintermedium-Thint.17G0231000.1            -------VMVELDESARERVKASSDWVMNSMMNGTDSYGVTTGFGATSHRRTKEGGALQR
Ecoracana-ELECO.r07.2AG0138010.1           -------ARVELDESARERVKASSDWVMDSMMKGTDSYGVTTGFGATSHRRTKEGGALQR
Ecoracana-ELECO.r07.2BG0193660.1           -------ARVELDESARERVKASSDWVMDSMMKGTDSYGVTTGFGATSHRRTKEGGALQR
Osativa-LOC_Os02g41650.3                   -------ARVELDESARGRVKASSDWVMNSMMNGTDSYGVTTGFGATSHRRTKEGGALQR
Pvaginatum-Pavag04G202800.1                -------ARVELDESARERVKASSDWVMSSMMNGTDSYGVTTGFGATSHRRTKEGGALQR
Pvaginatum-PavagK320900.1                  -------ARVELDESARERVKASSDWVMSSMMNGTDSYGVTTGFGATSHRRTKEGGALQR
Pvirgatum-Pavir.1NG356400.1                -------ARVELDESARGRVKASSDWVMSSMMNGTDSYGVTTGFGATSHRRTKEGGALQR
Sviridis-Sevir.1G245100.1                  -------ARVELDESARERVKASSDWVMNSMMNGTDSYGVTTGFGATSHRRTKEGGALQR
Sbicolor-Sobic.004G220400.1                -------ARVELDESARERVKASSDWVMSSMMNGTDSYGVTTGFGATSHRRTKEGGALQR
Zmays-Zm00001d017275_T001                  -------ARVELDESARGRVKASSDWVMSSMMNGTDSYGVTTGFGATSHRRTKEGGALQR
Zmays-Zm00001d051163_T002                  -------ARVELDESARERVRASSDWVMGSMMNGTDSYGVTTGFGATSHRRTKEGGALQR
Pvirgatum-Pavir.1KG386300.1                -------ARVELDESARGRVKASSDWVMSSMMNGTDSYGVTTGFGATSHRRTKEGGALQR
Platifolius-Phala.02G210600.1              -------AQVELDESARGRVKASSDWVMNSMMNGIDSYGVTTGFGATSHRRTKEGGALQR
Streptochaeta-strangu_020767-RA            ------AAKVELDESARGRVKASSDWVMNSMTKGTDSYGVTTGFGATSHRRTKEGGALQR
Bdistachyon-Bradi5g15830.1                 -------ARVELDESARGRVKESSDWVMNSMMNGTDSYGVTTGFGATSHRRTKEGGALQR
Hvulgare-HORVU2Hr1G089440.4                -------TRVELDESARGRVKESSDWVMNSMMNGTDSYGVTTGFGATSHRRTKEGGALQR
Taestivum-Traes_2AL_0DDC60D6E.1            ------------------------------------------------------------
Tintermedium-Thint.06G0475900.1            -------TRVELDESARGRVKESSDWVMNSMMNGTDSYGVTTGFGATSHRRTKEGGALQR
Tintermedium-Thint.06G0476200.1            -------TRVELDESARGRVKESSDWVMNSMMNGTDSYGVTTGFGATSHRRTKEGGALQR
Taestivum-Traes_2AL_2763280FC.1            ------------------------------------------------------------
Taestivum-Traes_2DL_8C21AAAEF.1            ------------------------------------------------------------
Tintermedium-Thint.V0383500.1              -------TRVELDESARGRVKESSDWVMNSMMNGTDSYGVTTGFGATSHRRTKEGGALQR
Taestivum-Traes_2AL_9EC3226F7.1            -------TRVELDESARGRVKESSDWVMNSMMNGTDSYGVTTGFGATSHRRTKEGGALQR
Tintermedium-Thint.05G0273800.1            -------TRVELDESARGRVKESSDWVMNSMMNGTDSYGVTTGFGATSHRRTKEGGALQR
Tintermedium-Thint.V0383400.1              -------TRVELDESARGRVKESSDWVMNSMMNGTDSYGVTTGFGATSHRRTKEGGVLQR
Taestivum-Traes_2BL_13D5272D7.1            -------TRVELDESARSRVKESSDWVMNSMMNGTDSYGVTTGFGATSHRRTKEGGALQR
Tintermedium-Thint.06G0475700.1            -------TRVELDESARGRVKESSDWVMNSMMNGTDSYGVTTGFGATSHRRTKEGGALQR
Taestivum-Traes_2BL_C051606EA.2            -------TRVELDESARGRVKESSDWVMNSMMNGTDSYGVTTGFGATSHRRTKEGGALQR
Tintermedium-Thint.V0383600.1              -------TRVELDESARGRVKESSDWVMNSMMNGTDSYGVTTGFGATSHRRTKEGGALQR
Taestivum-Traes_2AL_9D78F85E2.1            -------TRVELDESARGRVKESSDWVMNSMMNGTDSYGVTTGFGATSHRRTKEGGALQR
Tintermedium-Thint.05G0274100.1            -------TRVELDESARGRVKESSDWVMNSMMNGTDSYGVTTGFGATSHRRTKEGGALQR
Tintermedium-Thint.06G0476300.1            -------TRVELDESARGRVKESSDWVMNSMMNGTDSYGVTTGFGATSHRRTKEGGALQR
Taestivum-Traes_2BL_5A50FDA1A.1            -------TRVELDESARGRVKESSDWVMNSMMNGTDSYGVTTGFGATSHRRTKEGGALQR
Tintermedium-Thint.05G0273900.1            ------------------------------------------------------------
Taestivum-Traes_2DL_30F8555D4.1            ------------------------------------------------------------
Taestivum-Traes_2DL_83168C1E0.1            ------------------------------------------------------------
Taestivum-Traes_2DL_B978B4E22.1            ------------------------------------------------------------
Taestivum-Traes_1AS_0B9295A68.1            -------TRVELDESARGRVKESSDWVMNSMANGTDSYGVTTGFGATSHRRTKEGGALQR
Taestivum-Traes_1BS_7068F56E6.1            -------TRVELDESARGRVKESSDWVMNSMANGTDSYGVTTGFGATSHRRTKEGGALQR
Taestivum-Traes_1BS_AEA817616.1            ------------------------------------------------------------
Taestivum-Traes_4AL_892C47ED5.1            -------TRVELDESARCRVKESSDWVMNSMANGTDSYGVTTGFGATSHRRTKEGGALQR
Tintermedium-Thint.03G0055900.1            -------TRVELDESARGRVKESSDWVMNSMANGTDSYGVTTGFGATSHRRTKEGGALQR
Tintermedium-Thint.01G0012900.1            -------TRVELDESARGRVKESSDWVMNSMANGTDSYGVTTGFGATSHRRTKEGGALQR
Tintermedium-Thint.02G0158500.1            -------TRVELDESARGRVKESSDWVMSSMANGTDSYGVTTGFGATSHRRTKEGGALQR
Tintermedium-Thint.03G0055700.1            -------TRVELDESAHGRVKESSDWVMNSMANGTDSYGVTTGFGATSHRRTKEGGALQR
Taestivum-Traes_1BS_C3AA04888.1            ------------------------------------------------------------
Taestivum-Traes_1DS_A8BD91E4A.1            -------TRVELDESARGRVKESSDWVMNSMANGTDSYGVTTGFGATSHRRTKEGGALQR
Taestivum-Traes_1BS_C26CCE0E9.1            -------TRVELDESARGRVKESSDWVMNSMANGTDSYGVTTGFGATSHRRTKEGGALQR
Taestivum-Traes_1DS_A21FCBA3D.1            -------TRVELDESARDRVKESSDWVMNSMANGTDSYGVTTGFGATSHRRTKEGGALQR
Tintermedium-Thint.01G0012200.1            -------TRVELDESARGRVKESSDWVMNSMANGTDSYGVTTGFGATSHRRTKEGGALQR
Tintermedium-Thint.02G0158600.1            -------TRVELDESARGRVKESSDWVMNSMANGTDSYGVTTGFGATSHRRTKEGGALQR
Tintermedium-Thint.01G0012700.1            -------TRVELDESARGRVKESSDWVMNSMANGTDSYGVTTGFGATSHRRTKEGGALQR
Taestivum-Traes_5BL_F50A47503.1            -------TRVELDESARGRVKESSDWVMNSMANGTDSYGVTTGFGATSHRRTKEGGALQR
Tintermedium-Thint.02G0158700.1            -------TRVKLDESARGRVKESSDWVMKSMANGTDSYGVTTGFGATSHRRTKEGGALQR
Taestivum-Traes_5BL_AB528184A.1            -------TRVELDESARGRVKESSDWVMNSMANGTDSYGVTTGFGATSHRRTKEGGALQR
Tintermedium-Thint.03G0055800.1            -------TRVELDESTRGRVKESSDWVMNSMANGTDSYGVTTGFGATSHRRTKEGGALQR
Taestivum-Traes_1AS_F9013A945.1            -------TRVELDESARGRVKESSDWVMNSMANGTDSYGVTTGFGATSHRRTKEGGALQR
Taestivum-Traes_1DS_43DAC7A7F.1            -------TRVELDESARGRVKESSDWVMNSMANGTDSYGVTTGFGATSHRRTKEGGALQR
Taestivum-Traes_1BS_BD86C90A7.1            -------TRVELDESARGRVKESSDWVMNSMANGTDSYGVTTGFGATSHRRTKEGGALQR
Tintermedium-Thint.02G0158900.1            -------TRVELDESARSRVKESSDWVMNSMANGTDSYGVTTGFGATSHRRTKEGGALQR
Tintermedium-Thint.05G0274000.1            ------------------------------------------------------------
Osativa-LOC_Os04g43800.1                   -------ARVELDESARGRVKASSDWVMNSMMNGTDSYGVTTGFGATSHRRTKEGGALQR
Pvaginatum-Pavag06G160300.1                -------ARVELDESARGRVKASSDWVMNSMMNGTDSYGVTTGFGATSHRRTKEGGALQR
Sbicolor-Sobic.006G148900.1                -------ARVELDESARGRVKASSDWVMNSMMNGTDSYGVTTGFGATSHRRTKEGGALQR
Pvirgatum-Pavir.7KG237800.1                -------ARVELDESARGRVKASSDWVMSSMMNGTDSYGVTTGFGATSHRRTKEGGALQR
Pvirgatum-Pavir.7NG355800.1                -------ARVELDESARGRVKASSDWVMNSMMNGTDSYGVTTGFGATSHRRTKEGGALQR
Sviridis-Sevir.7G178300.1                  -------ARVELDESARGRVKASSDWVMNSMMNGTDSYGVTTGFGATSHRRTKEGGALQR
Zmays-Zm00001d003015_T001                  -------ARVELDESARGRVKASSDWVMDSMMNGTDSYGVTTGFGATSHRRTKEGGALQR
Hvulgare-HORVU1Hr1G022060.1                -------ASVELDESARGRVKESSDWVMASMANGVDSYGVTTGFGATSHRRTKEGGALQR
Taestivum-Traes_1BS_723922D171.2           -------ARVELDESARGRVKESSDWVMASMANGTDSYGVTTGFGATSHRRTKEGGALQR
Taestivum-Traes_1DS_A171C7D59.1            -------ARVELDESARGRVKESSDWVMASMANGTDSYGVTTGFGATSHRRTKEGGALQR
Tintermedium-Thint.01G0111500.1            -------ARVELDESARGRVKESSDWVMASMANGTDSYGVTTGFGATSHRRTKEGGALQR
Tintermedium-Thint.03G0265300.1            -------ARVELDESARGRVKESSDWVMASMANGTDSYGVTTGFGATSHRRTKEGGALQR
Hvulgare-HORVU2Hr1G038140.1                -------ARVEL----------------------------------------KEGGALQR
Tintermedium-Thint.04G0171200.1            ------------------------------------------------------------
Taestivum-Traes_2AS_958327519.3            ------------------------------------------------------------
Taestivum-Traes_2DS_28CA50371.1            ------------------------------------------------------------
Tintermedium-Thint.05G0160800.1            -------ARVELDESARGRVKESSDWVMTSMANGTDSYGVTTGFGATSHRRTKEGGALQR
Tintermedium-Thint.06G0299000.1            -------ARVELDESARGRVKESSDWVMTSMANGTDSYGVTTGFGATSHRRTKEGGALQR
Taestivum-Traes_2DS_3791A8A36.1            ------------------------------------------------------------
Tintermedium-Thint.06G0342800.1            -------ARVELDESARSRVKESSDWVMTSMANGTDSYGVTTGFGATSHRRTKEGGALQR
Tintermedium-Thint.05G0150900.1            ------------------------------------------------------------
Tintermedium-Thint.04G0242300.1            -------ARVELDESARGRVKESSDWVMTSMANGTDSYGVTTGFGATSHRRTKEGGALQR
Tintermedium-Thint.06G0298700.1            -------ARVELDESARGRVKESSDWVMTSMANGTDSYGVTTGFGATSHRRTKEGGALQR
Taestivum-Traes_2BS_88CF42F2E.1            -------ARVELDESARGRVKESSDWVMTSMANGTDSYGVTTGFGATSHRRTKEGGALQR
Tintermedium-Thint.04G0241900.1            -------ARVELDE-------------------------------------TKEGGALQR
Platifolius-Phala.06G165300.1              ----AGAARVELDESARGRVKASSDWVMNSMMIGTDSYGVTTGFGATSHRRTKEGGALQR
Streptochaeta-strangu_019385-RA            ----AAAAKVELDESARGRVKASSDWVMNSMMNDTDSYGVTTGFGATSHRRTKQGGALQR
Streptochaeta-strangu_028944-RA            ---GAAAARVELDESARGHVKASSDWVMNSMVNGVDTYGITTGFGGTSYRRTKEGGSLQR
Acomosus-Aco010091.1                       -----ATVTVELSEAARAGVKASSEWVMESIDKGKDSTGITAGFGAASHRRTEQGAALQN
Zmarina-Zosma03g22300.1                    ---SSGSVGVVVSQENRVVVAASSDWIMESMNKGTDSYGVTTGFGATSHRRTKQGGALQA
Ecdeiocolea-Emoptg0003741_1G000600.1       ----AAKARVELDEAARGRVQASSDWVMSSMMNGTDSYGVTTGFGATSHRRTKEGGALQR
Jascendens-Joasc.05G060500.1               ----AAAAKVELDESSRGRVKASSDWVMSSMMNGTDSYGVTTGFGATSHRRTKEGGALQR
Ecdeiocolea-Emoptg0003741_1G000630.1       ---AAAAAKVELDESARARVKASSDWIVDSVANGGDIYGVTTGFGGTSHRRTRDGPALQV
Jascendens-Joasc.05G060400.1               ----TGVAKVELDESARGRVKASSDWIVNSVANGGDIYGVTTGFGGTSHRRTRDGPALQI
Bdistachyon-Bradi3g49250.2                 ----AAGVSVELDEEARPRVKASSEWILSCLAAGGDIYGVTTGFGGTSHRRTKDGPALQV
Hvulgare-HORVU6Hr1G058820.1                ----ASGVSVELDEEARPRVKASSEWILSCLAGGGDIYGVTTGFGGTSHRRTKDGPALQV
Taestivum-Traes_6DL_1AEA7B869.1            ----ASGVSVELDEEARPRVKASSEWILSCLAGGGDIYGVTTGFGGTSHRRTKDGPALQV
Tintermedium-Thint.18G0192300.1            ----ASGVSVELDEEARPRVKASSEWILSCLAGGGDIYGVTTGFGGTSHRRTKDGPALQV
Tintermedium-Thint.17G0230800.1            ----ASGVSVELDEEARPRVKASSEWILSCLAGGGDIYGVTTGFGGTSHRRTKDGPALQV
Tintermedium-Thint.17G0230900.1            ----ASGVSVELDEEARPRVKASSEWILSCLAGGGDIYGVTTGFGGTSHRRTKDGPALQV
Ecoracana-ELECO.r07.2AG0138000.1           ----ASGVTVELDEAARPRVKASSEWILDCIAHGGDIYGVTTGFGGTSHRRTKDGPALQV
Ecoracana-ELECO.r07.2BG0193650.1           ----ASGVTVELDEAARPRVKASSEWILDCIAHGGDIYGVTTGFGGTSHRRTKDGPALQV
Pvaginatum-Pavag04G202700.1                ----ASGVAVQLDEEARPRVKASSEWILDCIAHGGDIYGVTTGFGGTSHRRTKDGPALQV
```

```
Sviridis-Sevir.1G245000.1                    ----ASGVAVELDEEARPRVKASSEWILDCIAHGGDIYGVTTGFGGTSHRRTKDGPALQV
Sviridis-Sevir.6G187100.1                    ----ASGVAVELDEEARPRVKASSEWILDCIAHGGDIYGVTTGFGGTSHRRTKDGPALQV
Pvirgatum-Pavir.1NG356200.1                  ----ASGVAVELDEEARPRVKASSEWILDCIAHGGDIYGVTTGFGGTSHRRTKDGPALQV
Sbicolor-Sobic.004G220300.1                  ----ASGVAVELDEEARPRVKASSEWILDCIAHGGDIYGVTTGFGGTSHRRTKDGPALQV
Zmays-Zm00001d017274_T001                    ----ASGVAVELDEEARPRVKASSEWILDCIAHGGDIYGVTTGFGGTSHRRTKDGPALQV
Zmays-Zm00001d051161_T003                    ----ASGVAVELDEEARPRVKASSEWILDCIAHGGDIYGVTTGFGGTSHRRTKDGPALQV
Sviridis-Sevir.7G178200.1                    ----ASGVAVELDEDARPRVKASSEWILDCIAHGGDIYGVTTGFGGTSHRRTKDGPALQV
Osativa-LOC_Os02g41630.2                     ----AAGVAVELDEEARPRVKASSEWILNCIAHGGDIYGVTTGFGGTSHRRTKDGPALQV
Ecoracana-ELECO.r07.4AG0310430.1             ----ASGVAVELDEEARPRVKASSEWILSCIANGGDIYGVTTGFGGTSHRRTKDGPALQV
Ecoracana-ELECO.r07.4BG0341620.1             ----ASGVAVELDEEARPRVKASSEWILSCIANGGDIYGVTTGFGGTSHRRTKDGPALQV
Pvaginatum-Pavag06G160200.1                  ----ASGVAVELDEEARPRVKASSEWILDCIAHGGDIYGVTTGFGGTSHRRTKDGPALQV
Pvirgatum-Pavir.7KG238255.1                  ----ASGVAVELDEAARPRVKASSEWILDCIAHGGDIYGVTTGFGGTSHRRTKDGPALQV
Sbicolor-Sobic.006G148800.1                  ----ASGVAVELDEEARPRVKASSEWILDCIAHGGDIYGVTTGFGGTSHRRTKDGPALQV
Zmays-Zm00001d003016_T001                    ----ASGVAVELDEEARPRVKASSEWILDCIAHGGDIYGVTTGFGGTSHRRTKDGAALQV
Osativa-LOC_Os04g43760.1                     ----ASGVVVELDEEARPRVKASSEWILNCIAHGGDIYGVTTGFGGTSHRRTKDGQALQV
Platifolius-Phala.02G210400.1                ----AAGVAVELNESARARVKESSEWILNCVASGGDIYGVTTGFGGTSHRRTKDGPALQV
Platifolius-Phala.06G165100.1                ----ASGAAVELDEAARGRVKASSEWILNCVATGGDIYGVTTGFGGTSHRRTKDGPALQV
Streptochaeta-strangu_019386-RA              ----GAGACVELDESARGRVKASSEWILDCIAHGGDIYGVTTGFGGTSHRRTKDGPALQV
Streptochaeta-strangu_020769-RA              ----AAGARVELDEAARGRVKESSEWILNCIATGGDIYGVTTGFGGTSHRRTKDGPALQV
Sviridis-Sevir.2G448300.1                    ----ASGVAVELDEEARLRVKASSEWVLSCIENGGDIYGVTTGFGGNSHRRTKDGHALQV
Sviridis-Sevir.7G177900.1                    ----ASGVAVELDEEARLRVKASSEWVLSCIENGGDIYGVTTGFGGNSHRRTKDGHALQV

Aamericanus-Acora.01G232100.1                ELIRFLNAGIFGSSN-DSSN-TLPPPATRAAMLVRTNTLLQGYSGIRFE-----------
Aamericanus-Acora.10G174800.1                ELIRFLNAGIFGASN-DTGN-TLPPPATRAAMLVRINTLLQGYSGIRFE-----------
Acomosus-Aco013943.1                         QLIRFLNAGIFGSGR-ESGQ-TLPAAATRAAMLVRINTLLQGYSGIRFE-----------
Aofficinalis-evm.model.AsparagusV1_01.827    ------------------------------------------------------------
Typha_latifolia-Tylat.14G083800.1            ELIRFLNAGIFGSGH-DSGH-TLPPSATRAAMLVRINTLLQGYSGIRFE-----------
Aofficinalis-evm.model.AsparagusV1_04.71     ----MLST-----------------SATRAAMLVRINTLLQGYSGIRFE-----------
Aofficinalis-evm.model.AsparagusV1_04.93     ----MLST-----------------SATRAAMLVRINTLLQGYSGIRFE-----------
Aofficinalis-evm.model.AsparagusV1_10.1948   --------------------------MVVEEMEVEMRNGIEG-SGDRDSGVGDAGDSSTL
Typha_latifolia-Tylat.03G127700.1            ELIRFLNAGIFGSGP-ESGH-TLPPAATRAAMIVRINTLLQGYSGIRFE-----------
Spolyrhiza-Spipo11G0025500                   ELIRFLNAGIFGSGD-DSANTTLPQSATRAAMLVRINTLLQGYSGIRFE-----------
Spolyrhiza-Spipo1G0003500                    ELIRFLNAGIFGSGA-DSGNTTLPPAATRAAMLVRVNTLLQGYSGIRFE-----------
Phoenix_dactylifera-LOC103718264             ELIRFLNAGIFGGGV-ESGH-TLPVSATRAAMVVRINTLLQGYSGIRFE-----------
Macuminata-GSMUA_Achr1T09070_001             ELIRFLNAGIFGSGP-ESGN-TLPSSAAKAAMLVRVNTLLQGYSGIRFE-----------
Acomosus-Aco006987.1                         ELIRFLNAGIFGSGR-ESGQ-TLPASATRAAMLVRINTLLQGYSGIRFE-----------
Acomosus-Aco027752.1                         ELIRFLNAGIFGSGR-ESGQ-TLPASATRAAMLVRINTLLQGYSGIRFE-----------
Cscoparia-Csco27g03980.1                     ELIRFLNAGIFGANK-ESGC-TLPSSTTRAAMLVRINTLLQGYSGIRFE-----------
Cscoparia-Csco27g03990.1                     ELIRFLNAGIFGANK-ESGC-TLPSSATRAAMLVRINTLLQGYSGIRFE-----------
Macuminata-GSMUA_Achr11T16380_001            ELIRFLNVGIFGSGA-ESGH-TLPPSATRAAMLVRINTLLQGYSGIRFE-----------
Macuminata-GSMUA_Achr11T22840_001            ELIRFLNAGIFGSGT-ESAH-TLPTPAARAAMLVRVNTLLQGYSGIRFE-----------
Macuminata-GSMUA_Achr2T00240_001             ELIRFLNAGIFGSGE-ESRH-TLPASAARAAMLVRVNTLLQGYSGIRFE-----------
Macuminata-GSMUA_Achr5T18560_001             ELIRFLNAGIFGSGA-ESGH-TLPTPAARAAMLVRINTLLQGYSGIRFE-----------
Macuminata-GSMUA_Achr8T18960_001             ELIRFLNAGIFGSGW-ESGH-TLPPSAARAAMLVRINTLLQGYSGIRFE-----------
Macuminata-GSMUA_Achr5T03950_001             ELIRFLNAGIFGSGSDQSGH-TLPPPATRAAMLVRINTLLQGYSGIRFE-----------
Phoenix_dactylifera-LOC103703950             ELIRFLNAGIFGSGP-ESGH-TLPPYATRAAMLVRINTLLQGYSGIRFE-----------
Acoerulea-Aqcoe1G140100.1                    ELIRFLNAGVFGNGT-ESSH-TLPYSATRAAMLVRINTLLQGYSGIRFE-----------
Acoerulea-Aqcoe2G144500.1                    ELIRFLNAGVFGNGT-ETSH-TLPHSATRAAMLVRINTLLQGYSGIRFE-----------
Typha_latifolia-Tylat.05G053500.1            ELIRFLNSGIFGSGD-DLAN-TLPASTTRAAMAVRVNTLLQGYSGIRFE-----------
Zmarina-Zosma05g29470.1                      ELIRFLNAGIFGGSK-DTAN-TLPSATTRAAMLVRANTLLQGYSGIRFE-----------
Acomosus-Aco007727.1                         ELIRFLNAGIFGSGP-KSEQ-TLPASATRAAMLVRINTLLQGYSGIRFE-----------
Bvulgarisssp-Bevul.7G054300.1                ELIRFLNAGVFGNGT-ESSH-TLPHTATRAAMLVRINTLLQGYSGIRFE-----------
Bdistachyon-Bradi3g47110.1                   ELIRFLNAGAFGTGS-D-GH-VLPAGATRAAMLVRINTLLQGYSGIRFE-----------
Bdistachyon-Bradi3g47120.1                   ELIRFLNAGAFGTGS-D-GH-VLPAGATRAAMLVRINTLLQGYSGIRFE-----------
Bdistachyon-Bradi3g48840.1                   ELIRFLNAGAFGTGS-D-GH-VLPAGATRAAMLVRINTLLQGYSGIRFE-----------
Bdistachyon-Bradi3g49280.1                   ELIRFLNAGAFGTGS-D-GH-VLPAGATRAAMLVRINTLLQGYSGIRFE-----------
Hvulgare-HORVU6Hr1G058840.1                  ELIRFLNAGAFGTGS-D-AH-VLPAATTRAAMLVRINTLLQGYSGIRFE-----------
Taestivum-Traes_6BL_C96F626D9.1              ELIRFLNAGAFGTGS-D-AH-VLPAATTRAAMLVRINTLLQGYSGIRFE-----------
Tintermedium-Thint.18G0192700.1              ELIRFLNAGAFGTGS-D-AH-VLPAATTRAAMLVRINTLLQGYSGIRFE-----------
Taestivum-Traes_6DL_04CD7E82C.1              ELIRFLNAGAFGTGS-D-AH-VLPAATTRAAMLVRINTLLQGYSGIRFE-----------
Bdistachyon-Bradi3g49270.1                   ELIRFLNAGAFGTGS-D-GH-VLPAGATRAAMLVRINTLLQGYSGIRFE-----------
Ecoracana-ELECO.r07.2AG0138020.1             ELIRFLNAGAFGTGT-D-GH-VLPAEATRAAMLVRINTLLQGYSGIRFE-----------
Ecoracana-ELECO.r07.2BG0193670.1             ELIRFLNAGAFGTGT-D-GH-VLPAEATRAAMLVRINTLLQGYSGIRFE-----------
Osativa-LOC_Os02g41670.1                     ELIRFLNAGAFGTGT-D-GH-VLSAEATRAAMLVRINTLLQGYSGIRFE-----------
Osativa-LOC_Os02g41680.1                     ELIRFLNAGAFGTGT-D-GH-VLPAEATRAAMLVRINTLLQGYSGIRFE-----------
Pvaginatum-Pavag04G202900.1                  ELIRFLNAGAFGTGA-D-GH-VLPAEATRAAMLVRINTLLQGYSGIRFE-----------
Pvaginatum-PavagK321000.1                    ELIRFLNAGAFGTGA-D-GH-VLPAEATRAAMLVRINTLLQGYSGIRFE-----------
Pvaginatum-PavagK321300.1                    ELIRFLNAGAFGTGA-D-GH-VLPAEATRAAMLVRINTLLQGYSGIRFE-----------
Pvaginatum-PavagK321100.1                    ELIRFLNAGAFGTGA-D-GH-VLPAEATRAAMLVRINTLLQGYSGIRFE-----------
Pvaginatum-PavagK321200.1                    ELIRFLNAGAFGTGA-D-GH-VLPAEATRAAMLVRINTLLQGYSGIRFE-----------
Pvirgatum-Pavir.1KG386500.1                  ELIRFLNAGAFGTGT-D-GH-VLPAEATRAAMLVRINTLLQGYSGIRFE-----------
Pvirgatum-Pavir.1NG356700.1                  ELIRFLNAGAFGTGT-D-GH-VLPAEATRAAMLVRINTLLQGYSGIRFE-----------
Sviridis-Sevir.1G245166.1                    ELIRFLNAGAFGTGT-D-GH-VLPAEATRAAMLVRINTLLQGYSGIRFE-----------
Sviridis-Sevir.1G245232.1                    ELIRFLNAGAFGTGT-D-GH-VLPAEATRAAMLVRINTLLQGYSGIRFE-----------
Pvirgatum-Pavir.1NG356800.1                  ELIRFLNAGAFGTGT-D-GH-VLPAEATRAAMLVRINTLLQGYSGIRFE-----------
Sbicolor-Sobic.004G220500.1                  ELIRFLNAGAFGTGT-D-GH-VLPAEATRAAMLVRINTLLQGYSGIRFE-----------
Sbicolor-Sobic.004G220600.2                  ELIRFLNAGAFGTGT-D-GH-VLPAEATRAAMLVRINTLLQGYSGIRFE-----------
Sbicolor-Sobic.004G220700.1                  ELIRFLNAGAFGTGT-D-GH-VLPAEATRAAMLVRINTLLQGYSGIRFE-----------
Sviridis-Sevir.1G245300.2                    ELIRFLNAGAFGTGT-D-GH-VLPAEATRAAMLVRINTLLQGYSGIRFE-----------
Zmays-Zm00001d017276_T001                    ELIRFLNAGAFGTGT-D-GH-VLPAEATRAAMLVRINTLLQGYSGIRFE-----------
Zmays-Zm00001d017279_T001                    ELIRFLNAGAFGTGA-D-GH-VLPAETTRAAMLVRINTLLQGYSGIRFE-----------
Zmays-Zm00001d051166_T001                    ELIRFLNAGAFGIGT-DAGH-VLPAEATRAAMLVRINTLLQGYSGIRFE-----------
Ecoracana-ELECO.r07.2AG0138030.1             ELIRFLNAGAFGTGA-D-GH-VLPAEATRAAMLVRINTLLQGYSGIRFE-----------
Ecoracana-ELECO.r07.2BG0193680.1             ELIRFLNAGAFGTGA-D-GH-VLPAEATRAAMLVRINTLLQGYSGIRFE-----------
Osativa-LOC_Os05g35290.1                     ELIRFLNAGAFGTGA-D-GH-VLPAGATRAAMLVRINTLLQGYSGIRFE-----------
Platifolius-Phala.02G210700.1                ELIRFLNAGAFGTSS-D-GH-MLPAEATRAAMLVRINTLLQGYSGIRFE-----------
Ecoracana-ELECO.r07.4AG0310420.1             ELIRFLNAGAFGTGE-D-GH-VLPSSATRAAMLVRINTLLQGYSGIRFE-----------
Ecoracana-ELECO.r07.4BG0341610.1             ELIRFLNAGAFGTGS-D-GH-VLPSSATRAAMLVRINTLLQGYSGIRFE-----------
Bdistachyon-Bradi3g49260.1                   ELIRFLNAGAFGTGT-D-GH-VLPAEATRAAMLVRINTLLQGYSGIRFE-----------
Hvulgare-HORVU0Hr1G016330.1                  ELIRFLNAGAFGTGT-D-GH-VLPAEATRAAMLVRINTLLQGYSGIRFE-----------
Tintermedium-Thint.17G0260200.1              ELIRFLNAGAFGTGT-D-GH-VLPAEATRAAMLVRINTLLQGYSGIRFE-----------
Tintermedium-Thint.18G0192600.1              ELIRFLNAGAFGTGT-D-GH-VLPAEATRAAMLVRINTLLQGYSGIRFE-----------
Tintermedium-Thint.17G0231000.1              ELIRFLNAGAFGTGS-D-AH-VLPAATTRAAMLVRINTLLQGYSGIRFE-----------
Ecoracana-ELECO.r07.2AG0138010.1             ELIRFLNAGAFGTGN-D-GY-VLPAEATRAAMLVRINTLLQGYSGIRFE-----------
Ecoracana-ELECO.r07.2BG0193660.1             ELIRFLNAGAFGTGN-D-GY-VLPAESTRAAMLVRINTLLQGYSGIRFE-----------
Osativa-LOC_Os02g41650.3                     ELIRFLNAGAFGTGT-D-GH-VLPAEATRAAMLVRINTLLQGYSGIRFE-----------
Pvaginatum-Pavag04G202800.1                  ELIRFLNAGAFGTGA-D-GH-VLPAEATRAAMLVRINTLLQGYSGIRFE-----------
Pvaginatum-PavagK320900.1                    ELIRFLNAGAFGTGA-D-GH-VLPAEATRAAMLVRINTLLQGYSGIRFE-----------
Pvirgatum-Pavir.1NG356400.1                  ELIRFLNAGAFGTGA-D-GH-VLPAEATRAAMLVRINTLLQGYSGIRFE-----------
Sviridis-Sevir.1G245100.1                    ELIRFLNAGAFGTGA-D-GH-VLPAEATRAAMLVRINTLLQGYSGIRFE-----------
Sbicolor-Sobic.004G220400.1                  ELIRFLNAGAFGTGA-D-GH-VLPAEATRAAMLVRINTLLQGYSGIRFE-----------
Zmays-Zm00001d017275_T001                    ELIRFLNAGAFGTGA-D-GH-VLPAEATRAAMLVRINTLLQGYSGIRFE-----------
Zmays-Zm00001d051163_T002                    ELIRFLNAGAFGTGA-D-GH-VLPAEATRAAMLVRVNTLLQGYSGIRFE-----------
Pvirgatum-Pavir.1KG386300.1                  ELIRFLNAGAFGTGA-D-GH-VLPAEATRAAMLVRINTLLQGYSGIRFE-----------
Platifolius-Phala.02G210600.1                ELIRFLNAGAFGTGS-D-GH-VLPAEATRAAMLVRINTLLQGYSGIRFE-----------
Streptochaeta-strangu_020767-RA              ELIRFLNAGAFGTGT-D-GH-VLPAAATRAAMLVRINTLLQGYSGIRFE-----------
Bdistachyon-Bradi5g15830.1                   ELIRFLNAGAFGTGE-D-GH-VLPAAATRAAMLVRVNTLLQGYSGIRFE-----------
Hvulgare-HORVU2Hr1G089440.4                  ELIRFLNAGAFGTGN-D-GH-VLPAAATRAAMLVRVNTLLQGYSGIRFE-----------
Taestivum-Traes_2AL_0DDC60D6E.1              -------------------------------MLVRVNTLLQGYSGIRFE-----------
Tintermedium-Thint.06G0475900.1              ELIRFLNAGAFGTGT-D-GH-VLPAAATRAAMLVRVNTLLQGYSGIRFE-----------
Tintermedium-Thint.06G0476200.1              ELIRFLNAGAFGTGT-D-GH-VLPAAATRAAMLVRVNTLLQGYSGIRFE-----------
Taestivum-Traes_2AL_2763280FC.1              -------------------------------MLVRVNTLLQGYSGIRFE-----------
Taestivum-Traes_2DL_8C21AAAEF.1              -------------------------------MLVRVNTLLQGYSGIRFE-----------
Tintermedium-Thint.V0383500.1                ELIRFLNAGAFGTGT-D-GH-VLPAAATRAAMLVRVNTLLQGYSGIRFE-----------
Taestivum-Traes_2AL_9EC3226F7.1              ELIRFLNAGAFGTGT-D-GH-VLPAAATRAAMLVRVNTLLQGYSGIRFE-----------
Tintermedium-Thint.05G0273800.1              ELIRFLNAGAFGTGT-D-GH-VLPAAATRAAMLVRVNTLLQGYSGIRFE-----------
Tintermedium-Thint.V0383400.1                ELIRFLNAGAFGTGT-D-GH-VLPAAATRAAMLVRVNTLLQGYSGIRFE-----------
Taestivum-Traes_2BL_13D5272D7.1              ELIRFLNAGAFGTGT-D-GH-VLPAAATRAAMLVRVNTLLQGYSGIRFE-----------
Tintermedium-Thint.06G0475700.1              ELIRFLNAGAFGTGT-D-GH-VLPAAATRAAMLVRVNTLLQGYSGIRFE-----------
Taestivum-Traes_2BL_C051606EA.2              ELIRFLNAGAFGTGT-D-GH-VLPAAATRAAMLVRVNTLLQGYSGIRFE-----------
Tintermedium-Thint.V0383600.1                ELIRFLNAGAFGTGN-D-GH-VLPAAATRAAMLVRVNTLLQGYSGIRFE-----------
Taestivum-Traes_2AL_9D78F85E2.1              ELIRFLNAGAFGTGN-D-GH-VLPAAATRAAMLVRVNTLLQGYSGIRFE-----------
Tintermedium-Thint.05G0274100.1              ELIRFLNAGAFGNGN-D-GH-VLPAAATRAAMLVRVNTLLQGYSGIRFE-----------
Tintermedium-Thint.06G0476300.1              ELIRFLNAGAFGNGN-D-GH-VLPAAATRAAMLVRVNTLLQGYSGIRFE-----------
Taestivum-Traes_2BL_5A50FDA1A.1              ELIRFLNAGAFGNGS-D-GH-VLPAAATRAAMLVRVNTLLQGYSGIRFE-----------
Tintermedium-Thint.05G0273900.1              ------------------------------------------------------------
Taestivum-Traes_2DL_30F8555D4.1              ------------------------------------------------------------
Taestivum-Traes_2DL_83168C1E0.1              ------------------------------------------------------------
```

```
Taestivum-Traes_2DL_B978B4E22.1                     ------------------------------------------------------------
Taestivum-Traes_1AS_0B9295A68.1                     ELIRFLNAGAFGTGG-R-EH-VLPAAATRAAMLVRVNTLLQGYSGIRFE-----------
Taestivum-Traes_1BS_7068F56E6.1                     ELIRFLNAGAFGTGG-H-EH-VLPAAATRAAMLVRVNTLLQGYSGIRFE-----------
Taestivum-Traes_1BS_AEA817616.1                     ------------------------------------------------------------
Taestivum-Traes_4AL_892C47ED5.1                     ELIRFLNAGAFGTGG-H-DH-VLPAAATRAAMLVRVNTLLQGYSGIRFE-----------
Tintermedium-Thint.03G0055900.1                     ELIRFLNAGAFGTGG-H-EH-VLPAAATRAAMLVRVNTLLQGYSGIRFE-----------
Tintermedium-Thint.01G0012900.1                     ELIRFLNAGAFGTGG-H-DH-VLPAAATRAAMLVRVNTLLQGYSGIRFE-----------
Tintermedium-Thint.02G0158500.1                     ELIRFLNAGAFGNGG-H-DH-VLPAAATRAAMLVRVNTLLQGYSGIRFE-----------
Tintermedium-Thint.03G0055700.1                     ELIRFLNAGAFGNGG-H-EH-VLPAATTRAAMLVRVNTLLQGYSGIRFE-----------
Taestivum-Traes_1BS_C3AA04888.1                     ------------------------------------------------------------
Taestivum-Traes_1DS_A8BD91E4A.1                     ELIRFLNAGAFGNGG-H-EH-VLPAAATRAAMLVRVNTLLQGYSGIRFE-----------
Taestivum-Traes_1BS_C26CCE0E9.1                     ELIRFLNAGVFGTGG-H-DH-VLPAAATRAAMLVRVNTLLQGYSGIRFE-----------
Taestivum-Traes_1DS_A21FCBA3D.1                     ELIRFLNAGVFGTGG-H-DH-VLPAAATRAAMLVRVNTLLQGYSGIRFE-----------
Tintermedium-Thint.01G0012200.1                     ELIRFLNAGAFGTGG-H-DH-VLPAAATRAAMLVRVNTLLQGYSGIRFE-----------
Tintermedium-Thint.02G0158600.1                     ELIRFLNAGAFGTGG-H-DH-VLPAAATRAAMLVRVNTLLQGYSGIRFE-----------
Tintermedium-Thint.01G0012700.1                     ELIRFLNAGAFGTGG-H-DH-VLPAAATRAAMLVRVNTLLQGYSGIRFE-----------
Taestivum-Traes_5BL_F50A47503.1                     ELIRFLNAGAFGTGG-H-DH-VLPAAATRAAMLVRVNTLLQGYSGIRFE-----------
Tintermedium-Thint.02G0158700.1                     ELIRFLNAGAFGTGG-H-EH-VLPAAATRAAMLVRVNTLLQGYSGIRFE-----------
Taestivum-Traes_5BL_AB528184A.1                     ELIRFLNAGAFGTGG-H-DH-VLPAAATRAAMLVRVNTLLQGYSGIRFE-----------
Tintermedium-Thint.03G0055800.1                     ELIRFLNAGAFGTGG-H-DH-VLPTAATRAAMLVRVNTLLQGYSGIRFE-----------
Taestivum-Traes_1AS_F9013A945.1                     ELIRFLNAGAFGTGG-H-NH-VLPAATTRAAMLVRVNTLLQGYSGIRFE-----------
Taestivum-Traes_1DS_43DAC7A7F.1                     ELIRFLNAGAFGTGG-H-NH-VLPAATTRAAMLVRVNTLLQGYSGIRFE-----------
Taestivum-Traes_1BS_BD86C90A7.1                     ELIRFLNAGAFGTGG-H-DH-VLPAATTRAAMLVRVNTLLQGYSGIRFE-----------
Tintermedium-Thint.02G0158900.1                     ELIRFLNAGAFGTGG-H-DH-VLPAAATRAAMLVRVNTLLQGYSGIRFE-----------
Tintermedium-Thint.05G0274000.1                     -------------------------------MLVCVNTLLQGYSGIRFE-----------
Osativa-LOC_Os04g43800.1                            ELIRFLNAGAFGNGD-D-GH-VLPAAATRAAMLVRINTLLQGYSGIRFE-----------
Pvaginatum-Pavag06G160300.1                         ELIRFLNAGAFGTGD-D-GH-VLPAAATRAAMLVRINTLLQGYSGIRFE-----------
Sbicolor-Sobic.006G148900.1                         ELIRFLNAGAFGTGD-D-GH-VLPAAATRAAMLVRINTLLQGYSGIRFE-----------
Pvirgatum-Pavir.7KG237800.1                         ELIRFLNAGAFGTGD-D-GH-VLPASATRAAMLVRINTLLQGYSGIRFE-----------
Pvirgatum-Pavir.7NG355800.1                         ELVRFLNAGAFGTGD-D-GH-VLPASATRAAMLVRINTLLQGYSGIRFE-----------
Sviridis-Sevir.7G178300.1                           ELIRFLNAGAFGTGD-D-GH-VLPASATRAAMLVRINTLLQGYSGIRFE-----------
Zmays-Zm00001d003015_T001                           ELIRFLNAGAFGTGD-D-GH-VLPAAATRAAMLVRINTLLQGYSGIRFE-----------
Hvulgare-HORVU1Hr1G022060.1                         ELIRFLNAGAFGTGT-D-GH-VLPAATTRAAMLVRVNTLLQGYSGIRFE-----------
Taestivum-Traes_1BS_723922D171.2                    ELIRFLNAGAFGTGS-D-GH-VLPAATTRAAMLVRVNTLLQGYSGIRFE-----------
Taestivum-Traes_1DS_A171C7D59.1                     ELIRFLNAGAFGTGS-D-GH-VLPAATTRAAMLVRVNTLLQGYSGIRFE-----------
Tintermedium-Thint.01G0111500.1                     ELIRFLNAGAFGTGS-D-GH-VLPAATTRAAMLVRVNTLLQGYSGIRFE-----------
Tintermedium-Thint.03G0265300.1                     ELIRFLNAGAFGTGS-D-GH-VLPAATTRAAMLVRVNTLLQGYSGIRFE-----------
Hvulgare-HORVU2Hr1G038140.1                         ELIRFLNAGAFGTGT-D-GH-VLPAATTRAAMLVRVNTLLQGYSGIRFE-----------
Tintermedium-Thint.04G0171200.1                     -------------------------------MLVRVNTLLQGYSGIRFE-----------
Taestivum-Traes_2AS_958327519.3                     ------------------------------------------------------------
Taestivum-Traes_2DS_28CA50371.1                     ------------------------------------------------------------
Tintermedium-Thint.05G0160800.1                     ELIRFLNAGAFGTGS-D-GH-VLPTATTRAAMLVRVNTLLQGYSGIRFE-----------
Tintermedium-Thint.06G0299000.1                     ELIRFLNAGAFGTGS-D-GH-VLPAATTRAAMLVRVNTLLQGYSGIRFE-----------
Taestivum-Traes_2DS_3791A8A36.1                     ------------------------------------------------------------
Tintermedium-Thint.06G0342800.1                     ELIRFLNAGAFGTGS-D-GH-VLPAATTRAAMLVRVNW----YSGIRFE-----------
Tintermedium-Thint.05G0150900.1                     ------------------------------------------------------------
Tintermedium-Thint.04G0242300.1                     ELIRFLNAGAFGTGS-D-GH-VLPAATTRAAMLVRVNTLLQGYSGIRFE-----------
Tintermedium-Thint.06G0298700.1                     ELIRFLNAGAFGTGS-D-GH-VLPAATTRAAMLVRVNTLLQGYSGIRFE-----------
Taestivum-Traes_2BS_88CF42F2E.1                     ELIRFLNAGAFGTGS-D-GH-VLPAATTRAAMLVRVNTLLQGYSGIRFE-----------
Tintermedium-Thint.04G0241900.1                     ELIRFLNAGAFGTGS-D-GH-VLPAATTRAAMLVRVNTLLQGYSGIRFE-----------
Platifolius-Phala.06G165300.1                       ELIRFLNAGAFGTGA-D-GH-VLPAAATRAAMLVRINTLLQGYSGIRFE-----------
Streptochaeta-strangu_019385-RA                     ELIRFLNAGAFGTGE-D-GH-VLPTAATRAAMLVRINTLLQGYSGIRFE-----------
Streptochaeta-strangu_028944-RA                     ELIRFLNAGVFGTGA-D-GH-VLPAATTRAAMLVRINTLLQGYSGIRFE-----------
Acomosus-Aco010091.1                                ELIRFLNVGIFGDGL-ESGH-MLPRAATRAAMLVRINSLLQGYSGIRFE-----------
Zmarina-Zosma03g22300.1                             ELIRFLNAGIFSSGNSDSCN-MLPASTTRAAMLVRINSLLQGYSGIRFE-----------
Ecdeiocolea-Emoptg0003741_1G000600.1                ELIRFLNAGVFGEDS-D-GH-VLPAPATRAAMLVRINTLLQGYSGIRFE-----------
Jascendens-Joasc.05G060500.1                        ELIRFLNAGVFGTGS-D-GH-VLPAAATRAAMLVRINTLLQGYSGIRFE-----------
Ecdeiocolea-Emoptg0003741_1G000630.1                ELLRHLNAGIFGTGK-D-GH-VLPSEVTRAAMVVRINTLLQGYSGIRFE-----------
Jascendens-Joasc.05G060400.1                        ELLRHLNAGIFGTGT-D-GH-VLPSAVTRAAMFVRINTLLQGYSGIRFE-----------
Bdistachyon-Bradi3g49250.2                          ELLRHLNAGIFGTGS-D-GH-SLPAEVTRAAMLVRINTLLQGYSGIRFE-----------
Hvulgare-HORVU6Hr1G058820.1                         ELLRHLNAGIFGPGA-D-GH-SLPAEVSRAAMLVRINTLLQGYSGIRFE-----------
Taestivum-Traes_6DL_1AEA7B869.1                     ELLRHLNAGIFGPGA-D-GH-SLPAEVSRAAMLVRINTLLQGYSGIRFE-----------
Tintermedium-Thint.18G0192300.1                     ELLRHLNAGIFGPGA-D-GH-SLPAEVSRAAMLVRINTLLQGYSGIRFE-----------
Tintermedium-Thint.17G0230800.1                     ELLRHLNAGIFGPGA-D-GH-SLPAEVSRAAMLVRINTLLQGYSGIRFE-----------
Tintermedium-Thint.17G0230900.1                     ELLRHLNAGIFGPGA-D-GH-SLPAEVSRAAMLVRINTLLQGYSGIRFE-----------
Ecoracana-ELECO.r07.2AG0138000.1                    ELLRHLNAGIFGTGS-D-GH-TLPSETVRAAMLVRINTLLQGYSGIRFE-----------
Ecoracana-ELECO.r07.2BG0193650.1                    ELLRHLNAGIFGTGS-D-GH-TLPSETVRAAMLVRINTLLQGYSGIRFE-----------
Pvaginatum-Pavag04G202700.1                         ELLRHLNAGIFGTGG-D-GH-TLPSEVTRAAMLVRINTLLQGYSGIRFE-----------
Sviridis-Sevir.1G245000.1                           ELLRHLNAGIFGTGS-D-GH-TLPSEVTRAAMLVRINTLLQGYSGIRFE-----------
Sviridis-Sevir.6G187100.1                           ELLRHLNAGIFGAGS-D-GH-TLPSEVTRAAMLVRINTLLQGYSGIRFE-----------
Pvirgatum-Pavir.1NG356200.1                         ELLRHLNAGIFGNDR-D-GH-TLPSETVRAAMLVRINTLLQGYSGIRFE-----------
Sbicolor-Sobic.004G220300.1                         ELLRHLNAGIFGTGS-D-GH-TLPSEVVRAAMLVRINTLLQGYSGIRFE-----------
Zmays-Zm00001d017274_T001                           ELLRHLNAGIFGTGS-D-GH-TLPSEVTRAAMLVRINTLLQGYSGIRFE-----------
Zmays-Zm00001d051161_T003                           ELLRHLNAGIFGTGS-D-GH-TLPSEVTRAAMLVRINTLLQGYSGIRFE-----------
Sviridis-Sevir.7G178200.1                           ELLRHLNAGIFGNGS-D-GH-TLPSEVSRAAMLVRINTLLQGYSGIRFE-----------
Osativa-LOC_Os02g41630.2                            ELLRHLNAGIFGTGS-D-GH-TLPSETVRAAMLVRINTLLQGYSGIRFE-----------
Ecoracana-ELECO.r07.4AG0310430.1                    ELLRHLNAGIFGTGS-D-GH-TLPSEVSRAAMLVRINTLLQGYSGIRFE-----------
Ecoracana-ELECO.r07.4BG0341620.1                    ELLRHLNAGIFGTGS-D-GH-TLPSEVSRAAMLVRINTLLQGYSGIRFE-----------
Pvaginatum-Pavag06G160200.1                         ELLRHLNAGIFGDGS-D-GH-TLPSEVSRAAMLVRINTLLQGYSGIRFE-----------
Pvirgatum-Pavir.7KG238255.1                         ELLRHLNAGIFGNGS-D-GH-TLPSEVSRAAMLVRINTLLQGYSGIRFE-----------
Sbicolor-Sobic.006G148800.1                         ELLRHLNAGIFGNGS-D-GH-TLPSEVSRAAMLVRINTLLQGYSGIRFE-----------
Zmays-Zm00001d003016_T001                           ELLRHLNAGIFGNGS-D-GH-TLPSEVSRAAMLVRINTLLQGYSGIRFE-----------
Osativa-LOC_Os04g43760.1                            ELLRHLNAGIFGNGS-D-GN-SLPSEVSRAAMLVRINTLLQGYSGIRFE-----------
Platifolius-Phala.02G210400.1                       ELLRHLNAGIFGTGS-D-GH-TLPSEVTRAAMLVRINTLLQGYSGIRFE-----------
Platifolius-Phala.06G165100.1                       ELLRHLNAGIFGTGS-D-GH-TLPSEVTRAAMLVRINTLLQGYSGIRFE-----------
Streptochaeta-strangu_019386-RA                     ELLRHLNAGIFGTGS-H-GH-TLPTEVVRAAMLVRTNTLLQGYSGIRFE-----------
Streptochaeta-strangu_020769-RA                     ELLRHLNAGIFGTGS-D-GH-TLPSEVARAAMLVRINTLLQGYSGIRFE-----------
Sviridis-Sevir.2G448300.1                           ELLRYLNAGIFGTGS-D-GH-TLPSQVSRAAMLVRINALMQGYSGIRFE-----------
Sviridis-Sevir.7G177900.1                           ELLRYLNAGIFGTGS-D-GH-TLPSQVSRAAMLVRINALMQGYSGIRFE-----------


Aamericanus-Acora.01G232100.1                       ---------ILESITALLNTGVTPCLPLRGTITASGDLVPLSYIAGMLTGRPNAKAIGPD
Aamericanus-Acora.10G174800.1                       ---------ILESITAFLNTGITPCLPLRGTITASGDLVPLSYIAGMLTGRPNSKAIGPD
Acomosus-Aco013943.1                                ---------ILESITALLNSHVTPCLPLRGTITASGDLVPLSYIAGILTGRPNSVAVAPD
Aofficinalis-evm.model.AsparagusV1_01.827           ------------------------------------------------------------
Typha_latifolia-Tylat.14G083800.1                   ---------ILEAITSLLNNNITPCLPLRGTITASGDLVPLSYIAGMLTGRPNSKAISPD
Aofficinalis-evm.model.AsparagusV1_04.71            ---------ILEAITSFLNANITPCLPLRGTISSSGDLVPLSYIAGMLTGRPNSKAIGPN
Aofficinalis-evm.model.AsparagusV1_04.93            ---------ILEAITSFLNANITPCLPLRGTISSSGDLVPLSYIAGMLTGRPNSKAIGPN
Aofficinalis-evm.model.AsparagusV1_10.1948          ESSAPANNPTIEAITSFLNTNITPCLPLRGTISSSGDLVPLSYIAGMLTGRPNSKAISPD
Typha_latifolia-Tylat.03G127700.1                   ---------ILEAMTSLLNSNVTPCLPLRGTITASGDLVPLSYIAGLLTGRPNSIAKTID
Spolyrhiza-Spipo11G0025500                          ---------IMEAITALLNHHVTPCLPLRGTITASGDLVPLSYIAGMLTGRPNARAVNAA
Spolyrhiza-Spipo1G0003500                           ---------ILEAITALLNNGVTPCLPLRGTITASGDLVPLSYIAGMLTGRPNAKAVTAE
Phoenix_dactylifera-LOC103718264                    ---------MLEAITSLLNSNITPCLPLRGTITASGDLVPLSYIAGLLTGRPNSKATTHD
Macuminata-GSMUA_Achr1T09070_001                    ---------ILEAIASLLNNGITPCLPLRGTITASGDLVPLSYIAGILTGRPNAKAVGPD
Acomosus-Aco006987.1                                ---------ILEAITALLNSRVTPCLPLRGTITASGDLVPLSYIAGILTGRPNAKAVGPD
Acomosus-Aco027752.1                                ---------ILEAITALLNSRVTPCLPLRGTITASGDLVPLSYIAGILTGRPNAKAVGPD
Cscoparia-Csco27g03980.1                            ---------ILEAITSLLNSNITPCLPLRGTITASGDLVPLSYIAGLITGRPNSKAIGPD
Cscoparia-Csco27g03990.1                            ---------ILEAITSLLNSNITPCLPLRGTITASGDLVPLSYIAGLITGRPNSKAIGPD
Macuminata-GSMUA_Achr11T16380_001                   ---------ILEAMASLLNSGITPCLPLRGTITASGDLVPLSYIAGVLTGRPNAKACDPA
Macuminata-GSMUA_Achr11T22840_001                   ---------ILEAMASLLNSGITPCLPLRGTITASGDLVPLSYIAGVLTGRPNAKAFAPG
Macuminata-GSMUA_Achr2T00240_001                    ---------ILEAMASLLNCGITPCLPLRGTITASGDLVPLSYIAGLLTGRPNAKALDPA
Macuminata-GSMUA_Achr5T18560_001                    ---------ILEAMASLLNSGITPCLPLRGTITASGDLVPLSYIAGLLTGRGNAKALAPG
Macuminata-GSMUA_Achr8T18960_001                    ---------ILEAMASLLNSGITPCIPLRGTITASGDLLPLSYIAALLTGRPNAKAFCPG
Macuminata-GSMUA_Achr5T03950_001                    ---------ILEAITSLLNSNITPCLPLRGTITASGDLVPLSYIAGILTGRPNSKAISPA
Phoenix_dactylifera-LOC103703950                    ---------ILEAMSSLLNSNITPCVPLRGSITASGDLMPLSYIAGILSGRPNAVAIGPT
Acoerulea-Aqcoe1G140100.1                           ---------ILEAITKFLNNNITPCLPLRGTITASGDLVPLSYIAGLLTGRPNSKAVGPN
Acoerulea-Aqcoe2G144500.1                           ---------ILEAITKFLNNNITPCLPLRGTITASGDLVPLSYIAGLLTGRPNSKAVGPS
Typha_latifolia-Tylat.05G053500.1                   ---------ILEAITNLLNSNITPCLPLRGTITASGDLIPLSYIAGILTGRPNSKAIGPD
Zmarina-Zosma05g29470.1                             ---------ILEAMTKFLNGNITPCLPLRGTITASGDLVPLSYIAGFLTGRTNSKAIGPD
Acomosus-Aco007727.1                                ---------ILEAIAAFLNSHITPCLPLRGTITASGDLVPLSYIAGLLTGRPNSIAVGPD
Bvulgarisssp-Bevul.7G054300.1                       ---------ILEAITGLLNNNITPCLPLRGTITASGDLVPLSYIAGLLTGRPNSKAVGPN
Bdistachyon-Bradi3g47110.1                          ---------ILEAIAKLLNANVTPCLPLRGTITASGDLVPLSYIAGLVTGRENSVAVAPD
Bdistachyon-Bradi3g47120.1                          ---------ILEAIAKLLNANVTPCLPLRGTITASGDLVPLSYIAGLVTGRENSVAVAPD
Bdistachyon-Bradi3g48840.1                          ---------ILEAIAKLLNANVTPCLPLRGTITASGDLVPLSYIAGLVTGRENSVAVTPD
Bdistachyon-Bradi3g49280.1                          ---------ILEAIAKLLNANVTPCLPLRGTITASGDLVPLSYIAGLVTGRENSVAVTPD
Hvulgare-HORVU6Hr1G058840.1                         ---------ILEAVAKLLNANVTPCLPLRGTITASGDLVPLSYIAGLITGRDNSVAIGPD
Taestivum-Traes_6BL_C96F626D9.1                     ---------ILEAVAKLLNANVTPCLPLRGTITASGDLVPLSYIAGLVTGRENSVAIGPD
Tintermedium-Thint.18G0192700.1                     ---------ILEAVAKLLNANVTPCLPLRGTITASGDLVPLSYIAGLVTGRENSVAIGPD
```

| | |
|---|---|
| Taestivum-Traes_6DL_04CD7E82C.1 | ---------ILEAVAKLLNANVMPCLPLRGTITASGDLVPLSYIAGLITGRENSVAIGPD |
| Bdistachyon-Bradi3g49270.1 | ---------ILEAIAKLLNANVTPCLPLRGTITASGDLVPLSYIAGLVTGRENSVAVAPD |
| Ecoracana-ELECO.r07.2AG0138020.1 | ---------ILEAVVKLLNANVTPCLPLRGTVTASGDLVPLSYIAGLITGRQNSVAVAPD |
| Ecoracana-ELECO.r07.2BG0193670.1 | ---------ILEAVVKLLNANVTPCLPLRGTVTASGDLVPLSYIAGLITGRQNSVAVAPD |
| Osativa-LOC_Os02g41670.1 | ---------ILEAITKLLNANVTPCLPLRGTVTASGDLVPLSYIAGLVTGRENSVAVAPD |
| Osativa-LOC_Os02g41680.1 | ---------ILEAIAKLLNANVTPCLPLRGTITASGDLVPLSYIAGLVTGRENAVAVAPD |
| Pvaginatum-Pavag04G202900.1 | ---------ILEAIVKLLNANVTPCLPLRGTVTASGDLVPLSYIAGLVTGRENSVAVAPD |
| Pvaginatum-PavagK321000.1 | ---------ILEAIVKLLNANVTPCLPLRGTVTASGDLVPLSYIAGLVTGRENSVAVAPD |
| Pvaginatum-PavagK321300.1 | ---------ILEAIVKLLNANVTPCLPLRGTVTASGDLVPLSYIAGLVTGRENSVAVAPD |
| Pvaginatum-PavagK321100.1 | ---------ILEAIVKLLNANVTPCLPLRGTVTASGDLVPLSYIAGLVTGRENSVAVAPD |
| Pvaginatum-PavagK321200.1 | ---------ILEAIVKLLNANVTPCLPLRGTVTASGDLVPLSYIAGLVTGRENSVAVAPD |
| Pvirgatum-Pavir.1KG386500.1 | ---------ILEAIAKLLNANVTPCLPLRGTITASGDLVPLSYIAGLVTGRENSVAVAPD |
| Pvirgatum-Pavir.1NG356700.1 | ---------ILEAIVKLLNANVTPCLPLRGTVTASGDLVPLSYIAGLVTGRENSVAVAPD |
| Sviridis-Sevir.1G245166.1 | ---------ILEAIVKLLNANVTPCLPLRGTVTASGDLVPLSYIAGLVTGRENSVAVAPD |
| Sviridis-Sevir.1G245232.1 | ---------ILEAIVKLLNANVTPCLPLRGTVTASGDLVPLSYIAGLVTGRENSVAVAPD |
| Pvirgatum-Pavir.1NG356800.1 | ---------ILEAIVKLLNANVTPCLPLRGTVTASGDLVPLSYIAGLVTGRENSVAVAPD |
| Sbicolor-Sobic.004G220500.1 | ---------ILEAIVKLLNANVTPCLPLRGTVTASGDLVPLSYIAGLVTGRENSVAVAPD |
| Sbicolor-Sobic.004G220600.2 | ---------ILEAIVKLLNANVTPCLPLRGTVTASGDLVPLSYIAGLVTGRENSVAVAPD |
| Sbicolor-Sobic.004G220700.1 | ---------ILEAIVKLLNANVTPCLPLRGTVTASGDLVPLSYIAGLVTGRENSVAVAPD |
| Sviridis-Sevir.1G245300.2 | ---------ILEAIVKLLNANVTPCLPLRGTVTASGDLVPLSYIAGLVTGRENSVAVAPD |
| Zmays-Zm00001d017276_T001 | ---------ILEAIVKLLNANVTPCLPLRGTVTASGDLVPLSYIAGLVTGRENSVAVAPD |
| Zmays-Zm00001d017279_T001 | ---------ILEAIVKLLNANVTPCLPLRGTVTASGDLVPLSYIAGLVTGRENSVAVAPD |
| Zmays-Zm00001d051166_T001 | ---------ILEAIVKLLNANVTPCLPLRGTVTASGDLVPLSYIAGLVTGRENAVAVAPD |
| Ecoracana-ELECO.r07.2AG0138030.1 | ---------ILEAIAKLLNANVTPCLPLRGTITASGDLVPLSYIAGLVTGRPNSVAVAPD |
| Ecoracana-ELECO.r07.2BG0193680.1 | ---------ILEAIAKLLNANVTPCLPLRGTITASGDLVPLSYIAGLVTGRPNSVAVAPD |
| Osativa-LOC_Os05g35290.1 | ---------ILEAVAKLLNANVTPCLPLRGTITASGDLVPLSYIAGLVTGRENSVAVAPD |
| Platifolius-Phala.02G210700.1 | ---------ILEAIAKLLNANVTPCLPLRGTITASGDLVPLSYIAGLITGRQNAVAVTPD |
| Ecoracana-ELECO.r07.4AG0310420.1 | ---------ILETIAQLLNAHVTPCLPLRGTITASGDLVPLSYIAGLVTGRPNAVATTAD |
| Ecoracana-ELECO.r07.4BG0341610.1 | ---------ILETIAQLLNAHVTPCLPLRGTITASGDLVPLSYIAGLVTGRPNAMATTAD |
| Bdistachyon-Bradi3g49260.1 | ---------ILEAITKLLNANVTPCLPLRGTITASGDLVPLSYIAGLITGRQNSVAVAPD |
| Hvulgare-HORVU0Hr1G016330.1 | ---------ILEAITKLLNANVTPCLPLRGTITASGDLVPLSYIAGLITGRQNSVAVAPD |
| Tintermedium-Thint.17G0260200.1 | ---------ILEAITKLLNANVTPCLPLRGTITASGDLVPLSYIAGLITGRQNSVAVGPD |
| Tintermedium-Thint.18G0192600.1 | ---------ILEAITKLLNANVTPCLPLRGTITASGDLVPLSYIAGLITGRQNSVAVAPD |
| Tintermedium-Thint.17G0231000.1 | ---------ILEAITKLLNANVTPCLPLRGTITASGDLVPLSYIAGLITGRQNSVAVAPD |
| Ecoracana-ELECO.r07.2AG0138010.1 | ---------ILEAIAKLLNANVTPCLPLRGTITASGDLVPLSYIAGLITGRQNAAAVAPD |
| Ecoracana-ELECO.r07.2BG0193660.1 | ---------ILEAIAKLLNANVTPCLPLRGTITASGDLVPLSYIAGLITGRQNAAAVAPD |
| Osativa-LOC_Os02g41650.3 | ---------ILEAITKLLNANVTPCLPLRGTITASGDLVPLSYIAGLITGRQNSVAVAPD |
| Pvaginatum-Pavag04G202800.1 | ---------ILEAIAKLLNANVTPCLPLRGTITASGDLVPLSYIAGLITGRANAAAVAPD |
| Pvaginatum-PavagK320900.1 | ---------ILEAIAKLLNANVTPCLPLRGTITASGDLVPLSYIAGLITGRANAAAVAPD |
| Pvirgatum-Pavir.1NG356400.1 | ---------ILEAIAKLLNANVTPCLPLRGTITASGDLVPLSYIAGLITGRQNSVAVAPD |
| Sviridis-Sevir.1G245100.1 | ---------ILETIAKLLNANVTPCLPLRGTITASGDLVPLSYIAGLITGRQNSVAVAPD |
| Sbicolor-Sobic.004G220400.1 | ---------ILEAIAKLLNANVTPCLPLRGTITASGDLVPLSYIAGLITGRQNSVAVAPD |
| Zmays-Zm00001d017275_T001 | ---------ILEAIAKLLNANVTPCLPLRGTITASGDLVPLSYIAGLITGRQNSVAVGPD |
| Zmays-Zm00001d051163_T002 | ---------ILEAIAKLLNANVTPCLPLRGTITASGDLVPLSYIAGLITGRQNSVAVDPD |
| Pvirgatum-Pavir.1KG386300.1 | ---------ILEAIAKLLNANVTPCLPLRGTITASGDLVPLSYIAGLITGRQNSVAVAPD |
| Platifolius-Phala.02G210600.1 | ---------ILEAIAKLLNANVTPCLPLRGTITASGDLVPLSYIAGLITGRQNAVALAPD |
| Streptochaeta-strangu_020767-RA | ---------ILEAIAALLNVNVTPCLPLRGTITASGDLVPLSYIAGLVTGRQNSVAVTTD |
| Bdistachyon-Bradi5g15830.1 | ---------ILETIATLLNANVTPCLPLRGTITASGDLVPLSYIAGLVTGRPNSVATAPD |
| Hvulgare-HORVU2Hr1G089440.4 | ---------ILETVATLLNANVTPCLPLRGTITASGDLVPLSYIAGLVTGRPNSMATAPD |
| Taestivum-Traes_2AL_0DDC60D6E.1 | ---------ILETVATLLNANVTPCLPLRGTITASGDLVPLSYIAGLVTGRPNSMATAPD |
| Tintermedium-Thint.06G0475900.1 | ---------ILETVATLLNANVTPCLPLRGTITASGDLVPLSYIAGLVTGRPNSMATAPD |
| Tintermedium-Thint.06G0476200.1 | ---------ILETVATLLNANVTPCLPLRGTITASGDLVPLSYIAGLVTGRPNSMATAPD |
| Taestivum-Traes_2AL_2763280FC.1 | ---------ILETVATLLNANVTPCLPLRGTITASGDLVPLSYIAGLVTGRPNSMATAPD |
| Taestivum-Traes_2DL_8C21AAAEF.1 | ---------ILETVATLLNANVTPCLPLRGTITASGDLVPLSYIAGLVTGRPNSMATAPD |
| Tintermedium-Thint.V0383500.1 | ---------ILETVATLLNANVTPCLPLRGTITASGDLVPLSYIAGLVTGRPNSMATAPD |
| Taestivum-Traes_2AL_9EC3226F7.1 | ---------ILETIATLLNANVTPCLPLRGTITASGDLVPLSYIAGLVTGRPNSMATAPD |
| Tintermedium-Thint.05G0273800.1 | ---------ILETIATLLNANVTPCLPLRGTITASGDLVPLSYIAGLVTGRPNSMATAPD |
| Tintermedium-Thint.V0383400.1 | ---------ILETIATLLNANVTPCLPLRGTITASGDLVPLSYIAGLVTGRPNSMATAPD |
| Taestivum-Traes_2BL_13D5272D7.1 | ---------ILETIATLLNANVTPCLPLRGTITASGDLVPLSYIAGLVTGRPNSMATAPD |
| Tintermedium-Thint.06G0475700.1 | ---------ILETIATLLNANVTPCLPLRGTITASGDLVPLSYIAGLVTGRPNSMATAPD |
| Taestivum-Traes_2BL_C051606EA.2 | ---------ILETVATLLNANVTPCLPLRGTITASGDLVPLSYIAGLVTGRPNSMATAPD |
| Tintermedium-Thint.V0383600.1 | ---------ILETIATLLNANVTPCLPLRGTITASGDLVPLSYIAGLVTGRPNSMATAPD |
| Taestivum-Traes_2AL_9D78F85E2.1 | ---------ILETIATLLNANVTPCLPLRGTITASGDLVPLSYIAGLVTGRPNSMATAPD |
| Tintermedium-Thint.05G0274100.1 | ---------ILETIATLLNANVTPCLPLRGTITASGDLVPLSYIAGLVTGRPNSMATAPD |
| Tintermedium-Thint.06G0476300.1 | ---------ILETIATLLNANVTPCLPLRGTITASGDLVPLSYIAGLVTGRPNSMATAPD |
| Taestivum-Traes_2BL_5A50FDA1A.1 | ---------ILETIATLLNANVTPCLPLRGTITASGDLVPLSYIAGLVTGRPNSIATAPD |
| Tintermedium-Thint.05G0273900.1 | ------------------------------------------------------------ |
| Taestivum-Traes_2DL_30F8555D4.1 | ------------------------------------------------------------ |
| Taestivum-Traes_2DL_83168C1E0.1 | ------------------------------------------------------------ |
| Taestivum-Traes_2DL_B978B4E22.1 | ------------------------------------------------------------ |
| Taestivum-Traes_1AS_0B9295A68.1 | ---------ILETVAALLNANVTPCLPLRGTITASGDLVPLSYIAGLVTGRANCVATAPD |
| Taestivum-Traes_1BS_7068F56E6.1 | ---------ILETVAALLNANVTPCLPLRGTITASGDLVPLSYIAGLVTGRANSVATAPD |
| Taestivum-Traes_1BS_AEA817616.1 | ------------------------------------------------------------ |
| Taestivum-Traes_4AL_892C47ED5.1 | ---------ILETVAALLNANVTPCLPLRGTITASGDLVPLSYIAGLVTGRANSVATAPD |
| Tintermedium-Thint.03G0055900.1 | ---------ILETVAALLNANVTPCLPLRGTITASGDLVPLSYIAGLVTGRANSVATAPD |
| Tintermedium-Thint.01G0012900.1 | ---------ILKTVAALLNANVTPCLPLRGTITASGDLVPLSYIAGLVTGRANSVATAPD |
| Tintermedium-Thint.02G0158500.1 | ---------ILETVAALLNANVTPCLPLRGTITASGDLVPLSYIAGLVTGRANSVATAPD |
| Tintermedium-Thint.03G0055700.1 | ---------ILETVAALLNANVTPCLPLRGTITASGDLVPLSYIAGLVTGRANSVATAPD |
| Taestivum-Traes_1BS_C3AA04888.1 | ------------------------------------------------------------ |
| Taestivum-Traes_1DS_A8BD91E4A.1 | ---------ILETVAALLNANVTPCLPLRGTITASGDLVPLSYIAGLVTGRANSVATAPD |
| Taestivum-Traes_1BS_C26CCE0E9.1 | ---------ILETVAALLNANVTPCLPLRGTITASGDLVPLSYIAGLVTGRANSVATAPD |
| Taestivum-Traes_1DS_A21FCBA3D.1 | ---------ILETVAALLNANVTPCLPLRGTITASGDLVPLSYIAGLVTGRANSVATAPD |
| Tintermedium-Thint.01G0012200.1 | ---------ILETVAALLNANVTPCLPLRGTITASGDLVPLSYIAGLVTGRANSVATAPD |
| Tintermedium-Thint.02G0158600.1 | ---------ILETVAALLNANVTPCLPLRGTITASGDLVPLSYIAGLVTGRANSVATAPD |
| Tintermedium-Thint.01G0012700.1 | ---------ILETVAALLNANVTPCLPLRGTITASGDLVPLSYIAGLVTGRANSVATAPD |
| Taestivum-Traes_5BL_F50A47503.1 | ---------ILETVAALLNANVTPCLPLRGTITASGDLVPLSYIAGLVTGRANSVATAPD |
| Tintermedium-Thint.02G0158700.1 | ---------ILETVAALLNANVTPCLPLRGTITASGDLVPLSYITGLVTGRANSVATAPD |
| Taestivum-Traes_5BL_AB528184A.1 | ---------ILETVAALLNANVTPCLPLRGTITASGDLVPLSYIAGLVTGRANSVATAPD |
| Tintermedium-Thint.03G0055800.1 | ---------ILETVAALLNANVTPCLPLRGTITASGDLVPLSYIAGLVTGRANSVATAPD |
| Taestivum-Traes_1AS_F9013A945.1 | ---------ILETIAKLLNTNVTPCLPLRGTITASGDLVPLSYIAGLVTGRANSVATAPD |
| Taestivum-Traes_1DS_43DAC7A7F.1 | ---------ILETIAKLLNANVTPCLPLRGTITASGDLVPLSYIAGLVTGRANSVATAPD |
| Taestivum-Traes_1BS_BD86C90A7.1 | ---------ILETISKLLNADVTPCLPLRGTITASGDLVPLSYIAGLVTGRANSVATALD |
| Tintermedium-Thint.02G0158900.1 | ---------ILETIATLLNANVTPCLPLRGTITASGDLVPLSYIAGLVTGRANSVATAPD |
| Tintermedium-Thint.05G0274000.1 | ---------ILETIATLLNANVTPCLPLRGTITASGDLVPLSYIVGLVTGRPNSMATAPD |
| Osativa-LOC_Os04g43800.1 | ---------ILETIATLLNANVTPCLPLRGTITASGDLVPLSYIAGLVTGRPNSVAVTPD |
| Pvaginatum-Pavag06G160300.1 | ---------ILETIAALLNANVTPCLPLRGTITASGDLVPLSYIAGLVTGRPNSTAVAPD |
| Sbicolor-Sobic.006G148900.1 | ---------ILETIAALLNANVTPCLPLRGTITASGDLVPLSYIAGLVTGRPNSTAVAPD |
| Pvirgatum-Pavir.7KG237800.1 | ---------ILEAIAALLNANVTPCLPLRGTITASGDLVPLSYIAGLVTGRPNSVAVTPD |
| Pvirgatum-Pavir.7NG355800.1 | ---------ILETIAALLNANVTPCLPLRGTITASGDLVPLSYIAGLVTGRPNSVAVTPD |
| Sviridis-Sevir.7G178300.1 | ---------ILETIASLLNANVTPCLPLRGTITASGDLVPLSYIAGLVTGRPNSVAVTAD |
| Zmays-Zm00001d003015_T001 | ---------ILEAIAALLNTNVTPCLPLRGTITASGDLVPLSYIAGLVTGRPNSTAVAPD |
| Hvulgare-HORVU1Hr1G022060.1 | ---------ILETIATLLNANVTPCLPLRGTITASGDLVPLSYIAGLVTGRPNAVAVAPD |
| Taestivum-Traes_1BS_723922D171.2 | ---------ILETIATLLNANVTPCLPLRGTITASGDLVPLSYIAGLVTGRPNAVAVAPD |
| Taestivum-Traes_1DS_A171C7D59.1 | ---------ILETIATLLNANVTPCLPLRGTITASGDLVPLSYIAGLVTGRPNAVAVAPD |
| Tintermedium-Thint.01G0111500.1 | ---------ILETIATLLNANVTPCLPLRGTITASGDLVPLSYIAGLVTGRPNAVAVAPD |
| Tintermedium-Thint.03G0265300.1 | ---------ILETIATLLNANVTPCLPLRGTITASGDLVPLSYIAGLVTGRPNAVAVAPD |
| Hvulgare-HORVU2Hr1G038140.1 | ---------ILETIATLLNANVTPCLPLRGTITASGDLVPLSYIAGLVTGHPNAVAVAPD |
| Tintermedium-Thint.04G0171200.1 | ---------ILETIATLLNAKVTPCLPLRGTITASGDLVPLSYIAGLVTGRPNAVAVAPD |
| Taestivum-Traes_2AS_958327519.3 | ------------------------------------------------------------ |
| Taestivum-Traes_2DS_28CA50371.1 | ------------------------------------------------------------ |
| Tintermedium-Thint.05G0160800.1 | ---------ILETIATLLNANVTPCLPLRGTITASGDLVPLSYIAGLVTGRPNAVAVAPD |
| Tintermedium-Thint.06G0299000.1 | ---------ILETIATLLNANVTPCLPLRGTITASGDLVPLSYIAGLVTGRPNAVAVAPD |
| Taestivum-Traes_2DS_3791A8A36.1 | ------------------------------------------------------------ |
| Tintermedium-Thint.06G0342800.1 | ---------ILETIATLLNANVTPCLPLRGTITASGDLVPLSYIAGLVTGRPNAVAVAPD |
| Tintermedium-Thint.05G0150900.1 | ------------------------------------------------------------ |
| Tintermedium-Thint.04G0242300.1 | ---------ILETIATLLNANVTPCLPLRGTITASGDLVPLSYIAGLVTGRPNAVAVAPD |
| Tintermedium-Thint.06G0298700.1 | ---------ILETIAMLLNANVTPCLPLRGTITASGDLVPLSYIAGLVTGRPNAVAVAPD |
| Taestivum-Traes_2BS_88CF42F2E.1 | ---------ILETIATLLNANVTPCLPLRGTITASGDLVPLSYIAGLVTGRPNAVAVAPD |
| Tintermedium-Thint.04G0241900.1 | ---------ILETIATLLNANVTPCLPLRGTITASGDLVPLSYIAGLVTGRPNAVAVAPD |
| Platifolius-Phala.06G165300.1 | ---------ILEAIAALLNANVTPCLPLRGTITASGDLVPLSYIAGLVTGRANSVAVAPD |
| Streptochaeta-strangu_019385-RA | ---------ILEAVAALLNANVTPCVPLRGTITASGDLVPLSYIAGLVTGRPNSVAVAPD |
| Streptochaeta-strangu_028944-RA | ---------ILEAIAALLNANVTPRLPLRGTLTASGDVIPLSYIAGLITGRQNSVAAAPD |
| Acomosus-Aco010091.1 | ---------ILEAIASLLNANVTPCVPLRGSITASGDLVPLAYIAGLLSGRPNSKAIGPD |
| Zmarina-Zosma03g22300.1 | ---------ILEALTDFLNHNITPCLPLRGSITASGDLIPLSYIVGLLIGRPNSKAIAPD |
| Ecdeiocolea-Emoptg0003741_1G000600.1 | ---------ILEAITALLNAGVTPCLPLRGTITASGDLVPLSYIAGLVTGRPNSVAVTPD |
| Jascendens-Joasc.05G060500.1 | ---------ILEAITALLNAGVTPCLPLRGTITASGDLVPLSYIAGLITGRPNSVAVAPD |
| Ecdeiocolea-Emoptg0003741_1G000630.1 | ---------ILEAITALLNSGVTPCLPLRGTITASGDLVPLSYIAGLITGRPNAMAVTPD |
| Jascendens-Joasc.05G060400.1 | ---------ILEAIAALLNAGVTPCLPLRGTITASGDLIPLSYIAGLITGRPNAVAIAPD |
| Bdistachyon-Bradi3g49250.2 | ---------ILEAITKLLNTGVSPCLPLRGTITASGDLVPLSYIAGLITGRPNAQATTAD |
| Hvulgare-HORVU6Hr1G058820.1 | ---------ILEAITKLINTGVSPCLPLRGTITASGDLVPLSYIAGLITGRPNAQATTVD |

```
Taestivum-Traes_6DL_1AEA7B869.1                    ---------ILEAITKLINTGVSPCLPLRGTITASGDLVPLSYIAGLITGRPNAQATTVD
Tintermedium-Thint.18G0192300.1                    ---------ILEAITKLINTGVSPCLPLRGTITASGDLVPLSYIAGLITGRPNAQATTVD
Tintermedium-Thint.17G0230800.1                    ---------ILEAITKLINTGVSPCLPLRGTITASGDLVPLSYIAGLITGRPNAQATTVD
Tintermedium-Thint.17G0230900.1                    ---------ILEAITKLINTGVSPCLPLRGTITASGDLVPLSYIAGLITGRPNAQATTVD
Ecoracana-ELECO.r07.2AG0138000.1                   ---------ILEAITKLINTGVSPCLPLRGTITASGDLVPLSYIAGLITGRPNAQATTVD
Ecoracana-ELECO.r07.2BG0193650.1                   ---------ILEAITKLINTGVSPCLPLRGTITASGDLVPLSYIAGLITGRPNAQATTVD
Pvaginatum-Pavag04G202700.1                        ---------ILEAITKLLNTGVSPCLPLRGTITASGDLVPLSYIAGLITGRPNAQAVTVD
Sviridis-Sevir.1G245000.1                          ---------ILEAITKLLNSGVSPCLPLRGTITASGDLVPLSYIAGLITGRPNAQAVTAD
Sviridis-Sevir.6G187100.1                          ---------ILEAITKLLNSGVSPCLPLRGTITASGDLVPLSYIAGLITGRPNAQAVTVD
Pvirgatum-Pavir.1NG356200.1                        ---------ILEALAKLLNTGVSPCLPLRGTITASGDLVPLSYIAGLITGRPNAQAVTAD
Sbicolor-Sobic.004G220300.1                        ---------ILEAITKLLNTGVSPCLPLRGTITASGDLVPLSYIAGLITGRPNAQATTVD
Zmays-Zm00001d017274_T001                          ---------ILEAITKLLNTGVSPCLPLRGTITASGDLVPLSYIAGLITGRPNAQAVTVD
Zmays-Zm00001d051161_T003                          ---------ILEAITKLLNTGVSPCLPLRGTITASGDLVPLSYIAGLITGRPNAQAVTVD
Sviridis-Sevir.7G178200.1                          ---------ILEAITKLINTGVSPCLPLRGTITASGDLVPLSYIAGLITGRPNAQAVTVD
Osativa-LOC_Os02g41630.2                           ---------ILEAITKLLNTGVTPCLPLRGTITASGDLVPLSYIAGLITGRPNAQAISPD
Ecoracana-ELECO.r07.4AG0310430.1                   ---------ILEAITKLINRGVSPCLPLRGTITASGDLVPLSYIAGLITGRPNAQAVTVD
Ecoracana-ELECO.r07.4BG0341620.1                   ---------ILEAITKLINTGVSPCLPLRGTITASGDLVPLSYIAGLITGRPNAQAVTVD
Pvaginatum-Pavag06G160200.1                        ---------ILEAITKLINTGVSPCLPLRGTITASGDLVPLSYIAGLITGRPNAQAVTVD
Pvirgatum-Pavir.7KG238255.1                        ---------ILEALAKLINTGVSPCLPLRGTITASGDLVPLSYIAGLITGRPNAQAVTVD
Sbicolor-Sobic.006G148800.1                        ---------ILEAITKLLNTGVSPCLPLRGTITASGDLVPLSYIAGLITGRPNAQAVTVD
Zmays-Zm00001d003016_T001                          ---------ILEAITKLINTGVSPCLPLRGTITASGDLVPLSYIAGLITGRPNAQAVTVD
Osativa-LOC_Os04g43760.1                           ---------ILEAITKLINTGVSPCLPLRGTITASGDLVPLSYIAGLITGRPNAQAVTVD
Platifolius-Phala.02G210400.1                      ---------ILEAITKLINTGVTPCLPLRGTITASGDLVPLSYIAGVITGRPNALAVAPD
Platifolius-Phala.06G165100.1                      ---------ILEAITKLLNSGVTPCLPLRGTITASGDLVPLSYIAGVITGRPNAQAVAPD
Streptochaeta-strangu_019386-RA                    ---------ILEAITKLLNSGVTPCLPLRGTITASGDLVPLSYIAGVLTGRPNAEAVTVD
Streptochaeta-strangu_020769-RA                    ---------ILEAIAKLLNTGVTPCLPLRGTITASGDLVPLSYIAGVITGRPNALAIAPD
Sviridis-Sevir.2G448300.1                          ---------ILEAIAKLINTGVSPCLPLRGSITASGDLVPLSYIAGLITGRPNAQAVTVD
Sviridis-Sevir.7G177900.1                          ---------ILEAIAKLINTGVSPCLPLRGSITASGDLVPLSYIAGLITGRPNAQAVTVD
```

```
Aamericanus-Acora.01G232100.1                      GRT-IDATEAFRAAGIQTGFFELQPKEGLALVNGTAVGSGLASMVLYEA-NVLSLLAEVL
Aamericanus-Acora.10G174800.1                      GQT-IDATEAFRLSGIQTGFFELQPKEGLALVNGTAVGSGLASMVLYEA-NILSVLAEVI
Acomosus-Aco013943.1                               GRS-VDAAEAFRLAGIPHGFFELQPKEGLALVNGTAVGSGLASIVLFEA-NILAVLSEVM
Aofficinalis-evm.model.AsparagusV1_01.827          ------------------------------------------------------------
Typha_latifolia-Tylat.14G083800.1                  GKT-IGAAEAFELAGIHSGFFELQPKEGLALVNGTAVGSGLASIVLFET-NVLAVLSEVL
Aofficinalis-evm.model.AsparagusV1_04.71           GSE-IDAAEAFKLAGIPSGFFELQPKEGLALVNGTAVGSALASTVLFEA-NILAVLAEII
Aofficinalis-evm.model.AsparagusV1_04.93           GSE-IDAAEAFKLAGIPSGFFELQPKEGLALVNGTAVGSALASTVLFEA-NILAVLAEII
Aofficinalis-evm.model.AsparagusV1_10.1948         GTH-IDAAEAFKLAGIPSGFFELQPKEGLALVNGTAVGSGLASTVLFEA-NILAVLSEII
Typha_latifolia-Tylat.03G127700.1                  GRS-VDAAEAFRLAGIANGFFELQPKEGLALVNGTAVGSGLASMVLFEA-NILVVLAEVI
Spolyrhiza-Spipo11G0025500                         GEK-VSAQEAFQQAGISGGFFELQPKEGLALVNGTAVGSGLASVVLFEA-NILAVLAEVL
Spolyrhiza-Spipo1G0003500                          GKQ-VSAEEAFRLAGIAGGFFELQPKEGLALVNGTAVGSGLASMVLFEA-NILAVLAEVL
Phoenix_dactylifera-LOC103718264                   GTA-IDASEAFKLAGITGGFFDLQPKEGLALVNGTAVGSGLASMVLFEA-NILAVLSEVL
Macuminata-GSMUA_Achr1T09070_001                   GKV-IGAAEAFRLAGIADGFFELQPKEGLALVNGTAVGSGLASMVLFEA-NVLAVLSEVL
Acomosus-Aco006987.1                               GEAELSAAEAFEVAGIGGGFFELQPKEGLALVNGTAVGSGLASIVLFEA-NILAVLSEVL
Acomosus-Aco027752.1                               GEAELSAAEAFEVAGIGGGFFELQPKEGLALVNGTAVGSGLASIVLFEA-NVLAVLSEVL
Cscoparia-Csco27g03980.1                           GQV-MNATEAFQHAGITTGFFELQPKEGLALVNGTAVGSGLASMVLYEA-NILSVLAEVI
Cscoparia-Csco27g03990.1                           GQV-MDATEAFKHAGITTGFFELQPKEGLALVNGTAVGSGLASMVLYEA-NILSVLAEVI
Macuminata-GSMUA_Achr11T16380_001                  GET-MDAAEAFRRAGIPHEFFELQPKEGLALVNGTTVGSGLASVVLYEA-NVLALLAEVL
Macuminata-GSMUA_Achr11T22840_001                  GEA-VDAAEAFRRSGIPHGFFELQPKEGLALVNGTAVGSGLASVVLYEA-NVLAVLAEVL
Macuminata-GSMUA_Achr2T00240_001                   GEA-IDAAEAFRRAGIPHGFFELQPKEGLALVNGTAVGSGLASVVLYEA-NVLAVLAEVL
Macuminata-GSMUA_Achr5T18560_001                   GEW-VDAAEAFRRAGIPHGFFELQPKEGLAMVNGTAVGSGLASLVLYEV-NVLAVLAEAL
Macuminata-GSMUA_Achr8T18960_001                   GET-VDAAEAFRRAGIPRGFFELQPKEGLALVNGTAVGSGLASVVLYEA-NVLAVLAEVL
Macuminata-GSMUA_Achr5T03950_001                   GEA-IDAAEAFRLAGIPHGFFELQPKEGLALVNGTAVGSGLASMVLFDA-NILAVLSEVL
Phoenix_dactylifera-LOC103703950                   GQP-IDATEAFRLAGIPHSFFELQPKEGLALVNGTAVGSGLASVVLFEA-NVLALLSEVL
Acoerulea-Aqcoe1G140100.1                          GES-LNAEQAFNLAGISSGFFELQPKEGLALVNGTAVGSGLASMVLFDA-NILAVMSEVL
Acoerulea-Aqcoe2G144500.1                          GDS-FDAMEAFRRAGIDGGFFELQPKEGLALVNGTSVGSGLASMVLFDA-NILAVLAEVI
Typha_latifolia-Tylat.05G053500.1                  GSL-HTAAEAFELAGISGGFFELQPKEGLALVNGTAVGSGLAAIVLFEA-NVLAVLAEIL
Zmarina-Zosma05g29470.1                            GKQ-ITSSEAFAIAGIQEGFFQLQPKEGLALVNGTAVGSGLASMVLFDA-NILALSAEVL
Acomosus-Aco007727.1                               GRS-LDASEAFRLAGIMQGFFDLQPKEGLALVNGTAVGSGLASIVLYEA-NILAVLSEVL
Bvulgarisssp-Bevul.7G054300.1                      GEV-LNAEQAFKAAQISSGFFELQPKEGLALVNGTAVGSGMASIVLFET-NILAVLAEVI
Bdistachyon-Bradi3g47110.1                         GSK-VNAAEAFKIAGIHGGFFELQPKEGLAMVNGTAVGSGLASTVLFDA-NVLAVMAEVI
Bdistachyon-Bradi3g47120.1                         GSK-VNAAEAFKIAGIHGGFFELQPKEGLAMVNGTAVGSGLASTVLFDA-NVLAVMAEVI
Bdistachyon-Bradi3g48840.1                         GRK-VNAAEAFKLAGIHGGFFELQPKEGLAMVNGTAVGSGLASTVLFEA-NILAVLAEVI
Bdistachyon-Bradi3g49280.1                         GRK-VNAAEAFKIAGIHGGFFELQPKEGLAMVNGTAVGSGLASTVLFEA-NILAVLAEVI
Hvulgare-HORVU6Hr1G058840.1                        GTK-VNAAEAFKIAGIHGGFFELQPKEGLAMVNGTAVGSALASTVLFEA-NILALLAEVI
Taestivum-Traes_6BL_C96F626D9.1                    GTK-VNAAEAFKIAGIHGGFFELQPKEGLAMVNGTAVGSALASTVLFEA-NILALLAEVI
Tintermedium-Thint.18G0192700.1                    GTK-VNAAEAFKIAGIHGGFFELQPKEGLAMVNGTAVGSALASTVLFEA-NILALLAEVI
Taestivum-Traes_6DL_04CD7E82C.1                    GTK-VNAAEAFKIAGIHGGFFELQPKEGLAMVNGTAVGSALASTVLFEA-NILALLAEVI
Bdistachyon-Bradi3g49270.1                         GRK-VNAAEAFKIAGIHGGFFELQPKEGLAMVNGTAVGSGLASTVLFEA-NILAVLAEVI
Ecoracana-ELECO.r07.2AG0138020.1                   GSK-VNAAEAFKIAGIQGGFFELQPKEGLAMVNGTAVGSGLASTVLFET-NVLAILAEVL
Ecoracana-ELECO.r07.2BG0193670.1                   GSK-VNAAEAFKIAGIQGGFFELQPKEGLAMVNGTVVGSGLASTVLFEA-NILAILAEVL
Osativa-LOC_Os02g41670.1                           GSK-VNAAEAFKIAGIQGGFFELQPKEGLAMVNGTAVGSGLASTVLFEA-NILAVLAEVL
Osativa-LOC_Os02g41680.1                           GSK-VNAAEAFKIAGIQGGFFELQPKEGLAMVNGTAVGSGLASTVLFEA-NILAILAEVL
Pvaginatum-Pavag04G202900.1                        GTK-VNAAEAFKIAGIQGGFFELQPKEGLAMVNGTAVGSGLASTVLFEA-NILAVLAEVL
Pvaginatum-PavagK321000.1                          GTK-VNAAEAFKIAGIQGGFFELQPKEGLAMVNGTAVGSGLASTVLFEA-NILAVLAEVL
Pvaginatum-PavagK321300.1                          GTK-VNAAEAFKIAGIQGGFFELQPKEGLAMVNGTAVGSGLASTVLFEA-NILAVLTEVL
Pvaginatum-PavagK321100.1                          GTK-VNAAEAFKIAGIQGGFFELQPKEGLAMVNGTAVGSGLASTVLFEA-NILAVLAEVL
Pvaginatum-PavagK321200.1                          GTK-VNAAEAFKIAGIQGGFFELQPKEGLAMVNGTAVGSGLASTVLFEA-NILAVLAEVL
Pvirgatum-Pavir.1KG386500.1                        GSK-VNAAEAFKIAGIQGGFFELQPKEGLAMVNGTAVGSGLASTVLFEA-NILAILAEVL
Pvirgatum-Pavir.1NG356700.1                        GTK-VNAAEAFKIAGIQGGFFELQPKEGLAMVNGTAVGSGLASTVLFEA-NILAILAEVL
Sviridis-Sevir.1G245166.1                          GTK-VNAAEAFKIAGIQGGFFELQPKEGLAMVNGTAVGSGLASTVLFEA-NVLAVLAEVL
Sviridis-Sevir.1G245232.1                          GTK-VNAAEAFKIAGIQGGFFELQPKEGLAMVNGTAVGSGLASTVLFEA-NVLAVLAEVL
Pvirgatum-Pavir.1NG356800.1                        GTK-VNAAEAFKIAGIQGGFFELQPKEGLAMVNGTAVGSGLASTVLFEA-NILAILAEVL
Sbicolor-Sobic.004G220500.1                        GTK-VNAAEAFKIAGIQGGFFELQPKEGLAMVNGTAVGSGLASTVLFEA-NVLAILAEVL
Sbicolor-Sobic.004G220600.2                        GTK-VNAAEAFKIAGIQGGFFELQPKEGLAMVNGTAVGSGLASTVLFEA-NVLAILAEVL
Sbicolor-Sobic.004G220700.1                        GTK-VNAAEAFKIAGIQGGFFELQPKEGLAMVNGTAVGSGLASTVLFEA-NVLAILAEVL
Sviridis-Sevir.1G245300.2                          GTK-VNAAEAFKIAGIQGGFFELQPKEGLAMVNGTAVGSGLASTVLFEA-NVLAVLAEVL
Zmays-Zm00001d017276_T001                          GSK-VNAAEAFKIAGIQGGFFELQPKEGLAMVNGTAVGSGLASTVLFEA-NILAILAEVL
Zmays-Zm00001d017279_T001                          GSK-VNAAEAFKIAGIQGGFFELQPKEGLAMVNGTAVGSGLASTVLFEA-NILAILAEVL
Zmays-Zm00001d051166_T001                          GTK-VNAAEAFRIADIQSGFFELQPKEGLAMVNGTAVGSGLASTVLFEA-NVLAVLAEVL
Ecoracana-ELECO.r07.2AG0138030.1                   GRK-VNAAEAFKIAGIEGGFFELQPKEGLAMVNGTAVGSGLASTVLFEA-NVLAILAEVL
Ecoracana-ELECO.r07.2BG0193680.1                   GRK-VNAAEAFKIAGIEGGFFELQPKEGLAMVNGTAVGSGLASTVLFEA-NVLAILAEVL
Osativa-LOC_Os05g35290.1                           GRK-VNAAEAFKIAGIQGGFFELQPKEGLAMVNGTAVGSGLASTVLFEA-NVLAILAEVL
Platifolius-Phala.02G210700.1                      GKK-VNAAEAFKIAGIQGGFFELQPKEGLAMVNGTAVGSGLASTVLFEA-NILAILAEVL
Ecoracana-ELECO.r07.4AG0310420.1                   GRK-VGAAEAFKIAGIERGFFELQPKEGLAMVNGTAVGSGLASTVLFEA-NILAILAEVL
Ecoracana-ELECO.r07.4BG0341610.1                   GRK-VGAAEAFKHAGIERGFFELQPKEGLAMVNGTAVGSGLASTVLFEA-NILAILAEVL
Bdistachyon-Bradi3g49260.1                         GSK-VSAAEAFKIAGIEHGFFELQPKEGLAMVNGTAVGSGLASTVLFEA-NIQAILAEVL
Hvulgare-HORVU0Hr1G016330.1                        GSK-VTAAEAFKIAGIEHGFFELQPKEGLAMVNGTAVGSGLASTVLFEA-NVLSVLAEVL
Tintermedium-Thint.17G0260200.1                    GSK-VTAAEAFKIAGIEHGFFELQPKEGLAMVNGTAVGSGLASTVLFEA-NVLSVLAEVL
Tintermedium-Thint.18G0192600.1                    GSK-VTAAEAFKIAGIEHGFFELQPKEGLAMVNGTAVGSGLASTVLFEA-NVLSVLAEVL
Tintermedium-Thint.17G0231000.1                    GSK-VTAAEAFKIAGIEHGFFELQPKEGLAMVNGTAVGSGLASTVLFEA-NVLSVLAEVL
Ecoracana-ELECO.r07.2AG0138010.1                   GRK-VNAAEAFKIAGIEHGFFELQPKEGLAMVNGTAVGSGLASTVLFEA-NVIAVLAEVM
Ecoracana-ELECO.r07.2BG0193660.1                   GRK-VNAAEAFKIAGIEHGFFELQPKEGLAMVNGTAVGSGLASTVLFEA-NVIAVLAEVM
Osativa-LOC_Os02g41650.3                           GRK-VTAAEAFKIAGIEHGFFELQPKEGLAMVNGTAVGSGLASTVLFEA-NVLAILAEVL
Pvaginatum-Pavag04G202800.1                        GTR-VGAAEAFKIAGIEHGFFELQPKEGLAMVNGTAVGSGLASTVLFEA-NVLAVLAEVL
Pvaginatum-PavagK320900.1                          GTR-VGAAEAFKIAGIEHGFFELQPKEGLAMVNGTAVGSGLASTVLFEA-NVLAVLAEVL
Pvirgatum-Pavir.1NG356400.1                        GRK-VDAAEAFKIAGIEHGFFELQPKEGLAMVNGTAVGSGLASTVLFEA-NVLAVMAEVI
Sviridis-Sevir.1G245100.1                          GRK-VDAAEAFKIAGIEHGFFELQPKEGLAMVNGTAVGSGLASTVLFEA-NVLAIMAEVI
Sbicolor-Sobic.004G220400.1                        GRK-VDAAEAFKIAGIEHGFFELQPKEGLAMVNGTAVGSGLASTVLFEA-NVLAIMAEVI
Zmays-Zm00001d017275_T001                          GRK-VGAAEAFRIAGIEHGFFELQPKEGLAMVNGTAVGSGLASTVLFEA-NVLAVMAEVI
Zmays-Zm00001d051163_T002                          GRK-VGAAEAFKIAGIEHGFFELQPKEGLAMVNGTAVGSGLASTVLFEA-NVLAVLAEVI
Pvirgatum-Pavir.1KG386300.1                        GRK-VDAAEAFKIAGIEHGFFELQPKEGLAMVNGTAVGSGLASTVLFEA-NVLAVMAEVI
Platifolius-Phala.02G210600.1                      GTK-VNAADAFKIAGIEHGFFELQPKEGLAMVNGTAVGSGLASTVLFEA-NILAILAEVL
Streptochaeta-strangu_020767-RA                    GRK-VNAAEAFKIAGIQHGFFELQPKEGLAMVNGTAVGSGLASTVLFEA-NILAILAEVL
Bdistachyon-Bradi5g15830.1                         GRK-VNAAEAFKIAGIQHGFFELQPKEGLAMVNGTAVGSGLASMVLFEA-NILGVLAEVL
Hvulgare-HORVU2Hr1G089440.4                        GTK-VNAAEAFKIAGIQHGFFELQPKEGLAMVNGTAVGSGLASMVLFEA-NILSLLAEVL
Taestivum-Traes_2AL_0DDC60D6E.1                    GTK-VNAAEAFKIAGIQHGFFELQPKEGLAMVNGTAVGSGLASMVLFEA-NILSLLAEVL
Tintermedium-Thint.06G0475900.1                    GTK-VNAVEAFKIAGIQHGFFELQPKEGLAMVNGTAVGSGLASMVLFEA-NILSLLAEVL
Tintermedium-Thint.06G0476200.1                    GTK-VNAAEAFKIAGIQHGFFELQPKEGLAMVNGTAVGSGLASMVLFEA-NILSLLAEVL
Taestivum-Traes_2AL_2763280FC.1                    GTK-VNAAEAFKIAGIQHGFFELQPKEGLAMVNGTAVGSGLASMVLFEA-NILSLLAEVL
Taestivum-Traes_2DL_8C21AAAEF.1                    GTK-VNAAEAFKIAGIQHGFFELQPKEGLAMVNGTAVGSGLASMVLFEA-NILSLLAEVL
Tintermedium-Thint.V0383500.1                      GTK-VNAAEAFKIAGIQHGFFELQPKEGLAMVNGTAVGSGLASMVLFEA-NILSLLAEVL
Taestivum-Traes_2AL_9EC3226F7.1                    GSK-VNAAEAFKIAGIQHGFFELQPKEGLAMVNGTAVGSGLASMVLFEA-NVLSLLAEVL
Tintermedium-Thint.05G0273800.1                    GTK-VNAAEAFKIAGIQHGFFELQPKEGLAMVNGTAVGSGLASMVLFEA-NVLSLLAEVL
Tintermedium-Thint.V0383400.1                      GTK-VNAAEAFKIAGIQHGFFELQPKEGLAMVNGTAVGSGLASMVLFEA-NVLSLLAEVL
Taestivum-Traes_2BL_13D5272D7.1                    GTK-VNAAKAFKIAGIQHGFFELQPKEGLAMVNGTAVGSGLASMVLFEA-NVLSLLAEVL
Tintermedium-Thint.06G0475700.1                    GTK-VNAAEAFKIAGIQHGFFELQPKEGLAMVNGTAVGSGLASMVLFEA-NVLSLLAEVL
Taestivum-Traes_2BL_C051606EA.2                    GTK-VNAAEAFKIAGIQHGFFELQPKEGLAMVNGTAVGSGLASMVLFEA-NILSLLVEVL
```

```
Tintermedium-Thint.V0383600.1                GTK-VNAAEAFKIAGIQHGFFELQPKEGLAMVNGTAVGSGLASMVLFEA-NILSLLAEVL
Taestivum-Traes_2AL_9D78F85E2.1              GTK-VNAAEAFKIAGIQHGFFELQPKEGLAMVNGTAVGSGLASMVLFEA-NILSLLAEVL
Tintermedium-Thint.05G0274100.1              GTK-VNAAEAFKIAGIQHGFFELQPKEGLAMVNGTAVGSGLASMVLFEA-NILSLLAEVL
Tintermedium-Thint.06G0476300.1              GTK-VNAAEAFKIAGIQHSFFELQPKEGLAMVNGTAVGSGLASMVLFEA-NILSLLAEVL
Taestivum-Traes_2BL_5A50FDA1A.1              GTK-VNAVEAFKIAGIQHGFFELQPKEGLAMVNGTAVGSGLASMVLFEA-NILSLLAEVL
Tintermedium-Thint.05G0273900.1              -------------------------------------------MVLFEA-NILSLLAEVL
Taestivum-Traes_2DL_30F8555D4.1              ------------------------------------------------------------
Taestivum-Traes_2DL_83168C1E0.1              ------------------------------------------------------------
Taestivum-Traes_2DL_B978B4E22.1              ------------------------------------------------------------
Taestivum-Traes_1AS_0B9295A68.1              GSK-VNAAEAFKIAGIQHGFFELQPKEGLAMVNGTAVGSGLASMVLFEA-NILSLLAEVL
Taestivum-Traes_1BS_7068F56E6.1              GSK-VNAAEAFKIAGIQHGFFELQPKEGLAMVNGTAVGSGLASMVLFEA-NILSLLAEVL
Taestivum-Traes_1BS_AEA817616.1              -------------------------------------------MVLFEA-NILSLLAEVL
Taestivum-Traes_4AL_892C47ED5.1              GSK-VNAAEAFKIAGIQHGFFELQPKEGLAMVNGTAVGSGLASMVLFEA-NILSLLAEVL
Tintermedium-Thint.03G0055900.1              GSK-VNAAEAFKIAGIQHGFFELQPKEGLAMVNGTAVGSGLASMVLFEA-NILSLLAEVL
Tintermedium-Thint.01G0012900.1              GSK-VNAAEAFKIAGIQHGFFELQPKEGLAMVNGTAVGSGLASMVLFEA-NILSLLAEIL
Tintermedium-Thint.02G0158500.1              GSK-VNAAEAFKIAGIQHGFFELQPKEGLAMVNGTAVGSGLASMVLFEA-NILSLLAEVL
Tintermedium-Thint.03G0055700.1              GSK-VNAAEAFKIAGIQHGFFELQPKEGLAMVNGTAVGSGLASMVLFEA-NILSLLAEVL
Taestivum-Traes_1BS_C3AA04888.1              -------------------------------------------MVLFEA-NILSLLAEVL
Taestivum-Traes_1DS_A8BD91E4A.1              GSK-VNAAEAFKIAGIQHGFFELQPKEGLAMVNGTAVGSGLASMVLFEA-NILSLLAEVL
Taestivum-Traes_1BS_C26CCE0E9.1              GSK-VNAAEAFKIAGIQHGFFELQPKEGLAMVNGTAVGSGLASMVLFEA-NILSLLAEVL
Taestivum-Traes_1DS_A21FCBA3D.1              GSK-VNAAEAFKIAGIQHGFFELQPKEGLAMVNGTAVGSGLASMVLFEA-NILSLLAEVL
Tintermedium-Thint.01G0012200.1              GSK-VNAAEAFKIAGIQHGFFELQPKEGLAMVNGTAVGSGLASMVLFEA-NILSLLAEVL
Tintermedium-Thint.02G0158600.1              GSK-VNAAKAFKIAGIQHGFFELQPKEGLAMVNGTAVGSGLASMVLFEA-NILSLLAEVL
Tintermedium-Thint.01G0012700.1              GSK-VNAAEAFKIAGIQHGFFELQPKEGLAMVNGTAVGSGLASMVLFEA-NILSLLAKVL
Taestivum-Traes_5BL_F50A47503.1              GSK-VNAAEAFKIAGIQHGFFELQPKEGLAMVNGTAVGSGLASMVLFEA-NILSLLAEVL
Tintermedium-Thint.02G0158700.1              GSK-VNAAEAFKIAGIQHGFFELQPKEGLAMVNGTAVGSGLASMVLFEA-NILSLLAEVL
Taestivum-Traes_5BL_AB528184A.1              GNK-VNAAEAFKIAGIQHGFFELQPKEGLAMVNGTAVGSGLASMVLFEA-NILSLLAEVL
Tintermedium-Thint.03G0055800.1              GSK-VNAAEAFKIAGIQHGFFELQPKEGLAMVNGTAVGSGLASMVLFEA-NILGLLAEVL
Taestivum-Traes_1AS_F9013A945.1              GSK-VNAAEAFKIAGIQHGFFELQPKEGLAMVNGTAVGSGLASMVLFEA-NILGVLAEVL
Taestivum-Traes_1DS_43DAC7A7F.1              GSK-VNAAGAFKIAGIEHGFFELQPKEGLAMVNGTAVGSGLASMVLFEA-NILGVLAEVL
Taestivum-Traes_1BS_BD86C90A7.1              GSK-VNAAEAFKIAGIQHGFFELQPKEGLAMVNGTAVGSGLASIVLFEA-NILGVLAEVL
Tintermedium-Thint.02G0158900.1              GSK-VNAAEAFKIAGIQHGFFELQPKEGLAMVNGTAVGSGLASMALFEA-NILGVLAEVL
Tintermedium-Thint.05G0274000.1              GTK-VNAAEAFKIAGIQHGFFELQPKEGLAMVNGTAVGSGLASMVLFEA-NILSLLAEVL
Osativa-LOC_Os04g43800.1                     GRK-VDAAEAFKIAGIQHGFFELQPKEGLAMVNGTAVGSGLASMVLFEA-NVLGVLAEVL
Pvaginatum-Pavag06G160300.1                  GRK-VGAAEAFNIAGIQHGFFELQPKEGLAMVNGTAVGSGLASMVLFDA-NVLAVLAEVL
Sbicolor-Sobic.006G148900.1                  GRK-VGAAEAFKIAGIQHGFFELQPKEGLAMVNGTAVGSGLASMVLFEA-NVLAVLAEVM
Pvirgatum-Pavir.7KG237800.1                  GRK-VDAAEAFKAAGIQHGFFELQPKEGLAMVNGTAVGSGLASPVLFEA-NVLAVLAEVL
Pvirgatum-Pavir.7NG355800.1                  GRK-VDAAEAFKVAGIQHGFFELQPKEGLAMVNGTAVGSGLASIVLFEA-NVLAVLAEVL
Sviridis-Sevir.7G178300.1                    GKK-VGAAEAFKAAGIQHGFFELQPKEGLAMVNGTAVGSGLASMVLFEA-NVLAVLAEVL
Zmays-Zm00001d003015_T001                    GRK-VGAAEAFEIAGIQHGFFELQPKEGLAMVNGTAVGSGLASMVLFEA-NVLAVLAEVM
Hvulgare-HORVU1Hr1G022060.1                  GTK-VNAAEAFKIAGIQHGFFELQPKEGLAMVNGTAVGSGLASIVLFEA-NILGVLAEVL
Taestivum-Traes_1BS_723922D171.2             GTK-VNAAEAFKIAGIQHGFFELQPKEGLAMVNGTAVGSGLASIVLFEA-NILGVLAEVL
Taestivum-Traes_1DS_A171C7D59.1              GTK-VNAAEAFKIAGIQHGFFELQPKEGLAMVNGTAVGSGLASIVLFEA-NILAVLAEVL
Tintermedium-Thint.01G0111500.1              GTK-VNAAEAFKIAGIQHSFFELQPKEGLAMVNGTAVGSGLASIVLFEA-NILAILAEVL
Tintermedium-Thint.03G0265300.1              GTK-VNAAEAFKIAGIQHGFFELQPKEGLAMVNGTAVGSGLASIVLFEA-NILGVLAEVL
Hvulgare-HORVU2Hr1G038140.1                  GTK-VNAAEAFKIAGIQHGFFELQPKEGLAMVNGTAVGSGLASIVLFEA-NVLGVLAEVL
Tintermedium-Thint.04G0171200.1              GTK-VNAAEAFKIAGIQHGFFELQPKEGLAMVNGTAVGSGLASIVLFEA-NVLGVLAEVL
Taestivum-Traes_2AS_958327519.3              ------------------------------MVNGTAVGSGLASIVLFEA-NVLGVLAEVL
Taestivum-Traes_2DS_28CA50371.1              ------------------------------MVNGTAVGSGLASIVLFEA-NVLGVLAEVL
Tintermedium-Thint.05G0160800.1              GTK-VNAAQAFKIAGIQHGFFELQPKEGLAMVNGTAVGSGLASIVLFEA-NVLGVLAEVL
Tintermedium-Thint.06G0299000.1              GTK-VNAAEAFKIAGIQHGFFELQPKEGLAMVNGTAVGSGLASIVLFEA-NVLGVLAEVL
Taestivum-Traes_2DS_3791A8A36.1              ------------------------------MVNGTAVGSGLASIVLFEA-NVLGVLAEVL
Tintermedium-Thint.06G0342800.1              GTK-VNAAEAFKIAGIQHGFFELQPKEGLAMVNGTAVGSGLASIVLFEA-NVLGVLAEVL
Tintermedium-Thint.05G0150900.1              ------------------------------------------------------------
Tintermedium-Thint.04G0242300.1              GTK-VNAAEAFKIAGIQHGFFELQPKEGLAMVNGTAVGSGLASIVLFEA-NILGVLAEVL
Tintermedium-Thint.06G0298700.1              GTK-VNAAEAFKIAGIQHGFFELQPKEGLAMVNGTAVGSGLASIVLFEA-NILGVLAEVL
Taestivum-Traes_2BS_88CF42F2E.1              GTK-VNAAEAFKIAGIQHGFFELQPKEGLAMVNGTAVGSGLASIVLFEA-NVLGVLAEVL
Tintermedium-Thint.04G0241900.1              GTK-VNAAEAFKIAGIQHGFFELQP-EGLAMVNGTAVGSGLASIVLFEATNVLGVLAEVL
Platifolius-Phala.06G165300.1                GRK-VNAAEAFKIAGIQHGFFELQPKEGLAMVNGTAVGSGLASTVLFEA-NILAVLAEVL
Streptochaeta-strangu_019385-RA              GRK-VDAAEAFKIAGIQHGFFELQPKEGLAMVNGTAVGSGLASTVLFEA-NILAVLAEVL
Streptochaeta-strangu_028944-RA              GRN-MDAAEAFKIAGIQHGFFELQPKEGLALVNGTAVGSGLASMVLFEA-NVLALLAEVL
Acomosus-Aco010091.1                         GMC-IDAAEAFQLARIIGGFFELRPKEGIALVNGTTVGSGLASIVLFEA-NILAVLSEVL
Zmarina-Zosma03g22300.1                      GTQ-ITAEKAFSLAGIHSGFFELQPKEGLALVNGTTVGSGLASTVLFDA-NILSILAEVL
Ecdeiocolea-Emoptg0003741_1G000600.1         GRK-VNAAEAFKIAGIEHGFFELQPKEGLAMVNGTAVGSGLASIVLFET-NILAVLAEVL
Jascendens-Joasc.05G060500.1                 GRK-VDAAEAFKIAGIQHGFFELQPKEGLAMVNGTAVGSGLASTVLFEA-NILTILAEVL
Ecdeiocolea-Emoptg0003741_1G000630.1         GRK-VDAAEAFKVAGIPHGFFKLQPKEGLAMVNGTSVGAALASMVLFDA-NILTVLAVVM
Jascendens-Joasc.05G060400.1                 GRK-VDAAEAFKIAGIPHGFFKLQPKEGLAIVNGTSVGAALASTVLFDA-NILAILAEVL
Bdistachyon-Bradi3g49250.2                   GRK-VDAAEAFKVAGIEGGFFTLNPKEGLAIVNGTSVGSALAATVLFDC-NVLAVLSEVL
Hvulgare-HORVU6Hr1G058820.1                  GKK-VDAAEAFKAAGIEGGFFKLNPKEGLAIVNGTSVGSALAATVLFDC-NVLAVLSEVL
Taestivum-Traes_6DL_1AEA7B869.1              GKK-VDAAEAFKAAGIEGGFFKLNPKEGLAIVNGTSVGSALAATVLFDC-NVLAVLSEVL
Tintermedium-Thint.18G0192300.1              GKK-VDAAEAFKAAGIEGGFFKLNPKEGLAIVNGTSVGSALAATVLFDC-NVLAVLSEVL
Tintermedium-Thint.17G0230800.1              GKK-VDAAEAFKVAGIEGGFFKLNPKEGLAIVNGTSVGSALAATVLFDC-NVLAVLSEVL
Tintermedium-Thint.17G0230900.1              GKK-VDAAEAFKVAGIEGGFFKLNPKEGLAIVNGTSVGSALAATVLFDC-NVLAVLSEVL
Ecoracana-ELECO.r07.2AG0138000.1             GRK-VDAAEAFKVAGIEGGFFKLNPKEGLAIVNGTSVGSALAATVMYDA-NVLAVLSEVL
Ecoracana-ELECO.r07.2BG0193650.1             GRK-VDAAEAFKVAGIEGGFFKLNPKEGLAIVNGTSVGSALAATVMYDA-NVLAVLSEVL
Pvaginatum-Pavag04G202700.1                  GRK-VDAAEAFKIAGIEGGFFRLNPKEGLAIVNGTSVGSALAATVLYDA-NLLTVLSEVL
Sviridis-Sevir.1G245000.1                    GRK-VDAAEAFKIAGIEGGFFKLNPKEGLAIVNGTSVGSALAATVLYDA-NVLAVLSEVL
Sviridis-Sevir.6G187100.1                    GRK-VDAAEAFKVAGIEGGFFRLNPKEGLAIVNGTSVGSALAATVLYDA-NLLAVLSEVL
Pvirgatum-Pavir.1NG356200.1                  GRK-VDAAEAFKLAGIDGGFFRLNPKEGLAIVNGTSVGSALAATVLYDA-NVLAVLSEVL
Sbicolor-Sobic.004G220300.1                  GRK-VDAAEAFKIAGIEGGFFKLNPKEGLAIVNGTSVGSALAATVMYDA-NVLAVLSEVL
Zmays-Zm00001d017274_T001                    GRK-VDAAEAFKIAGIEGGFFKLNPKEGLAIVNGTSVGSALAATVMYDA-NVLAVLSEVL
Zmays-Zm00001d051161_T003                    GRK-VDAAEAFKIAGIEGGFFKLNPKEGLAIVNGTSVGSALAATVMYDA-NVLAVLSEVL
Sviridis-Sevir.7G178200.1                    GRK-VDAAEAFKVAGIEGGFFKLNPKEGLAIVNGTSVGSALAAMVCFDA-NVLAVLSEVL
Osativa-LOC_Os02g41630.2                     GRK-VDAAEAFKLAGIEGGFFTLNPKEGLAIVNGTSVGSALAATVMFDA-NILAVLSEVL
Ecoracana-ELECO.r07.4AG0310430.1             GRK-VDAAEAFKVAGIEGGFFKLNPKEGLAIVNGTSVGSALAAMVCFDA-NVLAVLSEVL
Ecoracana-ELECO.r07.4BG0341620.1             GRK-VDAAEAFKVAGIEGGFFKLNPKEGLAIVNGTSVGSALAAMVCFDA-NVLAVLSEVL
Pvaginatum-Pavag06G160200.1                  GRK-VDAAEAFKAAGIEGGFFKLNPKEGLAIVNGTSVGSALAAMVCFDA-NVLAVLSEVL
Pvirgatum-Pavir.7KG238255.1                  GRK-VDAAEAFKVAGIEGGFFKLNPKEGLAIVNGTSVGSALAAVVCFDA-NVLAVLSEVL
Sbicolor-Sobic.006G148800.1                  GRK-VDAAEAFKVAGIQGGFFKLNPKEGLAIVNGTSVGSALAAMVCFDA-NVLAVLSSVL
Zmays-Zm00001d003016_T001                    GRK-VDAAEAFKAAGIEGGFFKLNPKEGLAIVNGTSVGSALAAMVCFDA-NVLAVLSEVL
Osativa-LOC_Os04g43760.1                     GKK-VDAAEAFKIAGIQGGFFRLEPKEGLAIVNGTSVGSALAAMVLYDA-NVLAVLSEVL
Platifolius-Phala.02G210400.1                GRK-VDAAEAFKIAGIQGGFFKLQPKEGLAIVNGTSVGAALAAMVLFDA-NILAVLSEVL
Platifolius-Phala.06G165100.1                GRK-VDAAEAFKIAGIQGGFFKLQPKEGLAIVNGTSVGAALAAMALFDA-NILAVLSEVL
Streptochaeta-strangu_019386-RA              GRK-VDAAEAFKVAGIEGGFFKLQPKEGLAIVNGTSVGAALAAMVLYDA-NILAVLSEVM
Streptochaeta-strangu_020769-RA              GRK-VDAAEAFKIAGIEGGFFKLQPKEGLAIVNGTSVGAALAAMVLYDA-NILAVLSEVM
Sviridis-Sevir.2G448300.1                    GRK-VDAAEAFKVAGIKGGFFKLNPKEGLAMVNGTSVGSALAAMVCFDA-NVLAVLAVVL
Sviridis-Sevir.7G177900.1                    GRK-VDAAEAFKVAGIEGGFFKLNPKEGLAMVNGTSVGSALAAMVCFDA-NVLAVLAVVL
```

```
Aamericanus-Acora.01G232100.1                SAVFCEVMQGKPEYTDHLTHKLKHHPGQIEAAAIMEHILDGSSYM-KLAQK----LHEQD
Aamericanus-Acora.10G174800.1                SAVFCEVMQGKPEYTDHLTHKLKHHPGQIEAAAIMEHILDGSSYM-KLAQK----LHEQD
Acomosus-Aco013943.1                         SAVFCEVMQGKPEFTDHLTHKLKHHPGQIEAAAIMEHVLEGSSYM-KMARK----LHELD
Aofficinalis-evm.model.AsparagusV1_01.827    -------MQGKPEYTDHLTHKLKHHPGQIEAAAIMEHILEGSSYM-KMAKK----LHEQD
Typha_latifolia-Tylat.14G083800.1            SAIFCEVMQGKPEYTDHLTHKLKHHPGQIEAAAIMEHILDGSSYM-KMAKK----IHELD
Aofficinalis-evm.model.AsparagusV1_04.71     SAVFCEVMQGKPEFTDHLTHKLKHHPGQIEAAAIMEHILEGSSYM-QMAKK----VHELD
Aofficinalis-evm.model.AsparagusV1_04.93     SAVFCEVMQGKPEFTDHLTHKLKHHPGQIEAAAIMEHILEGSSYM-QMAKK----VHELD
Aofficinalis-evm.model.AsparagusV1_10.1948   SAVFCEVMQGKPEFTDHLTHKLKHHPGQIEAAAIMEHILDGSSYM-QMAKK----VHELD
Typha_latifolia-Tylat.03G127700.1            SAIFCEVMQGKPEFTDHLTHKLKHHPGQIEAAAIMEHILEGSSYV-KMAKK----LHEVD
Spolyrhiza-Spipo11G0025500                   SAVFCEVMQGKPEFTDHLTHKLKHHPGQIEAAAIMEHILEGSSYM-KMAKK----IHELD
Spolyrhiza-Spipo1G0003500                    SAVFCEVMQGKPEFTDHLTHKLKHHPGQIEAAAIMEHILEGSSYM-KMAKK----LHELD
Phoenix_dactylifera-LOC103718264             SAIFCEVMQGKPEFTDHLTHKLKHHPGQIEAAAIMEYILEGSSYM-KMAKK----LHELD
Macuminata-GSMUA_Achr1T09070_001             SAVFCEVMQGKPEFTDHLTHKLKHHPGQIEAAAIMEHILEGSSYM-KMAQK----LHEQD
Acomosus-Aco006987.1                         SAVFCEVMQGKPEFTDHLTHKLKHHPGQIEAAAIMEHVLEGSSYM-RMAKK----LHELD
Acomosus-Aco027752.1                         SAVFCEVMQGKPEFTDHLTHKLKHHPGQIEAAAIMEHVLEGSSYM-RMAKK----LHELD
Cscoparia-Csco27g03980.1                     SAVFCEVMQGKPEYTDHLTHKLKHHPGQIEAAAIMEHVLDGSSYM-KMAKK----VHEQD
Cscoparia-Csco27g03990.1                     SAVFCEVMQGKPEYTDHLTHKLKHHPGQIEAAAIMEHVLDGSSYM-KMAKK----VHEQD
Macuminata-GSMUA_Achr11T16380_001            SAVFCEVMQGNPEFTDHLTHRLKHHPGQIEAAAIMEHILDGSSYM-KMAKK----LHEQD
Macuminata-GSMUA_Achr11T22840_001            SAVFCEVMQGKPEFTDHLTHKLKHHPGQIEAAAIMEHILDGSSYM-KMAKK----LHEQD
Macuminata-GSMUA_Achr2T00240_001             SAVFCEVMLGKPEFTDHLTHKLKHHPGQIEAAAIMEHILDGSSFM-KIAKK----LHEQD
Macuminata-GSMUA_Achr5T18560_001             SAVFCEVMQGKPEFTDHLTHKLKHHPGQIEAAAIMEHILDGSSYM-KMAKK----LHEQD
Macuminata-GSMUA_Achr8T18960_001             SAIFCEVMQGKPEFTDHLTHKLRHHPGQIEAAAIMEHILDGSSYM-KMAKN----LHEQD
Macuminata-GSMUA_Achr5T03950_001             SAIFCEVMQGKPEFTDHLTHKLKHHPGQIEAAAIMEHVLEGSSYM-KMAKK----LHEQD
Phoenix_dactylifera-LOC103703950             SAIFCEVMNGKPEFTDHLTHKLKHHPGQIEAAAIMEHVLDGSSYM-KMAKK----LHELD
Acoerulea-Aqcoe1G140100.1                    SAIFAEVMQGKPEFTDHLTHKLKHHPGQIEAAAIMEHILDGSSYV-KSAKK----LHDTD
Acoerulea-Aqcoe2G144500.1                    SAIFAEVMQGKPEFTDHLTHKLKHHPGQIEAAAIMEHILNGSSYV-KAAEK----LHEMD
Typha_latifolia-Tylat.05G053500.1            SAVFCEVMQGKPEFTDHLTHKLKHHPGQIEAAAVMEHILAGSSYM-SAAKK----LHELD
Zmarina-Zosma05g29470.1                      SAVFCEVMQGKPEFTDHLTHKLKHHPGQIEAAAIMEHILDGSSYM-KMAAK----IHEQD
Acomosus-Aco007727.1                         SAIFCEVMQGKPEFTDHLTHKLKHHPGQIEAAAIMEHILDGSSYM-KAAKK----LHELD
```

```
Bvulgarisssp-Bevul.7G054300.1          SAVFAEVMNGKPEFTDHLTHKLKHHPGQIEAAAIMEHILDGSSYM-KAAKA----LHELD
Bdistachyon-Bradi3g47110.1             SAVFCEVMNGKPEFTDHLTHKLKHHPGQIEAAAIMEHILEGSSYM-KLAKK----LGDLD
Bdistachyon-Bradi3g47120.1             SAVFCEVMNGKPEFTDHLTHKLKHHPGQIEAAAIMEHILEGSSYM-KLAKK----LGDLD
Bdistachyon-Bradi3g48840.1             SAVFCEVMNGKPEFTDHLTHKLKHHPGQIEAAAIMEHILEGSSYM-KLAKK----LGDLD
Bdistachyon-Bradi3g49280.1             SAVFCEVMNGKPEFTDHLTHKLKHHPGQIEAAAIMEHILEGSSYM-KLAKK----LGDLD
Hvulgare-HORVU6Hr1G058840.1            SAVFCEVMNGKPEFTDHLTHKLKHHPGQIEAAAIMEHILEGSSYM-KLAKK----LGDLD
Taestivum-Traes_6BL_C96F626D9.1        SAVFCEVMNGKPEFTDHLTHKLKHHPGQIEAAAIMEHILEGSSYM-KLAKK----LGDLD
Tintermedium-Thint.18G0192700.1        SAVFCEVMNGKPEFTDHLTHKLKHHPGQIEAAAIMEHILEGSSYM-KLAKK----LGDLD
Taestivum-Traes_6DL_04CD7E82C.1        SAVFCEVMNGKPEFTDHHTHKLKHHPGQIEAAAIMEHILEGSSYM-KLAKK----LGDLD
Bdistachyon-Bradi3g49270.1             SAVFCEVMNGKPEFTDHLTHKLKHHPGQIEAAAIMEHILEGSSYM-KLAKK----QGELD
Ecoracana-ELECO.r07.2AG0138020.1       SAVFCEVMNGKPEYTDHLTHKLKHHPGQIEAAAIMEHILEGSSYM-KLAKK----MGELD
Ecoracana-ELECO.r07.2BG0193670.1       SAVFCEVMHGKPEYTDHLTHKLKHHPGQIEAAAIMEHILEGSSYM-KLAKK----MGELD
Osativa-LOC_Os02g41670.1               SAVFCEVMNGKPEYTDHLTHKLKHHPGQIEAAAIMEHILEGSSYM-KLAKK----LGELD
Osativa-LOC_Os02g41680.1               SAVFCEVMNGKPEYTDHLTHKLKHHPGQIEAAAIMEHILEGSSYM-KHAKK----LGELD
Pvaginatum-Pavag04G202900.1            SAVFCEVMNGKPEYTDHLTHKLKHHPGQIEAAAIMEHILEGSSYM-KLAKK----LGELD
Pvaginatum-PavagK321000.1              SAVFCEVMNGKPEYTDHLTHKLKHHPGQIEAAAIMEHILEGSSYM-KLAKK----LGELD
Pvaginatum-PavagK321300.1              SAVFCEVMNGKPEYTDHLTHKLKHHPGQIEAAAIMEHILEGSSYM-KLAKK----LGELD
Pvaginatum-PavagK321100.1              SAVFCEVMNGKPEYTDHLTHKLKHHPGQIEAAAIMEHILEGSSYM-KLAKK----LGELD
Pvaginatum-PavagK321200.1              SAVFCEVMNGKPEYTDHLTHKLKHHPGQIEAAAIMEHILEGSSYM-KLAKK----LGELD
Pvirgatum-Pavir.1KG386500.1            SAVFCEVMNGKPEFTDHLTHKLKHHPGQIEAAAIMEHILEGSSYM-KHAKK----LGELD
Pvirgatum-Pavir.1NG356700.1            SAVFCEIMNGKPEFTDHLTHKLKHHPGQIEAAAIMEHILEGSSYM-KLAKK----LGELD
Sviridis-Sevir.1G245166.1              SAVFCEVMNGKPEYTDHLTHKLKHHPGQIEAAAIMEHILEGSSYM-KLAKK----LGELD
Sviridis-Sevir.1G245232.1              SAVFCEVMNGKPEYTDHLTHKLKHHPGQIEAAAIMEHILEGSSYM-KLAKK----LGELD
Pvirgatum-Pavir.1NG356800.1            SAVFCEVMNGKPEFTDHLTHKLKHHPGQIEAAAIMEHVLEGSSYM-KLAKK----LGELD
Sbicolor-Sobic.004G220500.1            SAVFCEVMNGKPEYTDHLTHKLKHHPGQIEAAAIMEHILEGSSYM-KLAKK----LGELD
Sbicolor-Sobic.004G220600.2            SAVFCEVMNGKPEYTDHLTHKLKHHPGQIEAAAIMEHILEGSSYM-KLAKK----LGELD
Sbicolor-Sobic.004G220700.1            SAVFCEVMNGKPEYTDHLTHKLKHHPGQIEAAAIMEHILEGSSYM-KLAKK----LGELD
Sviridis-Sevir.1G245300.2              SAVFCEVMNGKPEYTDHLTHKLKHHPGQIEAAAIMEHILEGSSYM-KLAKK----LGELD
Zmays-Zm00001d017276_T001              SAVFCEVMNGKPEYTDHLTHKLKHHPGQIEAAAIMEHILEGSSYM-KLAKK----LGELD
Zmays-Zm00001d017279_T001              SAVFCEVMNGKPEYTDHLTHKLKHHPGQIEAAAIMEHILEGSSYM-KLAKK----LGELD
Zmays-Zm00001d051166_T001              SAVFCEVMNGKPEYTDHLTHKLKHHPGQIEAAAIMEHILEGSSYM-KLAKK----LGELD
Ecoracana-ELECO.r07.2AG0138030.1       SAVFCEVMNGKPEYTDHLTHKLKHHPGQIEAAAIMEHILEGSSYM-KLAKK----LGELD
Ecoracana-ELECO.r07.2BG0193680.1       SAVFCEVMNGKPEYTDHLTHKLKHHPGQIEAAAIMEHILEGSSYM-KLAKK----MGELD
Osativa-LOC_Os05g35290.1               SAVFCEVMNGKPEYTDHLTHKLKHHPGQIEAAAIMEHILEGSSYM-KEAKR----LGELD
Platifolius-Phala.02G210700.1          SAVFCEVMNGKPEYTDHLTHKLKHHPGQIEAAAIMEHILEGSSYM-KLAKK----LGELD
Ecoracana-ELECO.r07.4AG0310420.1       SAVFCEVMNGKPEYTDHLTHKLKHHPGQIEAAAIMEHVLEGSSYM-ALAKK----MGELD
Ecoracana-ELECO.r07.4BG0341610.1       SAVFCEVMNGKPEYTDHLTHKLKHHPGQIEAAAIMEHVLEGSSYM-ALAKK----MGELD
Bdistachyon-Bradi3g49260.1             SAVFCEVMTGKPEFTDHLTHKLKHHPGQIEAAAIMEHILEGSSYM-KEAKK----QGELD
Hvulgare-HORVU0Hr1G016330.1            SAVFCEVMNGKPEFTDHLTHKLKHHPGQIEAAAIMEHILEGSSYM-KQAKK----LGELD
Tintermedium-Thint.17G0260200.1        SAVFCEVMNGKPEFTDHLTHKLKHHPGQIEAAAIMEHILEGSSYM-KQAKK----LGELD
Tintermedium-Thint.18G0192600.1        SAVFCEVMNGKPEFTDHLTHKLKHHPGQIEAAAIMEHILEGSSYM-KQAKK----LGELD
Tintermedium-Thint.17G0231000.1        SAVFCEVMNGKPEFTDHLTHKLKHHPGQIEAAAIMEHILEGSSYM-KQAKK----LGELD
Ecoracana-ELECO.r07.2AG0138010.1       SAVFCEVMNGKPEYTDHLTHKLKHHPGQIEAAAIMEHILEGSSYM-KLAKK----LGELD
Ecoracana-ELECO.r07.2BG0193660.1       SAVFCEVMNGKPEYTDHLTHKLKHHPGQIEAAAIMEHILEGSSYM-KLAKK----LGELD
Osativa-LOC_Os02g41650.3               SAVFCEVMTGKPEYTDHLTHKLKHHPGQIEAAAIMEHILEGSSYM-KLAKK----LGELD
Pvaginatum-Pavag04G202800.1            SAVFCEVMNGKPEYTDHLTHKLKHHPGQIEAAAIMEHILEGSSYM-KLAKK----LGELD
Pvaginatum-PavagK320900.1              SAVFCEVMNGKPEYTDHLTHKLKHHPGQIEAAAIMEHILEGSSYM-KLAKK----LGELD
Pvirgatum-Pavir.1NG356400.1            SAVFCEVMTGKPEFTDHLTHKLKHHPGQIEAAAIMEHILEGSSYM-KLAKK----LGELD
Sviridis-Sevir.1G245100.1              SAVFCEVMTGKPEFTDHLTHKLKHHPGQIEAAAIMEHILEGSSYM-KLAKK----LGELD
Sbicolor-Sobic.004G220400.1            SAVFCEVMTGKPEFTDHLTHKLKHHPGQIEAAAIMEHILEGSSYM-KLAKK----LGELD
Zmays-Zm00001d017275_T001              SAVFCEVMTGKPEFTDHLTHKLKHHPGQIEAAAIMEHVLEGSSYM-KLAKK----LGELD
Zmays-Zm00001d051163_T002              SAVFCEVMTGKPEFTDHLTHKLKHHPGQIEAAAVMEHILEGSSYM-KLAKR----LGELD
Pvirgatum-Pavir.1KG386300.1            SAVFCEVMTGKPEFTDHLTHKLKHHPGQIEAAAIMEHILEGSSYM-KLAKK----LGELD
Platifolius-Phala.02G210600.1          SAVFCEVMNGKPEYTDHLTHKLKHHPGQIEAAAIMEHILEGSSYM-KLAKK----LGELD
Streptochaeta-strangu_020767-RA        SAVFCEAMNGKPEYTDHLTHKLKHHPGQIEAAAIMEHILEGSSYM-KLAKK----LGELD
Bdistachyon-Bradi5g15830.1             SAVFCEVMNGKPEFTDHLTHKLKHHPGQIEAAAIMEHILEGSSYM-MLAKK----LGELD
Hvulgare-HORVU2Hr1G089440.4            SAVFCEVMNGKPEYTDHLTHKLKHHPGQIEAAAIMEHILEGSSYM-MLAKK----LGELD
Taestivum-Traes_2AL_0DDC60D6E.1        SAVFCEVMNGKPEYTDHLTHKLKHHPGQIEAAAIMEHILEGSSYM-MLAKK----LGELD
Tintermedium-Thint.06G0475900.1        SAVFCEVMNGKPEYTDHLTHKLKHHPGQIEAAAIMEHILEGSSYM-MLAKK----LGELD
Tintermedium-Thint.06G0476200.1        SAVFCEVMNGKPEYTDHLTHKLKHHPGQIEAAAIMEHILEGSSYM-MLAKK----LGELD
Taestivum-Traes_2AL_2763280FC.1        SAVFCEVMNGKPEYTDHLTHKLKHHPGQIEAAAIMEHILEGSSYM-MLAKK----LGELD
Taestivum-Traes_2DL_8C21AAAEF.1        SAVFCEVMNGKPEYTDHLTHKLKHHPGQIEAAAIMEHILEGSSYM-MLAKK----LGELD
Tintermedium-Thint.V0383500.1          SAVFCEVMNGKPEYTDHLTHKLKHHPGQIEAAAIMEHILEGSSYM-MLAKK----LGELD
Taestivum-Traes_2AL_9EC3226F7.1        SGVFCEVMNGKPEFTDHLTHKLKHHPGQIEAAAIMEHILEGSSYM-MLAKK----LGELD
Tintermedium-Thint.05G0273800.1        SAVFCEVMNGKPEYTDHLTHKLKHHPGQIEAAAIMEHILEGSSYM-MLAKK----LGELD
Tintermedium-Thint.V0383400.1          SAVFCEVMNGKPEYTDHLTHKLKHHPGQIEAATIMEHILEGSSYM-MLAKK----LGELD
Taestivum-Traes_2BL_13D5272D7.1        SAVFCEVMNGKPEYTDHLTHKLKHHPGQIEVAAIMEHILEGSSYM-MLAKK----LGELD
Tintermedium-Thint.06G0475700.1        SAVFCEVMNGKPEYTDHLTHKLKHHPGQIEAAAIMEHILEGSSYM-MLAKK----LGELD
Taestivum-Traes_2BL_C051606EA.2        SAVFCEVMNGKPEYTDHLTHKLKHHPGQIEAAAIMEHILEGSSYM-MLAKK----LGELD
Tintermedium-Thint.V0383600.1          SAVFCEVMNGKPEYTDHLTHKLKHHPGQIEAAAIMEHILEGSSYM-MLAKK----LGELD
Taestivum-Traes_2AL_9D78F85E2.1        SAVFCEVMNGKPEYTDHLTHKLKHHPGQIEAAAIMEHILEGSSYM-MLAKK----LGELD
Tintermedium-Thint.05G0274100.1        SAVFCEVMNGKPEYTDHLTHKLKHHPGQIEAAAIMEHILEGSSYM-MLAKK----LGELD
Tintermedium-Thint.06G0476300.1        SAVFCEVMNGKPEYTDHLTHKLKHHPGQIEAAAIMEHILEGSSYM-MLAKK----LGELD
Taestivum-Traes_2BL_5A50FDA1A.1        SAVFCEVMNGKPEYTDHLTHKLKHHPGQIEAAAIMEHILEGSSYM-MLAKK----LGELD
Tintermedium-Thint.05G0273900.1        SAVFCEVMNGKPEYTDHLTHKLKHHPGQIEAAAIMEHILEGSSYM-MLAKK----LGELD
Taestivum-Traes_2DL_30F8555D4.1        ------------------------------------------------------------
Taestivum-Traes_2DL_83168C1E0.1        ------------------------------------------------------------
Taestivum-Traes_2DL_B978B4E22.1        ------------------------------------------------------------
Taestivum-Traes_1AS_0B9295A68.1        SAVFCEVMNGKPEYTDQLTHKLKHHPGQIEAAAIMEHILEGSSYM-ALAKK----LGELD
Taestivum-Traes_1BS_7068F56E6.1        SAIFCEVMNGKPEYTDHLTHKLKHHPGQIEAAAIMEHILEGSSYM-ALAKK----LGELD
Taestivum-Traes_1BS_AEA817616.1        SAVFCEVMNGKPEYTDHLTHKLKHHPGQIEAAAIMEHILEGSSYM-ALAKK----LGELD
Taestivum-Traes_4AL_892C47ED5.1        SAVFCEVMNGKPEYTDHLTHKLKHHPGQIEAAAIMEHILEGSSYM-ALAKK----LGELD
Tintermedium-Thint.03G0055900.1        SAVFCEVMNGKPEYTDHLTHKLKHHPGQIEAAAIMEHILEGSSYM-ALAKK----LGELD
Tintermedium-Thint.01G0012900.1        SAVFCEVMNGKPEYTDHLTHKLKHHPGQIEAAAIMEHILEGSSYM-ALAKK----LGELD
Tintermedium-Thint.02G0158500.1        SAVFCEVMNGKPEYTDHLTHKLKHHPGQIEAAAIMEHILEGSSYM-ALAKK----LGELD
Tintermedium-Thint.03G0055700.1        SAVFCEVMNGKPEYTDHLTHKLKHHPGQIEAAAIMEHILEGSSYM-ALAKK----LGELD
Taestivum-Traes_1BS_C3AA04888.1        SAVFCEVMNGKPEYTDHLTHKLKHHPGQIEAAAIMEHILEGSSYM-ALAKK----LGELD
Taestivum-Traes_1DS_A8BD91E4A.1        SAVFCEVMNGKPEYTDHLTHKLKHHPGQIEAAAIMEHILEGSSYM-ALAKK----LGELD
Taestivum-Traes_1BS_C26CCE0E9.1        SAVFCEVMNGKPEYTDHLTHKLKHHPGQIEAAAIMEHILEGSSYM-ALAKK----LGELD
Taestivum-Traes_1DS_A21FCBA3D.1        SAVFCEVMNGKPEYTDHLTHKLKHHPGQIEAAAIMEHILEGSSYM-ALAKK----LGELD
Tintermedium-Thint.01G0012200.1        SAVFCEVMNGKPEYTDHLTHKLKHHPGQIEAAAIMEHILEGSSYM-ALAKK----LGELD
Tintermedium-Thint.02G0158600.1        SAVFCEVMNGKPEYTDHLTHKLKHHPGQIEAAAIMEHILEGSSYM-ALAKK----LGELD
Tintermedium-Thint.01G0012700.1        SAVFCEVMNGKPEYTDHLTHKLKHHPGQIEAAAIMEHILEGSSYM-ALAKK----LGELD
Taestivum-Traes_5BL_F50A47503.1        SAVFCEVMNGKPEYTDHLTHKLKHHPGQIEAAAIMEHILEGSSYM-MLAKK----LGELD
Tintermedium-Thint.02G0158700.1        SAVFCEVMNGKPEYTDHLTHKLKHHPGQIEAAAIMEHILEGSSYM-ALAKK----LGELD
Taestivum-Traes_5BL_AB528184A.1        SAVFCEVMNGKPEYTDHLTHKLKHHPGQIEAAAIMEHILEGSSYM-ALAKK----LGELD
Tintermedium-Thint.03G0055800.1        SAVFCEVMNGKPEYTDHLTHKLKHHPGQIEAAAIMEHILEGSSYM-ALAKK----LGELD
Taestivum-Traes_1AS_F9013A945.1        SAVFCEVMNGKPEYTDHLTHKLKHHPGQIEAAAIMEHILEGSSYM-ALAKK----LGELD
Taestivum-Traes_1DS_43DAC7A7F.1        SAIFCEVMNGKPEYTDHLTHKLKHHPGQIEAAAIMEHILEGSSYM-ALAKK----LGELD
Taestivum-Traes_1BS_BD86C90A7.1        SAVFCEVMNGKPEYTDHLTHKLKHHPGQIEAAAIMEHILEGSSYM-ALAKK----LGELD
Tintermedium-Thint.02G0158900.1        SAIFCEVMNGKPEYTDHLTHKLKHHPGQIEAAAIMEHILEGSSYM-ALAKK----LGELD
Tintermedium-Thint.05G0274000.1        SAVFCEVMNGKPEYTDHLTHKLKHHPGQIKAAAIMEHILEGSSYM-MLAKK----LGVLD
Osativa-LOC_Os04g43800.1               SAVFCEVMNGKPEYTDHLTHKLKHHPGQIEAAAIMEHILEGSSYM-MLAKK----LGELD
Pvaginatum-Pavag06G160300.1            SAVFCEVMNGKPEYTDHLTHKLKHHPGQIEAAAIMEHILEGSSYM-MLAKK----LGELD
Sbicolor-Sobic.006G148900.1            SAVFCEVMNGKPEYTDHLTHKLKHHPGQIESAAIMEHILDGSSYM-MLAKK----MGELD
Pvirgatum-Pavir.7KG237800.1            SAVFCEVMNGKPEYTDHLTHKLKHHPGQIEAAAIMEHILEGSSYM-ALAKK----LGELD
Pvirgatum-Pavir.7NG355800.1            SAVFCEVMNGKPEYTDHLTHKLKHHPGQIEAAAIMEHILDGSSYM-MLAKK----LGELD
Sviridis-Sevir.7G178300.1              SAVFCEVMNGKPEYTDHLTHKLKHHPGQIEAAAIMEHILEGSSYM-KLAKK----LGELD
Zmays-Zm00001d003015_T001              SAVFCEVMNGKPEYTDHLTHKLKHHPGQIESAAIMEHILEGSSYM-ALAKK----LGELD
Hvulgare-HORVU1Hr1G022060.1            SAVFCEVMNGKPEYTDHLTHKLKHHPGQIEAAAIMEHILEGSSYM-MLAKK----LGELD
Taestivum-Traes_1BS_723922D171.2       SAVFCEVMNGKPEYTDHLTHKLKHHPGQIEAAAIMEHILEGSSYM-MLAKK----LGELD
Taestivum-Traes_1DS_A171C7D59.1        SAVFCEVMNGKPEYTDHLTHKLKHHPGQIEAAAIMEHILEGSSYM-MLAKK----LGELD
Tintermedium-Thint.01G0111500.1        SAVFCEVMNGKPEYTDHLTHKLKHHPGQIEAAAIMEHILEGSSYM-MLAKK----LGELD
Tintermedium-Thint.03G0265300.1        SAVFCEVMNGKPEYTDHLTHKLKHHPGQIEAAAIMEHVLEGSSYM-MLAKK----LGELD
Hvulgare-HORVU2Hr1G038140.1            SAVFCEVMNGKPEYTDHLTHKLKHHPGQIEAAAIMEHILEGSSYM-MLAKK----LGELD
Tintermedium-Thint.04G0171200.1        SAVFCEVMNGKPEYIDHLTHKLKHHPGQIEAAAIMEHILEGSSYM-MLAKK----LGELD
Taestivum-Traes_2AS_958327519.3        SAVFCEVMNGKPEYTDHLTHKLKHHPGQIEAAAIMEHILEGSSYM-MLAKK----LGELD
Taestivum-Traes_2DS_28CA50371.1        SAVFCEVMNGKPEYTDHLTHKLKHHPGQIEAAAIMEHILEGSSYM-MLAKK----LGELD
Tintermedium-Thint.05G0160800.1        SAVFCEVMNGKPEYTDHLTHKLKHHPGQIEAAAIMEHILEGSSYM-MLAKK----LGELD
Tintermedium-Thint.06G0299000.1        SAVFCEVMNGKPEYTDHLTHKLKHHPGQIEAAAIMEHILEGSSYM-MLAKK----LGELD
Taestivum-Traes_2DS_3791A8A36.1        SAVFCEVMNGKPEYTDHLTHKLKHHPGQIEAAAIMEHILEGSSYM-MLAKK----LGELD
Tintermedium-Thint.06G0342800.1        SAVFCEVMNGKPEYTDHLTHKLKHHLGQIEAAAIMEHILEGSSYM-MLAKK----LGELD
Tintermedium-Thint.05G0150900.1        ----------------------------------MEHILEGSSYM-MLAKK----LGELD
Tintermedium-Thint.04G0242300.1        SAVFCEVMNGKPEYTDHLTHKLKHHPGQIEAAAIMEHILEGSSYM-MLAKK----LGELD
Tintermedium-Thint.06G0298700.1        SAVFCEVMNGKPEYTDHLTHKLKHHPGQIEAAAIMEHILEGSSYM-MLAKK----LGELD
Taestivum-Traes_2BS_88CF42F2E.1        SAVFCEVMNGKPEYTDHLTHKLKHHPGQIEAAAIMEHILEGSSYM-MLAKK----LGELD
Tintermedium-Thint.04G0241900.1        SAVFCEVMNGKPEYTDHLTHKLKHHPGQIEAAAIMEHILEGSSYM-MLAKK----LGELD
Platifolius-Phala.06G165300.1          SAVFCEVMNGKPEYTDHLTHKLKHHPGQIEAAAIMEHILEGSSYM-MLAKK----QGELD
Streptochaeta-strangu_019385-RA        SAVFCEVMNGKPEYTDHLTHKLKHHPGQIEAAAIMEHILEGSSYM-MLAKK----LGELD
Streptochaeta-strangu_028944-RA        AAVFCEVMHGKPEYTDHLTHKLKHHPGQIEAAAIMEHILEGSSYM-KLAKE----LAELD
```

```
Acomosus-Aco010091.1                            SAVFCEVMQGKPEFTDHLTHKLKHHPGQIEAAAIMEHILEGSSYM-KKAKE----LHELD
Zmarina-Zosma03g22300.1                         SAVFCEVMAGKPEFTDHLTHKLKHHPGQIEAAAIMEHILDGSSYM----KN----VNEHD
Ecdeiocolea-Emoptg0003741_1G000600.1            SAVFCEVMTGKPEYTDHLTHKLKHHPGQIEAAAIMEHILDGSSYM-KLAKK----LGELD
Jascendens-Joasc.05G060500.1                    SAVFCEVMTGKPEYTDHLTHKLKHHPGQIEAAAIMEHILEGSSYM-KLAKK----LGDLD
Ecdeiocolea-Emoptg0003741_1G000630.1            SAVFCEVMTGKPEYTDHLTHKLKHHPGQIEAAAIMEHILAGSSFM-KLAKK----LGELD
Jascendens-Joasc.05G060400.1                    SAVFCEVMTGKPEYTDHLTHRLKHHPGQIEAAAIMEHILEGSSFM-KLAKK----LGELD
Bdistachyon-Bradi3g49250.2                      SAVFCEVMNGKPEFTDHLTHKLKHHPGSIEAAAIMEHILAGSSFM-SHAKK----VNEID
Hvulgare-HORVU6Hr1G058820.1                     SAVFCEVMNGKPEFTDHLTHKLKHHPGSIEAAAIMEHILAGSAFM-GHAKK----VNEID
Taestivum-Traes_6DL_1AEA7B869.1                 SAVFCEVMNGKPEFTDHLTHKLKHHPGSIEAAAIMEHILAGSAFM-GHAKK----VNEID
Tintermedium-Thint.18G0192300.1                 SAVFCEVMNGKPEFTDHLTHKLKHHPGSIEAAAIMEHILAGSAFM-GHAKK----VNEID
Tintermedium-Thint.17G0230800.1                 SAVFCEVMNGKPEFTDHLTHKLKHHPGSIEAAAIMEHILAGSAFM-GHAKK----VNEID
Tintermedium-Thint.17G0230900.1                 SAVFCEVMNGKPEFTDHLTHKLKHHPGSIEAAAIMEHILAGSAFM-GHAKK----VNEID
Ecoracana-ELECO.r07.2AG0138000.1                SAVFCEVMNGKPEFADHLTHKLKHHPGSIEAAAIMEHILEGSSFM-KHAKK----VNEMD
Ecoracana-ELECO.r07.2BG0193650.1                SAVFCEVMNGKPEFADHLTHKLKHHPGSIEAAAIMEHILEGSSFM-KHAKK----VNEMD
Pvaginatum-Pavag04G202700.1                     SAVFCEVMNGKPEYTDHLTHKLKHHPGSIEAAAIMEHILEGSAFM-KHAKK----VNELD
Sviridis-Sevir.1G245000.1                       SAVFCEVMNGKPEYTDHLTHKLKHHPGSIEAAAIMEHILEGSEFM-KHAKK----VNELD
Sviridis-Sevir.6G187100.1                       SAVFCEVMNGKPEYTDHLTHKLKHHPGSIEAAAIMEHILEGSAFM-KHAKK----VNELD
Pvirgatum-Pavir.1NG356200.1                     SAVFCEVMNGKPEYTDHLTHKLKHHPGSIEAAAIMEHILEGSSFM-KQAKK----VNELD
Sbicolor-Sobic.004G220300.1                     SAIFCEVMNGKPEYTDHLTHKLKHHPGSIEAAAIMEHILDGSAFM-KHAKK----VNELD
Zmays-Zm00001d017274_T001                       SAVFCEVMNGKPEYTDHLTHKLKHHPGSIEAAAIMEHILDGSSFM-KQAKK----LNELD
Zmays-Zm00001d051161_T003                       SAVFCEVMNGKPEYTDHLTHKLKHHPGSIEAAAIMEHILEGSSFM-KHAKK----VNELD
Sviridis-Sevir.7G178200.1                       SAVFCEVMNGKPEYTDHLTHKLKHHPGSIEAAAIMEHILEGSSFM-KHAKK----VNELD
Osativa-LOC_Os02g41630.2                        SAVFCEVMNGKPEYTDHLTHKLKHHPGSIEAAAIMEHILAGSSFM-SHAKK----VNEMD
Ecoracana-ELECO.r07.4AG0310430.1                SAVFCEVMNGKPEFADHLTHKLKHHPGSIEAAAIMEHILEGSSFM-KHAKE----VNAMD
Ecoracana-ELECO.r07.4BG0341620.1                SAVFCEVMNGKPEFADHLTHKLKHHPGSIEAAAIMEHILEGSSFM-KHAKE----VNAMD
Pvaginatum-Pavag06G160200.1                     SAVFCEVMNGKPEYTDHLTHKLKHHPGSIEAAAIMEHILDGSSFM-KHAKE----VNAQD
Pvirgatum-Pavir.7KG238255.1                     SAVFCEVMNGKPEYTDHLTHKLKHHPGSIEAAAIMEHILDGSSFM-KHAKE----VNAMD
Sbicolor-Sobic.006G148800.1                     SAVFCEVMNGKPEYTDHLTHKLKHHPGSIESAAIMEHILDGSSFM-KHAKE----VNAMD
Zmays-Zm00001d003016_T001                       SAVFCEVMNGKPEYTDHLTHKLKHHPGSIEAAAIMEHILDGSSFM-KHAKE----VNAMD
Osativa-LOC_Os04g43760.1                        SAVFCEVMNGKPEYTDHLTHKLKHHPGSIEAAAIMEHILAGSAFM-PHAQK----VNEVD
Platifolius-Phala.02G210400.1                   SAVFCEVMNGKPEYTDHLTHKLKHHPGQIEAAAIMEHILDGSSFM-KHAKK----VNEMD
Platifolius-Phala.06G165100.1                   SAVFCEVMNGKPEYTDHLTHKLKHHPGQIEAAAIMEYILDGSSFM-KHAKK----VNEMD
Streptochaeta-strangu_019386-RA                 SAVFCEVMNGKPEYTDHLTHKLKHHPGQIEAAAIMEHILDGSSFM-KHAKK----VNELD
Streptochaeta-strangu_020769-RA                 SAVFCEVMNGKPEYTDHLTHKLKHHPGQIEAAAIMEHILAGSSFM-KHAKK----VNELD
Sviridis-Sevir.2G448300.1                       SAVFCEVMNGKPEYADHLIHKLKHHPGSIEAAAIMEHILDGSSLM-KHAKE----VNAMD
Sviridis-Sevir.7G177900.1                       SAVFCEVMNGKPEYADHLTHKLKHHPGSIEAAAIMEHILDGSSLM-KHAKE----VNAMD


Aamericanus-Acora.01G232100.1                   PLQKPKQDRYALRTSPQWLGPQIEVIRAATQSIQREINSVNDNPLIDVSRNKALHGGNFQ
Aamericanus-Acora.10G174800.1                   PLQKPKQDRYALRTSPQWLGPQIEVIRSATQSIQREINSVNDNPLIDVSRNKALHGGNFQ
Acomosus-Aco013943.1                            PLQKPKQDRYALRTSPQWLGPQIEVIRAATISIEREINSVNDNPLIDVSRNKALHGGNFQ
Aofficinalis-evm.model.AsparagusV1_01.827       PLQKPKQDRYALRTSPQWLGPLIEVIRSSTKSIEREINSVNDNPLIDVSRNKALHGGNFQ
Typha_latifolia-Tylat.14G083800.1               PLQKPKQDRYALRTSPQWLGPQIEVIRASTKSIEREINSVNDNPLIDVSRNKALHGGNFQ
Aofficinalis-evm.model.AsparagusV1_04.71        PLQKPKQDRYALRTSPQWLGPQIEVIRSSTKSIEREINSVNDNPLIDVSRNKALHGGNFQ
Aofficinalis-evm.model.AsparagusV1_04.93        PLQKPKQDRYALRTSPQWLGPQIEVIRSSTKSIEREINSVNDNPLIDVSRNKALHGGNFQ
Aofficinalis-evm.model.AsparagusV1_10.1948      PLQKPKQDRYALRTSPQWLGPQIEVIRSSTKSIEREINSVNDNPLIDVSRNKALHGGNFQ
Typha_latifolia-Tylat.03G127700.1               ALQKPKQDRYALRTSPQWLGPQIEVIRASTKSIEREINSVNDNPLIDVSRNKALHGGNFQ
Spolyrhiza-Spipo11G0025500                      PLQKPKQDRYALRTSPQWLGPQVEVLRHATKSIEREINSVNDNPLVDVSRNKALHGGNFQ
Spolyrhiza-Spipo1G0003500                       PLQKPKQDRYALRTSPQWLGPQVEVLRQATKSIEREINSVNDNPLIDVSRGKALHGGNFQ
Phoenix_dactylifera-LOC103718264                PLQKPKQDRYALRTSPQWLGPQIEVIRMSTKSIEREINSVNDNPLIDVSRSKALHGGNFQ
Macuminata-GSMUA_Achr1T09070_001                PLQKPKQDRYALRTSPQWLGPQIEVIRASTKSIEREINSVNDNPLIDVSRSKALHGGNFQ
Acomosus-Aco006987.1                            PLQKPKQDRYALRTSPQWLGPQIEVIRSSTRSIEREINSVNDNPLIDVARNKALHGGNFQ
Acomosus-Aco027752.1                            PLQKPKQDRYALRTSPQWLGPQIEVIRSSTRSIEREINSVNDNPLIDVARNKALHGGNFQ
Cscoparia-Csco27g03980.1                        PLQKPKQDRYALRTSPQWLGPQIEVIRAATKSIEREINSVNDNPLIDVSRNKAIHGGNFQ
Cscoparia-Csco27g03990.1                        PLQKPKQDRYALRTSPQWLGPQIEVIRAATKSIEREINSVNDNPLIDVSRNKAIHGGNFQ
Macuminata-GSMUA_Achr11T16380_001               PLQKPKKDRYALRTSPQWLGPQIEVIRSSTKSIEREINSVNDNPLIDVARNKTIYGGNFQ
Macuminata-GSMUA_Achr11T22840_001               PLQKPKQDRYALRTSPQWLGPQIEVIRSATKSIEREINSVNDNPLIDVSRNKALHGGNFQ
Macuminata-GSMUA_Achr2T00240_001                PLQKPKQDRYALRTSPQWLGPQIEVIRSSTKSIEREINSVNDNPLIDVSRNKALHGGNFQ
Macuminata-GSMUA_Achr5T18560_001                PLQKPKQDRYALRTSPQWLGPQIEVIRASTKSIEREINSVNDNPLIDVSRNKALHGGNFQ
Macuminata-GSMUA_Achr8T18960_001                PLQKPKQDRYALRTSPQWLGPQIEVIRSATKSIEREINSVNDNPLIDVSRNKALHGGNFQ
Macuminata-GSMUA_Achr5T03950_001                PLQKPKQDRYALRTSPQWLGPQIEVIRSSTKSIEREINSVNDNPLIDVSRNKALHGGNFQ
Phoenix_dactylifera-LOC103703950                PLQKPKQDRYALRTSPQWLGPQIEVIRSSTKSVEREINSVNDNPLIDVSRNKALHGGNFQ
Acoerulea-Aqcoe1G140100.1                       PLQKPKQDRYALRTSPQWLGPQIEVIRSSTKSIEREINSVNDNPLIDVSRNKAIHGGNFQ
Acoerulea-Aqcoe2G144500.1                       PLQKPKQDRYALRTSPQWLGPQIEVIRTSTKSIEREINSVNDNPLIDVSRNKAIHGGNFQ
Typha_latifolia-Tylat.05G053500.1               PLQKPKQDRYALRTSPQWLGPQIEVIRAATKSIEREINSVNDNPLIDVSRNKALHGGNFQ
Zmarina-Zosma05g29470.1                         PLQKPKQDRYALRTSPQWLGPQVEVIRASTKSIEREINSVNDNPLIDVSRNKALHGGNFQ
Acomosus-Aco007727.1                            PLQKPKQDRYALRTSPQWLGPQIEVIRAATKSIEREINSVNDNPLIDVSRNKALHGGNFQ
Bvulgarisssp-Bevul.7G054300.1                   PLQKPKQDRYALRTSPQWLGPQIEVIRFATKSIEREINSVNDNPLIDVSRNKALHGGNFQ
Bdistachyon-Bradi3g47110.1                      PLMKPKQDRYALRTSPQWLGPQIEVIRAATKSIEREINSVNDNPLIDVSRGKAIHGGNFQ
Bdistachyon-Bradi3g47120.1                      PLMKPKQDRYALRTSPQWLGPQIEVIRAATKSIEREINSVNDNPLIDVSRGKAIHGGNFQ
Bdistachyon-Bradi3g48840.1                      PLMKPKQDRYALRTSPQWLGPQIEVIRAATKSIEREINSVNDNPLIDVSRGKAIHGGNFQ
Bdistachyon-Bradi3g49280.1                      PLMKPKQDRYALRTSPQWLGPQIEVIRAATKSIEREINSVNDNPLIDVSRGKAIHGGNFQ
Hvulgare-HORVU6Hr1G058840.1                     PLMKPKQDRYALRTSPQWLGPQIEVIRAATKSIEREINSVNDNPLIDVSRGKAIHGGNFQ
Taestivum-Traes_6BL_C96F626D9.1                 PLMKPKQDRYALRTSPQWLGPQIEVIRAATKSIEREINSVNDNPLIDVSRGKAIHGGNFQ
Tintermedium-Thint.18G0192700.1                 PLMKPKQDRYALRTSPQWLGPQIEVIRAATKSIEREINSVNDNPLIDVSRGKAIHGGNFQ
Taestivum-Traes_6DL_04CD7E82C.1                 PLMKPKQDRYALRTSPQWLGPQIEVIRAATKSIEREINSVNDNPLIDVSRGKAIHGGNFQ
Bdistachyon-Bradi3g49270.1                      PLMKPKQDRYALRTSPQWLGPQIEVIRAATKSIEREINSVNDNPLIDVSRGKAIHGGNFQ
Ecoracana-ELECO.r07.2AG0138020.1                PLMKPKQDRYALRTSPQWLGPQIEVIRAATKSIEREINSVNDNPLIDVARSKALHGGNFQ
Ecoracana-ELECO.r07.2BG0193670.1                PLMKPKQDRYALRTSPQWLGPQIEVIRAATKSIEREINSVNDNPLIDVARSKALHGGNFQ
Osativa-LOC_Os02g41670.1                        PLMKPKQDRYALRTSPQWLGPQIEVIRAATKSIEREINSVNDNPLIDVSRDKALHGGNFQ
Osativa-LOC_Os02g41680.1                        PLMKPKQDRYALRTSPQWLGPQIEVIRAATKSIEREINSVNDNPLIDVSRGKALHGGNFQ
Pvaginatum-Pavag04G202900.1                     PLMKPKQDRYALRTSPQWLGPQIEVIRAATKSIEREINSVNDNPLIDVARSKALHGGNFQ
Pvaginatum-PavagK321000.1                       PLMKPKQDRYALRTSPQWLGPQIEVIRAATKSIEREINSVNDNPLIDVARSKALHGGNFQ
Pvaginatum-PavagK321300.1                       PLMKPKQDRYALRTSPQWLGPQIEVIRAATKSIEREINSVNDNPLIDVARSKALHGGNFQ
Pvaginatum-PavagK321100.1                       PLMKPKQDRYALRTSPQWLGPQIEVIRAATKSIEREINSVNDNPLIDVARSKALHGGNFQ
Pvaginatum-PavagK321200.1                       PLMKPKQDRYALRTSPQWLGPQIEVIRAATKSIEREINSVNDNPLIDVARSKALHGGNFQ
Pvirgatum-Pavir.1KG386500.1                     PLMKPKQDRYALRTSPQWLGPQIEVIRAATKSIEREINSVNDNPLIDVSRGKALHGGNFQ
Pvirgatum-Pavir.1NG356700.1                     PLMKPKQDRYALRTSPQWLGPQIEVIRAATKSIEREINSVNDNPLIDVARSKALHGGNFQ
Sviridis-Sevir.1G245166.1                       PLMKPKQDRYALRTSPQWLGPQIEVIRAATKSIEREINSVNDNPLIDVARSKALHGGNFQ
Sviridis-Sevir.1G245232.1                       PLMKPKQDRYALRTSPQWLGPQIEVIRAATKSIEREINSVNDNPLIDVARSKALHGGNFQ
Pvirgatum-Pavir.1NG356800.1                     PLMKPKQDRYALRTSPQWLGPQIEVIRAATKSIEREINSVNDNPLIDVARSKALHGGNFQ
Sbicolor-Sobic.004G220500.1                     PLMKPKQDRYALRTSPQWLGPQIEVIRAATKSIEREINSVNDNPLIDVARSKALHGGNFQ
Sbicolor-Sobic.004G220600.2                     PLMKPKQDRYALRTSPQWLGPQIEVIRAATKSIEREINSVNDNPLIDVARSKALHGGNFQ
Sbicolor-Sobic.004G220700.1                     PLMKPKQDRYALRTSPQWLGPQIEVIRAATKSIEREINSVNDNPLIDVARSKALHGGNFQ
Sviridis-Sevir.1G245300.2                       PLMKPKQDRYALRTSPQWLGPQIEVIRAATKSIEREINSVNDNPLIDVARSKALHGGNFQ
Zmays-Zm00001d017276_T001                       PLMKPKQDRYALRTSPQWLGPQIEVIRAATKSIEREINSVNDNPLIDVARSKALHGGNFQ
Zmays-Zm00001d017279_T001                       PLMKPKQDRYALRTSPQWLGPQIEVIRAATKSIEREINSVNDNPLIDVARSKALHGGNFQ
Zmays-Zm00001d051166_T001                       PLMKPKQDRYALRTSPQWLGPQIEVIRASTKSIEREINSVNDNPLIDVARSKALHGGNFQ
Ecoracana-ELECO.r07.2AG0138030.1                PLMKPKQDRYALRTSPQWLGPQIEVIRAATKSIEREINSVNDNPLIDVSRGKALHGGNFQ
Ecoracana-ELECO.r07.2BG0193680.1                PLMKPKQDRYALRTSPQWLGPQIEVIRAATKSIEREINSVNDNPLIDVSRGKALHGGNFQ
Osativa-LOC_Os05g35290.1                        PLMKPKQDRYALRTSPQWLGPQIEVIRAATKSIEREINSVNDNPLIDVSRGKALHGGNFQ
Platifolius-Phala.02G210700.1                   PLMKPKQDRYALRTSPQWLGPQIEVIRAATKSIEREINSVNDNPLIDVSRGKALHGGNFQ
Ecoracana-ELECO.r07.4AG0310420.1                PLMKPKQDRYALRTSPQWLGPQIEVIRAATTSIEREINSVNDNPLIDVSRGKALHGGNFQ
Ecoracana-ELECO.r07.4BG0341610.1                PLMKPKQDRYALRTSPQWLGPQIEVIRAATKSIEREINSVNDNPLIDVSRGKALHGGNFQ
Bdistachyon-Bradi3g49260.1                      PLMKPKQDRYALRTSPQWLGPQIEVIRFATKSIEREINSVNDNPLIDVSRGKAIHGGNFQ
Hvulgare-HORVU0Hr1G016330.1                     PLMKPKQDRYALRTSPQWLGPQIEVIRFATKSIEREINSVNDNPLIDVSRGKAIHGGNFQ
Tintermedium-Thint.17G0260200.1                 PLMKPKQDRYALRTSPQWLGPQIEVIRFATKSIEREINSVNDNPLIDVSRGKAIHGGNFQ
Tintermedium-Thint.18G0192600.1                 PLMKPKQDRYALRTSPQWLGPQIEVIRFATKSIEREINSVNDNPLIDVSRGKAIHGGNFQ
Tintermedium-Thint.17G0231000.1                 PLMKPKQDRYALRTSPQWLGPQIEVIRFATKSIEREINSVNDNPLIDVSRGKAIHGGNFQ
Ecoracana-ELECO.r07.2AG0138010.1                PLMKPKQDRYALRTSPQWLGPQIEVIRFATKSIEREINSVNDNPLIDVSRGKALHGGNFQ
Ecoracana-ELECO.r07.2BG0193660.1                PLMKPKQDRYALRTSPQWLGPQIEVIRFATKSIEREINSVNDNPLIDVSRGKALHGGNFQ
Osativa-LOC_Os02g41650.3                        PLMKPKQDRYALRTSPQWLGPQIEVIRFATKSIEREINSVNDNPLIDVSRGKALHGGNFQ
Pvaginatum-Pavag04G202800.1                     PLMKPKQDRYALRTSPQWLGPQIEVIRFATKSIEREINSVNDNPLIDVSRGKALHGGNFQ
Pvaginatum-PavagK320900.1                       PLMKPKQDRYALRTSPQWLGPQIEVIRFATKSIEREINSVNDNPLIDVSRGKALHGGNFQ
Pvirgatum-Pavir.1NG356400.1                     PLMKPKQDRYALRTSPQWLGPQIEVIRFATKSIEREINSVNDNPLIDVSRGKALHGGNFQ
Sviridis-Sevir.1G245100.1                       PLMKPKQDRYALRTSPQWLGPQIEVIRFATKSIEREINSVNDNPLIDVSRGKALHGGNFQ
Sbicolor-Sobic.004G220400.1                     PLMKPKQDRYALRTSPQWLGPQIEVIRFATKSIEREINSVNDNPLIDVSRGKALHGGNFQ
Zmays-Zm00001d017275_T001                       PLMKPKQDRYALRTSPQWLGPQIEVIRFATKSIEREINSVNDNPLIDVSRGKALHGGNFQ
Zmays-Zm00001d051163_T002                       PLMKPKQDRYALRTSPQWLGPQIEVIRFATKSIEREINSVNDNPLIDVSRGKALHGGNFQ
Pvirgatum-Pavir.1KG386300.1                     PLMKPKQDRYALRTSPQWLGPQIEVIRFATKSIEREINSVNDNPLIDVSRGKALHGGNFQ
Platifolius-Phala.02G210600.1                   PLMKPKQDRYALRTSPQWLGPQIEVIRAATKSIEREINSVNDNPLIDVSRGKALHGGNFQ
Streptochaeta-strangu_020767-RA                 PLMKPKQDRYALRTSPQWLGPQIEVIRAATKSIEREINSVNDNPLIDVSRGKALHGGNFQ
Bdistachyon-Bradi5g15830.1                      PLMKPKQDRYALRTSPQWLGPQIEVIRAATKSIEREINSVNDNPLIDVSRGKAIHGGNFQ
Hvulgare-HORVU2Hr1G089440.4                     PLMKPKQDRYALRTSPQWLGPQIEVIRAATKSIEREINSVNDNPLIDVSRGKAIHGGNFQ
Taestivum-Traes_2AL_0DDC60D6E.1                 PLMKPKQDRYALRTSPQWLGPQIEVIRAATKSIEREINSVNDNPLIDVSRGKAIHGGNFQ
Tintermedium-Thint.06G0475900.1                 PLMKPKQDRYALRTSPQWLGPQIEVIRAATKSIEREINSVNDNPLIDVSRGKAIHGGNFQ
Tintermedium-Thint.06G0476200.1                 PLMKPKQDRYALRTSPQWLGPQIEVIRAATKSIEREINSVNDNPLIDVSRGKAIHGGNFQ
Taestivum-Traes_2AL_2763280FC.1                 PLMKPKQDRYALRTSPQWLGPQIEVIRAATKSIEREINSVNDNPLIDVSRGKAIHGGNFQ
```

| | |
|---|---|
| Taestivum-Traes_2DL_8C21AAAEF.1 | PLMKPKQDRYALRTSPQWLGPQIEVIRASTKSIEREINSVNDNPLIDVSRGKAIHGGNFQ |
| Tintermedium-Thint.V0383500.1 | PLMKPKQDRYALRTSPQWLGPQIEVIRAATKSIEREINSVNDNPLIDVSRGKAIHGGNFQ |
| Taestivum-Traes_2AL_9EC3226F7.1 | PLMKPKQDRYALRTSPQWLGPQIEVIRAATKSIEREINSVNDNPLIDVSRGKAIHGGNFQ |
| Tintermedium-Thint.05G0273800.1 | PLMKPKQDRYALRTSPQWLGPQIEVIRAATKSIEREINSVNDNPLIDVSRGKAIHGGNFQ |
| Tintermedium-Thint.V0383400.1 | PLMKPKQDRYALRTSPQWLGPQIEVIRAATKSIEREINSVNDNPLIDVSRGKAIHGGNFQ |
| Taestivum-Traes_2BL_13D5272D7.1 | PLMKPKQDRYALRTSPQWLGPQIEVIRAATKSIEREINSVNDNPLIDVSRGKAIHGGNFQ |
| Tintermedium-Thint.06G0475700.1 | PLMKPKQDRYALRTSPQWLGPQIEVIRAATKSIEREINSVNDNPLIDVSRGKAIHGGNFQ |
| Taestivum-Traes_2BL_C051606EA.2 | PLMKPKQDRYALRTSPQWLGPQIEVIRAATKSIEREINSVNDNPLIDVSRGKAIHGGNFQ |
| Tintermedium-Thint.V0383600.1 | PLMKPKQDRYALRTSPQWLGPQIEVIRAATKSIEREINSVNDNPLIDVSRGKAIHGGNFQ |
| Taestivum-Traes_2AL_9D78F85E2.1 | PLMKPKQDRYALRTSPQWLGPQIEVIGAATKSIEREINSVNDNPLIDVSRGKAIHGGNFQ |
| Tintermedium-Thint.05G0274100.1 | PLMKPKQDRYALRTSPQWLGPQIEVIRAATKSIEREINSVNDNPLIDVSRGKAIHGGNFQ |
| Tintermedium-Thint.06G0476300.1 | PLMKPKQDRYALRTSPQWLGPQIEVIRAATKSIEREINSVNDNPLIDVSRGKAIHGGNFQ |
| Taestivum-Traes_2BL_5A50FDA1A.1 | PLMKPKQDRYALRTSPQWLGPQIEVIRAATKSIEREINSVNDNPLIDVSRDKAIHGGNFQ |
| Tintermedium-Thint.05G0273900.1 | PLMKPKQDRYALRTSPQWLGPQIEVIRAATKSIEREINSVNDNPLIDVSRGKAIHGGNFQ |
| Taestivum-Traes_2DL_30F8555D4.1 | ------------------------------------------------------------ |
| Taestivum-Traes_2DL_83168C1E0.1 | ------------------------------------------------------------ |
| Taestivum-Traes_2DL_B978B4E22.1 | ------------------------------------------------------------ |
| Taestivum-Traes_1AS_0B9295A68.1 | PLMKPKQDRYALRTSPQWLGPQIEVIRAATKSIEREINSVNDNPLIDVSRGKAIHGGNFQ |
| Taestivum-Traes_1BS_7068F56E6.1 | PLMKPKQDRYALRTSPQWLGPQIEVIRAATKSIEREINSVNDNPLIDVSRGKAIHGGNFQ |
| Taestivum-Traes_1BS_AEA817616.1 | PLMKPKQDRYALRTSPQWLGPQIEVIRAATKSIEREINSVNDNPLIDVSRGKAIHGGNFQ |
| Taestivum-Traes_4AL_892C47ED5.1 | PLMKPKQDRYALRTSPQWLGPQIEVIRAATKSIEREINSVNDNPLIDVSRGKAIHGGNFQ |
| Tintermedium-Thint.03G0055900.1 | PLMKPKQDRYALRTSPQWLGPQIEVIRAATKSIEREINSVNDNPLIDVSRGKAIHGGNFQ |
| Tintermedium-Thint.01G0012900.1 | PLMKPKQDRYALRTSPQWLGPQIEVIRAATKSIEREINSVNDNPLIDVSRGKAIHGGNFQ |
| Tintermedium-Thint.02G0158500.1 | PLMKPKQDRYALRTSPQWLGPQIEVIRAATKSIEREINSVNDNPLIDVSRGKAIHGGNFQ |
| Tintermedium-Thint.03G0055700.1 | PLMKPKQDRYALRTSPQWLGPQIEVIRAATKSIEREINSVNDNPLIDVSRGKAIHGGNFQ |
| Taestivum-Traes_1BS_C3AA04888.1 | PLMKPKQDRYALRTSPQWLGPQIEVIRAATKSIEREINSVNDNPLIDVSRGKAIHGGNFQ |
| Taestivum-Traes_1DS_A8BD91E4A.1 | PLMKPKQDRYALRTSPQWLGPQIEVIRAATKSIEREINSVNDNPLIDVSRGKAIHGGNFQ |
| Taestivum-Traes_1BS_C26CCE0E9.1 | PLMKPKQDRYALRTSPQWLGPQIEVIRAATKSIEREINSVNDNPLIDVSRGKAIHGGNFQ |
| Taestivum-Traes_1DS_A21FCBA3D.1 | PLMKPKQDRYALRTSPQWLGPQVEVIRAATKSIEREINSVNDNPLIDVSRGKAIHGGNFQ |
| Tintermedium-Thint.01G0012200.1 | PLMKPKQDRYALRTSPQWLGPQIEVIRAATKSIEREINSVNDNPLIDVSRGKAIHGGNFQ |
| Tintermedium-Thint.02G0158600.1 | PLMKPKQDRYALRTSPQWLGPQIEVIRAATKSIEREINSVNDNPLIDVSRGKAIHGGNFQ |
| Tintermedium-Thint.01G0012700.1 | PLMKPKQDRYALRTSPQWLGPQIEVIRAATKSIEREINSVNDNPLIDVSRGKAIHGGNFQ |
| Taestivum-Traes_5BL_F50A47503.1 | PLMKPKQDRYALRTSPQWLGPQIEVIRAATKSIEREINSVNDNPLIDVSRGKAIHGGNFQ |
| Tintermedium-Thint.02G0158700.1 | PLMKPKQDRYALRTSPQWLGPQIEVIRAATKSIEREINSVNDNPLIDVSRGKAIHGGNFQ |
| Taestivum-Traes_5BL_AB528184A.1 | PLMKPKQDRYALRTSPQWLGPQIEVIRAATKSIEREINSVNDNPLIDVSRGKAIHGGNFQ |
| Tintermedium-Thint.03G0055800.1 | PLMKPKQDRYALRTSPQWLGPQIEVIRAATKSIEREINSVNDNPLIDVSRGKAIHGGNFQ |
| Taestivum-Traes_1AS_F9013A945.1 | PLMKPKQDRYALRTSPQWLGPQIEVIRAATKSIEREINSVNDNPLIDVSRGKAIHGGNFQ |
| Taestivum-Traes_1DS_43DAC7A7F.1 | PLMKPKQDRYALRTSPQWLGPQIEVIRAATKSIEREINSVNDNPLIDVSRGKAIHGGNFQ |
| Taestivum-Traes_1BS_BD86C90A7.1 | PLMKPKQDRYALRTSPQWLGPQIEVIRAATKSIEREINSVNDNPLIDVSRGKAIHGGNFQ |
| Tintermedium-Thint.02G0158900.1 | PLMKPKQDRYALRTSPQWLGPQIEVIRAATKSIEREINSVNDNPLIDVSRGKAIHGGNFQ |
| Tintermedium-Thint.05G0274000.1 | PLMKPKQDRYALRTSPQWLGPQIEVIRAATKSIEREINSVNDNPLIDVSRGKAIHGGNFQ |
| Osativa-LOC_Os04g43800.1 | PLMKPKQDRYALRTSPQWLGPQIEVIRAATKSIEREINSVNDNPLIDVSRGKALHGGNFQ |
| Pvaginatum-Pavag06G160300.1 | PLMKPKQDRYALRTSPQWLGPQVEVIRAATKSIEREINSVNDNPLIDVSRGKALHGGNFQ |
| Sbicolor-Sobic.006G148900.1 | PLMKPKQDRYALRTSPQWLGPQIEVIRAATKSIEREINSVNDNPLIDVSRGKALHGGNFQ |
| Pvirgatum-Pavir.7KG237800.1 | PLMKPKQDRYALRTSPQWLGPQIEVIRAATKSIEREINSVNDNPLIDVSRGKALHGGNFQ |
| Pvirgatum-Pavir.7NG355800.1 | PLMKPKQDRYALRTSPQWLGPQIEVIRAATKSIEREINSVNDNPLIDVSRGKALHGGNFQ |
| Sviridis-Sevir.7G178300.1 | PLMKPKQDRYALRTSPQWLGPQIEVIRAATKSIEREINSVNDNPLIDVSRGKALHGGNFQ |
| Zmays-Zm00001d003015_T001 | PLMKPKQDRYALRTSPQWLGPQIEVIRSATKSIEREINSVNDNPLIDVSRGKALHGGNFQ |
| Hvulgare-HORVU1Hr1G022060.1 | PLMKPKQDRYALRTSPQWLGPQIEVIRAATKSIEREINSVNDNPLIDVSRGKAIHGGNFQ |
| Taestivum-Traes_1BS_723922D171.2 | PLMKPKQDRYALRTSPQWLGPQIEVIRAATKSIEREINSVNDNPLIDVSRGKAIHGGNFQ |
| Taestivum-Traes_1DS_A171C7D59.1 | PLMKPKQDRYALRTSPQWLGPQIEVIRAATKSIEREINSVNDNPLIDVSRGKAIHGGNFQ |
| Tintermedium-Thint.01G0111500.1 | PLMKPKQDRYALRTSPQWLGPQIEVIRAATKSIEREINSVNDNPLIDVSRGKAIHGGNFQ |
| Tintermedium-Thint.03G0265300.1 | PLMKPKQDRYALRTSPQWLGPQIEVIRAATKSIEREINSVNDNPLIDVSRGKAIHGGNFQ |
| Hvulgare-HORVU2Hr1G038140.1 | PLMKPKQDRYALRTSPQWLGPQIEVIRAATKSIEREINSVNDNPLIDVSRGKAIHGGNFQ |
| Tintermedium-Thint.04G0171200.1 | PLMKPKQDRYALRTSPQWLGPQIEVIRAATKSIEREINSVNDNPLIDVSRGKAIHGGNFQ |
| Taestivum-Traes_2AS_958327519.3 | PLMKPKQDRYALRTSPQWLGPQIEVIRAATKSIEREINSVNDNPLIDVSRGKAIHGGNFQ |
| Taestivum-Traes_2DS_28CA50371.1 | PLMKPKQDRYALRTSPQWLGPQIEVIRAATKSIEREINSVNDNPLIDVSRGKAIHGGNFQ |
| Tintermedium-Thint.05G0160800.1 | PLMKPKQDRYALRTSPQWLGPQIEVIRAATKSIEREINSVNDNPLIDVSRGKAIHGGNFQ |
| Tintermedium-Thint.06G0299000.1 | PLMKPKQDRYALRTSPQWLGPQIEVIRAATKSIEREINSVNDNPLIDVSRGKAIHGGNFQ |
| Taestivum-Traes_2DS_3791A8A36.1 | PLMKPKQDRYALRTSPQWLGPQIEVIRAATKSIEREINSVNDNPLIDVSRGKAIHGGNFQ |
| Tintermedium-Thint.06G0342800.1 | PLMKPKQDRYALRTSPQWLGPQIEVIRAATKSIEREINSVNDNPLIDVSRGKAIHGGNFQ |
| Tintermedium-Thint.05G0150900.1 | PLMKPKQDRYALRTSPQWLGPQIEVIRAATKSIEREINSVNDNPLIDVSRGKAIHGGNFQ |
| Tintermedium-Thint.04G0242300.1 | PLMKPKQDRYALRTSPQWLGPQIEVIRAATKSIEREINSVNDNPLIDVSRGKAIHGGNFQ |
| Tintermedium-Thint.06G0298700.1 | PLMKPKQDRYALRTSPQWLGPQIEVIRAATKSIEREINSVNDNPLIDVSRGKAIHGGNFQ |
| Taestivum-Traes_2BS_88CF42F2E.1 | PLMKPKQDRYALRTSPQWLGPQIEVIRAATKSIEREINSVNDNPLIDVSRGKAIHGGNFQ |
| Tintermedium-Thint.04G0241900.1 | PLMKPKQDRYALRTSPQWLGPQIEVIRAATKSIEREINSINDNPLIDVSRGKAIHGGNFQ |
| Platifolius-Phala.06G165300.1 | PLMKPKQDRYALRTSPQWLGPQIEVIRAATKSIEREINSVNDNPLIDVSRGKALHGGNFQ |
| Streptochaeta-strangu_019385-RA | PLMKPKQDRYALRTSPQWLGSQIEVIRAATKSIEREINSVNDNPLIDVSRGKALHGGNFQ |
| Streptochaeta-strangu_028944-RA | PLTKPKQDRYALRSSPQWLGPQVEVIRAATKSIEREINSVSDNPLIDVSRDKALHCGNFQ |
| Acomosus-Aco010091.1 | PLQKPKKDRYAVCTSPQWLGPQIEVIRSSTKSIEREINSVNDNPLIDVSRDKALHGGNFQ |
| Zmarina-Zosma03g22300.1 | PLQKPKQDRYALRTSPQWLGPQIEVIRASTKSIEREINSVNDNPLIDVKRGKAIHGGNFQ |
| Ecdeiocolea-Emoptg0003741_1G000600.1 | PLMKPKQDRYALRTSPQWLGPQIEVIRAATKSIEREINSVNDNPLIDVSRGKALHGGNFQ |
| Jascendens-Joasc.05G060500.1 | PLMKPKQDRYALRTSPQWLGPQIEVIRASTKSIEREINSVNDNPLIDVSRGKALHGGNFQ |
| Ecdeiocolea-Emoptg0003741_1G000630.1 | PLMKPKQDRYALRTSPQWLGPQIEVIRTATKSIEREINSVNDNPVIDVSRGKALHGGNFQ |
| Jascendens-Joasc.05G060400.1 | PLLKPKQDRYALRTAPQWLGPQIEVIRAATKSIEREINSVNDNPIIDVSRGKALHGGNFQ |
| Bdistachyon-Bradi3g49250.2 | PQLKPKQDRYALRTSPQWLGPQIEVIRSATKSIEREVNSVNDNPVIDVHRGKALHGGNFQ |
| Hvulgare-HORVU6Hr1G058820.1 | TQLKPKQDRYALRTSPQWLGPQIEVIRSATKSIEREVNSVNDNPVIDVHRGKALHGGNFQ |
| Taestivum-Traes_6DL_1AEA7B869.1 | TQLKPKQDRYALRTSPQWLGPQIEVIRSATKSIEREVNSVNDNPVIDVHRGKALHGGNFQ |
| Tintermedium-Thint.18G0192300.1 | TQLKPKQDRYALRTSPQWLGPQIEVIRSATKSIEREVNSVNDNPVIDVHRGKALHGGNFQ |
| Tintermedium-Thint.17G0230800.1 | TQLKPKQDRYALRTSPQWLGPQIEVIRSATKSIEREVNSVNDNPVIDVHRGKALHGGNFQ |
| Tintermedium-Thint.17G0230900.1 | TQLKPKQDRYALRTSPQWLGPQIEVIRSATKSIEREVNSVNDNPVIDVHRGKALHGGNFQ |
| Ecoracana-ELECO.r07.2AG0138000.1 | PLLKPKQDRYALRTSPQWLGPQIEVIRAATKSIEREVNSVNDNPVIDVHRGKALHGGNFQ |
| Ecoracana-ELECO.r07.2BG0193650.1 | PLLKPKQDRYALRTSPQWLGPQIEVIRAATKSIEREVNSVNDNPVIDVHRGKALHGGNFQ |
| Pvaginatum-Pavag04G202700.1 | PLLKPKQDRYALRTSPQWLGPQIEVIRAATKSIEREVNSVNDNPVIDVHRGKALHGGNFQ |
| Sviridis-Sevir.1G245000.1 | PLLKPKQDRYALRTSPQWLGPQIEVIRAATKSIEREVNSVNDNPVIDVHRGKALHGGNFQ |
| Sviridis-Sevir.6G187100.1 | PLLKPKQDRYALRTSPQWLGPQIEVIRAATKSIEREVNSVNDNPVIDVHRGKALHGGNFQ |
| Pvirgatum-Pavir.1NG356200.1 | PLLKPKQDRYALRTSPQWLGPQIEVIRAATKSIEREVNSVNDNPVIDVHRGKALHGGNFQ |
| Sbicolor-Sobic.004G220300.1 | PLLKPKQDRYALRTSPQWLGPQIEVLRAATKSIEREVNSVNDNPVIDVHRGKALHGGNFQ |
| Zmays-Zm00001d017274_T001 | PLLKPKQDRYALRTSPQWLGPQIEVIRAATKSIEREVNSVNDNPVIDVHRGKALHGGNFQ |
| Zmays-Zm00001d051161_T003 | PLLKPKQDRYALRTSPQWLGPQIEVIRAATKSIEREVNSVNDNPVIDVHRGKALHGGNFQ |
| Sviridis-Sevir.7G178200.1 | PLLKPKQDRYALRTSPQWLGPQIEVIRAATKSIEREVNSVNDNPVIDVHRGKALHGGNFQ |
| Osativa-LOC_Os02g41630.2 | PLLKPKQDRYALRTSPQWLGPQIEVIRAATKSIEREVNSVNDNPVIDVHRGKALHGGNFQ |
| Ecoracana-ELECO.r07.4AG0310430.1 | PLLKPKQDRYALRTSPQWLGPQIEVIRAATKSIEREVNSVNDNPVIDVHRGKALHGGNFQ |
| Ecoracana-ELECO.r07.4BG0341620.1 | PLLKPKQDRYALRTSPQWLGPQIEVIRAATKSIEREVNSVNDNPVIDVHRGKALHGGNFQ |
| Pvaginatum-Pavag06G160200.1 | PLLKPKQDRYALRTSPQWLGPQIEVIRAATKSIEREVNSVNDNPVIDVHRGKALHGGNFQ |
| Pvirgatum-Pavir.7KG238255.1 | PLLKPKQDRYALRTSPQWLGPQIEVIRAATKSIEREVNSVNDNPVIDVHRGKALHGGNFQ |
| Sbicolor-Sobic.006G148800.1 | PLLKPKQDRYALRTSPQWLGPQIEVIRAATKSIEREVNSVNDNPVIDVHRGKALHGGNFQ |
| Zmays-Zm00001d003016_T001 | PLLKPKQDRYALRTSPQWLGPQIEVIRAATKSIEREVNSVNDNPVIDVHRGKALHGGNFQ |
| Osativa-LOC_Os04g43760.1 | PLLKPKQDRYALRTSPQWLGPQIEVIRAATKSIEREVNSVNDNPVIDVHRGKALHGGNFQ |
| Platifolius-Phala.02G210400.1 | PLLKPKQDRYALRTSPQWLGPQIEVIRSATKSIEREVNSVNDNPVIDVHRGKALHGGNFQ |
| Platifolius-Phala.06G165100.1 | PLLKPKQDRYALRTSPQWLGPQIEVIRAATKSIEREVNSVNDNPVIDVHRGKALHGGNFQ |
| Streptochaeta-strangu_019386-RA | PLLKPKQDRYALRTSPQWLGPQIEVIRAATKSIEREVNSVNDNPVIDVSRGKALHGGNFQ |
| Streptochaeta-strangu_020769-RA | PLLKPKQDRYALRTAPQWLGPQIEVIRAATKSIEREVNSVNDNPVIDVKRGKALHGGNFQ |
| Sviridis-Sevir.2G448300.1 | PLLKPKQDRYALRTSPQWLGPQIEVIRAATKAIEREINSVSDNPVIDVNRGKALHGGNFQ |
| Sviridis-Sevir.7G177900.1 | PLLKPKQDQYALRTSPQWLGPQIEVIRAATKAIEREINSVSDNPVIDVHRGKALHGGNFQ |

| | |
|---|---|
| Aamericanus-Acora.01G232100.1 | GTPIGVSMDNTRLAIAAIGKLMFAQFSELVNDFYNNGLPSNLSGGRNPSLDYGFKGAEIA |
| Aamericanus-Acora.10G174800.1 | GTPIGVSMDNTRLAIAAIGKLMFAQFSELVNDFYNNGLPSNLSGGRNPSLDYGFKGAEIA |
| Acomosus-Aco013943.1 | GTPIGVSMDNTRLAIAAIGKLMFAQLSELVNDFYNNGLPSNLSGGRNPSLDYGFKGAEIA |
| Aofficinalis-evm.model.AsparagusV1_01.827 | GTPIGVSMDNTRLAIAAIGKLMFAQFSELVNDLYNNGLPSNLSGGRNPSLDYGFKGAEIA |
| Typha_latifolia-Tylat.14G083800.1 | GTPIGVSMDNTRLAIAAIGKLMFAQFSELVNDFYNNGLPSNLSGGRNPSLDYGFKGAEIA |
| Aofficinalis-evm.model.AsparagusV1_04.71 | GTPIGVSMDNTRLAIAAIGKLMFAQVSELVNDFYNNGLPSNLSGGRNPSLDYGFKGAEIA |
| Aofficinalis-evm.model.AsparagusV1_04.93 | GTPIGVSMDNTRLAIAAIGKLMFAQVSELVNDFYNNGLPSNLSGGRNPSLDYGFKGAEIA |
| Aofficinalis-evm.model.AsparagusV1_10.1948 | GTPIGVSMDNTRLAIAAIGKLMFAQLSELVNDFYNNGLPSNLSGGRNPSLDYGFKGAEIA |
| Typha_latifolia-Tylat.03G127700.1 | GTPIGVSMDNTRLAIAAIGKLMFAQFSELVNDFYNNGLPSNLSGGRNPSLDYGFKGAEIA |
| Spolyrhiza-Spipo11G0025500 | GTPIGVSMDNARLALAAIGKLMFAQLSELVNDFYNNGLPSNLSGGRNPSLDYGFKGAEIA |
| Spolyrhiza-Spipo1G0003500 | GTPIGVSMDNSRLAIAAIGKLMFAQISELVNDFYNNGLPSNLTAGRNPSLDYGFKGAEIA |
| Phoenix_dactylifera-LOC103718264 | GTPIGVSMDNTRLAIAAIGKLMFAQFSELVNDFYNNGLPSNLSGGRNPSLDYGFKGAEIA |
| Macuminata-GSMUA_Achr1T09070_001 | GTPIGVSMDNTRLAVAAIGKLMFAQFSELVNDFYNNGLPSNLSGGRNPSLDYGFKGAEIA |
| Acomosus-Aco006987.1 | GTPIGVSMDNTRLALAAIGKLMFAQFSELVNDFYNNGLPSNLSGGRNPSLDYGFKGAEIA |
| Acomosus-Aco027752.1 | GTPIGVSMDNTRLALAAIGKLMFAQFSELVNDFYNNGLPSNLSGGRNPSLDYGFKGAEIA |
| Cscoparia-Csco27g03980.1 | GTPIGVSMDNTRLAIAAIGKLLFAQFSELVNDFYNNGLPSNLSGGRNPSLDYGFKGAEIA |
| Cscoparia-Csco27g03990.1 | GTPIGVSMDNTRLAIAAIGKLLFAQFSELVNDFYNNGLPSNLSGGRNPSLDYGFKGAEIA |
| Macuminata-GSMUA_Achr11T16380_001 | GTPIGVSMDNARLALAAVGKLMFAQFSELVNDLYNNGLPSNLSGGRNPSLDYGFKGAEIA |
| Macuminata-GSMUA_Achr11T22840_001 | GTPIGVSMDNARLALAAIGKLMFAQFSELVNDFYNNGLPSNLSGGRNPSLDYGFKGAEIA |
| Macuminata-GSMUA_Achr2T00240_001 | GTPIGVSMDNTRLALAAIGKLMFAQFSELVNDFYNNGLPSNLSGGRTPSLDYGFKGAEIA |
| Macuminata-GSMUA_Achr5T18560_001 | GTPIGVSMDNTRLAVAAIGKLMFAQFSELVNDFYNNGLPSNLSGGRNPSLDYGFKGAEIA |

```
Macuminata-GSMUA_Achr8T18960_001          GTPIGVSMDNARLALATIGKLLFAQFSELDNDLYNNGLPSNLSGGRNPSLDYGFKGAEIA
Macuminata-GSMUA_Achr5T03950_001          GTPIGVSMDNTRLAIAAIGKLMFAQFSELVNDFYNNGLPSNLSGGRNPSLDYGFKGAEIA
Phoenix_dactylifera-LOC103703950          GTPVGVSMDNTRLAIAAIGKLMFAQFSELVNDFYNNGLPSNLSGGRNPSLDYGFKGAEIA
Acoerulea-Aqcoe1G140100.1                 GTPIGVSMDNTRLAIASIGKLMFAQFSELVNDFYNNGLPSNLSGGRNPSLDYGFKGAEIA
Acoerulea-Aqcoe2G144500.1                 GTPIGVSMDNTRLAIAAIGKLMFAQFSELVNDFYNNGLPSNLSGGRNPSLDYGFKGAEIA
Typha_latifolia-Tylat.05G053500.1         GTPIGVSMDNTRLAIAAIGKLIFAQLSELVNDFYNNGLPSNLSGGRNPSLDYGFKGAEIA
Zmarina-Zosma05g29470.1                   GTPIGVSMDNTRLAIAAIGKLMFAQFSELVNDFYNNGLPSNLSGGRNPSLDYGFKGGEIA
Acomosus-Aco007727.1                      GTPIGVSMDNTRLAIAAIGKLMFAQFSELVNDFYNNGLPSNLSGGRNPSLDYGFKGAEIA
Bvulgarisssp-Bevul.7G054300.1             GTPIGVSMDNARLAIAAIGKLLFAQFSELVNDYYNNGLPSNLTASRDPSLDYGFKGAEIA
Bdistachyon-Bradi3g47110.1                GTPIGVSMDNTRLALAAIGKLMFAQFSELVNDFYNNGLPSNLSGGRNPSLDYGFKGAEIA
Bdistachyon-Bradi3g47120.1                GTPIGVSMDNTRLALAAIGKLMFAQFSELVNDFYNNGLPSNLSGGRNPSLDYGFKGAEIA
Bdistachyon-Bradi3g48840.1                GTPIGVSMDNTRLALAAIGKLMFAQFSELVNDFYNNGLPSNLSGSRNPSLDYGFKGAEIA
Bdistachyon-Bradi3g49280.1                GTPIGVSMDNTRLALAAIGKLMFAQFSELVNDFYNNGLPSNLSGSRNPSLDYGFKGAEIA
Hvulgare-HORVU6Hr1G058840.1               GTPIGVSMDNTRLAIAAIGKLMFAQFSELVNDLYNNGLPSNLSGGRNPSLDYGFKGAEIA
Taestivum-Traes_6BL_C96F626D9.1           GTPIGVSMDNTRLAIAAIGKLMFAQFSELVNDLYNNGLPSNLSGGRNPSLDYGFKGAEIA
Tintermedium-Thint.18G0192700.1           GTPIGVSMDNTRLAIAAIGKLMFAQFSELVNDLYNNGLPSNLSGGRNPSLDYGFKGAEIA
Taestivum-Traes_6DL_04CD7E82C.1           GTPIGVSMDNTRLAIAAIGKLMFAQFSELVNDLYNNGLPSNLSGGHNPSLDYGFKGAEIA
Bdistachyon-Bradi3g49270.1                GTPIGVSMDNTRLALAAIGKLMFAQFSELVNDLYNNGLPSNLSGGRNPSLDYGLKGAEIA
Ecoracana-ELECO.r07.2AG0138020.1          GTPIGVSMDNTRLAIAAIGKLMFAQFSELVNDFYNNGLPSNLSGGRNPSLDYGFKGAEIA
Ecoracana-ELECO.r07.2BG0193670.1          GTPIGVSMDNTRLAIAAIGKLMFAQFSELVNDFYNNGLPSNLSGGRNPSLDYGFKGAEIA
Osativa-LOC_Os02g41670.1                  GTPIGVSMDNTRLAIAAIGKLMFAQFSELVNDYYNNGLPSNLSGGRNPSLDYGFKGAEIA
Osativa-LOC_Os02g41680.1                  GTPIGVSMDNTRLAIAAIGKLMFAQFSELVNDFYNNGLPSNLSGGRNPSLDYGFKGAEIA
Pvaginatum-Pavag04G202900.1               GTPIGVSMDNTRLAIAAIGKLMFAQFSELVNDYYNNGLPSNLSGGRNPSLDYGFKGAEIA
Pvaginatum-PavagK321000.1                 GTPIGVSMDNTRLAIAAIGKLMFAQFSELVNDYYNNGLPSNLSGGRNPSLDYGFKGAEIA
Pvaginatum-PavagK321300.1                 GTPIGVSMDNTRLAIAAIGKLMFAQFSELVNDYYNNGLPSNLSGGRNPSLDYGFKGAEIA
Pvaginatum-PavagK321100.1                 GTPIGVSMDNTRLAIAAIGKLMFAQFSELVNDYYNNGLPSNLSGGRNPSLDYGFKGAEIA
Pvaginatum-PavagK321200.1                 GTPIGVSMDNTRLAIAAIGKLMFAQFSELVNDYYNNGLPSNLSGGRNPSLDYGFKGAEIA
Pvirgatum-Pavir.1KG386500.1               GTPIGVSMDNTRLALAAIGKLMFAQFSELVNDYYNNGLPSNLSGGRNPSLDYGFKGAEIA
Pvirgatum-Pavir.1NG356700.1               GTPIGVSMDNTRLALAAIGKLMFAQFSELVNDYYNNGLPSNLSGGRNPSLDYGFKGAEIA
Sviridis-Sevir.1G245166.1                 GTPIGVSMDNTRLAIAAIGKLMFAQFSELVNDYYNNGLPSNLSGGRNPSLDYGFKGAEIA
Sviridis-Sevir.1G245232.1                 GTPIGVSMDNTRLAIAAIGKLMFAQFSELVNDYYNNGLPSNLSGGRNPSLDYGFKGAEIA
Pvirgatum-Pavir.1NG356800.1               GTPIGVSMDNTRLALAAIGKLMFAQFSELVNDYYNNGLPSNLSGGRNPSLDYGFKGAEIA
Sbicolor-Sobic.004G220500.1               GTPIGVSMDNTRLAIAAIGKLMFAQFSELVNDYYNNGLPSNLSGGRNPSLDYGFKGAEIA
Sbicolor-Sobic.004G220600.2               GTPIGVSMDNTRLAIAAIGKLMFAQFSELVNDYYNNGLPSNLSGGRNPSLDYGFKGAEIA
Sbicolor-Sobic.004G220700.1               GTPIGVSMDNTRLAIAAIGKLMFAQFSELVNDYYNNGLPSNLSGGRNPSLDYGFKGAEIA
Sviridis-Sevir.1G245300.2                 GTPIGVSMDNTRLALAAIGKLMFAQFSELVNDYYNNGLPSNLSGGRNPSLDYGFKGAEIA
Zmays-Zm00001d017276_T001                 GTPIGVSMDNTRLAIAAIGKLMFAQFSELVNDYYNNGLPSNLSGGRNPSLDYGFKGAEIA
Zmays-Zm00001d017279_T001                 GTPIGVSMDNTRLAIAAIGKLMFAQFSELVNDYYNNGLPSNLSGGRNPSLDYGFKGAEIA
Zmays-Zm00001d051166_T001                 GTPIGVSMDNTRLAVAAIGKLMFAQFSELVNDYYNNGLPSNLSGGRNPSLDYGFKGAEIA
Ecoracana-ELECO.r07.2AG0138030.1          GTPIGVSMDNTRLAIAAMGKLMFAQFSELVNDFYNNGLPSNLSGGRNPSLDYGFKGAEIA
Ecoracana-ELECO.r07.2BG0193680.1          GTPIGVSMDNTRLAIAAIGKLMFAQFSELVNDFYNNGLPSNLSGGRNPSLDYGFKGAEIA
Osativa-LOC_Os05g35290.1                  GTPIGVSMDNTRLAIAAIGKLMFAQFSELVNDFYNNGLPSNLSGGRNPSLDYGFKGAEIA
Platifolius-Phala.02G210700.1             GTPIGVSMDNTRLAIAAIGKLMFAQFSELVNDFYNNGLPSNLSGGRNPSLDYGFKGAEIA
Ecoracana-ELECO.r07.4AG0310420.1          GTPIGVSMDNTRLAVAAIGKLMFAQFSELVNDFYNNGLPSNLSGGRNPSLDYGFKGAEIA
Ecoracana-ELECO.r07.4BG0341610.1          GTPIGVSMDNTRLAVAAIGKLMFAQFSELVNDFYNNGLPSNLSGGRNPSLDYGFKGAEIA
Bdistachyon-Bradi3g49260.1                GTPIGVSMDNTRLAIAAIGKLMFAQFSELVNDFYNNGLPSNLSGGRNPSLDYGFKGAEIA
Hvulgare-HORVU0Hr1G016330.1               GTPIGVSMDNTRLAIAAIGKLMFAQFSELVNDFYNNGLPSNLSGGRNPSLDYGFKGAEIA
Tintermedium-Thint.17G0260200.1           GTPIGVSMDNTRLAIAAIGKLMFAQFSELVNDFYNNGLPSNLSGGRNPSLDYGFKGAEIA
Tintermedium-Thint.18G0192600.1           GTPIGVSMDNTRLAIAAIGKLMFAQFSELVNDFYNNGLPSNLSGGRNPSLDYGFKGAEIA
Tintermedium-Thint.17G0231000.1           GTPIGVSMDNTRLAIAAIGKLMFAQFSELVNDFYNNGLPSNLSGGRNPSLDYGFKGAEIA
Ecoracana-ELECO.r07.2AG0138010.1          GTPIGVSMDNTRLALAAIGKLMFAQFSELVNDFYNNGLPSNLSGGRNPSLDYGFKGAEIA
Ecoracana-ELECO.r07.2BG0193660.1          GTPIGVSMDNTRLALAAIGKLMFAQFSELVNDFYNNGLPSNLSGGRNPSLDYGFKGAEIA
Osativa-LOC_Os02g41650.3                  GTPIGVSMDNTRLALAAIGKLMFAQFSELVNDFYNNGLPSNLSGGRNPSLDYGFKGAEIA
Pvaginatum-Pavag04G202800.1               GTPIGVSMDNTRLALAAIGKLMFAQFSELVNDFYNNGLPSNLSGGRNPSLDYGFKGAEIA
Pvaginatum-PavagK320900.1                 GTPIGVSMDNTRLALAAIGKLMFAQFSELVNDFYNNGLPSNLSGGRNPSLDYGFKGAEIA
Pvirgatum-Pavir.1NG356400.1               GTPIGVSMDNTRLALAAIGKLMFAQFSELVNDFYNNGLPSNLSGGRNPSLDYGFKGAEIA
Sviridis-Sevir.1G245100.1                 GTPIGVSMDNTRLALAAIGKLMFAQFSELVNDFYNNGLPSNLSGGRNPSLDYGFKGAEIA
Sbicolor-Sobic.004G220400.1               GTPIGVSMDNTRLALAAIGKLMFAQFSELVNDYYNNGLPSNLSGGRNPSLDYGFKGAEIA
Zmays-Zm00001d017275_T001                 GTPIGVSMDNTRLALAAIGKLMFAQFSELVNDYYNNGLPSNLSGGRNPSLDYGFKGAEIA
Zmays-Zm00001d051163_T002                 GTPIGVSMDNTRLALAAIGKLMFAQFSELVNDYYNNGLPSNLSGGRNPSLDYGFKGAEIA
Pvirgatum-Pavir.1KG386300.1               GTPIGVSMDNTRLALAAIGKLMFAQFSELVNDYYNNGLPSNLSGGRNPSLDYGFKGAEIA
Platifolius-Phala.02G210600.1             GTPIGVSMDNTRLAIAAIGKLMFAQFSELVNDFYNNGLPSNLSGGRNPSLDYGFKGAEIA
Streptochaeta-strangu_020767-RA           GTPIGVSMDNTRLAIAAIGKLMFAQFSELVNDFYNNGLPSNLSGGRNPSLDYGFKGAEIA
Bdistachyon-Bradi5g15830.1                GTPIGVSMDNTRLAIAAIGKLMFAQFSELVNDFYNNGLPSNLSGGRNPSLDYGFKGAEIA
Hvulgare-HORVU2Hr1G089440.4               GTPIGVSMDNTRLAIAAIGKLMFAQFSELVNDFYNNGLPSNLSGGRNPSLDYGFKGAEIA
Taestivum-Traes_2AL_0DDC60D6E.1           GTPIGVSMDNTRLAIAAIGKLMFAQFSELVNDFYNNGLPSNLSGGRNPSLDYGFKGAEIA
Tintermedium-Thint.06G0475900.1           GTPIGVSMDNTRLAIAAIGKLMFAQFSELVNDFYNNGLPSNLSGGRNPSLDYGFKGAEIA
Tintermedium-Thint.06G0476200.1           GTPIGVSMDNTRLAIAAIGKLMFAQFSELVNDFYNNGLPSNLSGGRNPSLDYGFKGAEIA
Taestivum-Traes_2AL_2763280FC.1           GTPIGVSMDNTRLAIAAIGKLMFAQFSELVNDFYNNGLPSNLSGGRNPSLDYGFKGAEIA
Taestivum-Traes_2DL_8C21AAAEF.1           GTPIGVSMDNTRLAIAAIGKLMFAQFSELVNDFYNNGLPSNLSGGRNPSLDYGFKGAEIA
Tintermedium-Thint.V0383500.1             GTPIGVSMDNTRLAIAVIGKLMFAQFSELVNDFYNNGLPSNLSGGRNPSLDYGFKGAEIA
Taestivum-Traes_2AL_9EC3226F7.1           GTPIGVSMDNTRLAIAAIGKLMFAQFSELVNDFYNNGLPSNLSGGRNPSLDYGFKGAEIA
Tintermedium-Thint.05G0273800.1           GTPIGVSMDNTRLAIAAIGKLMFAQFSELVNDFYNNGLPSNLSGGRNPSLDYGFKGAEIA
Tintermedium-Thint.V0383400.1             GTPIGVSMDNTRLAIAATGKLMFAQFSELVNDFYNNGLPSNLSGGRNPSLDYGFKGAEIA
Taestivum-Traes_2BL_13D5272D7.1           GTPIGVSMDNTRLAIAAIGKLMFAQFSELVNDFYNNGLPSNLSGGRNPSLDYGFKGAEIA
Tintermedium-Thint.06G0475700.1           GTPIGVSMDNTRLAIAAIGKLMFAQFSELVNDFYNNGLPSNLSGGRNPSLDYGFKGAEIA
Taestivum-Traes_2BL_C051606EA.2           GTPIGVSMDNTRLAIAAIGKLMFAQFSELVNDFYNNGLPSNLSGGRNPSLDYGFKGAEIA
Tintermedium-Thint.V0383600.1             GTPIGVSMDNTRLAIAAIGKLMFAQFSELVNDFYNNGLPSNLSGGRNPSLDYGFKGAEIA
Taestivum-Traes_2AL_9D78F85E2.1           GTPIGVSMDNTRLAIAAIGKLMFAQFSELVNDFYNNGLPSNLSGGRNPSLDYGFKGAEIA
Tintermedium-Thint.05G0274100.1           GTPIGVSMDNTRLAIAAIGKLMFAQFSELVNDFYNNGLPSNLSGGRNPSLDYGFKGAEIA
Tintermedium-Thint.06G0476300.1           GTPIGVSMDNTRLAIAAIGKLMFAQFSELVNDFYNNGLPSNLSGGRNPSLDYGFKGAEIA
Taestivum-Traes_2BL_5A50FDA1A.1           GTPIGVSMDNTRLAIAAIGKLMFAQFSELVNDFYNNGLPSNLSGGRNPSLDYGFKGAEIA
Tintermedium-Thint.05G0273900.1           GTPIGVSMDNTRLAIAAIGKLMFAQFSELVNDFYNNGLPSNLSGGRNPSLDYGFKGAEIA
Taestivum-Traes_2DL_30F8555D4.1           ------------------------------------------------------------
Taestivum-Traes_2DL_83168C1E0.1           ------------------------------------------------------------
Taestivum-Traes_2DL_B978B4E22.1           ------------------------------------------------------------
Taestivum-Traes_1AS_0B9295A68.1           GTPIGVSMDNTRLAIAAIGKLMFAQFSELVNDLYNNGLPSNLSGGRNPSLDYGFKGAEIA
Taestivum-Traes_1BS_7068F56E6.1           GTPIGVSMDNTRLAIAAIGKLMFAQFSELVNDLYNNGLPSNLSGGRNPSLDYGFKGAEIA
Taestivum-Traes_1BS_AEA817616.1           GTPIGVSMDNTRLAIAAIGKLMFAQFSELVNDLYNNGLPSNLSGGRNPSLDYGFKGAEIA
Taestivum-Traes_4AL_892C47ED5.1           GTPIGVSMDNTRLAIAAIGKLMFAQFSELVNDLYNNGLPSNLSGGRNPSLDYGFKGAEIA
Tintermedium-Thint.03G0055900.1           GTPIGVSMDNTRLAIAAIGKLMFAQFSELVNDLYNNGLPSNLSGGRNPSLDYGFKGAEIA
Tintermedium-Thint.01G0012900.1           GTPIGVSMDNTRLAIAAIGKLMFAQFSELVNDLYNNGLPSNLSGGRNPSLDYGFKGAEIA
Tintermedium-Thint.02G0158500.1           GTPIGVSMDNTRLAIAAIGKLMFAQFSELVNDLYNNGLPSNLSGGRNPSLDYGFKGAEIA
Tintermedium-Thint.03G0055700.1           GTPIGVSMDNTRLAIAAIGKLMFAQFSELVNDLYNNGLPSNLSGGRNPSLDYGFKGAEIA
Taestivum-Traes_1BS_C3AA04888.1           GTPIGVSMDNTRLAIAAIGKLMFAQFSELVNDLYNNGLPSNLSGGRNPSLDYGFKGAEIA
Taestivum-Traes_1DS_A8BD91E4A.1           GTPIGVSMDNTRLAIAAIGKLMFAQFSELVNDLYNNGLPSNLSGGRNPSLDYGFKGAEIA
Taestivum-Traes_1BS_C26CCE0E9.1           GTPIGVSMDNTRLAIAAIGKLMFAQFSELVNDFYNNGLPSNLSGGRNPSLDYGFKGAEIA
Taestivum-Traes_1DS_A21FCBA3D.1           GTPIGVSMDNTRLAIAAIGKLMFAQFSELVNDFYNNGLPSNLSGGRNPSLDYGFKGAEIA
Tintermedium-Thint.01G0012200.1           GTPIGVSMDNTRLAIAAIGKLMFAQFSELVNDFYNNGLPSNLSGGRNPSLDYGFKGAEIA
Tintermedium-Thint.02G0158600.1           GTPIGVSMDNTRLAIAAIGKLMFAQFSELVNDFYNNGLPSNLSGGRNPSLDYGFKGAEIA
Tintermedium-Thint.01G0012700.1           GTPIGVSMDNTRLAIAAIGKLMFAQFSELVNDFYNNGLPSNLSGGRNPSLDYGFKGAEIA
Taestivum-Traes_5BL_F50A47503.1           GTPIGVSMDNTRLAIAAIGKLMFAQFSELVNDFYNNGLPSNLSGGRNPSLDYGFKGAEIA
Tintermedium-Thint.02G0158700.1           GTPIGVSMDNTRLAIAAIGKLMFAQFSELVNDFYNNGLPSNLSGGRNPSLDYGFKGAEIA
Taestivum-Traes_5BL_AB528184A.1           GTPIGVSMDNTRLAIAAIGKLMFAQFSELVNDFYNNGLPSNLSGGRNPSLDYGFKGAEIA
Tintermedium-Thint.03G0055800.1           GTPIGVSMDNTRLAIAAIGKLMFAQFSELVNDFYNNGLPSNLSGGRNPSLDYGFKGAEIA
Taestivum-Traes_1AS_F9013A945.1           GTPIGVSMDNTRLAIAAIGKLMFAQFSELVNDFYNNGLPSNLSGGRNPSLDYGLKGAEIA
Taestivum-Traes_1DS_43DAC7A7F.1           GTPIGVSMDNTRLAIAAIGKLMFAQFSELVNDFYNNGLPSNLSGGRNPSLDYGLKGAEIA
Taestivum-Traes_1BS_BD86C90A7.1           GTPIGVSMDNTRLAIAAIGKLMFAQFSELVNDFYNNGLPSNLSGGRNPSLDYGLKGAEIA
Tintermedium-Thint.02G0158900.1           GTPIGVSMDNTRLAIAAIGKLMFAQFSELVNDFYNNGLPSNLSGGRNPSLDYGFKGAEIA
Tintermedium-Thint.05G0274000.1           GTPIGVSMDNTRLAIAAIGKLMFAQFSELVNDFYNNGLPSNLSGGRNPSFDYGFKGAEIA
Osativa-LOC_Os04g43800.1                  GTPIGVSMDNTRLAIAAIGKLMFAQFSELVNDFYNNGLPSNLSGGRNPSLDYGFKGAEIA
Pvaginatum-Pavag06G160300.1               GTPIGVSMDNTRLAVAAIGKLMFAQFSELVNDFYNNGLPSNLSGGRNPSLDYGFKGAEIA
Sbicolor-Sobic.006G148900.1               GTPIGVSMDNTRLAVAAIGKLMFAQFSELVNDFYNNGLPSNLSGGRNPSLDYGFKGAEIA
Pvirgatum-Pavir.7KG237800.1               GTPIGVSMDNTRLAVAAIGKLMFAQFSELVNDFYNNGLPSNLSGGRNPSLDYGFKGAEIA
Pvirgatum-Pavir.7NG355800.1               GTPIGVSMDNTRLAIAAIGKLMFAQFSELVNDFYNNGLPSNLSGGRNPSLDYGFKGAEIA
Sviridis-Sevir.7G178300.1                 GTPIGVSMDNTRLAIAAIGKLMFAQFSELVNDFYNNGLPSNLSGGRNPSLDYGFKGAEIA
Zmays-Zm00001d003015_T001                 GTPIGVSMDNTRLAVAAIGKLMFAQFSELVNDFYNNGLPSNLSGGRNPSLDYGFKGAEIA
Hvulgare-HORVU1Hr1G022060.1               GTPIGVSMDNTRLAIAAIGKLMFAQFSELVNDFYNNGLPSNLSGGRNPSLDYGFKGAEIA
Taestivum-Traes_1BS_723922D171.2          GTPIGVSMDNTRLAIAAIGKLMFAQFSELVNDYYNNGLPSNLSGGRNPSLDYGFKGAEIA
Taestivum-Traes_1DS_A171C7D59.1           GTPIGVSMDNTRLAIAAIGKLMFAQFSELVNDFYNNGLPSNLSGGRNPSLDYGFKGAEIA
Tintermedium-Thint.01G0111500.1           GTPIGVSMDNTRLAIAAIGKLMFAQFSELVNDFYNNGLPSNLSGGRNPSLDYGFKGAEIA
Tintermedium-Thint.03G0265300.1           GTPIGVSMDNTRLAIAAIGKLMFAQFSELVNDLYNNGLPSNLSGGRNPSLDYGFKGAEIA
Hvulgare-HORVU2Hr1G038140.1               GTPIGVSMDNTRLAIAAIGKLMFAQFSELVNDFYNNGLPSNLSGGRNPSLDYGFKGAEIA
Tintermedium-Thint.04G0171200.1           GTPIGVSMDNTRLAIAAIGKLMFAQFSELVNDFYNNGLPSNLSGGRNPSLDYGLKGAEIA
Taestivum-Traes_2AS_958327519.3           GTPIGVSMDNTRLAIAAIGKLMFAQFSELVNDFYNNGLPSNLSGGRNPSLDYGFKGAEIA
Taestivum-Traes_2DS_28CA50371.1           GTPIGVSMDNTRLAIAAIGKLMFAQFSELVNDFYNNGLPSNLSGGRNPSLDYGFKGAEIA
Tintermedium-Thint.05G0160800.1           GTPIGVSMDNTRLAIAAIGKLMFAQFSELVNDFYNNGLPSNLSGGRNPSLDYGFKGAEIA
Tintermedium-Thint.06G0299000.1           GTPIGVSMDNTRLAIAAIGKLMFAQFSELVNDFYNNGLPSNLSGGRNPSLDYGFKGAEIA
Taestivum-Traes_2DS_3791A8A36.1           GTPIGVSMDNTRLAVAAIGKLMFAQFSELVNDFYNNGLPSNLSGGRNPSLDYGFKGAEIA
Tintermedium-Thint.06G0342800.1           GTPIGVSMDNTRLAVAAIGKLMFAQFSELVNDFYNNGLPSNLSGGRNPSLDYGFKGAEIA
```

```
Tintermedium-Thint.05G0150900.1                   GTPIGVSMDNTRLAIAAIGKLMFAQFSELVNDFYNNGLPSNLSGGRNPSLDYGFKGAEIA
Tintermedium-Thint.04G0242300.1                   GTPIGVSMDNTRLAIAAIGKLMFAQFSELVNDFYNNGLPSNLSGGRNPSLDYGFKGAEIA
Tintermedium-Thint.06G0298700.1                   GTPIGVSMDNTRLAIAAIGKLMFAQFSELVNDFYNNGLPSNLSGGRNPSLDYGFKGAEIA
Taestivum-Traes_2BS_88CF42F2E.1                   GTPIGVSMDNTRLAIAAIGKLMFAQFSELVNDFYNNGLPSNLSGGRNPSLDYGFKGAEIA
Tintermedium-Thint.04G0241900.1                   GTPIGVSMDNTRLAIAAIGKLMFAQFSELVNDFYNNGLPSNLSGGRNPSLDYGFKGAEIA
Platifolius-Phala.06G165300.1                     GTPIGVSMDNTRLAIAAIGKLMFAQFSELVNDFYNNGLPSNLSGGRNPSLDYGFKGAEIA
Streptochaeta-strangu_019385-RA                   GTPIGVSMDNARLAIAAIGKLMFAQFSELVNDIYNNGLPSNLSGGRNPSLDYGFKGAEIA
Streptochaeta-strangu_028944-RA                   GTPIGVSMDNTRLAVAAISRLMFAQFSELVNDFYNNGLPANLSGGRDPSLDFGFKGADIA
Acomosus-Aco010091.1                              GTPIGVSMDNTRLAVAAIGKLMFAQFSELVNDLYNNGLPSNLSGGRNPSLDYGFKGAEIA
Zmarina-Zosma03g22300.1                           GTPIGVSMDNTRLAIAAIGKLVFAQFSELVNHFYNNGLPSNLSGGSNPSLDYGFKGGEIA
Ecdeiocolea-Emoptg0003741_1G000600.1              GTPIGVSMDNTRLAIAAIGKLMFAQFSELVNDFYNNGLPSNLSGGRNPSLDYGFKGAEIA
Jascendens-Joasc.05G060500.1                      GTPIGVSMDNTRLAIAAIGKLMFAQFSELVNDFYNNGLPSNLSGGRNPSLDYGFKGAEIA
Ecdeiocolea-Emoptg0003741_1G000630.1              GTPIGVSMDNARLAIANIGKLMFAQFSELVNEFYNNGLTSNLAGSRNPSLDYGFKGAEIA
Jascendens-Joasc.05G060400.1                      GTPIGVSMDNTRLAIANIGKLMFAQFSELVNDFYNNGLTSNLAGGRNPSLDYGFKGTEIA
Bdistachyon-Bradi3g49250.2                        GTPIGVSMDNTRLAIANIGKLMFAQFSELVNEFYNNGLTSNLAGSRNPSLDYGFKGTEIA
Hvulgare-HORVU6Hr1G058820.1                       GTPIGVSMDNTRLAIANIGKLMFAQFSELVNEFYNNGLTSNLAGSRNPSLDYGFKGTEIA
Taestivum-Traes_6DL_1AEA7B869.1                   GTPIGVSMDNTRLAIANIGKLMFAQFSELVNEFYNNGLTSNLAGSRNPSLDYGFKGTEIA
Tintermedium-Thint.18G0192300.1                   GTPIGVSMDNTRLAIANIGKLMFAQFSELVNEFYNNGLTSNLAGSRNPSLDYGFKGTEIA
Tintermedium-Thint.17G0230800.1                   GTPIGVSMDNTRLAIANIGKLMFAQFSELVNEFYNNGLTSNLAGSRNPSLDYGFKGTEIA
Tintermedium-Thint.17G0230900.1                   GTPIGVSMDNTRLAIANIGKLMFAQFSELVNEFYNNGLTSNLAGSRNPSLDYGFKGTEIA
Ecoracana-ELECO.r07.2AG0138000.1                  GTPIGVSMDNARLAIANIGKLMFAQFSELVNEFYNNGLTSNLAGSRNPSLDYGFKGTEIA
Ecoracana-ELECO.r07.2BG0193650.1                  GTPIGVSMDNARLAIANIGKLMFAQFSELVNEFYNNGLTSNLAGSRNPSLDYGFKGTEIA
Pvaginatum-Pavag04G202700.1                       GTPIGVSMDNARLAIANIGKLMFAQFSELVNEFYNNGLTSNLAGSRNPSLDYGFKGTEIA
Sviridis-Sevir.1G245000.1                         GTPIGVSMDNARLAIANIGKLMFAQFSELVNEFYNNGLTSNLAGSRNPSLDYGFKGTEIA
Sviridis-Sevir.6G187100.1                         GTPIGVSMDNARLAIANIGKLMFAQFSELVNEFYNNGLTSNLAGSRNPSLDYGFKGTEIA
Pvirgatum-Pavir.1NG356200.1                       GTPIGVSMDNARLAIANIGKLMFAQFSELVNEFYNNGLTSNLAGSRNPSLDYGFKGTEIA
Sbicolor-Sobic.004G220300.1                       GTPIGVSMDNARLAIANIGKLMFAQFSELVNEFYNNGLTSNLAGSRNPSLDYGFKGTEIA
Zmays-Zm00001d017274_T001                         GTPIGVSMDNARLAIANIGKLMFAQFSELVNEFYNNGLTSNLAGSRNPSLDYGFKGTEIA
Zmays-Zm00001d051161_T003                         GTPIGVSMDNARLAIANIGKLMFAQFSELVNEFYNNGLTSNLAGSRNPSLDYGFKGTEIA
Sviridis-Sevir.7G178200.1                         GTPIGVSMDNARLAIANIGKLMFAQFSELVNEFYNNGLTSNLAGSRNPSLDYGFKGTEIA
Osativa-LOC_Os02g41630.2                          GTPIGVSMDNARLAIANIGKLMFAQFSELVNEFYNNGLTSNLAGSRNPSLDYGFKGTEIA
Ecoracana-ELECO.r07.4AG0310430.1                  GTPIGVSMDNARLALANIGKLMFAQFSELVNEFYNNGLTSNLAGSRNPSLDYGFKGTEIA
Ecoracana-ELECO.r07.4BG0341620.1                  GTPIGVSMDNARLAIANIGKLMFAQFSELVNEFYNNGLTSNLAGSRNPSLDYGFKGTEIA
Pvaginatum-Pavag06G160200.1                       GTPIGVSMDNARLAIANIGKLMFAQFSELVNEFYNNGLTSNLAGSRNPSLDYGFKGTEIA
Pvirgatum-Pavir.7KG238255.1                       GTPIGVSMDNARLAIANIGKLMFAQFSELVNEFYNNGLTSNLAGSRNPSLDYGFKGTEIA
Sbicolor-Sobic.006G148800.1                       GTPIGVSMDNARLAIANIGKLMFAQFSELVNEFYNNGLTSNLAGSRNPSLDYGFKGTEIA
Zmays-Zm00001d003016_T001                         GTPIGVSMDNARLAVANIGKLMFAQFSELVNEFYNNGLTSNLAGSRNPSLDYGFKGTEIA
Osativa-LOC_Os04g43760.1                          GTPIGVSMDNTRLAIANIGKLMFAQFSELVNEFYNNGLTSNLAGSRNPSLDYGFKGTEIA
Platifolius-Phala.02G210400.1                     GTPIGVSMDNTRLAIANIGKLMFAQFSELVNEFYNNGLTSNLAGSRNPSLDYGFKGTEIA
Platifolius-Phala.06G165100.1                     GTPIGVSMDNTRLAIANIGKLMFAQFSELVNEFYNNGLTSNLAGSRNPSLDYGFKGTEIA
Streptochaeta-strangu_019386-RA                   GTPIGVSMDNARLAIASIGKLMFAQFSELVNEFYNNGLTSNLAGSRNPSLDYGFKGTEIA
Streptochaeta-strangu_020769-RA                   GTPIGVSMDNARLAIANIGKLMFAQFSELVNEFYNNGLTSNLAGSRNPSLDYGFKGTEIA
Sviridis-Sevir.2G448300.1                         GTPIGVSMDNARLAVASIGRLMFAQFTELVIDFYNNGLPSNLAGSRNLSLDFGLKGAEIA
Sviridis-Sevir.7G177900.1                         GTPIGVSMDNARLAVASIGRLMFAQFTELVIDFYNNGLPSNLAGSRNLSLDFGLKGAEIA
```

```
Aamericanus-Acora.01G232100.1                     MAAYCSELQFLGNPVTNHVQSAEQHNQDVNSLGLISSRKTAEAVEILKLMSATYLVGLCQ
Aamericanus-Acora.10G174800.1                     MAAYCSELQFLGNPVTNHVQSAEQHNQDVNSLGLISSRKTAEAVEILKLMSATYLVGLCQ
Acomosus-Aco013943.1                              MAAYCSELQFLANPVTNHVQSAEQHNQDVNSLGLISARKTAEAVEILKLMSSTFLIALCQ
Aofficinalis-evm.model.AsparagusV1_01.827         MASYCSELQFLANPVTNHVQSAEQHNQDVNSLGLISSRKTAEAVHILKLMSSTFLVGLCQ
Typha_latifolia-Tylat.14G083800.1                 MASYCSELQFLGNPVTNHVQSAEQHNQDVNSLGLISSRKTAEAVDILKLMSSTYLVALCQ
Aofficinalis-evm.model.AsparagusV1_04.71          MASYCSELQYLANPVTNHVESAEQHNQDVNSLGLISSRKTAEAVEILKLMTSTFLVALCQ
Aofficinalis-evm.model.AsparagusV1_04.93          MASYCSELQYLANPVTNHVESAEQHNQDVNSLGLISSRKTAEAVEILKLMTSTFLVALCQ
Aofficinalis-evm.model.AsparagusV1_10.1948        MASYCSELQYLANPVTNHVQSAEQHNQDVNSLGLISSRKTAEAVEILKLMTSTFLVALCQ
Typha_latifolia-Tylat.03G127700.1                 MASYCSELQFLANPVTNHVQSAEQHNQDVNSLGLISSRKTAEAVDILKLMSSTYLIALCQ
Spolyrhiza-Spipo11G0025500                        MASYCSELQFLGNPVTNHVQSAEQHNQDVNSLGLISARKTAEAVEILKLMTSTYLVALCQ
Spolyrhiza-Spipo1G0003500                         MAAYCSELQFLANPVTNHVQSAEQHNQDVNSLGLISARKTAEAVEILKLMTSTYLVALCQ
Phoenix_dactylifera-LOC103718264                  MAAYCSEIQFLGNPVTNHVQSAEQHNQDVNSLGLISARKTAEAVETLKLMSSTYLVALCQ
Macuminata-GSMUA_Achr1T09070_001                  MAAYCSELQFLANPVTNHVQSAEQHNQDVNSLGLISARKTAEAVDILKLMSTTYLVALCQ
Acomosus-Aco006987.1                              MAAYCSELQFLGNPVTNHVQSAEQHNQDVNSLGLISARKTAEAVEILKLMTSTYLVALCQ
Acomosus-Aco027752.1                              MAAYCSELQFLGNPVTNHVQSAEQHNQDVNSLGLISARKTAEAVEILKLMTSTYLVALCQ
Cscoparia-Csco27g03980.1                          MASYCSELQFLGNPVTNHVQSAEQHNQDVNSLGLISSRKTAEAVEILKLMSSTFMVALCQ
Cscoparia-Csco27g03990.1                          MASYCSELQFLGNPVTNHVQSAEQHNQDVNSLGLISSRKTAEAVEILKLMSSTFLVALCQ
Macuminata-GSMUA_Achr11T16380_001                 MAAYCSELQFLANPVTNHVQSAEQHNQDVNSLGFISSRKAAEAVDILKLMSATYLLALCQ
Macuminata-GSMUA_Achr11T22840_001                 MAAYCSELQFLANPVTNHVQSAEQHNQDVNSLGLISSRKTAEAVDILKLMSTTYLVALCQ
Macuminata-GSMUA_Achr2T00240_001                  MAAYCSELQFLANPVTNHVQSAEQHNQDVNSLGLISSRKTADAVDILKLMSATYLVALCQ
Macuminata-GSMUA_Achr5T18560_001                  MAAYCSELQFLANPVTNHVQSAEQHNQDVNSLGLISSRKTAEAVDILKLMSATYLVALCQ
Macuminata-GSMUA_Achr8T18960_001                  MAAYCSELQFLANPVTNHVQSAEQHNQDVNSLGLISSRKTAEAVDILKLMSATYLVALCQ
Macuminata-GSMUA_Achr5T03950_001                  MAAYCSELQFLANPVTNHVQSAEQHNQDVNSLGLISSRKTAEAVDILKLMSSTYLVALCQ
Phoenix_dactylifera-LOC103703950                  MAAYCSELQFLGNPVTNHVQSAEQHNQDVNSLGLISARKTQEAVEILKLMSSTYLVAICQ
Acoerulea-Aqcoe1G140100.1                         MASYCSELQFLANPVTNHVQSAEQHNQDVNSLGLISSRKTAEAVDILKLMSSTFLVALCQ
Acoerulea-Aqcoe2G144500.1                         MASYCSELQFLANPVTNHVQSAEQHNQDVNSLGLISSRKTAEAVDILKLMSSTFLVALCQ
Typha_latifolia-Tylat.05G053500.1                 MAAYCSELQFLANPVTNHVQSAEQHNQDVNSLGLISSRKTAEAVEILKLMSSTYLVALCQ
Zmarina-Zosma05g29470.1                           MASYCSELQFLANPVTNHVQSAEQHNQDVNSLGLISARKTAESIEILKLMTSTFLVGICQ
Acomosus-Aco007727.1                              MAAYCSELQFLANPVTNHVQSAEQHNQDVNSLGLISSRKTAEAVEILKLMSSTYLIALCQ
Bvulgarisssp-Bevul.7G054300.1                     MASYCSELQFLGNPVTNHVQSAEQHNQDVNSLGLISARKTAEAVDILKLMSSTFLVALCQ
Bdistachyon-Bradi3g47110.1                        MASYCSELQFLGNPVTNHVQSAEQHNQDVNSLGLISARKTAEAIEILKLMTSTFLVALCQ
Bdistachyon-Bradi3g47120.1                        MASYCSELQFLGNPVTNHVQSAEQHNQDVNSLGLISARKTAEAIEILKLMTSTFLVALCQ
Bdistachyon-Bradi3g48840.1                        MASYCSELQFLGNPVTNHVQSAEQHNQDVNSLGLISSRKTAEAVEILKLMSSTFLVALCQ
Bdistachyon-Bradi3g49280.1                        MASYCSELQFLGNPVTNHVQSAEQHNQDVNSLGLISSRKTAEAVEILKLMSSTFLVALCQ
Hvulgare-HORVU6Hr1G058840.1                       MASYCSELQFLGNPVTNHVQSAEQHNQDVNSLGLISSRKTAEAIEILKLMSSTFLVALCQ
Taestivum-Traes_6BL_C96F626D9.1                   MASYCSELQFLGNPVTNHVQSAEQHNQDVNSLGLISSRKTAEAIEILKLMSSTFLVALCQ
Tintermedium-Thint.18G0192700.1                   MASYCSELQFLGNPVTNHVQSAEQHNQDVNSLGLISSRKTAEAIEILKLMSSTFLVALCQ
Taestivum-Traes_6DL_04CD7E82C.1                   MASYCSELQFLGNPVTNHVQSAEQHNQDVNSLGLISFRKTAEAIEILKLMSSTFLVALCQ
Bdistachyon-Bradi3g49270.1                        MASYCSELQFLGNPVTNHVQSAEQHNQDVNSLGLISSRKTAEAIEILKLMSSTFLVALCQ
Ecoracana-ELECO.r07.2AG0138020.1                  MASYCSELQFLGNPVTNHVQSAEQHNQDVNSLGLISSRKTAEAIEILKLMSSTFLVALCQ
Ecoracana-ELECO.r07.2BG0193670.1                  MASYCSELQFLGNPVTNHVQSAEQHNQDVNSLGLISSRKTAEAIEILKLMSSTFLVALCQ
Osativa-LOC_Os02g41670.1                          MASYCSELQFLGNPVTNHVQSAEQHNQDVNSLGLISSRKTAEAVEILKLMSSTFLVALCQ
Osativa-LOC_Os02g41680.1                          MASYCSELQFLGNPVTNHVQSAEQHNQDVNSLGLISSRKTDEAIDILKLMSSTFLIALCQ
Pvaginatum-Pavag04G202900.1                       MASYCSELQFLGNPVTNHVQSAEQHNQDVNSLGLISSRKTAEAIEILKLMSSTFLIALCQ
Pvaginatum-PavagK321000.1                         MASYCSELQFLGNPVTNHVQSAEQHNQDVNSLGLISSRKTAEAIEILKLMSSTFLIALCQ
Pvaginatum-PavagK321300.1                         MASYCSELQFLGNPVTNHVQSAEQHNQDVNSLGLISSRKTAEAIEILKLMSSTFLIALCQ
Pvaginatum-PavagK321100.1                         MASYCSELQFLGNPVTNHVQSAEQHNQDVNSLGLISSRKTAEAIEILKLMSSTFLIALCQ
Pvaginatum-PavagK321200.1                         MASYCSELQFLGNPVTNHVQSAEQHNQDVNSLGLISSRKTAEAIEILKLMSSTFLIALCQ
Pvirgatum-Pavir.1KG386500.1                       MASYCSELQFLGNPVTNHVQSAEQHNQDVNSLGLISSRKTAEAIEILKLMSSTFLIALCQ
Pvirgatum-Pavir.1NG356700.1                       MASYCSELQFLGNPVTNHVQSAEQHNQDVNSLGLISSRKTAEAIEILKLMSSTFLIALCQ
Sviridis-Sevir.1G245166.1                         MASYCSELQFLGNPVTNHVQSAEQHNQDVNSLGLISSRKTAEAIEILKLMTSTFLIALCQ
Sviridis-Sevir.1G245232.1                         MASYCSELQFLGNPVTNHVQSAEQHNQDVNSLGLISSRKTAEAIEILKLMTSTFLIALCQ
Pvirgatum-Pavir.1NG356800.1                       MASYCSELQFLGNPVTNHVQSAEQHNQDVNSLGLISSRKTAEAIEILKLMSSTFLIALCQ
Sbicolor-Sobic.004G220500.1                       MASYCSELQFLGNPVTNHVQSAEQHNQDVNSLGLISSRKTAEAIEILKLMSSTFLIALCQ
Sbicolor-Sobic.004G220600.2                       MASYCSELQFLGNPVTNHVQSAEQHNQDVNSLGLISSRKTAEAIEILKLMSSTFLIALCQ
Sbicolor-Sobic.004G220700.1                       MASYCSELQFLGNPVTNHVQSAEQHNQDVNSLGLISSRKTAEAIEILKLMSSTFLIALCQ
Sviridis-Sevir.1G245300.2                         MASYCSELQFLGNPVTNHVQSAEQHNQDVNSLGLISSRKTAEAIEILKLMTSSFLIALCQ
Zmays-Zm00001d017276_T001                         MASYCSELQFLGNSVTNHVQSAEQHNQDVNSLGLISSRKTAEAIEILKIMSSTFLIALCQ
Zmays-Zm00001d017279_T001                         MASYCSELQFLGNPVTNHVQSAEQHNQDVNSLGLISSRKTAEAIEILKLMSSTFLIALCQ
Zmays-Zm00001d051166_T001                         MASYCSELQFLGNPVTNHVQSAEQHNQDVNSLGLISSRKTAEAIEILKLMSSTFLIALCQ
Ecoracana-ELECO.r07.2AG0138030.1                  MASYCSELQFLGNPVTNHVQSAEQHNQDVNSLGLISSRKTAEAIDILKLMSSTFLIALCQ
Ecoracana-ELECO.r07.2BG0193680.1                  MASYCSELQFLGNPVTNHVQSAEQHNQDVNSLGLISSRKTAEAIDILKLMSSTFLIALCQ
Osativa-LOC_Os05g35290.1                          MASYCSELQFLGNPVTNHVQSAEQHNQDVNSLGLISSRKTAEAIDILKLMSSTFLVALCQ
Platifolius-Phala.02G210700.1                     MASYCSELQFLGNPVTNHVQSAEQHNQDVNSLGLISSRKTAEAIDILKLMSSTFLIALCQ
Ecoracana-ELECO.r07.4AG0310420.1                  MASYCSELQFLGNPVTNHVQSAEQHNQDVNSLGLISSRKTAEAVDILRLMSSTFLIALCQ
Ecoracana-ELECO.r07.4BG0341610.1                  MASYCSELQFLGNPVTNHVQSAEQHNQDVNSLGLISSRKTAEAIDILKLMSSTFLIALCQ
Bdistachyon-Bradi3g49260.1                        MASYCSELQFLGNPVTNHVQSAEQHNQDVNSLGLISSRKTAEAIDILKIMSSTFLIALCQ
Hvulgare-HORVU0Hr1G016330.1                       MASYCSELQFLGNPVTNHVQSAEQHNQDVNSLGLISSRKTAEAIDILKIMSSTFLVALCQ
Tintermedium-Thint.17G0260200.1                   MASYCSELQFLGNPVTNHVQSAEQHNQDVNSLGLISSRKTAEAIDILKIMSSTFLVALCQ
Tintermedium-Thint.18G0192600.1                   MASYCSELQFLGNPVTNHVQSAEQHNQDVNSLGLISSRKTAEAIDILKIMSSTFLVALCQ
Tintermedium-Thint.17G0231000.1                   MASYCSELQFLGNPVTNHVQSAEQHNQDVNSLGLISSRKTAEAIDILKIMSSTFLVALCQ
Ecoracana-ELECO.r07.2AG0138010.1                  MASYCSELQFLANPVTNHVQSAEQHNQDVNSLGLISSRKTAEAIDILKLMSSTYLIALCQ
Ecoracana-ELECO.r07.2BG0193660.1                  MASYCSELQFLANPVTNHVQSAEQHNQDVNSLGLISSRKTAEAIDILKLMSSTYLIALCQ
Osativa-LOC_Os02g41650.3                          MASYCSELQFLGNPVTNHVQSAEQHNQDVNSLGLISSRKTAEAIDILKLMSSTFLIALCQ
Pvaginatum-Pavag04G202800.1                       MASYCSELQFLGNPVTNHVQSAEQHNQDVNSLGLISSRKTAEAIDILKLMSSTFLIALCQ
Pvaginatum-PavagK320900.1                         MASYCSELQFLGNPVTNHVQSAEQHNQDVNSLGLISSRKTAEAIDILKLMSSTFLIALCQ
Pvirgatum-Pavir.1NG356400.1                       MASYCSELQFLGNPVTNHVQSAEQHNQDVNSLGLISSRKTAEAIDILKLMTSTFLIALCQ
Sviridis-Sevir.1G245100.1                         MASYCSELQFLGNPVTNHVQSAEQHNQDVNSLGLISSRKTAEAIEILKLMTSTFLIALCQ
Sbicolor-Sobic.004G220400.1                       MASYCSELQFLGNPVTNHVQSAEQHNQDVNSLGLISSRKTAEAIDILKLMTSTFLIALCQ
Zmays-Zm00001d017275_T001                         MASYCSELQFLGNPVTNHVQSAEQHNQDVNSLGLISSRKTAEAVDILKLMTSTFLIALCQ
Zmays-Zm00001d051163_T002                         MASYCSELQFLGNPVTNHVQSAEQHNQDVNSLGLISSRKTAEAIDILKLMSSTFLIALCQ
Pvirgatum-Pavir.1KG386300.1                       MASYCSELQFLGNPVTNHVQSAEQHNQDVNSLGLISSRKTAEAIDILKLMTSTFLIALCQ
```

```
Platifolius-Phala.02G210600.1                 MASYCSELQFLANPVTNHVQSAEQHNQDVNSLGLISSRKTAEAIDILKLMSSTFLIALCQ
Streptochaeta-strangu_020767-RA               MASYCSELQFLGNPVTNHVQSAEQHNQDVNSLGLISSRKTAEAIDILKLMSSTFLIALCQ
Bdistachyon-Bradi5g15830.1                    MASYCSELQFLGNPVTNHVQSAEQHNQDVNSLGLISSRKTAEAIDILKLMSSTFLVALCQ
Hvulgare-HORVU2Hr1G089440.4                   MASYCSELQFLGNPVTNHVQSAEQHNQDVNSLGLISSRKTAEAIDILKLMSSTFLVALCQ
Taestivum-Traes_2AL_0DDC60D6E.1               MASYCSELQFLGNPVTNHVQSAEQHNQDVNSLGLISSRKTAEAIDILKLMSSTFLVALCQ
Tintermedium-Thint.06G0475900.1               MASYCSELQFLGNPVTNHVQSAEQHNQDVNSLGLISSRKTAEAIDILKLMSSTFLVALCQ
Tintermedium-Thint.06G0476200.1               MASYCSELQFLGNPVTNHVQSAEQHNQDVNSLGLISSRKTAEAIDILKLMSSTFLVALCQ
Taestivum-Traes_2AL_2763280FC.1               MASYCSELQFLGNPVTNHVQSAEQHNQDVNSLGLISSRKTAEAIDILKLMSSTFLVALCQ
Taestivum-Traes_2DL_8C21AAAEF.1               MASYCSELQFLGNPVTNHVQSAEQHNQDVNSLGLISSRKTAEAIDILKLMSSTFLVALCQ
Tintermedium-Thint.V0383500.1                 MASYCSELQFLGNPVTNHVQSAEQHNQDVNSLGLISSRKTAEAIDILKLMSSTLLVALCQ
Taestivum-Traes_2AL_9EC3226F7.1               MASYCSELQFLGNPVTNHVQSAEQHNQDVNSLGLISSRKTAEAIDILKLMSSTFLVALCQ
Tintermedium-Thint.05G0273800.1               MASYCSELQFLGNPVTNHVQSAEQHNQDVNSLGLISSRKTAEAIDILKLMSSTFLVALCQ
Tintermedium-Thint.V0383400.1                 MASYCSELQFLGNPVTNHVQSAEQHNQDVNSLGLISSRKTAEAIDILKLMSSTFLVALCQ
Taestivum-Traes_2BL_13D5272D7.1               MASYCSELQFLGNPVTNHVQSAEQHNQDVNSLGLISSRKTAEAIDILKLMSSTFLVALCQ
Tintermedium-Thint.06G0475700.1               MASYCSELQFLGNPVTNHVQSAEQHNQDVNSLGLISSRKTAEAIDILKLMSSTFLVALCQ
Taestivum-Traes_2BL_C051606EA.2               MASYCSELQFLGNPVTNHVQSAEQHNQDVNSLGLISSRKTAEAIDILKLMSSTFLVALCQ
Tintermedium-Thint.V0383600.1                 MASYCSELQFLGNPVTNHVQSAEQHNQDVNSLGLISSRKTAEAIDILKLMSSTFLVALCQ
Taestivum-Traes_2AL_9D78F85E2.1               MTSYCSELQFLGNPVTNHVQSAEQHNQDVNSLGLISSRKTAEAIDILKLMSSTFLVALCQ
Tintermedium-Thint.05G0274100.1               MASYCSELQFLGNPVTNHVQSAEQHNQDVNSLGLISSRKTAEAIDILKLMSSTFLVALCQ
Tintermedium-Thint.06G0476300.1               MASYCSELQFLGNPVTNHVQSAEQHNQDVNSLGLISSRKTAEAIDILKLMSSTFLVALCQ
Taestivum-Traes_2BL_5A50FDA1A.1               MASYCSELQFLGNPVTNHVQSAEQHNQDVNSLGLISSRKTAEAIDILKLMSSTFLVALCQ
Tintermedium-Thint.05G0273900.1               MASYCSELQFLGNPVTNHVQSAEQHNQDVNSLGLISSRKTAEAIDILKLMSSTFLVALCQ
Taestivum-Traes_2DL_30F8555D4.1               MASYCSELQFLGNPVTNHVQSAEQHNQDVNSLGLISSRKTAEAIDILKLMSSTFLVALCQ
Taestivum-Traes_2DL_83168C1E0.1               ----------------------------------------------MSSTFLVALCQ
Taestivum-Traes_2DL_B978B4E22.1               ----------------------------------------------MSSTFLVALCQ
Taestivum-Traes_1AS_0B9295A68.1               MASYCSELQFLGNPVTNHVQSAEQHNQDVNSLGLISSRKTAEAIDILKLMSSTFMVALCQ
Taestivum-Traes_1BS_7068F56E6.1               MASYCSELQFLGNPVTNHVQSAEQHNQDINSLGLISSRKTAEAIDILKLMSSTFMVALCQ
Taestivum-Traes_1BS_AEA817616.1               MASYCSELQFLGNPVTNHVQSAEQHNQDVNSLGLISSRKTAEAIDILKLMSSTFMVALCQ
Taestivum-Traes_4AL_892C47ED5.1               MASYCSELQFLGNPVTNHVQSAEQHNQDVNSLGLISSRKTAEAIDILKLMSSTFMVALCQ
Tintermedium-Thint.03G0055900.1               MASYCSELQFLGNPVTNHVQSAEQHNQDVNSLGLISSRKTAEAIDILKLMSSTFMVALCQ
Tintermedium-Thint.01G0012900.1               MASYCSELQFLGNPVTNHVQSAEQHNQDVNSLGLISSRKTAEAIDILKLMSSTFMVALCQ
Tintermedium-Thint.02G0158500.1               MASYCSELQFLGNPVTNHVQSAEQHNQDVNSLGLISSRKTAEAIDILKLMSSTFMVALCQ
Tintermedium-Thint.03G0055700.1               MASYCSELQFLGNPVTNHVQSAEQHNQDVNSLGLISSRKTAEAIDILKLMSSTFMVALCQ
Taestivum-Traes_1BS_C3AA04888.1               MASYCSELQFLGNPVTNHVQSAEQHNQDVNSLGLISSRKTAEAIDILKLMSSTFMVALCQ
Taestivum-Traes_1DS_A8BD91E4A.1               MASYCSELQFLGNPVTNHVQSAEQHNQDVNSLGLISSRKTAEAIDILKLMSSTFMVALCQ
Taestivum-Traes_1BS_C26CCE0E9.1               MASYCSELQFLGNPVTNHVQSAEQHNQDVNSLGLISSRKTAEAIDILKLMSSTFMVALCQ
Taestivum-Traes_1DS_A21FCBA3D.1               MASYCSELQFLGNPVTNHVQSAEQHNQDVNSLGLISSRKTAEAIDILKLMSSTFMVALCQ
Tintermedium-Thint.01G0012200.1               MASYCSELQFLGNPVTNHVQSAEQHNQDVNSLGLISSRKTAEAIDILKLMSSTFMVALCQ
Tintermedium-Thint.02G0158600.1               MASYCSELQFLGNPVTNHVQSAEQHNQDVNSLGLISSRKTAEAIDILKLMSSTFLVALCQ
Tintermedium-Thint.01G0012700.1               MASYCSELQFLGNPVANHVQSAEQHNQDVNSLGLISARKTAEAIDILKLMSSTFLVALCQ
Taestivum-Traes_5BL_F50A47503.1               MASYCSELQFLGNPVTNHVQSAEQHNQDVNSLGLISSRKTAEAIDILKLMSSTFLVALCQ
Tintermedium-Thint.02G0158700.1               MASYCSELQFLGNPVTNHVQSAEQHNQDVNSLGLISSRKTAEAIDILKLMSSTFLVALCQ
Taestivum-Traes_5BL_AB528184A.1               MASYCSELQFLGNPVTNHVQSAEQHNQDVNSLGLISSRKTAEAIDILKLMSSTFMVALCQ
Tintermedium-Thint.03G0055800.1               MASYCSELQFLGNPVTNHVQSAEQHNQDVNSLGLISSRKTAEAIDILKLMSSTFMVALCQ
Taestivum-Traes_1AS_F9013A945.1               MASYCSELQFLGNPVTNHVQSAEQHNQDVNSLGLMSARKTAESIDILKLMSSTFLVALCQ
Taestivum-Traes_1DS_43DAC7A7F.1               MASYCSELQFLGNPVTNHVQSAEQHNQDVNSLGLISARKTAESIDILKLMSSTFLVALCQ
Taestivum-Traes_1BS_BD86C90A7.1               MASYCSELQFLGNPVTNHVQSAEQHNQDVNSLGLISARKTAESIDILKLMSSTFLVALCQ
Tintermedium-Thint.02G0158900.1               MASYCSELQFLGNPVTNHVQSAEQHNQDVNSLGLISSRKTAEAIDILKLMSSTFLVALCQ
Tintermedium-Thint.05G0274000.1               MASYCSELQFLGNPVTNHVHSVEQHNQDINSLGLISSRKTAEAIDILKLMSSTFLVALCQ
Osativa-LOC_Os04g43800.1                      MASYCSELQFLANPVTNHVQSAEQHNQDVNSLGLISSRKTAEAIDVLKLMSSTFLIALCQ
Pvaginatum-Pavag06G160300.1                   MASYCSELQFLANPVTNHVQSAEQHNQDVNSLGLISSRKTAEAVDVLKLMSSTFLVALCQ
Sbicolor-Sobic.006G148900.1                   MASYCSELQFLANPVTNHVQSAEQHNQDVNSLGLISSRKTAEAVDVLKLMSSTFLIALCQ
Pvirgatum-Pavir.7KG237800.1                   MASYCSELQFLANPVTNHVQSAEQHNQDVNSLGLISSRKTAEAVDILKLMSSTFLVALCQ
Pvirgatum-Pavir.7NG355800.1                   MASYCSELQFLANPVTNHVQSAEQHNQDVNSLGLISSRKTAEAVDILKLMSSTFLIALCQ
Sviridis-Sevir.7G178300.1                     MASYCSELQFLGNPVTNHVQSAEQHNQDVNSLGLISSRKTAEAVDILKLMSSTFLIALCQ
Zmays-Zm00001d003015_T001                     MASYCSELQFLANPVTNHVQSAEQHNQDVNSLGLISSRKTAEAVDVLKLMSSTFLVALCQ
Hvulgare-HORVU1Hr1G022060.1                   MASYCSELQFLGNPVTNHVQSAEQHNQDVNSLGLISSRKTAEAIDILKLMSSTFLVALCQ
Taestivum-Traes_1BS_723922D171.2              MASYCSELQFLGNPVTNHVQSAEQHNQDVNSLGLISSRKTAEAIDILKLMSSTFLVALCQ
Taestivum-Traes_1DS_A171C7D59.1               MASYCSELQFLGNPVTNHVQSAEQHNQDVNSLGLISSRKTAEAIDILKLMSSTFLVALCQ
Tintermedium-Thint.01G0111500.1               MASYCSELQFLGNPVTNHVQSAEQHNQDVNSLGLISSRKTAEAIDILKLMSSTFLVALCQ
Tintermedium-Thint.03G0265300.1               MASYCSELQFLGNPVTNHVQSAEQHNQDVNSLGLISSRKTAEAIDILKLMSSTFLVALCQ
Hvulgare-HORVU2Hr1G038140.1                   MASYCSELQFLGNPVTNHVQSAEQHNQDVNSLGLISARKTAEAIDILKLMSSTFLVALCQ
Tintermedium-Thint.04G0171200.1               MASYCSELQFLGNPVTNHVQSAEQHNQDVNSLGLISARKTAEAIDILKLMSSTFLVALCQ
Taestivum-Traes_2AS_958327519.3               MASYCSELQFLGNPVTNHVQSAEQHNQDVNSLGLISARKTAEAIDILKLMSSTFLVALCQ
Taestivum-Traes_2DS_28CA50371.1               MASYCSELQFLGNPVTNHVQSAEQHNQDVNSLGLISARKTAEAIDILKLMSSTFLVALCQ
Tintermedium-Thint.05G0160800.1               MASYCSELQFLGNPVTNHVQSAEQHNQDVNSLGLISARKTAEAIDILKLMSSTFLVALCQ
Tintermedium-Thint.06G0299000.1               MASYCSELQFLGNPVTNHVQSAEQHNQDVNSLGLISARKTAEAIDILKLMSSTFLVALCQ
Taestivum-Traes_2DS_3791A8A36.1               MASYCSELQFLGNPVTNHVQSAEQHNQDVNSLGLISARKTAEAIDILKLMSSTFLVALCQ
Tintermedium-Thint.06G0342800.1               MASYCSELQFLGNPVTNHVQSAEQHNQDVNSLGLISARKTAEAIDILKLMSSTFLVALCQ
Tintermedium-Thint.05G0150900.1               MASYCSELQFLGNPVTNHVQSAEQHNQDVNSLGLISSRKTAEAIDILKLMSSTFLVALCQ
Tintermedium-Thint.04G0242300.1               MASYCSELQFLGNPVTNHVQSAEQHNQDVNSLGLISSRKTAEAIDILKLMSSTFLVALCQ
Tintermedium-Thint.06G0298700.1               MASYCSELQFLGNPVTNHVQSAEQHNQDVNSLGLISSRKTAEAIDILKLMSSTFLVALCQ
Taestivum-Traes_2BS_88CF42F2E.1               MASYCSELQFLGNPVTNHVQSAEQHNQDVNSLGLISARKTAEAIDILKLMSSTFLVALCQ
Tintermedium-Thint.04G0241900.1               MASYCSELQFLGNPVTNHVQSAEQHNQDVNSLGLISARKTAEAIDILKLMSSTFLVALCQ
Platifolius-Phala.06G165300.1                 MASYCSELQFLANPVTNHVQSAEQHNQDVNSLGLISSRKTAEAIDILKLMSSTFLIALCQ
Streptochaeta-strangu_019385-RA               MASYCSELQFLANPVTNHVQSAEQHNQDVNSLGLISSRKTAEAIDILKLMSSTYLIALCQ
Streptochaeta-strangu_028944-RA               IASYCSELQFLANPVTSHVQSAERHNQDINSLGFISARMTAEAIDLLKLMSSTFLVALCQ
Acomosus-Aco010091.1                          MAAYCSELQFLANPVTTHVQSAEQHNQDVNSLGLISARKTAEAVDVLKLMSSTYVVALCQ
Zmarina-Zosma03g22300.1                       MASYCSELQFLGNPVTNHVQSAEQHNQDVNSLGLISARKTAESVEILKMMSATFLVALVQ
Ecdeiocolea-Emoptg0003741_1G000600.1          MASYCSELQFLANPVTNHVQSAEQHNQDVNSLGLISSRKTAEAVDILKIMSSTFLVALCQ
Jascendens-Joasc.05G060500.1                  MASYCSELQFLANPVTNHVQSAEQHNQDVNSLGLISSRKTAEAVDILKLMSSTFLIALCQ
Ecdeiocolea-Emoptg0003741_1G000630.1          MASYCSELQFLANPVTNHVQSAEQHNQDVNSLGLVSARKTAEAIDILKLMSSTYLMALCQ
Jascendens-Joasc.05G060400.1                  MASYCSELQFLANPVTNHVQSAEQHNQDVNSLGLVSARKTAEAVDILKLMSSTFLIALCQ
Bdistachyon-Bradi3g49250.2                    MASYCSELQYLANPVTNHVQSAEQHNQDVNSLGLVSARKTAEAVDILKLMSSTYMVALCQ
Hvulgare-HORVU6Hr1G058820.1                   MASYCSELQYLANPVTNHVQSAEQHNQDVNSLGLVSARKTAEAVDILKLMSSTYMVALCQ
Taestivum-Traes_6DL_1AEA7B869.1               MASYCSELQYLANPVTNHVQSAEQHNQDVNSLGLVSARKTAEAVDILKLMSSTYMVALCQ
Tintermedium-Thint.18G0192300.1               MASYCSELQYLANPVTNHVQSAEQHNQDVNSLGLVSARKTAEAVDILKLMSSTYMVALCQ
Tintermedium-Thint.17G0230800.1               MASYCSELQYLANPVTNHVQSAEQHNQDVNSLGLVSARKTAEAVDILKLMSSTYMVALCQ
Tintermedium-Thint.17G0230900.1               MASYCSELQYLANPVTNHVQSAEQHNQDVNSLGLVSARKTAEAVDILKLMSSTYMVALCQ
Ecoracana-ELECO.r07.2AG0138000.1              MASYCSELQYLANPITNHVQSAEQHNQDVNSLGLVSARKTAEAVDILKLMSSTYIVALCQ
Ecoracana-ELECO.r07.2BG0193650.1              MASYCSELQYLANPITNHVQSAEQHNQDVNSLGLVSARKTAEAVDILKLMSSTYIVALCQ
Pvaginatum-Pavag04G202700.1                   MASYCSELQYLGNPITNHVQSAEQHNQDVNSLGLVSARKTAEAIDILKLMSSTYIVALCQ
Sviridis-Sevir.1G245000.1                     MASYCSELQYLGNPITNHVQSAEQHNQDVNSLGLVSARKTAEAIDILKLMSSTYMVALCQ
Sviridis-Sevir.6G187100.1                     MASYCSELQYLGNPITNHVQSAEQHNQDVNSLGLVSARKTAEAIDILKLMSSTYIVALCQ
Pvirgatum-Pavir.1NG356200.1                   MASYCSELQYLGNPITNHVQSAEQHNQDVNSLGLVSARKTAEAVDILKLMSSTYIVALCQ
Sbicolor-Sobic.004G220300.1                   MASYCSELQYLGNPITNHVQSAEQHNQDVNSLGLVSARKTAEAIDILKLMSSTYIVALCQ
Zmays-Zm00001d017274_T001                     MASYCSELQYLGNPITNHVQSAEQHNQDVNSLGLVSARKTAEAIDILKLMSSTYIVALCQ
Zmays-Zm00001d051161_T003                     MASYCSELQYLGNPITNHVQSAEQHNQDVNSLGLVSARKTAEAIDILKLMSSTYIVALCQ
Sviridis-Sevir.7G178200.1                     MASYCSELQYLGNPITNHVQSAEQHNQDVNSLGLVSARKTAEAIDILKLMSSTYMVALCQ
Osativa-LOC_Os02g41630.2                      MASYCSELQYLANPITNHVQSAEQHNQDVNSLGLVSARKTLEAVDILKLMTSTYIVALCQ
Ecoracana-ELECO.r07.4AG0310430.1              MASYCSELQFLANPVTNHVQSAEQHNQDVNSLGLVSARKTAEAVDILKLMSATYMVALCQ
Ecoracana-ELECO.r07.4BG0341620.1              MASYCSELQFLANPVTNHVQSAEQHNQDVNSLGLVSARKTAEAVDILKLMSATYMVALCQ
Pvaginatum-Pavag06G160200.1                   MASYCSELQYLANPITNHVQSAEQHNQDVNSLGLVSARKTAEAVDILKLMTSTYMVALCQ
Pvirgatum-Pavir.7KG238255.1                   MASYCSELQYLANPITNHVQSAEQHNQDVNSLGLVSARKTAEAIDILKLMSSTYMVALCQ
Sbicolor-Sobic.006G148800.1                   MASYCSELQYLANPITNHVQSAEQHNQDVNSLGLVSARKTAEAVDILKLMSSTYMVALCQ
Zmays-Zm00001d003016_T001                     MASYCSELQYLANPITNHVQSAEQHNQDVNSLGLVSARKTAEAVDILKLMSSTYMVALCQ
Osativa-LOC_Os04g43760.1                      MASYCSELQFLANPVTNHVQSAEQHNQDVNSLGLVSARKTAEAVDILKLMSSTYLVALCQ
Platifolius-Phala.02G210400.1                 MASYCSELQYLANPITNHVQSAEQHNQDVNSLGLVSARKTAEAVDILKLMSSTYLVALCQ
Platifolius-Phala.06G165100.1                 MASYCSELQFLANPVTNHVQSAEQHNQDVNSLGLVSARKTAEAVDILKLMSSTYLVALCQ
Streptochaeta-strangu_019386-RA               MASYCSELQYLGNPITNHVQSAEQHNQDVNSLGLVSARKTAEAIDILKLMSSTYLVALCQ
Streptochaeta-strangu_020769-RA               MASYCSELQFLANPVTNHVQSAEQHNQDVNSLGLVSARKTAEAVDILKLMSSTYLVALCQ
Sviridis-Sevir.2G448300.1                     MASYCSELQYLANPVTNHVQSAEQHTQDVNSLGLISARKTAEAVEILKLMSSTFMIALCQ
Sviridis-Sevir.7G177900.1                     MASYCSELQYLANPVTNHVQSAEQHTQDVNSLGLISARKTAEAVEILKLMSSTFMIALCQ
```

```
Aamericanus-Acora.01G232100.1                 AIDLRHLEEILRNTVRNTVMQVAKRVLTMGGNGELHPSRFCEKDLIKVVDRESVYAYVDE
Aamericanus-Acora.10G174800.1                 AIDLRHLEEIMRNTVRNTVSQVAKRTLTMGVNGELHPSRFCEKDLIKVIDREYVFAYVDD
Acomosus-Aco013943.1                          AIDLRHLEENLKNAIKNAVSQIAKRVLTMGPNGELHPSRFCEKDLITVIDREHVFSYADD
Aofficinalis-evm.model.AsparagusV1_01.827     AIDLRHLEENLRSTVKNTVSQVAKRTLTMGANGELHPSRFCEKDLLKVIDREYVFAYIDE
Typha_latifolia-Tylat.14G083800.1             AIDLRHLEENLKSTVKNTVSQVAKRVLTMGANGELHPSRFCEKDLIKVIDREYVFSYIDD
Aofficinalis-evm.model.AsparagusV1_04.71      AIDLRHLEENLKQAVKNTVSQVSKRVLTTGVNGELHPSRFCEKELIKVIDREYVFAYIDD
Aofficinalis-evm.model.AsparagusV1_04.93      AIDLRHLEENLKQAVKNTVSQVSKRVLTTGVNGELHPSRFCEKELIKVIDREYVFAYIDD
Aofficinalis-evm.model.AsparagusV1_10.1948    AIDLRHLEENLKQAVKNTVSQVSKRVLTTGVNGELHPSRFCELELIKMIDREYVFAYIDE
Typha_latifolia-Tylat.03G127700.1             AIDLRHLEENLRSSVKNTVTQVAKKVLTMGVNGELHPSRFCEKDLIKIIDHEYVFSYVDD
Spolyrhiza-Spipo11G0025500                    AVDLRHLEENLKAAVKNTVSQVAKRVLTMGVNGELHPSRFCEKDLLKVVDREYVFAYADD
Spolyrhiza-Spipo1G0003500                     AVDLRHLEENLKNAVKNTVSQVAKRVLTIGVNGELHPSRFCEKDLIRAVDREHVFTYADD
Phoenix_dactylifera-LOC103718264              AIDLRHLEENLKNTVKNTVTQAAKRVLTMGANGELHPSRFCEKDLIKVIDGEYVFSYIDD
Macuminata-GSMUA_Achr1T09070_001              AVDLRHLEENLKSAVKSTVSQVAKRVLTMGANGELHPARFCEKELIKVVDREHVFTYVDD
```

```
Acomosus-Aco006987.1                       AIDLRHLEENLKSAVKNAVGQAAKRVLTVGAGGELHPARFCEKDLIKVVDRDHVFAYADD
Acomosus-Aco027752.1                       AIDLRHLEENLKSAVKNAVGQAAKRVLTVGAGGELHPARFCEKDLIKVVDRDHVFAYADD
Cscoparia-Csco27g03980.1                   AIDLRHLEENLKATVKNTVTQVAKKVLSLGVNGELHPSRFCEKDLIKVIDREYVFAYADD
Cscoparia-Csco27g03990.1                   AIDLRHLEENLKATVKNTVTQVAKKVLSLGVNGELHPSRFCEKDLIKVIDREYVFAYADD
Macuminata-GSMUA_Achr11T16380_001          AIDLRHLEENLKNAVKNTVSQVAKRVLTTGA--------FCEKNLITVADREHIFGYIDD
Macuminata-GSMUA_Achr11T22840_001          AIDLRHLEENLKQAVKNTVSQVAKRVLTTAANGELHPSRFCEKDLITVIDREHVFSYIDD
Macuminata-GSMUA_Achr2T00240_001           AIDLRHLEENLKNAVKNTVSQVAKRVLTAGVNGELQPSRFCEKDLITIVDREHVFSYIDD
Macuminata-GSMUA_Achr5T18560_001           AIDLRHLEENLKHAVKNTVGQVAKRVLTTGANGELHPSRFCEKDLIMVVDRENVFSYIDD
Macuminata-GSMUA_Achr8T18960_001           AIDLRHLEENLKKAVTNTVSQVAKR-----------------------------------
Macuminata-GSMUA_Achr5T03950_001           AIDLRHLEENLKNAVKNTVIQVAKKTLTTGVNGELHPSRFCEKDLITIIDREHVFSYVDD
Phoenix_dactylifera-LOC103703950           AIDLRHLEENLKNTVKNTASQVAKRVLTTGVNGELHPSRFCEKDLIKVIDREYVFSYADD
Acoerulea-Aqcoe1G140100.1                  SIDLRHLEENLKSTVKNTVSQVAKRVLTMGVNGELHPSRFCEKDLLKVVDREYVFSYIDD
Acoerulea-Aqcoe2G144500.1                  AIDLRHLEENLKLTVKNTVSQVAKKVLTMGVNGELHPSRFCEKEMLKVVDREYLFAYIDD
Typha_latifolia-Tylat.05G053500.1          AIDLRHLEENLKAAVRNTVSQVAKKVLTMGVNGELHPSRFCEKELIRAVEVEDVFAYADD
Zmarina-Zosma05g29470.1                    AIDLRHMEENLKASVKNTVSQVAKRVLTMTANGELHPSRFCEKDLLKVVDREYVFSYIDD
Acomosus-Aco007727.1                       AIDLRHLEENLMHAVKNAVSQVAKRVLTMDTAGELHPARFCVKDLIKVVDCEAAFSYADD
Bvulgarisssp-Bevul.7G054300.1              AIDLRHLEENVKHAVKNTVSQVCRRVLITGSNGELHPGRFCEKELIRVVEREYVFAYADD
Bdistachyon-Bradi3g47110.1                 AIDLRHIEENVKSAVTSCVRAVAKKTLSTNSAGGLHVARFSEKDLIQEIDREAVFAYADD
Bdistachyon-Bradi3g47120.1                 AIDLRHIEENVKSAVTSCVRAVAKKTLSTNSAGGLHVARFSEKDLIQEIDREAVFAYADD
Bdistachyon-Bradi3g48840.1                 AIDLRHIEENVKIAVKSCVMTVAKKTLSTNSTGGLHVARFCEKDLLQEIEREAVFAYADD
Bdistachyon-Bradi3g49280.1                 AIDLRHIEENVKSAVKSCVMTVAKRTLSTNSTGGLHVARFCEKDLLQEIEREAVFAYADD
Hvulgare-HORVU6Hr1G058840.1                AIDLRHIEENVKSAVKSCVMTVAKKTLSTNSTGGLHVARFCEKDLLQEIEREAVFAYADD
Taestivum-Traes_6BL_C96F626D9.1            AIDLRHIEVNVKSAVKSCVIMVAKKTLSTNSTGGLHVARFCEKDLLQEIEREAVFAYADD
Tintermedium-Thint.18G0192700.1            AIDLRHIEENVKSAVKSCVMMVAKKTLSTNSTGGLHVARFCEKDLLQEIEREAVFAYADD
Taestivum-Traes_6DL_04CD7E82C.1            AIDLRHIEENVKSAVKSCVLTVAKKTLSTNSTSGLHVARFCEKDLLQEIEREAVFAYADD
Bdistachyon-Bradi3g49270.1                 AIDLRHIEENVKSAVKSCVMTVAKKTLSTNSTGDLHVSRFCEKDMLQEIDREAVFAYADD
Ecoracana-ELECO.r07.2AG0138020.1           AVDLRHIEENVKSAVKSCVMTVAKRTLCTNSTGGLHVARFCEKDMLQEIEREAVFAYSDD
Ecoracana-ELECO.r07.2BG0193670.1           AVDLRHIEENVKSAVKSCVMTVAKRTLCTNSTGGLHVARFCEKDMLQEIEREAVFAYADD
Osativa-LOC_Os02g41670.1                   AIDLRQIEENVKSAVKSCVMTVARKTLSTSATGGLHAARFCEKDLLQEIDREAVFAYADD
Osativa-LOC_Os02g41680.1                   AVDLRHIEENVKSAVKSCVMTVAKKTLSTNSTGDLHVARFCEKDLLKEIDREAVFAYADD
Pvaginatum-Pavag04G202900.1                AVDLRHIEENVKSAVKSCVMTVAKKTLSTNSTGGLHDARFCEKDLLQEIEREAVFAYADD
Pvaginatum-PavagK321000.1                  AVDLRHIEENVKSAVKSCVMTVAKKTLSTNSTGGLHDARFCEKDLLQEIEREAVFAYADD
Pvaginatum-PavagK321300.1                  AVDLRHIEENVKSAVKSCVMTVAKKTLSTNSTGGLHDARFCEKDLLQEIEREAVFAYADD
Pvaginatum-PavagK321100.1                  AVDLRHIEENVKSAVKSCVMTVAKKTLSTNSTGGLHDARFCEKDLLQEIEREAVFAYADD
Pvaginatum-PavagK321200.1                  AVDLRHIEENVKSAVKSCVMTVAKKTLSTNSTGGLHDARFCEKDLLQEIEREAVFAYADD
Pvirgatum-Pavir.1KG386500.1                AVDLRHIEENVKSAVKSCVMTVAKRTLSTNSSGGLHVARFCEKDLLQEIEREAVFAYADD
Pvirgatum-Pavir.1NG356700.1                AVDLRHIEENVKSAVKSCVMTVAKKTLSTNATGGLHVARFCEKDLLQEIEREAVFAYADD
Sviridis-Sevir.1G245166.1                  AVDLRHIEENVKSAVKSCVMTVAKKTLSTNSTGGLHVARFCEKDLLQEIEREAVFAYADD
Sviridis-Sevir.1G245232.1                  AVDLRHIEENVKSAVKSCVMTVAKKTLSTNSTGGLHVARFCEKDLLQEIEREAVFAYADD
Pvirgatum-Pavir.1NG356800.1                AVDLRHIEENVKSAVKSCVMTVAKKTLSTNSTGGLHVARFCEKDLLQEIEREAVFAYADD
Sbicolor-Sobic.004G220500.1                AVDLRHIEENVKSAVKSCVMTVAKKTLSTNSTGGLHVARFCEKDLLQEIEREAVFAYADD
Sbicolor-Sobic.004G220600.2                AVDLRHIEENVKSAVKSCVMTVAKKTLSTNSTGGLHVARFCEKDLLQEIEREAVFAYADD
Sbicolor-Sobic.004G220700.1                AVDLRHIEENVKSAVKSCVMTVAKKTLSTNSTGGLHVARFCEKDLLQEIEREAVFAYADD
Sviridis-Sevir.1G245300.2                  AVDLRHIEENVKSAVKGCVMTVAKKTLSTNSAGGLHVARFCEKDLLQEIEREAVFAYADD
Zmays-Zm00001d017276_T001                  AVDLRHIEENVKSAVKSCVMTVAKKTLSTDSTGGLHVARFCEKDLLQEIEREAVFAYADD
Zmays-Zm00001d017279_T001                  AVDLRHIEENVKSAVKSCVMTVAKKTLSTNSTGGLHVARFCEKDLLQEIEREAVFAYADD
Zmays-Zm00001d051166_T001                  AVDLRHIEENVKSAVKSCVMTVAKKTLSTNSTGGLHVARFCEKDLLQEIEREAVFAYADD
Ecoracana-ELECO.r07.2AG0138030.1           AVDLRHIEENVRSAVKTCVMTVAKKTLSTNSTGDLHAARFCEKDMLMEIEREAVFAYADD
Ecoracana-ELECO.r07.2BG0193680.1           AVDLRHIEENVKAAVKTCVMTVAKKTLSTNSTGDLHAARFCEKDMLMEIEREAVFAYADD
Osativa-LOC_Os05g35290.1                   AIDLRHIEENVKGAVKTCVMTVAKKTLSTNSTGGLHVARFCEKDLLSEIDREAVFAYADD
Platifolius-Phala.02G210700.1              AIDLRHLEENVKSAVKSCVMMVAKKTLSRNAAGDLHIARFCDKDLLTTIDREAVFAYADD
Ecoracana-ELECO.r07.4AG0310420.1           AVDLRHVEENVKAAVKSCVTAAAKRTLSTNSTGDLHAARFCEKDMLLEIEREAVFAYADD
Ecoracana-ELECO.r07.4BG0341610.1           AVDLRHAEENIKAAVKTCVTAAAKRTLSTNSTGDLHAARFCEKDMLLEIEREAVFAYADD
Bdistachyon-Bradi3g49260.1                 AIDLRHLEENMKTAVRNCVMQVAKKTLSMNAMGGLHIARFCEKDLLTAIDREAVFAYADD
Hvulgare-HORVU0Hr1G016330.1                AIDLRHIEENMKTAVRNCVMQVAKKTLSMNNMGGLHIARFCEKDLLTAIDREAVFAYADD
Tintermedium-Thint.17G0260200.1            AIDLRHIEENMKTAVRNCVMQVAKKTLSMNNMGGLHIARFCEKDLLTAIDREAVFAYADD
Tintermedium-Thint.18G0192600.1            AIDLRHIEENMKTAVRNCVMQVAKKTLSMNNMGGLHIARFCEKDLLTAIDREAVFAYADD
Tintermedium-Thint.17G0231000.1            AIDLRHIEENMKTAVRNCVMQVAKKTLSMNNMGGLHIARFCEKDLLTAIDREAVFAYADD
Ecoracana-ELECO.r07.2AG0138010.1           AADLRHLEENMKAAVKSCVMQIAKKSLSLNSMGGLHVARFCEKDLVTAIDREAVFAYADD
Ecoracana-ELECO.r07.2BG0193660.1           AADLRHLEENMKAAVKSCVMQIAKKSLSLNSMGGLHVARFCEKDLVTAIDREAVFAYADD
Osativa-LOC_Os02g41650.3                   AIDLRHLEENMKTAVKNCVMQVAKKSLSMNHMGGLHIARFCEKDLLTAIDREAVFAYADD
Pvaginatum-Pavag04G202800.1                AVDLRHLEENMKAAVKNCVTQVAKKSLSLNARGGLHIARFCEKDLQTAIDREAVFAYADD
Pvaginatum-PavagK320900.1                  AVDLRHLEENMKAAVKNCVTQVAKKSLSLNARGGLHIARFCEKDLQTAIDREAVFAYADD
Pvirgatum-Pavir.1NG356400.1                AIDLRHLEENMKAAVKNCVTQVAKKTLSMNAMGGLHIARFCEKDLQTAIDREAVFAYADD
Sviridis-Sevir.1G245100.1                  AIDLRHLEENMKAAVKNCVTQVAKKTLSMNAMGGLHIARFCEKDLQTAIDREAVFAYADD
Sbicolor-Sobic.004G220400.1                AIDLRHIEENVKAAVKNCVTQVAKKSLSLNARGGLHNARFCEKDLQTAIDREAVFAYADD
Zmays-Zm00001d017275_T001                  AIDLRHLEENVKAAVKNCVTQVAKKSLSLNARGGLHNARFCEKDLQTAIDREAVFAYADD
Zmays-Zm00001d051163_T002                  AIDLRHLEENVKAAVKNCVTQVAKKSLSLNARGGLHNARFCEKDLQTAIDREAVFAYADD
Pvirgatum-Pavir.1KG386300.1                AIDLRHLEENMKAAVKNCVTQVAKKTLSMNAMGGLHIARFCEKDLQTAIDREAVFAYADD
Platifolius-Phala.02G210600.1              AIDLRHLEENVKSAVKNCVMQVARKALSMSAMGGLHIARFCEKDLLTAIDREDVFAYADD
Streptochaeta-strangu_020767-RA            AIDLRHIEENVKSAVKNCVMQVSKRVLSLNATGNLHIARFCEKDLLTAIDREAVFAYADD
Bdistachyon-Bradi5g15830.1                 AIDLRHLEENVRSAVKNCVTTVARKTLSTNVNGHLHNARFCEKDLLLTIDREAVFAYADD
Hvulgare-HORVU2Hr1G089440.4                AIDLRHLEENVKNAVKSCVKTVARKTLSTNSNGHLHSARFCEKDLLLTIDREAVFAYADD
Taestivum-Traes_2AL_0DDC60D6E.1            AIDLRHLEENVKNAIKRCVKTVARKTLSTDTNGHLHNARFCEKDLLLTIDREAVFAYADD
Tintermedium-Thint.06G0475900.1            AIDLRHLEENVKNAVKSCVKTVARKTLSTDTNGHLHNARFCEKDLLLTIDREAVFAYADD
Tintermedium-Thint.06G0476200.1            AIDLRHLEENVKNAIKSCVKTVARKTLSTDTNGHLHNARFCEKDLLLTIDREAVFAYADD
Taestivum-Traes_2AL_2763280FC.1            AIDLRHLEENVKNAVKSCVKTVARKTLSTDTNGHLHNARFCEKDLLLTIDREAVFAYADD
Taestivum-Traes_2DL_8C21AAAEF.1            AIDLRHLEENVKNAVKSCVKTVARKTLSTDTNGHLHNARFCEKDLLLTIDREAVFAYADD
Tintermedium-Thint.V0383500.1              AIDLRHLEENVKNAVKSCVKTVARKTLSTDNNGHLHNARFCEKDLLLTIDREAVFAYADD
Taestivum-Traes_2AL_9EC3226F7.1            AIDLRHLEENVKNAVKSCVKTVARKTLSTDNNGHLHNARFCEKDLLLTIDREAVFAYADD
Tintermedium-Thint.05G0273800.1            AIDLRHLEENVKNAVKSCVKTVARKTLSTDNNGHLHNARFCEKDLLLTIDREAVFAYADD
Tintermedium-Thint.V0383400.1              AIDLRHLEENVKNAVKSCVKTVARKTLSTDNNGHLHNARFCEKDLLLTIDREAVFAYADD
Taestivum-Traes_2BL_13D5272D7.1            AIDLRHLEENVKNAVKSCVKTVARKTLSTDNNGHLHKARFCEKDLLLTIDREAVFAYADD
Tintermedium-Thint.06G0475700.1            AIDLRHLEENVKNAVKSCVKTVARKTLSTDNNGHLHNARFCEKDLLLTIDREAVFAYADD
Taestivum-Traes_2BL_C051606EA.2            AIDLRHLEENVKNAVKSCVKTVARKILSTDNNGHLHNARFCEKDLLLTIDRDAVFAYADD
Tintermedium-Thint.V0383600.1              AIDLRHLEENVKNAVKSCVKTVARKTLSTNSNGHLHNARFCEKDLLLTIDREAVFAYADD
Taestivum-Traes_2AL_9D78F85E2.1            AIDLRHLEENVKNAVKSCVKTVARKTLSTNNNGHLHNARFCEKDLLLTIDREAVFAYADD
Tintermedium-Thint.05G0274100.1            AIDLRHLEENVKNAVKSCVKMVARKTLSTNNNGHLHNARFCEKDLLLTIDREAVFAYADD
Tintermedium-Thint.06G0476300.1            AIDLRHLEENVKNAVKSCVKTVARKTLSTNNSGHLHNARFCEKDLLLTIDREAVFAYADD
Taestivum-Traes_2BL_5A50FDA1A.1            AIDLRHLEENVKNAVKSCVKTVARKTLSTNNNGHLHNARFCEKDLLLTIDREAVFAYADD
Tintermedium-Thint.05G0273900.1            AIDLRHLEENVKNAVKSCVKTVARKTLSTDNNGHLHNARFCEKDLLLTIDREAVFAYADD
Taestivum-Traes_2DL_30F8555D4.1            AIDLRHLEENVKNAVKSCVKTVARKTLSTDNNGHLHKARFCEKDLLLIIDREAVFVYADD
Taestivum-Traes_2DL_83168C1E0.1            AIDLRHLEENVKNAVKSCVKTVARKTLSTNNNGHLHNARFCEKDLLLTIDREAVFAYADD
Taestivum-Traes_2DL_B978B4E22.1            AIDLRHLEENVKNAVKSCVKTVARKTLSTDTNGNLHNARFCEKDLLLTIDREAVFAYADD
Taestivum-Traes_1AS_0B9295A68.1            AIDLRHLEENVKDAVKSCVKMTARKTLSTTNNGHLHNARFCEKDLLLTIDREVVFAYADD
Taestivum-Traes_1BS_7068F56E6.1            AIDLRHLEENVKDAVKNCVKMTARKTLSTTNNGHLHNARFCEKDLLLTIDREAVFAYADD
Taestivum-Traes_1BS_AEA817616.1            AIDLRHLEENVKDAVKSCVKMTARKTLSTTNNGHLHNARFCEKDLLLTIDREAVFAYADD
Taestivum-Traes_4AL_892C47ED5.1            AIDLRHLEENVKDAVKSCVKMTARKTLSTTNNGHLHNARFCEKDLLLTIDREAVFAYADD
Tintermedium-Thint.03G0055900.1            AIDLRHLEENVKDAVKSCVKMTARKTLSTTNNGHLHNARFCEKDLLLTIDREAVFAYADD
Tintermedium-Thint.01G0012900.1            AIDLRHLEENVKDAVRSCVKMTARKTLSTTNNGHLHNARFCEKDLLLTIDREAVFAYADD
Tintermedium-Thint.02G0158500.1            AIDLRHLEENVKDAVKSCVKMTARKTLSTTNNGHLHNARFCEKDLLLTIDREAVFAYADD
Tintermedium-Thint.03G0055700.1            AIDLRHLEENVKDAVKSCVKMTARKTLSTTNNGHLHNARFCEKDLLLTIDHEAVFAYADD
Taestivum-Traes_1BS_C3AA04888.1            AIDLRHLEENVKDAVKSCVKMTARKTLSTTNNGHLHNARFCEKDLLLTIDREAVFAYADD
Taestivum-Traes_1DS_A8BD91E4A.1            AIDLRHLEENVKDAVKSCVKMTARKTLSTTNNGHLHNARFCEKDLLLTIDREAVFAYADD
Taestivum-Traes_1BS_C26CCE0E9.1            AIDLRHLEENIKDAVKSCVKMAARKTLSTNHNGHLHNARFCEKDLLLTIDREAVFAYADD
Taestivum-Traes_1DS_A21FCBA3D.1            AIDLRHLEENIKDAVKSCVKMAARKTLSTNNNGHLHNARFCEKDLLLTIDREAVFAYADD
Tintermedium-Thint.01G0012200.1            AIDLRHLEENIKDAVKSCVKMAARKTLSTNNNGHLHNARFCEKDLLLTIDREAVFAYADD
Tintermedium-Thint.02G0158600.1            AIDLRHLEENIKDAVKSCVKMAARKTLSTNNNGHLHNARFCEKDLLLTIDREAVFAYADD
Tintermedium-Thint.01G0012700.1            AIDLRHLEENIKEAVKSCVKMAARKTLSTNNNGHLHNTRFCEKDLLLTIDREAVFAYADD
Taestivum-Traes_5BL_F50A47503.1            AIDLRHLEENIKEAVKSCVKMAARKTLSTNNNGHLHNARFCEKDLLLTIDREAVFAYADD
Tintermedium-Thint.02G0158700.1            AIDLRHLEENIKDAVKSCVKMAARKTLSTNNNGHLHNARFCEKDLLLTIDREAVFAYADD
Taestivum-Traes_5BL_AB528184A.1            AIDLRHLEENIKDAVKRCVNMAARKTLSTNNNGHLHNARFCEKDLLLTIDREAVFAYADN
Tintermedium-Thint.03G0055800.1            AIDLRHLEENIKDAVKSCVKMAARKTLSTTNNGHLHNARFCEKDLLLTIDREAVFTYADD
Taestivum-Traes_1AS_F9013A945.1            AIDLRHLEENVKDAVKSCVKTVARKTLSTNNNGHLHNARFCEKDLLLTIDREAVFTYADD
Taestivum-Traes_1DS_43DAC7A7F.1            AIDLRHLEENVKDAVKSCVKTVARKTLSTNNNGHLHNARFCEKDLLLTIDREAVFTYADD
Taestivum-Traes_1BS_BD86C90A7.1            AIDLRHLEENVKDAVKSCVKTVAKKTLSTNNNGHLHNARFCEKDLLLTIDREAVFTYADD
Tintermedium-Thint.02G0158900.1            AIDLRHLEENVKNAVKSCVKTVARKTLSTNNNGHLHNARFCEKDLLLTIDREAVFTYADD
Tintermedium-Thint.05G0274000.1            AIDLRHLEENVKNAVKSCVKTVARKTLSTNSNGHLHNARFYEKDLLLTIDREEVFAYADD
Osativa-LOC_Os04g43800.1                   AIDLRHLEENVRSAVKGCVTTVARKTLSTSATGDLHKARFCEKDLLQAIDREAVFAYADD
Pvaginatum-Pavag06G160300.1                AIDLRHLEENLRSAVKRCVMTVSKKTLSTSATGELHRARFCEKGLMTAIDREAVFAYADD
Sbicolor-Sobic.006G148900.1                AVDLRHLEENLKSAVKSCVTTVAKKTLSTSATGGLHNARFCEKDLLTAIDREAVFAYADD
Pvirgatum-Pavir.7KG237800.1                AIDLRHLEENLKSAVKSCVMTVAKKTLSTGATGALHSARFCEKDLLTAIDREAVFAYADD
Pvirgatum-Pavir.7NG355800.1                AIDLRHLEENLKSAVKSCVMTVAKKTLSTSATGALHSARFCEKDLLTAIDREAVFAYADD
Sviridis-Sevir.7G178300.1                  AIDLRHLEENLKGAVKSCVMTVAKKTLSTGATGALHNARFCEKDLLTAIDREAVFAYADD
Zmays-Zm00001d003015_T001                  AIDLRHLEENLRSAVKRCVTVVARKTLSTGATGALHDARFCEKDLLTAIEREAVFAYADD
Hvulgare-HORVU1Hr1G022060.1                AIDLRHLEENVKNAVKNCVTRVARKTLITNDMGGLHNARFCEKDLLQTIDREAVFAYADD
Taestivum-Traes_1BS_723922D171.2           AIDLRHLEENVKNAVKNCVTRVARKTLITNDMGGLHNARFCEKDLLQTIDREAVFAYADD
Taestivum-Traes_1DS_A171C7D59.1            AIDLRHLEENVKNAVKNCVTRVARKTLITNDMGGLHNARFCEKDLLQTIDREAVFAYADD
Tintermedium-Thint.01G0111500.1            AIDLRHLEENVKNAVKNCVTRVARKTLITNDMGGLHNARFCEKDLLQTIDREAVFAYADD
Tintermedium-Thint.03G0265300.1            AIDLRHLEENVKNAVKNCVTRVARKTLITNDMGGLHNARFCEKDLLQTIDREAVFAYADD
```

```
Hvulgare-HORVU2Hr1G038140.1                    AIDLRHLEENVKNAVKSCVTRVARKTLITNDMGGLHNARFCEKDLLQTIDREAVFAYADD
Tintermedium-Thint.04G0171200.1                AIDLRHLEENVKNAVKNCVTRVARKTLITNDMGGLHNARFCEKDLLQTIDREAVFAYADD
Taestivum-Traes_2AS_958327519.3                AIDLRHLEENVKNAVKDCVTRVVRKTLITNDMGGLHNARFCEKDLIRTIDREAVFAYADD
Taestivum-Traes_2DS_28CA50371.1                AIDLRHLEENVKNAVKNCVTRVARKTLITNDMGGLHNARFCEKDLLQTIDREAVFAYADD
Tintermedium-Thint.05G0160800.1                AIDLRHLEENVKNAVKNCVTRVARKTLITNDMGGLHNARFCEKDLLQTIDREAVFAYADD
Tintermedium-Thint.06G0299000.1                AIDLRYLEENVKNAVKNCVTRVARKTLITNDMGGLHNARFCEKDLLQTIDREAVFAYADD
Taestivum-Traes_2DS_3791A8A36.1                AIDLRHLEENVKNAVKNCVTRVARKTLITNDMGGLHNARFCEKDLLQTIDREAVFAYADD
Tintermedium-Thint.06G0342800.1                AIDLRHLEENVKNAVKNCVTRVARKTLITNDMGGLHNARFCEKDLLQTIDREAVFAYADD
Tintermedium-Thint.05G0150900.1                AIDLRHLEENVKNAVKNCVTRVARKTLITNDMGGLHNARFCEKDLLQTIDREAVFAYADD
Tintermedium-Thint.04G0242300.1                AIDLRHLEENVKNAVKNCVTRVARKTLITNDMGGLHNARFCEKDLLQTIDREAVFAYADD
Tintermedium-Thint.06G0298700.1                AIDLRHLEENVKNVVKNCVTRVARKTLITNDMGGLQNARFCEKDLLQTIDREAVFAYADD
Taestivum-Traes_2BS_88CF42F2E.1                AIDLRHLEENVKNAVKNCVTRVARKT-ITNDMGGLHNTRFCEKDLLQTIDREAVFAYADD
Tintermedium-Thint.04G0241900.1                AIDLRHLEENVKNAVKNCVTRVARKTLITNDMGGLHNARFCEKDLLQTIDREAVFAYADD
Platifolius-Phala.06G165300.1                  AIELRHIEENVKTAVKNCVTTVARKTLSTSPTGDLHIARFCEKDLLTVIDREAVFHYADD
Streptochaeta-strangu_019385-RA                AVDLRHLEENIKSAVKKCVTAAAKKALNTSTTGDLHIARFCEKDLLTAIDREAVFAYADD
Streptochaeta-strangu_028944-RA                AVDLRHLEENIKSTVKNCVTTAATKTL-----GTTHTVRFSE-DLLTAIDREAVFAYADD
Acomosus-Aco010091.1                           AIDLRHLEENLKAAVKHTVSQAAKRVLMMGADGGLQPSMFCVEGLIKVIDREHVFSYIDD
Zmarina-Zosma03g22300.1                        AVNLRHLEENMKRTVKNVVGQVAKRVLTLGVNSELHPSRFCEKDLIKVIDREQVFDYIDD
Ecdeiocolea-Emoptg0003741_1G000600.1           AIDLRHLEENLKSAIKNCVAQVAKKALTLNTAGDLHNARFSEKDLLTAIDREAVFAYADD
Jascendens-Joasc.05G060500.1                   AIDLRHLEENLKSAVKNCVAQVAKKALTLNTVGDLHNARFSEKDLLTAIDREALFAYADD
Ecdeiocolea-Emoptg0003741_1G000630.1           AIDLRHLEENLKSTVKNCVAQVAKKVLTMSTAGDLHSARFSEKGLLTAIDREAVFAYADD
Jascendens-Joasc.05G060400.1                   AIDLRHLEENLKSAVKNCVAQVAKKVLTVNTVGDLHNARFSEKDLLTAIDREAVFTYADD
Bdistachyon-Bradi3g49250.2                     AVDLRHLEENIKASVKNCVTQVSKKVLTMNPTGDLSSARFSEKSLLTAIDREAVFSYADD
Hvulgare-HORVU6Hr1G058820.1                    AIDLRHLEENIKTSVKNCVTQVSKKVLTMNPTGDLSSARFSEKSLITAIDREAVFSYADD
Taestivum-Traes_6DL_1AEA7B869.1                AIDLRHLEENIKTSVKNCVTQVSKKVLTMNPTGDLSSARFSEKSLITAIDREAVFSYADD
Tintermedium-Thint.18G0192300.1                AIDLRHLEENIKTSVKNCVTQVSKKVLTMNPTGDLSSARFSEKSLITAIDREAVFSYADD
Tintermedium-Thint.17G0230800.1                AIDLRHLEENIKTSVKNCVTQVSKKVLTMNPTGDLSSARFSEKSLITAIDREAVFSYADD
Tintermedium-Thint.17G0230900.1                AIDLRHLEENIKTSVKNCVTQVSKKVLTMNPTGDLSSARFSEKSLITAIDREAVFSYADD
Ecoracana-ELECO.r07.2AG0138000.1               AVDLRHLEENIKESVKSCVTQVAKKVLTMSPTGDLSVARFSEKNLLTAIDREGVFTYAED
Ecoracana-ELECO.r07.2BG0193650.1               AVDLRHLEENIKESVKSCVTQVAKKVLTMSPTGDLSVARFSEKNLLTAIDREGVFTYAED
Pvaginatum-Pavag04G202700.1                    AIDLRHLEENIKTSVKNTVTQVAKKVLTMNPTGDLSSARFSEKDLITAIDREGVFTYAED
Sviridis-Sevir.1G245000.1                      AIDLRHLEENIKTSVKNTVTQVAKKVLTMNPTGDLSSARFSEKDLITAIDREGVFTYAED
Sviridis-Sevir.6G187100.1                      AIDLRHLEENIKTSVKNTVTQVAKKVLTMNPAGELSSARFSEKDLLTAIDREGVFTYAED
Pvirgatum-Pavir.1NG356200.1                    AVDLRHLEENIKTSVKNTVTQVAKKVLTMNPTGDLSTARFSEKDLITAIDREAVFTYAED
Sbicolor-Sobic.004G220300.1                    AIDLRHLEENIKTSVKNTVTQVAKKVLTMNPSGDLSSARFSEKELITAIDREGVFTYAED
Zmays-Zm00001d017274_T001                      AVDLRHLEENIKASVKNTVTQVAKKVLTMNPSGELSSARFSEKELISAIDREAVFTYAED
Zmays-Zm00001d051161_T003                      AVDLRHLEENIKTSVKNTVTQVAKKVLTMNPSGDLSSARFSEKELISAIDREAVFTYAED
Sviridis-Sevir.7G178200.1                      AIDLRHLEENIKTSVKNTVTQVAKKVLTMNPVGELSSARFSEKDIITAIDREGVFTYAED
Osativa-LOC_Os02g41630.2                       AVDLRHLEENIKSSVKNCVTQVAKKVLTMNPTGDLSSARFSEKNLLTAIDREAVFSYADD
Ecoracana-ELECO.r07.4AG0310430.1               AVDLRHLEENLKAAVKGIVTSVARKVLTTSPDGELHSARFSEKALLTGIDREPVYGYYDD
Ecoracana-ELECO.r07.4BG0341620.1               AVDLRHLEENLKAAVKGTVTSVARKVLTTSPDGELHSARFSEKALLTGIDREPVYGYYDD
Pvaginatum-Pavag06G160200.1                    AVDLRHLEENLKSAVTHCVTAVARKVLTTSPDGDLHNARFSEKALLTAIDHEAVYGYYDD
Pvirgatum-Pavir.7KG238255.1                    AVDLRHLEENLKSAVKSCVTAVARRVLTTRPDGELHSARFSEKSLLTAIDREAVYGYYDD
Sbicolor-Sobic.006G148800.1                    AVDLRHLEENLKSAVKNCVMAAARKVLTTSLDGDLHSARFSEKALLTAIDREAVYGYYDD
Zmays-Zm00001d003016_T001                      AVDLRHLEENLKSAVKSCVMAVARKVLTTSLGGDLHSARFSEKALLTAIDREAVYGYYDD
Osativa-LOC_Os04g43760.1                       AVDLRHLEENLKSAVKNCVTTVAKKVLTTGPAGGLHSARFSEKALLTAIDREAVYSYADD
Platifolius-Phala.02G210400.1                  AVDLRHLEENLKNAIKNCVTQTAKKVLTMNPAGDLHNARFSEKDLLTAIDREAVFSYADD
Platifolius-Phala.06G165100.1                  AIDLRHLEENLKNSVKNCVTQVAKKVLTTGLAGDLHNARFSEKALLTAIDREAVFSYADD
Streptochaeta-strangu_019386-RA                AIDLRHLEENLKNAVKNCVMQVAKKVLTTNPAGDLHNARFSEKALLTAIDREAVFSYADD
Streptochaeta-strangu_020769-RA                AVDLRHLEENLKNAVKNCVTQVAKKVLTTNPTGDLSNARFSEKALITSIDREAVFSYADD
Sviridis-Sevir.2G448300.1                      AVDLRHLEENLKSAVKNCVKTVALKVLTTSPDGEHCSARFSEKALLAAIDRKAVYSYYDD
Sviridis-Sevir.7G177900.1                      AVDLRHLEENLKSAVKNCVKTVALKVLTTSPDGEHCSARFSEKALLAAIDRKAVYSYYDD
```

```
Aamericanus-Acora.01G232100.1                  PCSATYPLMQKLRQVLVEHAIN-NG--EKEREVGTSIF--QKIGVFEEEIKAVLPKEVEA
Aamericanus-Acora.10G174800.1                  PCSATYPLMQKLRQVLVEHALN-NG--EKEKEWSTSIF--QKISLFEEELKAVLPKEVEA
Acomosus-Aco013943.1                           PCSATYPLMQKLRQVLVEHALG-NG--EKEKATETSIF--QKIFAFEEELKAQLPKEVEA
Aofficinalis-evm.model.AsparagusV1_01.827      PCSATYPLMQKLRQVLVEHALN-NG--EKEKEASTAIF--HKILEFEEELKATLPKEVEG
Typha_latifolia-Tylat.14G083800.1              PCSATYPLMQKLRQVLVEHALN-NG--EKEKDSNCSIF--QKIALFEEELKTHLPKEVEA
Aofficinalis-evm.model.AsparagusV1_04.71       PCSYTYPLMQKVRQVLVEHALN-NG--EKERDLNTSIF--QKIAVFEEELKAVLPKEVEA
Aofficinalis-evm.model.AsparagusV1_04.93       PCSYTYPLMQKVRQVLVEHALN-NG--EKERDLNTSIF--QKIAVFEEELKAVLPKEVEA
Aofficinalis-evm.model.AsparagusV1_10.1948     PCSYNYPLMQKLRQVLVEHALN-NG--EKEKDVKTSIF--QKIAAFEEELKAVLPKEVEA
Typha_latifolia-Tylat.03G127700.1              PCSSTYPLMQKLRQVLVDHALN-NG--EKEKDAATSIF--QKITAFEEELKTQLPKEVEA
Spolyrhiza-Spipo11G0025500                     PCLSTYPLMQKLRQVLVEHAIA-NG--EKEKDSATSIF--QKVTAFEEELKMALPKEVEA
Spolyrhiza-Spipo1G0003500                      PCSATYPLMQKLRQVLVEHALT-NG--EKEKDSNTSIF--QKVAAFEEELKMTLPKEVEA
Phoenix_dactylifera-LOC103718264               PCSSTYPLMQKLRQVLVEHALN-NG--EKEKDPSSSIF--QKISVFEEELKTLLPKEVEG
Macuminata-GSMUA_Achr1T09070_001               PCSATYPLMQKLRQ-------------------------------------------E-
Acomosus-Aco006987.1                           PCSSTYPLMQKLRQVLVEQALG-NG--EKEKDPNNSIF--QKIAAFEEELKAQLPKEVEA
Acomosus-Aco027752.1                           PCSSTYPLMQKLRQVLVEQALG-NG--EKEKDPNNSIF--QKIAAFEEELKAQLPKEVEA
Cscoparia-Csco27g03980.1                       ACSATYPLMQKLRQVLVDHALS-NG--NRELDANTSIF--QKINSFEEELKIQLPKDVEA
Cscoparia-Csco27g03990.1                       ACSATYPLMQKLRQVLVDHALS-NG--DKELDANTSIF--QKINSFEEELKIHLPKDVEA
Macuminata-GSMUA_Achr11T16380_001              PCSSTYALMPKLRMAL--------------------------------------------
Macuminata-GSMUA_Achr11T22840_001              PCSSTYALMPKLRME---------------------------------------------
Macuminata-GSMUA_Achr2T00240_001               PCSSTYALMPKLRME---------------------------------------------
Macuminata-GSMUA_Achr5T18560_001               PCSSTYALMPKLRMVL--------------------------------------------
Macuminata-GSMUA_Achr8T18960_001               ------------------------------------------------------------
Macuminata-GSMUA_Achr5T03950_001               PCSSTYPLMQKLRQE---------------------------------------------
Phoenix_dactylifera-LOC103703950               PCSSTYILMEKLRQVLVEHALN-NG--EKEKDASTSIF--QKITAFEQELKAVLPKEVEA
Acoerulea-Aqcoe1G140100.1                      PCSATYPLMQKLRQVLVEHALL-NG--EKEKTSSNSIF--QKINAFEEELKNLLPKEVES
Acoerulea-Aqcoe2G144500.1                      PCLGTYPLMQKLRQVLVDHALI-NN--GKEKDFNTSIF--QKISAFEEELKATLPKEVES
Typha_latifolia-Tylat.05G053500.1              PCSSTYPLMQKLRQVLVEHALL-NS--EKE-EANTSIF--QKITAFEEELKTQLPKEVES
Zmarina-Zosma05g29470.1                        PCSATYPLMQKLRSVLVDHALN-NG--DKEKDEAMSIF--QKIAVFEEELIAALPKEVEA
Acomosus-Aco007727.1                           PCSPSYLLMQKLRQVLVEHALK-NG--EKEKNSNSSIF--QKITAFEEELKANLPKEVDA
Bvulgarisssp-Bevul.7G054300.1                  PCSVTYPLMQKLRQVLVDQALA-NG--DSEKNVSTSIF--QKIGAFEEELKARLPKEVEA
Bdistachyon-Bradi3g47110.1                     PCNPNYPLMKKLRGVLVERALA-NG--VAEFDAETSVF--AKVARFEEELRAALPVAVEA
Bdistachyon-Bradi3g47120.1                     PCNPNYPLMKKLRGVLVERALA-NG--VAEFDAETSVF--AKVARFEEELRAALPVAVEA
Bdistachyon-Bradi3g48840.1                     PCSANYPLMKKLRSVLVERALS-NG--MAEFNAETSVF--AKVAQFEEELRTVLPKAVEA
Bdistachyon-Bradi3g49280.1                     PCSANYPLMKKLRGVLVERALS-NG--KAEFNAETSVF--AKVAQFEEELRTALPKAVEA
Hvulgare-HORVU6Hr1G058840.1                    PCSPNYPLMKKLRGVLVERALS-NG--MAEFDAETSVL--AKVALFEEELRKTLPRAVEA
Taestivum-Traes_6BL_C96F626D9.1                PCSPNYPLMKKLRGVLVERALS-NG--MGEFDAETSVF--AKVALFEEELRKTLPPAVEA
Tintermedium-Thint.18G0192700.1                PCSPNYPLTKKLRGVLVERALS-NG--MAEFDAETSVF--AKVALFEEELRKTLPPAVEA
Taestivum-Traes_6DL_04CD7E82C.1                PCSPNYPLMKKLRGVLVERALS-NG--MAEFDAETSVF--AKVALFEEELRKTLPPAIEA
Bdistachyon-Bradi3g49270.1                     PCNHNYPLMKKLRGVLVESALA-NG--VAEYNVETSVF--AKVAQFEEELRAALPKAVEA
Ecoracana-ELECO.r07.2AG0138020.1               PCSANYPLMKKLRNVLVERALA-NG--NAEFNAETSVF--AKVAQFENELRQALPKAVEA
Ecoracana-ELECO.r07.2BG0193670.1               PCSENYPLMKKLRNVLVERALA-NG--PAEFNADTSVF--AKVAQFENELRQALPKAIEA
Osativa-LOC_Os02g41670.1                       PCSANYPLMKKLRNVLVERALA-NG--AAEFNAETSVF--AKVAQFEEELRAALPAAVLA
Osativa-LOC_Os02g41680.1                       PCSHNYPLMKKLRNVLVERALA-NG--AAEFNADTSVF--AKVAQFEEELRATLPGAIEA
Pvaginatum-Pavag04G202900.1                    PCSANYPLMKKLRNVLVERALA-NG--MAEFNAETSVF--AKVAQFEEELRAALPNAVEA
Pvaginatum-PavagK321000.1                      PCSANYPLMKKLRNVLVERALA-NG--MAEFNAETSVF--AKVAQFEEELRAALPNAVEA
Pvaginatum-PavagK321300.1                      PCSANYPLMKKLRNVLVERALA-NG--MAEFNAETSVF--AKVAQFEEELRAALPNAVEA
Pvaginatum-PavagK321100.1                      PCSANYPLMKKLRNVLVERALA-NG--MAEFNAETSVF--AKVAQFEEELRAALPNAVEA
Pvaginatum-PavagK321200.1                      PCSANYPLMKKLRNVLVERALA-NG--MAEFNAETSVF--AKVAQFEEELRAALPNAVEA
Pvirgatum-Pavir.1KG386500.1                    PCSANYPLMKKLRNVLVERALA-NG--AAEFNAETSVF--AKVAQFEEELRAALPKAVEA
Pvirgatum-Pavir.1NG356700.1                    PCSANYPLMKKLRNVLVERALA-NG--AAEFSAETSVF--AKVAQFEEELRAALPKAVEA
Sviridis-Sevir.1G245166.1                      PCSANYPLMKKLRNVLVERALA-NG--TAEFNAETSVF--AKVAQFEEELRAALPKAVEA
Sviridis-Sevir.1G245232.1                      PCSANYPLMKKLRNVLVERALA-NG--TAEFNAETSVF--AKVAQFEEELRAALPKAVEA
Pvirgatum-Pavir.1NG356800.1                    PCSANYPLMKKLRNVLVERALA-NG--AAEFSAETSVF--AKVAQFEEELRAALPKAVEA
Sbicolor-Sobic.004G220500.1                    PCSANYPLMKKLRNVLVERALA-NG--AAEFDAETSVF--AKVAQFEEELRAALPKAVEA
Sbicolor-Sobic.004G220600.2                    PCSANYPLMKKLRNVLVERALA-NG--AAEFDAETSVF--AKVAQFEEELRAALPKAVEA
Sbicolor-Sobic.004G220700.1                    PCSANYPLMKKLRNVLVERALA-NG--AAEFDAETSVF--AKVAQFEEELRAALPKAVEA
Sviridis-Sevir.1G245300.2                      PCSANYPLMKKLRNVLVERALA-NG--TAEFNADTSVL--AKVAQFEEELRAALPNAVEA
Zmays-Zm00001d017276_T001                      PCSANYPLMKKLRNVLVERALA-NG--AAEFNAETSVF--AKVAQFEEDLRAALPKAVEA
Zmays-Zm00001d017279_T001                      PCSANYPLMKKLRNVLVERALA-NG--AAEFNAETSVF--AKVAQFEEDLRAALPKAVEA
Zmays-Zm00001d051166_T001                      PCSANYPLMKKLRNVLVERALA-NG--TAEFDAETSVF--AKVAQFEEELRTALPSAVEA
Ecoracana-ELECO.r07.2AG0138030.1               PCSANYPLMKKLRNVLVERALA-NG--NAEFNAETSVF--AKVAQFEEELRATLPKAVEA
Ecoracana-ELECO.r07.2BG0193680.1               PCSANYPLMKKLRNVLVERALD-NG--NAEFNAETSVF--AKVAQFEEELRATLPKAVEA
Osativa-LOC_Os05g35290.1                       PCSANYPLMKKLRSVLVERALA-NG--AAEFDAETSVL--AKVARFEEELRAALPRAVEA
Platifolius-Phala.02G210700.1                  PCSANYPLMQKLRAVLVERALA-NG--KDEFNVETSVF--AKVAEFEEELRAALPKAVEG
Ecoracana-ELECO.r07.4AG0310420.1               PCSAAYPLMKKLRNVLVERALA-NG--MDEFNAETSVF--AKVAQFEEELRATLPKAVEA
Ecoracana-ELECO.r07.4BG0341610.1               PCSAAYPLMKKLRNVLVERALA-NG--MDEFNAETSVF--AKVAQFEEELRATLPKAVEA
Bdistachyon-Bradi3g49260.1                     PCSPNYPLMQKLRAVLIEHALA-NG--DGERALETSIF--AKVAEFEQNLRAALPKEVEA
Hvulgare-HORVU0Hr1G016330.1                    PCSPNYPLMQKLRAVLVEHALA-NG--DGERALETSIF--AKVAEFEQNIRAALPKEVEA
Tintermedium-Thint.17G0260200.1                PCSPNYPLMQKLRAVLVEHALA-NG--DGERALETSIF--AKVAEFEQNIRAALPKEVEA
Tintermedium-Thint.18G0192600.1                PCSPNYPLMQKLRAVLVEHALA-NG--DGERALETSIF--AKVAEFEQNIRAALPKEVEA
Tintermedium-Thint.17G0231000.1                PCSPNYPLMQKLRAVLVEHALA-NG--DGERALETSIF--AKVAEFEQNIRAALPKEVEA
Ecoracana-ELECO.r07.2AG0138010.1               PCSPNYPLMQKLRAVLIEHALA-NG--DAERVAETSIF--AKVAEFEQQVRAALPKEIEA
Ecoracana-ELECO.r07.2BG0193660.1               PCSPNYPLMQKLRAVLIEHALA-NG--DAERVAETSIF--AKVAEFEQQVRAALPKEIEA
Osativa-LOC_Os02g41650.3                       PCSANYPLMQKLRAVLIEHALA-NG--DAERVLETSIF--AKVAEFEQHVRAALPKEVEA
```

```
Pvaginatum-Pavag04G202800.1                   PCSPNYPLMQKLRAVLIEHALA-NG--AAERDVETSIF--AKVAEFEQQVRAALPKEVEA
Pvaginatum-PavagK320900.1                     PCSPNYPLMQKLRAVLIEHALA-NG--AAERDVETSIF--AKVAEFEQQVRAALPKEVEA
Pvirgatum-Pavir.1NG356400.1                   PCSPNYPLMQKLRAVLIEHALA-NG--DAERVVETSIF--AKVAEFEQQVRAALPKGVEA
Sviridis-Sevir.1G245100.1                     PCSPNYPLMQKLRAVLIEHALA-NG--DAERVVETSIF--AKVAEFEQQVRAALPKEVEA
Sbicolor-Sobic.004G220400.1                   PCSPNYPLMQKLRAVLIEHALA-NG--DAERVVETSIF--AKVAEFEQQVRAALPKEVEA
Zmays-Zm00001d017275_T001                     PCSPNYPLMQKLRAVLVEHALA-NG--DAERAAETSIF--AKVAEFEQQVRAALPKEVEA
Zmays-Zm00001d051163_T002                     PCSPNYALMQKLRAVLVEHALA-NG--DAERDVDTSIF--AKVAEFEQQVRAALPKEVEA
Pvirgatum-Pavir.1KG386300.1                   PCSPNYPLMQKLRAVLIEHALA-NG--DAERVAETSIF--AKITKFEEELRAVLPREVEA
Platifolius-Phala.02G210600.1                 PCSASYPLMQKLRAVLVEHALA-NG--QAERVLETSIF--AKVAEFEQQLRAALPKEVEA
Streptochaeta-strangu_020767-RA               PCNASYPLMKNLRAVLVEHALA-NG--EAERNVDTSVF--TKVAEFEQQLRAALPKEVEA
Bdistachyon-Bradi5g15830.1                    PCSANYPLMQKMRAVLVEHALA-NG--EAERDVETSVF--AKLAAFEQELRAVLPKEVEA
Hvulgare-HORVU2Hr1G089440.4                   PCSANYPLMQKMRAVLVEHALA-NG--EAEHDVETSVF--AKLATFEQELRAVLPKEVEA
Taestivum-Traes_2AL_0DDC60D6E.1               PCSANYPLMQKMRAVLVEHALA-NG--EAERDVETSVF--AKLATFEQELRAVLPKEVEA
Tintermedium-Thint.06G0475900.1               PCSANYPLMQKMRAVLVEHALA-NG--EAERDVETSVF--AKLATFEQELRAVLPKEVEA
Tintermedium-Thint.06G0476200.1               PCSANYPLMQKMRAVLVEHALA-NG--EAERDVETSVF--AKLATFEQELRAVLPKEVEA
Taestivum-Traes_2AL_2763280FC.1               PCSANYPLMQKMRAVLVEHALA-NG--EAERDVETSVF--AKLATFEQELRAVLPKEVEA
Taestivum-Traes_2DL_8C21AAAEF.1               PCSANYPLMQKMRAVLVEHALA-NG--EAERDVETSVF--AKLATFEQELRAVLPKEVEA
Tintermedium-Thint.V0383500.1                 PCSANYPLMQKMRAVLVEHALA-NG--EAERDVETSVF--AKLATFEQELRAVLPKEVEA
Taestivum-Traes_2AL_9EC3226F7.1               PCSANYPLMQKMRAVLVEHALA-NG--EAERDVETSVF--AKLAMFEQELRAVLPKEVEA
Tintermedium-Thint.05G0273800.1               PCSANYPLMQKMRAVLVEHALA-NG--EAERDVETSVF--AKLAMFKQELRAVLPKEVEA
Tintermedium-Thint.V0383400.1                 PCSANYPLMQKMRAVLVEHALA-NG--EAERDVETSVF--AKLAMFEQELRAVLPKEVEA
Taestivum-Traes_2BL_13D5272D7.1               PCSANYPLMQKMRAVLVEHALA-NG--EAERDVETSVF--AKLAMFEQELRAVLPKEVEA
Tintermedium-Thint.06G0475700.1               PCSANYPLMQKMRAVLVEHALA-NG--EAEREVETSVF--AKLAMFEQELRAVLPKEVEA
Taestivum-Traes_2BL_C051606EA.2               PCSANYPLMQKMRAVLVEHALA-NG--EAEHDMETSVF--AKLAMFEQELRAVLPKEVEA
Tintermedium-Thint.V0383600.1                 PCSANYPLMQKMRAVLVEHALA-NG--EAERDVETSVF--AKLAMFEQELRAVLPKEVEA
Taestivum-Traes_2AL_9D78F85E2.1               PCSANYPLMQKMRAVLVEHALA-NG--EAERDVETSVF--AKLAIFEQELRAVLPKEVEA
Tintermedium-Thint.05G0274100.1               PCSANYPLMQKMRAVLVEHALA-NG--EAERDVETSVF--AKLAMFEHELRAVLPKEVEA
Tintermedium-Thint.06G0476300.1               PCSANYPLMQKMRAVLVEHALA-NG--EAERDVETSVF--AKLAIFEQELRAVLPKEVEA
Taestivum-Traes_2BL_5A50FDA1A.1               PCSANYPLMQKMRAVLVEHALA-NG--EAERDAETSVF--AKLAIFEQELRAVLPKEVEA
Tintermedium-Thint.05G0273900.1               PCSANYPLTQKMRAVLVEHALA-NG--EAERDVETSVF--AKLAMFEQELRAVLSKEVEA
Taestivum-Traes_2DL_30F8555D4.1               PCSANYPLMQKMRAVLVEHALA-NG--EAERDVETSVF--AKLAMFEQELRAVLPKEVEA
Taestivum-Traes_2DL_83168C1E0.1               PCSANYPLMQKMRAVLVEHALA-NG--EAERDVETSVF--AKLAIFEQELRAVLPKEVEA
Taestivum-Traes_2DL_B978B4E22.1               PCSANYPLMQKMRAVLVEHALA-NG--EAERDVETSVF--AKLATFEQELRAVLPKEVEA
Taestivum-Traes_1AS_0B9295A68.1               PCSANYPLMQKMRAVLVEHTLA-NG--EAELDVETSVF--AKLAAFEQELRAVLPKEVEA
Taestivum-Traes_1BS_7068F56E6.1               PCSANYPLMQKMRAVLVEHALA-NG--EAELDVESSVF--AKLAAFEQELRAELPKEVEA
Taestivum-Traes_1BS_AEA817616.1               PCSANYPLMQKMRAVLVEHALA-NG--EAERDVETSVF--AKLAAFEQELRAVLPKEVEA
Taestivum-Traes_4AL_892C47ED5.1               PCSANYPLMQKMRAVLVEHALA-NG--EAERDVETSVF--AKLAAFEQELRAVLPKEVEA
Tintermedium-Thint.03G0055900.1               PCSANYPLMQKMRAVLVEHALA-NG--EAERDVETSVF--AKLAAFEQELRAVLPKEVEA
Tintermedium-Thint.01G0012900.1               PCSANYPLMQKMRAVLVEHALA-NG--EAERDVETSVF--AKLAAFEQELRAVLPKEVEA
Tintermedium-Thint.02G0158500.1               PCSANYPLMQKMRAVLVEHALA-NG--EAERDVETSVF--AKLAAFKQELRAVLPKEVEA
Tintermedium-Thint.03G0055700.1               PCSANYPLMQKLRAVLVEHALA-NG--EAEHDVETSVF--AKLAAFEHELRAVLPKEVEA
Taestivum-Traes_1BS_C3AA04888.1               PCSPNYPLMQKMRVVLVEHALA-NG--EAERDTETSVF--AKLAAFEQELRAVLPKEVEA
Taestivum-Traes_1DS_A8BD91E4A.1               PCSANYPLMQKMRAVLVEHALA-NG--EAEHDMETSVF--AKLAAFEQELRAVLPKEVEA
Taestivum-Traes_1BS_C26CCE0E9.1               PCSSNYPLMQKMRAVLVEHALA-NG--EAERDVETSVF--AKLAAFEQELRAVLPREVEA
Taestivum-Traes_1DS_A21FCBA3D.1               PCSSNYPLMQKMRAVLVEHALA-NG--EAERDVETSVF--AKLATFEQELRAVLPREVEA
Tintermedium-Thint.01G0012200.1               PCSFNYPLMQKMRAVLVEHALA-NG--EAERDVETSVF--AKLAAFEQELRAVLPREVEA
Tintermedium-Thint.02G0158600.1               PCSSNYPLMQKMRAVLVEHALA-NG--EAERDVETSVF--AKLAAFEQELRAVLPKEVEA
Tintermedium-Thint.01G0012700.1               PCSANYPLMQKMRAVLVEHALA-NG--EAERDVETSVF--AKLAAFEQELSAVLPKEVEA
Taestivum-Traes_5BL_F50A47503.1               PCSANYPLMQKMRAVLVEHALA-NG--EAERNVETSVF--AKLAAFEQELRAVLPKEVEA
Tintermedium-Thint.02G0158700.1               PCSSNYPLMQKMRAVLVEHALA-NG--EAERDVETSVF--AKLAAFEQELRAVLPKEVEA
Taestivum-Traes_5BL_AB528184A.1               PCSSNYPLMQKMRAVLVEHALA-NG--EAERNVETSVF--AKLAAFEQELHAVLPKEVEA
Tintermedium-Thint.03G0055800.1               PCSVNYPLMQKMRAVLVEHALA-NG--EAERDMETSVF--AKLATFEQELHAVLPKEVEA
Taestivum-Traes_1AS_F9013A945.1               PCSANYPLMQKMRAVLVEHALA-NG--DAEHDVETSVF--AKLAMFEQELRVVLPKEVEA
Taestivum-Traes_1DS_43DAC7A7F.1               PCSANYPLMQKMRAVLVEHALA-NG--DAEHDVETSVF--AKLAMFEQELRVVLPKEVEA
Taestivum-Traes_1BS_BD86C90A7.1               PCSANYPLMQKMRAVLVEHALA-NG--DAEHDVETSVF--AKLAMFEQELRVVLPKEVEA
Tintermedium-Thint.02G0158900.1               PCSANYPLMQKMRAVLVEHALA-NG--EAERDVETSVF--AKLAAFEEELRAVLPTEVEA
Tintermedium-Thint.05G0274000.1               PCSANYPLMQKMRAVLVEHALA-ND--EAEHDVETSVF--AKLAMFEQELRAVLPKEVEA
Osativa-LOC_Os04g43800.1                      PCSANYPLMQKMRAVLIEHALA-NG--EAERNVDTSVF--AKVATFEEELRVALPREVEA
Pvaginatum-Pavag06G160300.1                   PCSASYPLMQKMRSVLVEHALA-NG--DAERDPDTSVF--AKVATFEEELRAALPREVEA
Sbicolor-Sobic.006G148900.1                   PCSANYPLMQKMRSVLVEHALA-NG--EAERDPETSVF--AKVATFEEELRAALPREVDA
Pvirgatum-Pavir.7KG237800.1                   PCSANYPLMQKMRSVLVEHALA-NG--EAEHDPDTSVF--AKLATFEEELRAALPREVDA
Pvirgatum-Pavir.7NG355800.1                   PCSTNYPLMQKMRSVLVEHALA-NG--EAERDPDTSVF--GKVATFEEELRAALPPEVEA
Sviridis-Sevir.7G178300.1                     SCSANYPLMQKMRSVLVEHALA-NG--EAERDPETSVF--AKVATFEEEIRAALPREVEA
Zmays-Zm00001d003015_T001                     PCSATYPLMQKMRSVLVEHALA-NG--EAERDPDTSVF--AKVATFEEELRAALPREVDA
Hvulgare-HORVU1Hr1G022060.1                   PCSANYPLMKKMRAVLVEHALA-NG--EAERNAETSVF--AKVATFEQELCAALPQEVEA
Taestivum-Traes_1BS_723922D171.2              PCSANYPLMKKMRAVLVEHALA-NG--EAERNAETSVF--AKVATFEQELCAALPQEVEA
Taestivum-Traes_1DS_A171C7D59.1               PCSANYPLMKKMRAVLVEHALA-NG--EAERNKETSVF--AKVATFEQELCAALPQEVEA
Tintermedium-Thint.01G0111500.1               PCSANYPLMKKMRAVLVEHALA-NG--EAERNKETSVF--AKVATFEQELCAALPQEVEA
Tintermedium-Thint.03G0265300.1               PCSANYPLMKKLRAVLVEHALA-NG--EAERTVETSVF--AKVATFEQELCAALPQEAEA
Hvulgare-HORVU2Hr1G038140.1                   PCSANYPLMKKMRAVLVEHALA-NG--EAERNVETSVF--AKVAKFEQELCATLPQEVEA
Tintermedium-Thint.04G0171200.1               PCSANYPLMKKMRAVLVEHALA-NG--EAERNVETSVF--AKVATFEQELCAALPQEVEA
Taestivum-Traes_2AS_958327519.3               PCSANYPLMKKMRAVLVEHALA-NG--EAERNAETSVF--AKVAKFEQELCATLPQEVEA
Taestivum-Traes_2DS_28CA50371.1               PCSANYPLMKKMRAVLVEHALA-NG--EAERNVETSVF--AKVAKFEQELCATLPQEVEA
Tintermedium-Thint.05G0160800.1               PCSANYPLMKKMRAVLVEHALA-NG--EAERNVETSVF--AKVAKFEQELCATLPQEVEA
Tintermedium-Thint.06G0299000.1               PCSANYPLMKKMRAVLVEHALA-NG--EAERNVETSVF--AKVTKFEQELCATLPQEVEA
Taestivum-Traes_2DS_3791A8A36.1               PCSANYPLMKKMRAVLVEHALA-NG--EAERNVETSVF--AKVAKFEQEVCATLPQEVEA
Tintermedium-Thint.06G0342800.1               PCSANYPLMKKMRAVLVEHALA-NG--EAERNVETSVF--AKVAKFEQELCATLPQEVEA
Tintermedium-Thint.05G0150900.1               PCSANYPLMKKMRAVLVEHALA-NG--EAERNVETSVF--AKVAKFEQELCATLPQEVEG
Tintermedium-Thint.04G0242300.1               PCSANYPLMKKMRAVLVEHALA-NG--EAERNVKTSVF--AKVAKFEQELCATLPQEVEA
Tintermedium-Thint.06G0298700.1               PCSANYPLMKKMRAVLVEHALA-NG--EAERNVETSVF--AKVAKFEQELCATLPQEVEA
Taestivum-Traes_2BS_88CF42F2E.1               PCSANYPLMKKMRAVLVEHALA-NG--EAERNVETSVF--AKVAKFEQELCATLPQEVEA
Tintermedium-Thint.04G0241900.1               PCSANYPLMKKMRAMLVEHALA-NG--EAERNVETSVF--AKVAKFEQELCATLPQEVEA
Platifolius-Phala.06G165300.1                 ACNANYPLMQKLRAVLVEHALA-NG--QAERDVNSSVF--AKIAQFEEEMEAALPKEVEA
Streptochaeta-strangu_019385-RA               PCSNNYPLMQKLREVLVEHALA-NG--EAERNVETSVF--AKIAMFEEELRAVLPREVEA
Streptochaeta-strangu_028944-RA               PCSPKYPLMKNLRAVLLERALA-NG--EAERDVETSVL--AKLASFEEELHAELPKAVEA
Acomosus-Aco010091.1                          PCSSAYPLVQKLRQALVEHALS-RG--EKEKDNDSSTFFLHRITAFEEELSAHLPKDVEA
Zmarina-Zosma03g22300.1                       PCNATYPLMEKLRGVLVDHALG-NW--EKETDVSTSIF--QKIPTFEAELKLTLPKEVDA
Ecdeiocolea-Emoptg0003741_1G000600.1          PCSPNYPLMKKLRSVLVEHALA-NG--EAELVAETSVF--AKIAKFEEELRAALPSEVEA
Jascendens-Joasc.05G060500.1                  PCNPNYPLMQKLRAVLVEHALA-NG--EAEHVATTSVF--AKITKFEEELRATLPKEVEA
Ecdeiocolea-Emoptg0003741_1G000630.1          PCSPNYPLMQKVRAVLVEHGLA-NG--EAERVTDTSVF--AKITKFEDELRAALPKEVEA
Jascendens-Joasc.05G060400.1                  PCSPNYPLMQKVRAVLVEHALA-NG--EAERVANTSVF--AKITKFEEELRATLPKEVEA
Bdistachyon-Bradi3g49250.2                    ACSANYPLMQKLRAVLVDHALT-SSGVDNAGESEATVF--SKINKFEEELRAALPREIEA
Hvulgare-HORVU6Hr1G058820.1                   ACSANYPLMQKLRAVLVDHALT-AG--PGEAESEATVF--SKINKFEEELRSALPREIEA
Taestivum-Traes_6DL_1AEA7B869.1               ACSANYPLMQKLRAVLVDHALT-AG--PGEAESEATVF--SKINKFEEELRSALPREIEA
Tintermedium-Thint.18G0192300.1               ACSANYPLMQKLRAVLVDHALT-AG--PGEAESEATVF--SKINKFEEELRSALPREIEA
Tintermedium-Thint.17G0230800.1               ACSANYPLMQKLRAVLVEHALT-AG--PGEAESEATVF--SKINKFEEELRSALPREIEA
Tintermedium-Thint.17G0230900.1               ACSANYPLMQKLRAVLVEHALT-AG--PGEAESEATVF--SKINKFEEELRSALPREIEA
Ecoracana-ELECO.r07.2AG0138000.1              PCCGTLPLMTKLRAVLVDHALT-SG--EAEREP--SVF--SKITKFEEELRAVLPQEVEA
Ecoracana-ELECO.r07.2BG0193650.1              PCCGTLPLMTKLRAVLVDHALT-SG--EAEREP--AVF--SKITKFEEELRAVLPQEVEA
Pvaginatum-Pavag04G202700.1                   AASASLPLMQKLRAVLVDHALS-SG--EAEREP--SVF--SKITKFEEELRAVLPREVEA
Sviridis-Sevir.1G245000.1                     AASASLPLMQKLRAVLVDHAFS-SG--D---EP--SMF--SKITKFEEELRAVLPQEVEA
Sviridis-Sevir.6G187100.1                     AASASLPLMQKLRAVLVDHAFS-SG--D---EP--SVF--SKITKFEEELRAVLPQEVEA
Pvirgatum-Pavir.1NG356200.1                   AASASLPLMQKLRAVLVDHALS-SG--DAEREP--SVF--SKITKFEEELRAVLPREVEA
Sbicolor-Sobic.004G220300.1                   PASASLPLMTKLRAVLVDHALS-SG--DAEREP--SVF--SKITKFEEELRAVLPREVEA
Zmays-Zm00001d017274_T001                     AASGSLPLMQKLRAVLVDHALS-SG--DAEREP--SVF--SKITRFEEELRAVLPQEVEA
Zmays-Zm00001d051161_T003                     AASASLPLMQKLRAVLVDHALS-SG--DAEREP--SVF--SRITKFEEELRAVLPQEVEA
Sviridis-Sevir.7G178200.1                     AASASLPLMQKLRAVLVDHALS-SG--DAEREP--SVF--SKITKFEEELRAVLPREVEA
Osativa-LOC_Os02g41630.2                      PCSANYPLMQKLRAVLVEHALT-SG--DAEPEA--SVF--SKITKFEEELRSALPREIEA
Ecoracana-ELECO.r07.4AG0310430.1              PCGANSALMKKIRAVLVDHALA-NG--EADKEPSASVF--SKITKFEEELREALPREMEA
Ecoracana-ELECO.r07.4BG0341620.1              PCGANSALMKKIRAVLVDHALA-NG--EADKEPSASVF--SKITKFEEELREALPREMEA
Pvaginatum-Pavag06G160200.1                   PCSASSSLMKKIRAVLVDHALA-NG--EAEKDASSSVL--SKINRFEAELRESLPRDFEA
Pvirgatum-Pavir.7KG238255.1                   PCGANSPLMKKIRAVLVDHALA-NG--EAEKDASASVL--SKINKLEEELREALPREMEA
Sbicolor-Sobic.006G148800.1                   PCSANSPLMKKIRAVLVDHALA-SG--EAEKDASASVF--SKINRFEEALREALPREMEA
Zmays-Zm00001d003016_T001                     PCSANSPLMKKIRAVLVDHALA-SG--EAEKDASASVF--SKINRFEEELREALPREMEA
Osativa-LOC_Os04g43760.1                      PCSANYPLMTKIRAVLVEHALA-NG--PAEKDDGSSVF--SKITAFEEELREALPREMEA
Platifolius-Phala.02G210400.1                 PCSSNYPLMQKLRGVLVDHAIA-NG--EAEKEPATSVF--AKITKFEEELRSALPREIED
Platifolius-Phala.06G165100.1                 PCSANYPLMQKLRSVLVDHAIA-NG--EAEKEPGTSVF--AKITKFEEELRTALPREMEA
Streptochaeta-strangu_019386-RA               PCSANYPLMQKLRSVLVDHAIA-NG--EDEKEPATSVF--AKITKFEDELRTALPREMEA
Streptochaeta-strangu_020769-RA               PCSANYPLMQKIRAVLVDHAIT-NG--DGENEPAAAVF--ARITKFEDELRSALPREMEA
Sviridis-Sevir.2G448300.1                     PCSASSSLMMTIRAVLVDHALA-NG--EAENEARAPIF--SKITKFEEELREALPREMEK
Sviridis-Sevir.7G177900.1                     PCSASSSLMMTIRAVLVDHALA-NG--EAENEAGAPIF--SKITKFEEELREALPREMEK

Aamericanus-Acora.01G232100.1                 ARVAYEAG-NASVGNRIAECRSYPLYRFVREELKTGLLTGEKV---------------RS
Aamericanus-Acora.10G174800.1                 ARAAYEAG-NSSVSNRIEECRSYPLYKFVREELKTGLLTGEKV---------------RS
Acomosus-Aco013943.1                          VRAAVESG-SSAIANRIKECRSYPLYRFVREELGTEYLTGEKV---------------RS
Aofficinalis-evm.model.AsparagusV1_01.827     VRVAFENE-CLAIPNRIKECRSYPLYRLVREELRAGYLTGEKV---------------TS
Typha_latifolia-Tylat.14G083800.1             TRIAFDNG-SLAIPNRIKECRSYPLYRLVREELKTGYLTGEKV---------------RS
```

```
Aofficinalis-evm.model.AsparagusV1_04.71           ARVAVENG-SAAIANRIKECRSYPLYRFVREELGTELLTGEKV---------------RS
Aofficinalis-evm.model.AsparagusV1_04.93           ARVAVENG-SAAIANRIKECRSYPLYRFVREELGTELLTGEKV---------------RS
Aofficinalis-evm.model.AsparagusV1_10.1948         ARVAVEAG-NAQIGNRIKECRSYPLYKFVREELGTGLLTGEKV---------------RS
Typha_latifolia-Tylat.03G127700.1                  ARVAVENG-SSAIPNRIKECRSYPLYRFVREELGTEYLTGEKT---------------RS
Spolyrhiza-Spipo11G0025500                         AREAVEKG-APAIPNRIEDCRSYPLYKFVREELGTAMLTGEKV---------------RS
Spolyrhiza-Spipo1G0003500                          VRAAFDNG-TAVIPNRIKECRSYPLYKFVREELGTSLLTGEKV---------------RS
Phoenix_dactylifera-LOC103718264                   VRVAFENG-NSAIANRIKECRSYPLYRFVREELNTGFLTGEKV---------------RS
Macuminata-GSMUA_Achr1T09070_001                   -------------------CRSYPLYRLVREELKTGFLTGEKV---------------TS
Acomosus-Aco006987.1                               ARVAFENG-TSAIPNRIQECRSYPLYRFVREELGAGYLTGEKV---------------RS
Acomosus-Aco027752.1                               ARVAFENG-TSAIANRIQECRSYPLYRFVREELGAGYLTGEKV---------------RS
Cscoparia-Csco27g03980.1                           TRVAFDKG-TLAIPNRIKECRSYPLYRFVREELCSGYLSGENV---------------VS
Cscoparia-Csco27g03990.1                           TRVAFDKG-TLVIPNRIKECRSYPLYRFVREELGSEYLSGEKV---------------VS
Macuminata-GSMUA_Achr11T16380_001                  -------------------CRSYPLYRFVREELGTEYLTGEKV---------------RS
Macuminata-GSMUA_Achr11T22840_001                  -------------------CRSYPLYRFVRKELGAAYLTGEKV---------------RS
Macuminata-GSMUA_Achr2T00240_001                   -------------------CRSYPLYRFVREELGTACLTGEKV---------------RS
Macuminata-GSMUA_Achr5T18560_001                   -------------------CRSYPLYRLVREELGVAYLTGEKA---------------RS
Macuminata-GSMUA_Achr8T18960_001                   -------------------CRSYPLYRFVREELGTAYLTGEKV---------------RS
Macuminata-GSMUA_Achr5T03950_001                   -------------------CRSYPLYRFVRAELSTGYLTGEKV---------------RS
Phoenix_dactylifera-LOC103703950                   TRVAFENG-NLPIENRIKECRSYPLYRFVREELGAGYLTGEKV---------------RS
Acoerulea-Aqcoe1G140100.1                          TRTLYETG-TLPIANQIKECRSYPLYKFVREELDTGLLTGEKT---------------LS
Acoerulea-Aqcoe2G144500.1                          ARSACESG-NPTIPNRIKECRSYPLYRFVREELGTGLLTGEKV---------------LS
Typha_latifolia-Tylat.05G053500.1                  GRIAFENGTTSAIKNRIEECRSYPLYKFVREELGTGYLTGEKV---------------KS
Zmarina-Zosma05g29470.1                            ARLAVEAG-KPAIANRIEECRSYPLYKFVREELRTGLLTGEKV---------------RS
Acomosus-Aco007727.1                               ARVAFETG-SSAIPNRVKECRSYPLYRFIREELGTEYLTGERV---------------RS
Bvulgarisssp-Bevul.7G054300.1                      ARAAYESG-NSVIPNRIKECRSYPVYKFIREELNTNLLTGEKV---------------IS
Bdistachyon-Bradi3g47110.1                         ARAAVESG-TAEAPNRIAECRSYPLYRFVRQELGTVYLTGEKT---------------RS
Bdistachyon-Bradi3g47120.1                         ARAAVESG-TAEAPNRIAECRSYPLYRFVRQELGTVYLTGEKT---------------RS
Bdistachyon-Bradi3g48840.1                         ARAAVESG-TAATPNRIKECRSYPLYRFVREELGTAYLTGEKT---------------RS
Bdistachyon-Bradi3g49280.1                         ARSAVESG-TAATPNRIKECRSYPLYRFVREELGTAYLTGEKT---------------RS
Hvulgare-HORVU6Hr1G058840.1                        ARAAVESG-TAEMPNRITECRSYPLYRFVREELGTVYLTGEKT---------------RS
Taestivum-Traes_6BL_C96F626D9.1                    ARAAVESG-TAEMPNRIIECRSYPLYRFVREELGTVYLTGEKT---------------LS
Tintermedium-Thint.18G0192700.1                    ARAAVESG-TAEMPNRITECRSYPLYRFVREELGTVYLTGEKT---------------LS
Taestivum-Traes_6DL_04CD7E82C.1                    ARAAVESG-TAEMPNRITECCSYPLYRFVREELGTVYLTGEKT---------------LS
Bdistachyon-Bradi3g49270.1                         ARAAVESG-TAATPNRITECRSYPLYRFVREELGTVFLTGEKT---------------RS
Ecoracana-ELECO.r07.2AG0138020.1                   ARAAVENG-TAAIPNRIKECRSYPLYRFVREELGTEYLTGEKT---------------RS
Ecoracana-ELECO.r07.2BG0193670.1                   ARAAVENG-AAAIPNRIKECRSYPLYRFVREELGTEYLTGEKT---------------RS
Osativa-LOC_Os02g41670.1                           ARAAVENG-TAATPNRITECRSYPLYRFVREELGTAYLTGEKT---------------RS
Osativa-LOC_Os02g41680.1                           ARAAVENG-TAAIPSRITECRSYPLYRFVREELGTKYLTGEKT---------------RS
Pvaginatum-Pavag04G202900.1                        ARAAVESG-TAAIPNRIAECRSYPLYRFVREELGAVYLTGEKT---------------RS
Pvaginatum-PavagK321000.1                          ARAAVESG-TAAIPNRIAECRSYPLYRFVREELGAVYLTGEKT---------------RS
Pvaginatum-PavagK321300.1                          ARAAVESG-TAAIPNRIAECRSYPLYRFVREELGAVYLTGEKT---------------RS
Pvaginatum-PavagK321100.1                          ARAAVESG-TAAIPNRIAECRSYPLYRFVREELGAVYLTGEKT---------------RS
Pvaginatum-PavagK321200.1                          ARAAVESG-TAAIPNRIAECRSYPLYRFVREELGAVYLTGEKT---------------RS
Pvirgatum-Pavir.1KG386500.1                        ARASVENG-TAAIPNRIAECRSYPLYRFVREELGAVYLTGEKA---------------RS
Pvirgatum-Pavir.1NG356700.1                        ARAAVENG-TAAIPNRIAECRSYPLYRFVREELGAVYLTGEKT---------------RS
Sviridis-Sevir.1G245166.1                          ARAAVENG-TAAIPIRITECRSYPLYRFVREELGAVYLTGEKT---------------RS
Sviridis-Sevir.1G245232.1                          ARAAVENG-TAAIPIRITECRSYPLYRFVREELGAVYLTGEKT---------------RS
Pvirgatum-Pavir.1NG356800.1                        ARAAVENG-TAAIPNRIAECRSYPLYRFVREELGAVYLTGEKT---------------RS
Sbicolor-Sobic.004G220500.1                        ARAAVENG-TAAIPNRITECRSYPLYRFVRQEVGAVYLTGEKT---------------RS
Sbicolor-Sobic.004G220600.2                        ARAAVENG-TAAIPNRITECRSYPLYRFVRQEVGAVYLTGEKT---------------RS
Sbicolor-Sobic.004G220700.1                        ARAAVENG-TAAIPNRITECRSYPLYRFVRQEVGAVYLTGEKT---------------RS
Sviridis-Sevir.1G245300.2                          ARAAVENG-TAAIPNRITECRSYPLYRFVREELGAVYLTGEKT---------------RS
Zmays-Zm00001d017276_T001                          ARAAVENG-TAAIPNRITDCRSYPLYRFVREELGAVYLTGEKT---------------RS
Zmays-Zm00001d017279_T001                          ARAAVENG-TAGIPNRIAECRSYPLYRFVREELGAVYLTGEKT---------------RS
Zmays-Zm00001d051166_T001                          ARAAVENG-TAAIPNRIAECRSYPLYRFVREELGAVYLTGEKT---------------RS
Ecoracana-ELECO.r07.2AG0138030.1                   ARAAVEKG-TAPMPNRIKECRSYPLYRFVREELGTEYLTGEKT---------------RS
Ecoracana-ELECO.r07.2BG0193680.1                   ARAAVEKG-TAPIPNRIKECRSYPLYRFVREELGTEYLTGEKT---------------RS
Osativa-LOC_Os05g35290.1                           ARAAVESG-TAAAPNRIAECRSYPLYRFVREELGTAYLTGEKT---------------RS
Platifolius-Phala.02G210700.1                      ARAVVENG-IAPTPNRIIECRSYPLYRFVREELGTEYLTGEKT---------------RS
Ecoracana-ELECO.r07.4AG0310420.1                   ARAAVESG-AAPIPNRIRECRSYPVYRFVREELGTEYLTGEKT---------------RS
Ecoracana-ELECO.r07.4BG0341610.1                   ARAAVESG-AAPIPNRIRECRSYPLYRFVREELGTEYLTGEKT---------------RS
Bdistachyon-Bradi3g49260.1                         ARASVENG-TPLAPNRIKDCRSYPLYRFVREELGTEYLTGEKT---------------RS
Hvulgare-HORVU0Hr1G016330.1                        ARASVENG-TPLAPNRIKDCRSYPLYQFVREVCGTEYLTGEKT---------------RS
Tintermedium-Thint.17G0260200.1                    ARASVENG-TPLAPNRIKDCRSYPLYQFVREVCGTEYLTGEKT---------------RS
Tintermedium-Thint.18G0192600.1                    ARASVENG-TPLAPNRIKDCRSYPLYQFVREVCGTEYLTGEKT---------------RS
Tintermedium-Thint.17G0231000.1                    ARASVENG-TPLAPNRIKDCRSYPLYQFVREVCGTEYLTGEKT---------------RS
Ecoracana-ELECO.r07.2AG0138010.1                   ARAAVENG-TPLVPNRIKECRSYPLYRFVREEVGTEYLTGEKT---------------RS
Ecoracana-ELECO.r07.2BG0193660.1                   ARAAVENG-TPLVPNRIKECRSYPLYRFVREEVGTEYLTGEKT---------------RS
Osativa-LOC_Os02g41650.3                           ARAAVENG-TPLVPNRIKECRSYPLYRFVREEVGTEYLTGEKT---------------RS
Pvaginatum-Pavag04G202800.1                        ARAAVESG-NPLVPNRIKECRSYPLYRFVREEVGTEYLTGEKA---------------RS
Pvaginatum-PavagK320900.1                          ARAAVESG-NPLVPNRIKECRSYPLYRFVREEVGTEYLTGEKA---------------RS
Pvirgatum-Pavir.1NG356400.1                        ARAAVESG-NPMVPNRIKECRSYPLYRFVREELGTEYLTGEKT---------------RS
Sviridis-Sevir.1G245100.1                          ARAAVESG-RPMVPNRIKECRSYPLYRFVREELGAEYLTGEKT---------------RS
Sbicolor-Sobic.004G220400.1                        ARAAVESG-NPLVPNRIKECRSYPLYRFVREEVGTQYLTGEKT---------------RS
Zmays-Zm00001d017275_T001                          ARAAVESG-SPLVPNRIRECRSYPLYRFVREDVGTEYLTGERT---------------RS
Zmays-Zm00001d051163_T002                          ARAAVENG-SPLVPNRIKECRSYPLYRFVREEVGTKYLTGEKT---------------RS
Pvirgatum-Pavir.1KG386300.1                        ARVAVAEG-AAPLPNRIKDSRSFPLYRFVREELGCVFLTGEKL---------------KS
Platifolius-Phala.02G210600.1                      ARNAVENG-TAFVPNRIKECRSYPLYRFVREELGTEYLTGEKT---------------RS
Streptochaeta-strangu_020767-RA                    ARAAVENG-AATVPNRIKECRSYPLYRFAREELGTDYLTGEKA---------------RS
Bdistachyon-Bradi5g15830.1                         ARAAVENG-TATKQNRIAECRSYPLYRFVREELGTEYLTGEKT---------------RS
Hvulgare-HORVU2Hr1G089440.4                        ARSAVENG-TAAQQNRISECRSYPLYRFVRKELGTEYLTGEKT---------------RS
Taestivum-Traes_2AL_0DDC60D6E.1                    ARSAVENG-TAAQQNRIAECRSYPLYRFVRKELGTEYLTGEKT---------------RS
Tintermedium-Thint.06G0475900.1                    ARSAVENG-TAAQQNRIAECRSYPLYRFVRKELGTKYLTGEKT---------------RS
Tintermedium-Thint.06G0476200.1                    ARSAVENG-TAAQQNRIAECRSYPLYRFVRKELGTEYLTGEKT---------------RS
Taestivum-Traes_2AL_2763280FC.1                    ARSAVENG-TAAQQNRIAECRSYPLYRFVRKELGTEYLTGEKT---------------RS
Taestivum-Traes_2DL_8C21AAAEF.1                    ARSAVENG-TAAQQNRIAECRSYPLYRFVRKELGTEYLTGEKT---------------RS
Tintermedium-Thint.V0383500.1                      ARSAVENG-TAAQQNRIAECRSYPLYRFVRKELGTEYLTGEKT---------------RS
Taestivum-Traes_2AL_9EC3226F7.1                    ARSAVENG-TAAQQNRIAECRSYPLYRFVRKELGTEYLTGEKT---------------RS
Tintermedium-Thint.05G0273800.1                    ARSAVENG-TAAQQNRIAECRSYPLYRFVRKELGTEYLTGEKT---------------RS
Tintermedium-Thint.V0383400.1                      ARSAVENG-TAEQQNRITECRSYPLYRFVRKELGTEYLTGEKT---------------RS
Taestivum-Traes_2BL_13D5272D7.1                    ARSAVENG-TAAQQNRIADCRSYPLYRFVRKELGTEYLTGEKT---------------RS
Tintermedium-Thint.06G0475700.1                    ARSAVENG-TAAQQNRIADCRSYPLYRFVRKELGTEYLTGEKT---------------RS
Taestivum-Traes_2BL_C051606EA.2                    ARTAVENG-TAAQQNRIAECRSYPLYRFVRKELGTEYLTGEKT---------------RS
Tintermedium-Thint.V0383600.1                      ARSAVENG-TATQQNRITECRSYPLYRFVRKELGTEYLTGEKT---------------TS
Taestivum-Traes_2AL_9D78F85E2.1                    ARSAVENG-TATQQNRIAECRSYPLYRFVRKELGTVYLTGEKT---------------RS
Tintermedium-Thint.05G0274100.1                    ARSAVENG-TATQQNRITECRSYPLYRLVRKELGTEYLTGEKT---------------RS
Tintermedium-Thint.06G0476300.1                    ARSAVENG-TATQQNRITECRSYPLYRFVRKELGTEYLTGEKT---------------RS
Taestivum-Traes_2BL_5A50FDA1A.1                    ARSAVENG-TATQQNRITECRSYPLYRFVRKELGTEYLTGEKT---------------RS
Tintermedium-Thint.05G0273900.1                    ARSAVENG-TAAQQNRIAECRSYPLYRFVRKELGTEYLTGEKT---------------RS
Taestivum-Traes_2DL_30F8555D4.1                    ARSAVENG-TATQQNRIADCRSYPLYRFVRKELGTEYLTGEKT---------------RS
Taestivum-Traes_2DL_83168C1E0.1                    ARGAVENG-TATQQNRIAECRSYPLYRFVRKELGTEYLTGEKT---------------RS
Taestivum-Traes_2DL_B978B4E22.1                    ARSAVENG-TAAQQNRIAECRSYPLYRFVRKELGTEYLTGEKT---------------RS
Taestivum-Traes_1AS_0B9295A68.1                    ARTSVENG-TATQQNRITECRSYPLYRFVRKELGAEYLTGEKT---------------RC
Taestivum-Traes_1BS_7068F56E6.1                    ARISVENG-TAALQNRITECRSYPLYRLVRKELGTEYLTGEKT---------------RS
Taestivum-Traes_1BS_AEA817616.1                    AWTSVENG-TAAQQNRITECRSYPLYRFVRKELGTEYLTGEKT---------------RS
Taestivum-Traes_4AL_892C47ED5.1                    ARTSVENG-TATQQNRITECRSYPLYRFVRKELGTEYLTGEKT---------------RS
Tintermedium-Thint.03G0055900.1                    ARTSVENG-IAAQKNRITECRSYPLYRFMRKELGTEFLTGEKT---------------RS
Tintermedium-Thint.01G0012900.1                    ARTSVENG-TATQQNRINECRSYPLYRFVRKELGTEYLTGEKT---------------RS
Tintermedium-Thint.02G0158500.1                    ARTSVENG-TAAQQNRITECRSYPLYRFVRKELGTEYLTGEKT---------------RS
Tintermedium-Thint.03G0055700.1                    ARTSVENG-TAAQQNRITECRSYPLYRFVRKELGTEYLTGEKT---------------RS
Taestivum-Traes_1BS_C3AA04888.1                    ARTSVENG-TAAQQNRITECRSYPLYRFVRKELGTEYLTGEKT---------------RS
Taestivum-Traes_1DS_A8BD91E4A.1                    ARSSVENG-TATQHNRITGCRSYPLYRFVRKELGTEYLTGEKT---------------RS
Taestivum-Traes_1BS_C26CCE0E9.1                    ARTLVENG-TAAQQNRISECRSYPLYRFVRKELGTEYLTGEKT---------------RS
Taestivum-Traes_1DS_A21FCBA3D.1                    ARTSVENG-TAAQQNRITECRSYPLYRFVRKELGTEYLTGEMT---------------RS
Tintermedium-Thint.01G0012200.1                    ARTSVENG-TAAQQNRITECRSYPLYRFVRKELGTEYLTGEKT---------------RS
Tintermedium-Thint.02G0158600.1                    ARTSVENG-TAVQQNRITECRSYPLYRLVRKELGTEYLTGEKT---------------RS
Tintermedium-Thint.01G0012700.1                    ARASVENG-TAAQQNRITECRSYPLYRLVRKELGTEYLTGEKT---------------RS
Taestivum-Traes_5BL_F50A47503.1                    ARTSVENG-TAAQQNRITECRSYPLYRLVRKELGTEYLTGEKT---------------RS
Tintermedium-Thint.02G0158700.1                    ARTSMENG-TAAQQNRITECRSYPLYRLVRKELGTEYLTGEKT---------------RS
Taestivum-Traes_5BL_AB528184A.1                    ARTSVENG-TAAQQNRITGCRSYPLYRLVRKELGTEYLTGEKT---------------RS
Tintermedium-Thint.03G0055800.1                    ARTSVENG-TAVQKNRITECRSYPLYRFVRKELGTEYLTGEKT---------------RS
Taestivum-Traes_1AS_F9013A945.1                    ARSAVENG-IAAQQNRITGCRSYPLYRFVRNELGTEYLTGEKT---------------RS
Taestivum-Traes_1DS_43DAC7A7F.1                    ARSAVENG-TAAQQNRITECRSYPLYRFVRKELGTEYLTGEKT---------------RS
Taestivum-Traes_1BS_BD86C90A7.1                    ARSAVENG-TAEQQNRITECRSYPLYRFVRKELGTEYLTGEKT---------------RS
Tintermedium-Thint.02G0158900.1                    ARSAVENG-TAAQQNRITECRSYPLYQFVREELGTVYLTGEKA---------------RS
Tintermedium-Thint.05G0274000.1                    ARSAVENG-TATQQNRITECRSYPLYRFVRKELGTEYLTGEKT---------------TS
Osativa-LOC_Os04g43800.1                           ARAAVENG-TAAKANRITECRSYPLYRFVREELGTEYLTGEKT---------------RS
Pvaginatum-Pavag06G160300.1                        ARAAVENG-TAEVPNRIAECRSYPLYRFVREELGTQYLTGEKT---------------RS
Sbicolor-Sobic.006G148900.1                        ARAAVENG-TAAVPNRIAECRSYPLYRFVREELGTEYLTGEKT---------------RS
Pvirgatum-Pavir.7KG237800.1                        ARAAVENG-TAAIPNRIAECRSYPLYRFVRQELGTEYLTGEKT---------------RS
```

```
Pvirgatum-Pavir.7NG355800.1                        SRAGVEKG-AAAIPNRIAECRSYPLYRFVRQELGTEYLTGEKT---------------RS
Sviridis-Sevir.7G178300.1                          ARAAVENG-TAAIPNRIAECRSYPLYRFVREELGTQYLTGEKT---------------RS
Zmays-Zm00001d003015_T001                          ARAAVESG-TAAIRNRIAECRSYPLYRFVREELGTEYLTGEKA---------------RS
Hvulgare-HORVU1Hr1G022060.1                        ARGAVENG-TATEPNRIADCRSYPLYRFVRKELGTVYLTGEKT---------------RS
Taestivum-Traes_1BS_723922D171.2                   ARGAVENG-TAAEPNRIADCRSYPLYRFVRKELGTVYLTGEKT---------------RS
Taestivum-Traes_1DS_A171C7D59.1                    ARGAVENG-TAAEPNRIADCRSYPLYRFVRKELGTVYLTGEKT---------------RS
Tintermedium-Thint.U1G0111500.1                    ARGAVENG-TAAEPNRIADCRSYPLYRFVRKELGTVYLTGEKT---------------RS
Tintermedium-Thint.03G0265300.1                    ARGAVENG-TATEPNRIADCRSYPLYHFVRKELGTVYLTGEKT---------------RS
Hvulgare-HORVU2Hr1G038140.1                        ARGAVENG-TAAEPNRIVDCRSYPLYRFVREELGTVYLTGEKT---------------RS
Tintermedium-Thint.04G0171200.1                    ARGAVENG-TAAEPNRITDCRSYPLYRFVREELGTVYLTGEKT---------------RS
Taestivum-Traes_2AS_958327519.3                    ARGAVENG-TTAEPNRIVDCRSYPLYRFVRKELGTVYLTGEKT---------------RS
Taestivum-Traes_2DS_28CA50371.1                    ARGAVENG-TAAEPNRIVDCRSYPLYRFVRKELGTVYLTGEKT---------------RS
Tintermedium-Thint.U5G0160800.1                    ARGAVENG-TAAEPNRIVDCRSYPLYRFVRKELGTVYLTGEKT---------------RS
Tintermedium-Thint.06G0299000.1                    ARGAVENG-TAAEPNRIVDCRSYPLYRFVRKELGTVYLTGEKT---------------RS
Taestivum-Traes_2DS_3791A8A36.1                    ARGAVENG-TAAEPNRITDCRSYPLYRFVREELGTVYLTGEKT---------------RS
Tintermedium-Thint.U6G0342800.1                    ARGAVENG-TAAEPNRITDCRSYPLYRFVREELGTVYLTGEKT---------------RS
Tintermedium-Thint.05G0150900.1                    ARGAVENG-TAAEPNRIVDCRSYPLYRFVRKELGTVYLTGEKT---------------RS
Tintermedium-Thint.04G0242300.1                    ARGAVENG-TAEEPNRIVDCRSYPLYRFVREELGTVYLTGEKT---------------RS
Tintermedium-Thint.06G0298700.1                    ARGAVENG-TAAEPNRIVDCRSYPLYRFVREELGTVYLTGEKT---------------RS
Taestivum-Traes_2BS_88CF42F2E.1                    ARGAVENG-TAAEPNRIVDCRSYPLYRFVREELGTVYLTGEKT---------------RS
Tintermedium-Thint.U4G0241900.1                    ARGAVENG-TAAEPNRIVDCRSYPLYRFVRKELGTVYLTGEKT---------------RS
Platifolius-Phala.06G165300.1                      ARAAVENG-AAAKPNRITECRSYPLYRFVREELGTQYLTGEKT---------------RS
Streptochaeta-strangu_019385-RA                    ARAAVENG-AAAKPNRITECRSYPLYRFVREELGAKYLTGEKT---------------RS
Streptochaeta-strangu_028944-RA                    ARVAVENG-GAPTPNRVKECRSYPLYRFVRDEAGAEYLTGEKA---------------RS
Acomosus-Aco010091.1                               AWVAFGSG-KSAIQNRIKECRSYPLYKFVKEELGTECLTGEKV---------------RS
Zmarina-Zosma03g22300.1                            ARLAHENG-NSVVGNRIKDCRSYPLYRFVTEELGVGLLTGEKI---------------RS
Ecdeiocolea-Emoptg0003741_1G000600.1               AWAAVENG-TAATPNRIKECRSYPLYRFVREELGSGYLTGEKV---------------RS
Jascendens-Joasc.05G060500.1                       ARVAVENG-TAPTPNRIKECRSYPLYRFVREELGTEYLTGEKL---------------RS
Ecdeiocolea-Emoptg0003741_1G000630.1               VRAAVENG-NAPIPNRVKECRSYPLYRFLREELGTVYLTGEKL---------------RS
Jascendens-Joasc.05G060400.1                       ARVAVENR-TAPTQNRIKESRSYPLYRFVREDLGTAYLTGEKL---------------RS
Bdistachyon-Bradi3g49250.2                         ARVAFEKG-TAPIPNLIKDSRSFPLYRFVREELGCVYLTGEKL---------------LS
Hvulgare-HORVU6Hr1G058820.1                        ARVAFENG-TAPIPNLIKESRSFPLYRFVREELGCVYLTGEKL---------------LS
Taestivum-Traes_6DL_1AEA7B869.1                    ARVAFENG-TAPIPNLIKESRSFPLYRFVREELGCVYLTGEKL---------------LS
Tintermedium-Thint.I8G0192300.1                    ARVAFENG-TAPIPNLIKESRSFPLYRFVREELGCVYLTGEKL---------------LS
Tintermedium-Thint.17G0230800.1                    ARVAFENG-TAPIPNLIKESRSFPLYRFVREELGCVYLTGEKL---------------LS
Tintermedium-Thint.17G0230900.1                    ARVAFENG-TAPIPNLIKESRSFPLYRFVREELGCVYLTGEKL---------------LS
Ecoracana-ELECO.r07.2AG0138000.1                   ARVAVAEG-TAPVANRIKDSRSYPLYRFVREELGCVFLTGEKL---------------KS
Ecoracana-ELECO.r07.2BG0193650.1                   ARVAVAEG-TAPVANRIKDSRSYPLYRFVREELGCVFLTGEKL---------------KS
Pvaginatum-Pavag04G202700.1                        ARVAVAEG-AAAVDNRIKDSRSFPLYRFVREELGCVFLTGEKL---------------KS
Sviridis-Sevir.1G245000.1                          ARVAVAEG-TAAVENRIKDSRSFPLYRFVREELGCVFLTGEKL---------------KS
Sviridis-Sevir.6G187100.1                          ARVAVAEG-TAAVENRIKDSRSFPLYRFVREELGCVFLTGEKL---------------KS
Pvirgatum-Pavir.1NG356200.1                        ARVAVAEG-AAAVPNRIKDSRSFPLYRFVREELGCVFLTGEKL---------------KS
Sbicolor-Sobic.004G220300.1                        ARVAVAEG-TAPVANRIADSRSFPLYRFVREELGCVFLTGEKL---------------KS
Zmays-Zm00001d017274_T001                          ARVAVAEG-TAPVANRIADSRSFPLYRFVREELGCVFLTGERL---------------KS
Zmays-Zm00001d051161_T003                          ARVAVAEG-TAPVANRIADSRSFPLYRFVREELGCVFLTGEKL---------------KS
Sviridis-Sevir.7G178200.1                          ARVAVAEG-TAPVANRIKDSRSFPVYRFVREELGCVFLTGEKL---------------KS
Osativa-LOC_Os02g41630.2                           ARVAVANG-TAPVANRIVESRSFPLYRFVREELGCVFLTGEKL---------------KS
Ecoracana-ELECO.r07.4AG0310430.1                   ARVAFENG-TAPIGNRIQESRSYPLYRFIRQDLGAVYLTGEKL---------------KS
Ecoracana-ELECO.r07.4BG0341620.1                   ARVAFENG-TAPIGNRIQESRSYPLYRFIRQDLGAVYLTGEKL---------------KS
Pvaginatum-Pavag06G160200.1                        ARVAFETG-TAPIANRIKESRSYPLYRFIRQDLGAVFLTGEKL---------------KS
Pvirgatum-Pavir.7KG238255.1                        ARVAFETG-AAPIANRIEESRSYPLYRFIRQDLGAVYLTGEKL---------------KS
Sbicolor-Sobic.006G148800.1                        ARVAFETG-TAPIGNRIKDSRSYPLYRFIRQDLGAVYLTGEKL---------------KS
Zmays-Zm00001d003016_T001                          ARVAFETG-AAPIANRIKESRSYPLYRFIRQDLGAVYLTGEKL---------------KS
Osativa-LOC_Os04g43760.1                           ARVAFETG-TAPITNRIKESRSFPLYRFVREELGCVYLTGEKL---------------KS
Platifolius-Phala.02G210400.1                      ARVAVEKG-AAPTANRIKESRSYPLYRFIREDLGAVYLTGEKL---------------RS
Platifolius-Phala.06G165100.1                      SRLAVENG-AAPTANRIKESRSYPLYRFIREDLGTVYLTGEKL---------------RS
Streptochaeta-strangu_019386-RA                    ARVAVENG-TAPTANRIKESRSYPLYRFIREDLGAVYLTGEKL---------------RS
Streptochaeta-strangu_020769-RA                    ARVAFENG-ASQTPNRMKDSRSYPLYRFIREDLGAVYLTGEKL---------------RS
Sviridis-Sevir.2G448300.1                          TRVAFETG-TAPIGNRIKESRSYPLYRFIREDLGAVYLTGEKL---------------KS
Sviridis-Sevir.7G177900.1                          TRVAFETG-TAPIGNRIKESRSYPLYRFIREDLGAVYLTGEKL---------------KS
                                                                      .  :                                    .


Aamericanus-Acora.01G232100.1                      PGE-EFDKVFVAISEGRVIDPLLECLS------------------------DWNG-----
Aamericanus-Acora.10G174800.1                      PGE-EFDKVFVAISQGKVIDPLLECLS------------------------GWNG-----
Acomosus-Aco013943.1                               PGE-EFDKVFVAINEGKLIDPLLECLK------------------------EWNG-----
Aofficinalis-evm.model.AsparagusV1_01.827          PGE-EFDKVFDAICDGRFIDPMLECLS------------------------EWNG-----
Typha_latifolia-Tylat.14G083800.1                  PGE-EFDKVFNAISQGKVIDPMLECIK------------------------EWNG-----
Aofficinalis-evm.model.AsparagusV1_04.71           PGE-EFDKVFVAISQGKMIDAVLACLK------------------------DWNG-----
Aofficinalis-evm.model.AsparagusV1_04.93           PGE-EFDKVFVAISQGKMIDAVLACLK------------------------DWNG-----
Aofficinalis-evm.model.AsparagusV1_10.1948         PGE-EFDKVFVAISEGKVIDPMLECLR------------------------DWNG-----
Typha_latifolia-Tylat.03G127700.1                  PGE-ECDKVFMAINRGKVIDPLIECLK------------------------EWNG-----
Spolyrhiza-Spipo11G0025500                         PGE-EFDKVFTAISQGKVIDPLFNCLS------------------------GWNG-----
Spolyrhiza-Spipo1G0003500                          PGE-EFDKVYVAISQGKVVDPLFECLR------------------------DWNG-----
Phoenix_dactylifera-LOC103718264                   PGE-EFGKVFNGICEGKVIDPLLDCLK------------------------DWNG-----
Macuminata-GSMUA_Achr1T09070_001                   PGE-EFDKVFDAICQGKVIDPLLECLK------------------------EWDG-----
Acomosus-Aco006987.1                               PGE-EFNKVFNAICKGKAIDPMLECLK------------------------EWNG-----
Acomosus-Aco027752.1                               PGE-EFNKVFNAICKGKAIDPMLECLK------------------------EWNG-----
Cscoparia-Csco27g03980.1                           PGE-ECNKVFNAICDGKVVDPLLECLK------------------------EWNG-----
Cscoparia-Csco27g03990.1                           PGE-ECNKVFNAICDGKVIDPLLECLK------------------------EWNG-----
Macuminata-GSMUA_Achr11T16380_001                  PGE-EFDKVFVAIDRGLVIDPLLECLK------------------------EWNG-----
Macuminata-GSMUA_Achr11T22840_001                  PGE-EFDKVFAAINKGLLIDPLMECLK------------------------EWNG-----
Macuminata-GSMUA_Achr2T00240_001                   PGE-EFDKVFVAINGGSLIDPLLECLK------------------------EWNG-----
Macuminata-GSMUA_Achr5T18560_001                   PGE-EFDKVFMAVNEGLLIDPLLECLK------------------------EWNG-----
Macuminata-GSMUA_Achr8T18960_001                   PGE-EFDKVFAAINRGQLIDPLLECLK------------------------EWNG-----
Macuminata-GSMUA_Achr5T03950_001                   PGE-DFDKIFAAINKGMVIDPLLECLK------------------------EWNG-----
Phoenix_dactylifera-LOC103703950                   PGE-EFDKVFVAISRGMVIDPLLECLK------------------------GWDG-----
Acoerulea-Aqcoe1G140100.1                          PGE-VFDKVFTAMCNGKMIDPLLECLS------------------------CWNG-----
Acoerulea-Aqcoe2G144500.1                          PGE-EFDQVFSAICQGKMIDPLLDCLK------------------------EWNG-----
Typha_latifolia-Tylat.05G053500.1                  PGE-EFDKVFAAICEGKVFAPMLECLK------------------------EWNG-----
Zmarina-Zosma05g29470.1                            PGE-EFDKVFVAINEGELVGPLMECLK------------------------EWNG-----
Acomosus-Aco007727.1                               PGE-ECDKVFAAINGGLLIDPLLECLK------------------------DWDG-----
Bvulgarisssp-Bevul.7G054300.1                      PGE-EIDKVFTAICQGKIIDPLLDCLK------------------------EWNG-----
Bdistachyon-Bradi3g47110.1                         PGE-ELNKVLVAINQGKHIDPLLECLK------------------------EWNG-----
Bdistachyon-Bradi3g47120.1                         PGE-ELNKVLVAINQGKHIDPLLECLK------------------------EWNG-----
Bdistachyon-Bradi3g48840.1                         PGE-ELNKVLVAINQGKHIDPLLECLK------------------------EWNG-----
Bdistachyon-Bradi3g49280.1                         PGE-ELNKVLVAINQGKHIDPLLECLK------------------------EWNG-----
Hvulgare-HORVU6Hr1G058840.1                        PGE-ELNKVLVAINQGKHIDALLECLK------------------------EWNG-----
Taestivum-Traes_6BL_C96F626D9.1                    PGE-ELNKVLVAMNQGKHIDALLECLK------------------------EWNG-----
Tintermedium-Thint.I8G0192700.1                    PGE-ELNKVLVAMNQGKHIDALLECLK------------------------EWNG-----
Taestivum-Traes_6DL_04CD7E82C.1                    PGE-ELNKVFVAMNQGKHIDALLECLK------------------------EWNG-----
Bdistachyon-Bradi3g49270.1                         PGE-ELNKVLMAINQGKHIDPLLECLK------------------------EWNG-----
Ecoracana-ELECO.r07.2AG0138020.1                   PGE-ELNKVLVAINQDKHIDPLLECLK------------------------EWNG-----
Ecoracana-ELECO.r07.2BG0193670.1                   PGE-ELNKVLVAMNEGKHIDPLLECLK------------------------EWNG-----
Osativa-LOC_Os02g41670.1                           PGE-ELNKVLVAINQGKHIDPLLECLK------------------------EWNG-----
Osativa-LOC_Os02g41680.1                           PGE-ELNKVLVAINEGKHIDPLLECLK------------------------EWNG-----
Pvaginatum-Pavag04G202900.1                        PGE-ELNKVLLAINQGKHIDPLLECLK------------------------EWNG-----
Pvaginatum-PavagK321000.1                          PGE-ELNKVLLAINQGKHIDPLLECLK------------------------EWNG-----
Pvaginatum-PavagK321300.1                          PGE-ELNKVLLAINQGKHIDPLLECLK------------------------EWNG-----
Pvaginatum-PavagK321100.1                          PGE-ELNKVLLAINQGKHIDPLLECLK------------------------EWNG-----
Pvaginatum-PavagK321200.1                          PGE-ELNKVLLAINQGKHIDPLLECLK------------------------EWNG-----
Pvirgatum-Pavir.1KG386500.1                        PGE-ELNKVLVAINQGKHIDPLLECLK------------------------EWNG-----
Pvirgatum-Pavir.1NG356700.1                        PGE-ELNKVLVAINQGKHIDPLLECLK------------------------EWNG-----
Sviridis-Sevir.1G245166.1                          PGE-ELNKVLVAINQGKHIDPLLECLK------------------------EWNG-----
Sviridis-Sevir.1G245232.1                          PGE-ELNKVLVAINQGKHIDPLLECLK------------------------EWNG-----
Pvirgatum-Pavir.1NG356800.1                        PGE-ELNKVLVAINLGKHIDPLLECLK------------------------EWNG-----
Sbicolor-Sobic.004G220500.1                        PGE-ELNKVLVAINQGKHIDPLLECLK------------------------EWNG-----
Sbicolor-Sobic.004G220600.2                        PGE-ELNKVLVAINQGKHIDPLLECLK------------------------EWNG-----
Sbicolor-Sobic.004G220700.1                        PGE-ELNKVLVAINQGKHIDPLLECLK------------------------EWNG-----
Sviridis-Sevir.1G245300.2                          PGE-ELNKVLVAINQGKHIDPLLECLK------------------------EWNG-----
Zmays-Zm00001d017276_T001                          PGE-ELNKVLVAINQGKHIDPLLECLK------------------------EWNG-----
Zmays-Zm00001d017279_T001                          PGE-ELNKVLVAINQGKHIDPLLECLK------------------------EWNG-----
Zmays-Zm00001d051166_T001                          PGE-ELNKVLVAINQGKHIDPLLECLK------------------------EWNG-----
Ecoracana-ELECO.r07.2AG0138030.1                   PGE-ELNKVLVAINEGKHIDPLLECLK------------------------EWNG-----
Ecoracana-ELECO.r07.2BG0193680.1                   PGE-ELNKVLVAMNEGKHIDPLLECLN------------------------EWNG-----
Osativa-LOC_Os05g35290.1                           PGE-ELNKVLVAINQGKHIDPLLECLK------------------------EWNG-----
Platifolius-Phala.02G210700.1                      PGE-EVNKVFVAMNQGKHIDALLECLK------------------------EWNG-----
Ecoracana-ELECO.r07.4AG0310420.1                   PGE-ELNKVLVAINQGKHIDPLLECLK------------------------EWNG-----
Ecoracana-ELECO.r07.4BG0341610.1                   PGE-ELNKVLVAINQGKHIDPLLECLK------------------------EWDG-----
```

```
Bdistachyon-Bradi3g49260.1                PGE-ELNKVLVAMNQRKHIDPLLECLK-----------------------EWNG-----
Hvulgare-HORVU0Hr1G016330.1               PGE-ELNKVLVAMNERKHIDPLLECLK-----------------------EWNG-----
Tintermedium-Thint.17G0260200.1           PGE-ELNKVLVAMNERKHIDPLLKCLK-----------------------EWNG-----
Tintermedium-Thint.18G0192600.1           PGE-ELNKVLVAMNERKHIDPLLECLK-----------------------EWNG-----
Tintermedium-Thint.17G0231000.1           PGE-ELNKVLVAMNERKHIDPLLECLK-----------------------EWNG-----
Ecoracana-ELECO.r07.2AG0138010.1          PGE-ELNKVLVAMNERKHIDPLLECLK-----------------------EWNG-----
Ecoracana-ELECO.r07.2BG0193660.1          PGE-ELNKVLVAMNERKHIDPLLECLK-----------------------EWNG-----
Osativa-LOC_Os02g41650.3                  PGE-ELNKVLVAINERKHIDPLLECLK-----------------------EWNG-----
Pvaginatum-Pavag04G202800.1               PGE-ELNKVLVAINQGKHIDPLLECLK-----------------------EWNG-----
Pvaginatum-PavagK320900.1                 PGE-ELNKVLVAINQGKHIDPLLECLK-----------------------EWNG-----
Pvirgatum-Pavir.1NG356400.1               PGE-ELNKVLVAINQRKHIDPLLECLK-----------------------EWNG-----
Sviridis-Sevir.1G245100.1                 PGE-ELNKVLVAINERKHIDPLLECLK-----------------------EWNG-----
Sbicolor-Sobic.004G220400.1               PGE-ELNKVLVAINQRKHVDPLLECLK-----------------------EWNG-----
Zmays-Zm00001d017275_T001                 PGE-ELNKVLVAINQRKHIDPLLECLK-----------------------EWNG-----
Zmays-Zm00001d051163_T002                 PGE-ELNKVLVAINQRKHIDPLLECLK-----------------------EWNG-----
Pvirgatum-Pavir.1KG386300.1               PGE-ECNKVFVGISEGKLVDPMLECLK-----------------------EWDG-----
Platifolius-Phala.02G210600.1             PGE-EVNKVLVAMNQSKHIDPLLECLK-----------------------EWNG-----
Streptochaeta-strangu_020767-RA           PGE-EVDKVLVAINQGKHIDPLLECLK-----------------------EWNG-----
Bdistachyon-Bradi5g15830.1                PGE-EVDKVFVAMNQGKHIDALLECLK-----------------------EWNG-----
Hvulgare-HORVU2Hr1G089440.4               PGE-EVDKVFVAMNQGKHIDALLECLK-----------------------EWNG-----
Taestivum-Traes_2AL_0DDC60D6E.1           PGE-EVDKVFIAMNQGKHIDALLECLK-----------------------EWNG-----
Tintermedium-Thint.06G0475900.1           PGE-EVDKVFIAMNQGKHIDALLECLK-----------------------EWNG-----
Tintermedium-Thint.06G0476200.1           PGE-EVDKVFIAMNQGKHIDALLECLK-----------------------EWNG-----
Taestivum-Traes_2AL_2763280FC.1           PGE-EVDKVF-------------------------------------------------
Taestivum-Traes_2DL_8C21AAAEF.1           PGE-E------------------------------------------------------
Tintermedium-Thint.V0383500.1             PGE-EVDKVFIAMNQGKHIDALLECLK-----------------------EWNG-----
Taestivum-Traes_2AL_9EC3226F7.1           PGE-EVDKVFVAMNQGKHIDALLECLK-----------------------EWNG-----
Tintermedium-Thint.05G0273800.1           PGE-EVDKVFVAMNQGKHIDALLECLK-----------------------EWNG-----
Tintermedium-Thint.V0383400.1             PGE-EVDKVFVAMNHGKHIDALLECLK-----------------------EWNG-----
Taestivum-Traes_2BL_13D5272D7.1           PGE-EVDKVFVAMNQGKHIDALLECLK-----------------------EWNG-----
Tintermedium-Thint.06G0475700.1           PGE-EVDKVFVAMNQGKHIDALLECLK-----------------------EWNG-----
Taestivum-Traes_2BL_C051606EA.2           PGE-EVDKVFIAMNQGKHIDALLECLK-----------------------EWNG-----
Tintermedium-Thint.V0383600.1             PGE-EVDKVFVAMNQGKHIGALLECLK-----------------------RWNG-----
Taestivum-Traes_2AL_9D78F85E2.1           PGE-EVDKVFVAMNQGKHIDALLECLK-----------------------GWNG-----
Tintermedium-Thint.05G0274100.1           PGE-EVDKVFIAMNQGKHIDALLECLK-----------------------GWNG-----
Tintermedium-Thint.06G0476300.1           PGE-EVDKVFVAMNQGKHIDALLECLK-----------------------GWNG-----
Taestivum-Traes_2BL_5A50FDA1A.1           PGE-EVDKVFVAMNQGKHIDALLECLK-----------------------GWNG-----
Tintermedium-Thint.05G0273900.1           PGE-EVDKVFNAMNQGKHIDALLECLK-----------------------EWNG-----
Taestivum-Traes_2DL_30F8555D4.1           PGE-EVDKVFVAMNQGKHIDALLECLK-----------------------EWNG-----
Taestivum-Traes_2DL_83168C1E0.1           PGE-EVDKVFVAMNQGKHIDALLECLK-----------------------GWNG-----
Taestivum-Traes_2DL_B978B4E22.1           PGE-EVDKVFIAMNQGKHIDALLECLK-----------------------EWNG-----
Taestivum-Traes_1AS_0B9295A68.1           PGE-EVDKVFVAMNQGKHIDALLECLK-----------------------EWNG-----
Taestivum-Traes_1BS_7068F56E6.1           PGE-EVDKVFIAMNQGKHIDALLECLK-----------------------EWNG-----
Taestivum-Traes_1BS_AEA817616.1           PGE-EVDKVFVAMNQGKHIDALLECLK-----------------------EWNG-----
Taestivum-Traes_4AL_892C47ED5.1           PGE-EVDKVFVAMNQGKHIDALLECLK-----------------------EWNG-----
Tintermedium-Thint.03G0055900.1           PGE-EVDKVFVAMNQGKHIDALLECLK-----------------------EWNG-----
Tintermedium-Thint.01G0012900.1           PGE-EVDKVFVAMNQGKHIDALLECLK-----------------------EWNG-----
Tintermedium-Thint.02G0158500.1           PGE-EVDKVFVAMNQGKHIDALLECLK-----------------------EWKG-----
Tintermedium-Thint.03G0055700.1           PGE-EVDKVFVAMNQGKHIDALLECLK-----------------------EWNG-----
Taestivum-Traes_1BS_C3AA04888.1           PGE-EVDKVFVAMNQGKHIDALLECLK-----------------------EW-------
Taestivum-Traes_1DS_A8BD91E4A.1           PGE-EVDKVFVAMNQGKHIDALLECLE-----------------------EWNG-----
Taestivum-Traes_1BS_C26CCE0E9.1           PGE-EVDKVFVAMNQGKHIDALLECLK-----------------------EWNG-----
Taestivum-Traes_1DS_A21FCBA3D.1           PGE-EVDKVFVAMNQGKHIDALLECLK-----------------------EWNG-----
Tintermedium-Thint.01G0012200.1           PGE-EVDKVFVAMNQGKHIDALLECLK-----------------------EWNG-----
Tintermedium-Thint.02G0158600.1           PGE-EVDKVFVAMNQGKHIDALLECLK-----------------------EWNG-----
Tintermedium-Thint.01G0012700.1           PGE-EVDKVFVAMNQGKHIDALLECLK-----------------------EWNG-----
Taestivum-Traes_5BL_F50A47503.1           PGE-EVDKVFVAMNQGKHIDALLECLK-----------------------EWNG-----
Tintermedium-Thint.02G0158700.1           PGE-EVDKVFVAMNQGKHIDALLECLK-----------------------EWNG-----
Taestivum-Traes_5BL_AB528184A.1           PGE-EVDKVFAAMNQGKHIDALLECLK-----------------------EWNG-----
Tintermedium-Thint.03G0055800.1           PGE-EVDKVFVAMNQGKYIDALLECLK-----------------------EWNG-----
Taestivum-Traes_1AS_F9013A945.1           PGE-EVDKVFIAMNQGKHIDALLECLK-----------------------EWNG-----
Taestivum-Traes_1DS_43DAC7A7F.1           PGE-EVDKVFIAMNQGKHIDALLECLK-----------------------EWNG-----
Taestivum-Traes_1BS_BD86C90A7.1           PGE-EVDKVFIAMNQGKHIDALLECLK-----------------------EWNG-----
Tintermedium-Thint.02G0158900.1           PGE-EVDKVFIAMNQGKHIDALLECLK-----------------------EWNG-----
Tintermedium-Thint.05G0274000.1           PGE-VVNKVFVAMNQGKHIDALLECLM-----------------------GWNG-----
Osativa-LOC_Os04g43800.1                  PGE-EVNKVFVAMNQGKHIDALLECLK-----------------------EWNG-----
Pvaginatum-Pavag06G160300.1               PGE-EVDKVFVAMNQGRHIDAVLECLK-----------------------EWNG-----
Sbicolor-Sobic.006G148900.1               PGE-EVNKVFVAMNLGKHIDAVLECLK-----------------------EWNG-----
Pvirgatum-Pavir.7KG237800.1               PGE-EVNKVFVAMNAGKHIDAVLECLK-----------------------GWNG-----
Pvirgatum-Pavir.7NG355800.1               PGE-EVNKVFVAMNAGKHIDAVLECLK-----------------------EWNG-----
Sviridis-Sevir.7G178300.1                 PGE-EVDKVFVAMNLGKHIDAVLECLK-----------------------EWNG-----
Zmays-Zm00001d003015_T001                 PGE-EVDKVFVAMNLGKHIDAVLECLK-----------------------EWNG-----
Hvulgare-HORVU1Hr1G022060.1               PSE-EVDKVFIAMNQGKHIDALLECLK-----------------------EWNG-----
Taestivum-Traes_1BS_723922D171.2          PGE-EVDKVFIAMNQGKHIDALLECLK-----------------------EWNG-----
Taestivum-Traes_1DS_A171C7D59.1           PGE-EVDKVFIAMNQGKHINALLECLK-----------------------EWND-----
Tintermedium-Thint.01G0111500.1           PGE-EVDKVFIAISQGKHIDALLECLK-----------------------EWNG-----
Tintermedium-Thint.03G0265300.1           PGE-EVDKVFVAMNQGKHIDALLECLK-----------------------EWNG-----
Hvulgare-HORVU2Hr1G038140.1               PGE-EVDKVFVAMNQGKHIDALLECLK-----------------------EWNG-----
Tintermedium-Thint.04G0171200.1           PGE-EVDKVFVAMNQGKHIDALLECLK-----------------------EWNG-----
Taestivum-Traes_2AS_958327519.3           PGE-EVDKVFVAMNQGKHIDALLECLK-----------------------EWNG-----
Taestivum-Traes_2DS_28CA50371.1           PGE-EVDKVFVAMNQGKHIDALLECLK-----------------------EWNG-----
Tintermedium-Thint.05G0160800.1           PGE-EVDKVFVAMNQGKHIDALLECLK-----------------------EWNG-----
Tintermedium-Thint.06G0299000.1           PGE-EVDKVFVAMNQGKHIDALLECLK-----------------------EWNG-----
Taestivum-Traes_2DS_3791A8A36.1           PGE-EVDKVFIAMNQGKHIDALLECLK-----------------------EWNG-----
Tintermedium-Thint.06G0342800.1           PGE-EVDKVFIAMNQGKHIDALLECLK-----------------------EWNG-----
Tintermedium-Thint.05G0150900.1           PGE-EVDKVFVAMNQGKHIDALLECLK-----------------------EWNG-----
Tintermedium-Thint.04G0242300.1           PGE-EVDKVFVAMNQGKHIDALLECLK-----------------------EWNG-----
Tintermedium-Thint.06G0298700.1           PGE-EVDKVFVAMNQGKHIDALLECLK-----------------------EWNG-----
Taestivum-Traes_2BS_88CF42F2E.1           PGE-EVDKVFVAMNQGKHIDALLECLE-----------------------EWNG-----
Tintermedium-Thint.04G0241900.1           PGE-EVDKVFVGMNQGKHIDALLECLK-----------------------EWNG-----
Platifolius-Phala.06G165300.1             PGE-EVNKVFVAMNQGKLIDPLLECLK-----------------------EWNG-----
Streptochaeta-strangu_019385-RA           PGE-EVDKVFVAINQGKHIDPLLECLK-----------------------EWNG-----
Streptochaeta-strangu_028944-RA           PGE-EVDKVFVAMNLGKHIDPLLDCLK-----------------------EWNG-----
Acomosus-Aco010091.1                      PGE-EFDKVFDAICQARMIDPLLECLK-----------------------DWNG-----
Zmarina-Zosma03g22300.1                   PGE-EFDKVFAAINRGELIDPLFHCLA-----------------------DWNG-----
Ecdeiocolea-Emoptg0003741_1G000600.1      PGE-ECNRVFVAINKGKLIDPLLDCLN-----------------------EWNG-----
Jascendens-Joasc.05G060500.1              PGE-ECNKVFVAINQGKLIDPLLECLK-----------------------EWNG-----
Ecdeiocolea-Emoptg0003741_1G000630.1      PGE-ECNKVFVAINEGKLIDPLLECLK-----------------------EWNG-----
Jascendens-Joasc.05G060400.1              PGE-ECNKVFVAINQGKLIDPLLECLK-----------------------KWNG-----
Bdistachyon-Bradi3g49250.2                PGE-ECNKVFIGISQGKLIDPMLECLK-----------------------EWNG-----
Hvulgare-HORVU6Hr1G058820.1               PGE-ECTKVFLGISQNKLIDPMLECLK-----------------------EWNG-----
Taestivum-Traes_6DL_1AEA7B869.1           PGE-ECTKVFLGISQNKLIDPMLECLK-----------------------EWNG-----
Tintermedium-Thint.18G0192300.1           PGE-ECTKVFLGISQNKLIDPMLECLK-----------------------EWNG-----
Tintermedium-Thint.17G0230800.1           PGE-ECTKVFLGISQNKLIDPMLECLQ-----------------------EWNG-----
Tintermedium-Thint.17G0230900.1           PGE-ECTKVFLGISQNKLIDPMLECLK-----------------------EWNG-----
Ecoracana-ELECO.r07.2AG0138000.1          PGE-ECNKVFVGISQGKLVDPMLDCLK-----------------------EWDG-----
Ecoracana-ELECO.r07.2BG0193650.1          PGE-ECNKVFIGISQGKLVDPMLDCLK-----------------------EWDG-----
Pvaginatum-Pavag04G202700.1               PGE-ECNKVFIGISQGKLVDPMLECLK-----------------------EWDG-----
Sviridis-Sevir.1G245000.1                 PGE-ECNKVFIGISQGKLIDPMLECLK-----------------------EWDG-----
Sviridis-Sevir.6G187100.1                 PGE-ECNKVFVGISQGKLVDPMLECLK-----------------------EWDG-----
Pvirgatum-Pavir.1NG356200.1               PGE-ECNKVFIGISEGKLVDPMLECLK-----------------------EWDG-----
Sbicolor-Sobic.004G220300.1               PGE-ECTKVFNGINQGKLVDPMLECLK-----------------------EWDG-----
Zmays-Zm00001d017274_T001                 PGE-ECNKVFVGISQGKLVDPMLECLK-----------------------EWDG-----
Zmays-Zm00001d051161_T003                 PGE-ECNKVFLGISQGKLVDPMLECLK-----------------------EWDG-----
Sviridis-Sevir.7G178200.1                 PGE-ECNKVFIGISQGKLIDPMLECLK-----------------------EWDG-----
Osativa-LOC_Os02g41630.2                  PGE-ECNKVFLGISQGKLIDPMLDCLK-----------------------EWNG-----
Ecoracana-ELECO.r07.4AG0310430.1          PGE-ECNKVFLAINQGKIIDPMLDCLK-----------------------EWDG-----
Ecoracana-ELECO.r07.4BG0341620.1          PGE-ECNKVFLAINQDKIIDPMLDCLK-----------------------EWDG-----
Pvaginatum-Pavag06G160200.1               PGE-ECNKVFLALSEGKLIDPMLECLK-----------------------EWNG-----
Pvirgatum-Pavir.7KG238255.1               PGE-ECNKVFLALSEGKLIDPMLECLK-----------------------EWDG-----
Sbicolor-Sobic.006G148800.1               PGE-ECNKVFLALSEGKLIDPMLECLK-----------------------EWDG-----
Zmays-Zm00001d003016_T001                 PGE-ECNKVFLALSEGKLIDPMLDCLK-----------------------EWDG-----
Osativa-LOC_Os04g43760.1                  PGE-ECNKVFLAISERKLIDPMLECLK-----------------------EWNG-----
Platifolius-Phala.02G210400.1             PGE-ECNKVFLAISQGKLIDPLLECLK-----------------------EWNG-----
Platifolius-Phala.06G165100.1             PGE-ECNKVFVAISQGKLIDPLLECLK-----------------------EWNG-----
Streptochaeta-strangu_019386-RA           PGE-ECNKVFVAINQGKLVDPLLECLK-----------------------EWKG-----
Streptochaeta-strangu_020769-RA           PGE-ECNKVFVAISQGKLVDPLLDCLK-----------------------EWNG-----
Sviridis-Sevir.2G448300.1                 AGE-ECNKVFLALSEGKLIDPMLGCLK-----------------------EWNG-----
Sviridis-Sevir.7G177900.1                 AGE-ECNKVFLALSEGKLIDPMLGCLK-----------------------EWNG-----
```

```
Aamericanus-Acora.01G232100.1                        ---AP-------IPIC-----------------------
Aamericanus-Acora.10G174800.1                        ---AP-------IPIC-----------------------
Acomosus-Aco013943.1                                 ---EP-------LPIC-----------------------
Aofficinalis-evm.model.AsparagusV1_01.827            ---AP-------IPIC-----------------------
Typha_latifolia-Tylat.14G083800.1                    ---AP-------IPIC-----------------------
Aofficinalis-evm.model.AsparagusV1_04.71             ---AP-------LPIC-----------------------
Aofficinalis-evm.model.AsparagusV1_04.93             ---AP-------LPIC-----------------------
Aofficinalis-evm.model.AsparagusV1_10.1948           ---AP-------LPIC-----------------------
Typha_latifolia-Tylat.03G127700.1                    ---AP-------LPIC-----------------------
Spolyrhiza-Spipo11G0025500                           ---SP-------VPIC-----------------------
Spolyrhiza-Spipo1G0003500                            ---AP-------VPIC-----------------------
Phoenix_dactylifera-LOC103718264                     ---AP-------LPIC-----------------------
Macuminata-GSMUA_Achr1T09070_001                     ---AP-------IPIC-----------------------
Acomosus-Aco006987.1                                 ---AP-------LPLC-----------------------
Acomosus-Aco027752.1                                 ---AP-------LPLC-----------------------
Cscoparia-Csco27g03980.1                             ---EP-------LPIC-----------------------
Cscoparia-Csco27g03990.1                             ---EP-------LPIC-----------------------
Macuminata-GSMUA_Achr11T16380_001                    ---VP-------LPMC-----------------------
Macuminata-GSMUA_Achr11T22840_001                    ---AP-------LPIC-----------------------
Macuminata-GSMUA_Achr2T00240_001                     ---AP-------LPIC-----------------------
Macuminata-GSMUA_Achr5T18560_001                     ---AP-------LPIC-----------------------
Macuminata-GSMUA_Achr8T18960_001                     ---AP-------LPIC-----------------------
Macuminata-GSMUA_Achr5T03950_001                     ---AP-------LPIC-----------------------
Phoenix_dactylifera-LOC103703950                     ---AP-------LPLH-----------------------
Acoerulea-Aqcoe1G140100.1                            ---EP-------LPIC-----------------------
Acoerulea-Aqcoe2G144500.1                            ---AP-------LPIR-----------------------
Typha_latifolia-Tylat.05G053500.1                    ---AP-------LPIG-----------------------
Zmarina-Zosma05g29470.1                              ---AP-------IPIC-----------------------
Acomosus-Aco007727.1                                 ---AP-------LPIC-----------------------
Bvulgarisssp-Bevul.7G054300.1                        ---AP-------LPIC-----------------------
Bdistachyon-Bradi3g47110.1                           ---EP-------LPIC-----------------------
Bdistachyon-Bradi3g47120.1                           ---EP-------LPIC-----------------------
Bdistachyon-Bradi3g48840.1                           ---EP-------LPIC-----------------------
Bdistachyon-Bradi3g49280.1                           ---EP-------LPIC-----------------------
Hvulgare-HORVU6Hr1G058840.1                          ---EP-------LPIC-----------------------
Taestivum-Traes_6BL_C96F626D9.1                      ---KP-------LPIC-----------------------
Tintermedium-Thint.18G0192700.1                      ---EP-------LPIC-----------------------
Taestivum-Traes_6DL_04CD7E82C.1                      ---EP-------LPIC-----------------------
Bdistachyon-Bradi3g49270.1                           ---EP-------LPIC-----------------------
Ecoracana-ELECO.r07.2AG0138020.1                     ---EP-------LPIC-----------------------
Ecoracana-ELECO.r07.2BG0193670.1                     ---EP-------LPIC-----------------------
Osativa-LOC_Os02g41670.1                             ---AP-------LPIC-----------------------
Osativa-LOC_Os02g41680.1                             ---EP-------LPIC-----------------------
Pvaginatum-Pavag04G202900.1                          ---EP-------LPIC-----------------------
Pvaginatum-PavagK321000.1                            ---EP-------LPIC-----------------------
Pvaginatum-PavagK321300.1                            ---EP-------LPIC-----------------------
Pvaginatum-PavagK321100.1                            ---EP-------LPIC-----------------------
Pvaginatum-PavagK321200.1                            ---EP-------LPIC-----------------------
Pvirgatum-Pavir.1KG386500.1                          ---EP-------LPIC-----------------------
Pvirgatum-Pavir.1NG356700.1                          ---EP-------LPIC-----------------------
Sviridis-Sevir.1G245166.1                            ---EP-------LPIC-----------------------
Sviridis-Sevir.1G245232.1                            ---EP-------LPIC-----------------------
Pvirgatum-Pavir.1NG356800.1                          ---EP-------LPIC-----------------------
Sbicolor-Sobic.004G220500.1                          ---EP-------LPIC-----------------------
Sbicolor-Sobic.004G220600.2                          ---EP-------LPIC-----------------------
Sbicolor-Sobic.004G220700.1                          ---EP-------LPIC-----------------------
Sviridis-Sevir.1G245300.2                            ---EP-------LPIS-----------------------
Zmays-Zm00001d017276_T001                            ---EP-------LPIC-----------------------
Zmays-Zm00001d017279_T001                            ---EP-------LPIC-----------------------
Zmays-Zm00001d051166_T001                            ---EP-------LPIC-----------------------
Ecoracana-ELECO.r07.2AG0138030.1                     ---EP-------LPIC-----------------------
Ecoracana-ELECO.r07.2BG0193680.1                     ---EP-------LPIC-----------------------
Osativa-LOC_Os05g35290.1                             ---AP-------LPIC-----------------------
Platifolius-Phala.02G210700.1                        ---EP-------LPIC-----------------------
Ecoracana-ELECO.r07.4AG0310420.1                     ---EP-------LPIC-----------------------
Ecoracana-ELECO.r07.4BG0341610.1                     ---EP-------LPIC-----------------------
Bdistachyon-Bradi3g49260.1                           ---EP-------LPLC-----------------------
Hvulgare-HORVU0Hr1G016330.1                          ---AP-------LPLC-----------------------
Tintermedium-Thint.17G0260200.1                      ---EP-------LPLC-----------------------
Tintermedium-Thint.18G0192600.1                      ---EP-------LPLC-----------------------
Tintermedium-Thint.17G0231000.1                      ---EP-------LPLC-----------------------
Ecoracana-ELECO.r07.2AG0138010.1                     ---EP-------LPLC-----------------------
Ecoracana-ELECO.r07.2BG0193660.1                     ---EP-------LPLC-----------------------
Osativa-LOC_Os02g41650.3                             ---AP-------LPLC-----------------------
Pvaginatum-Pavag04G202800.1                          ---EP-------LPLC-----------------------
Pvaginatum-PavagK320900.1                            ---EP-------LPLC-----------------------
Pvirgatum-Pavir.1NG356400.1                          ---EP-------LPLC-----------------------
Sviridis-Sevir.1G245100.1                            ---EP-------LPLC-----------------------
Sbicolor-Sobic.004G220400.1                          ---EP-------LPLC-----------------------
Zmays-Zm00001d017275_T001                            ---AP-------LPLC-----------------------
Zmays-Zm00001d051163_T002                            ---EP-------LPLC-----------------------
Pvirgatum-Pavir.1KG386300.1                          ---KP-------LPIN-----------------------
Platifolius-Phala.02G210600.1                        ---KP-------LPIC-----------------------
Streptochaeta-strangu_020767-RA                      ---EP-------LPIC-----------------------
Bdistachyon-Bradi5g15830.1                           ---EP-------LPLC-----------------------
Hvulgare-HORVU2Hr1G089440.4                          ---EP-------LPIC-----------------------
Taestivum-Traes_2AL_0DDC60D6E.1                      ---EP-------LPLC-----------------------
Tintermedium-Thint.06G0475900.1                      ---EP-------LPLC-----------------------
Tintermedium-Thint.06G0476200.1                      ---EP-------LPLC-----------------------
Taestivum-Traes_2AL_2763280FC.1                      ---------------------------------------
Taestivum-Traes_2DL_8C21AAAEF.1                      ---------------------------------------
Tintermedium-Thint.V0383500.1                        ---EP-------LPIC-----------------------
Taestivum-Traes_2AL_9EC3226F7.1                      ---EP-------LPLC-----------------------
Tintermedium-Thint.05G0273800.1                      ---EP-------LPIC-----------------------
Tintermedium-Thint.V0383400.1                        ---EP-------LPIC-----------------------
Taestivum-Traes_2BL_13D5272D7.1                      ---EP-------LPIC-----------------------
Tintermedium-Thint.06G0475700.1                      ---EP-------LPIC-----------------------
Taestivum-Traes_2BL_C051606EA.2                      ---EP-------LPIC-----------------------
Tintermedium-Thint.V0383600.1                        ---EP-------LPIC-----------------------
Taestivum-Traes_2AL_9D78F85E2.1                      ---EP-------LPIC-----------------------
Tintermedium-Thint.05G0274100.1                      ---EP-------LPIC-----------------------
Tintermedium-Thint.06G0476300.1                      ---EP-------LPIC-----------------------
Taestivum-Traes_2BL_5A50FDA1A.1                      ---EP-------LPIC-----------------------
Tintermedium-Thint.05G0273900.1                      ---EP-------LPIC-----------------------
Taestivum-Traes_2DL_30F8555D4.1                      ---EP-------LPIC-----------------------
Taestivum-Traes_2DL_83168C1E0.1                      ---EP-------LPIC-----------------------
Taestivum-Traes_2DL_B978B4E22.1                      ---EP-------LPIC-----------------------
Taestivum-Traes_1AS_0B9295A68.1                      ---EP-------LPIC-----------------------
Taestivum-Traes_1BS_7068F56E6.1                      ---EP-------LPIC-----------------------
Taestivum-Traes_1BS_AEA817616.1                      ---EP-------LPIC-----------------------
Taestivum-Traes_4AL_892C47ED5.1                      ---EP-------LPLC-----------------------
Tintermedium-Thint.03G0055900.1                      ---EP-------LPIC-----------------------
Tintermedium-Thint.01G0012900.1                      ---EP-------LPIC-----------------------
Tintermedium-Thint.02G0158500.1                      ---EP-------LPIC-----------------------
Tintermedium-Thint.03G0055700.1                      ---EP-------LPIC-----------------------
Taestivum-Traes_1BS_C3AA04888.1                      ---------------------------------------
Taestivum-Traes_1DS_A8BD91E4A.1                      ---EP-------LPLC-----------------------
Taestivum-Traes_1BS_C26CCE0E9.1                      ---EP-------LPIC-----------------------
Taestivum-Traes_1DS_A21FCBA3D.1                      ---EP-------LPIC-----------------------
Tintermedium-Thint.01G0012200.1                      ---EP-------LPIC-----------------------
Tintermedium-Thint.02G0158600.1                      ---EP-------LPIC-----------------------
Tintermedium-Thint.01G0012700.1                      ---EP-------LPIC-----------------------
Taestivum-Traes_5BL_F50A47503.1                      ---EP-------LPIC-----------------------
Tintermedium-Thint.02G0158700.1                      ---EP-------LPIC-----------------------
Taestivum-Traes_5BL_AB528184A.1                      ---EP-------LPIC-----------------------
Tintermedium-Thint.03G0055800.1                      ---EP-------LPIC-----------------------
Taestivum-Traes_1AS_F9013A945.1                      ---EP-------LPIC-----------------------
```

```
Taestivum-Traes_1DS_43DAC7A7F.1              ---EP-------LPIC-----------------------
Taestivum-Traes_1BS_BD86C90A7.1              ---EP-------LPIC-----------------------
Tintermedium-Thint.02G0158900.1              ---EP-------LPLC-----------------------
Tintermedium-Thint.05G0274000.1              ---EP-------LPIC-----------------------
Osativa-LOC_Os04g43800.1                     ---EP-------LPIC-----------------------
Pvaginatum-Pavag06G160300.1                  ---EP-------LPIC-----------------------
Sbicolor-Sobic.006G148900.1                  ---EP-------LPIC-----------------------
Pvirgatum-Pavir.7KG237800.1                  ---EP-------LPIC-----------------------
Pvirgatum-Pavir.7NG355800.1                  ---EP-------LPIC-----------------------
Sviridis-Sevir.7G178300.1                    ---EP-------LSIC-----------------------
Zmays-Zm00001d003015_T001                    ---EP-------LPIC-----------------------
Hvulgare-HORVU1Hr1G022060.1                  ---EP-------LPIC-----------------------
Taestivum-Traes_1BS_723922D171.2             ---EP-------LPIC-----------------------
Taestivum-Traes_1DS_A171C7D59.1              ---EP-------LPIC-----------------------
Tintermedium-Thint.01G0111500.1              ---EP-------LPIC-----------------------
Tintermedium-Thint.03G0265300.1              ---EP-------LPIC-----------------------
Hvulgare-HORVU2Hr1G038140.1                  ---EP-------LPLC-----------------------
Tintermedium-Thint.04G0171200.1              ---EP-------LPLC-----------------------
Taestivum-Traes_2AS_958327519.3              ---EP-------LPLC-----------------------
Taestivum-Traes_2DS_28CA50371.1              ---EP-------LPLC-----------------------
Tintermedium-Thint.05G0160800.1              ---EP-------LPLC-----------------------
Tintermedium-Thint.06G0299000.1              ---EP-------LPLC-----------------------
Taestivum-Traes_2DS_3791A8A36.1              ---EP-------LPIC-----------------------
Tintermedium-Thint.06G0342800.1              ---EP-------LPIC-----------------------
Tintermedium-Thint.05G0150900.1              ---KP-------LPIC-----------------------
Tintermedium-Thint.04G0242300.1              ---EP-------LPIC-----------------------
Tintermedium-Thint.06G0298700.1              ---EP-------LPIC-----------------------
Taestivum-Traes_2BS_88CF42F2E.1              ---EP-------LPIC-----------------------
Tintermedium-Thint.04G0241900.1              ---EP-------LPLC-----------------------
Platifolius-Phala.06G165300.1                ---EP-------LPIC-----------------------
Streptochaeta-strangu_019385-RA              ---EP-------LPIC-----------------------
Streptochaeta-strangu_028944-RA              ---EP-------LSICRT---------------------
Acomosus-Aco010091.1                         ---AP-------LPIS-----------------------
Zmarina-Zosma03g22300.1                      ---SP-------LPIS-----------------------
Ecdeiocolea-Emoptg0003741_1G000600.1         ---EP-------LPIC-----------------------
Jascendens-Joasc.05G060500.1                 ---EP-------LPIC-----------------------
```

```
Ecdeiocolea-Emoptg0003741_1G000630.1         ---EP-------LPIC-----------------------
Jascendens-Joasc.05G060400.1                 ---EP-------LPIC-----------------------
Bdistachyon-Bradi3g49250.2                   ---EP-------LPINVV---------------------
Hvulgare-HORVU6Hr1G058820.1                  ---EP-------LPINV----------------------
Taestivum-Traes_6DL_1AEA7B869.1              ---EP-------LPINV----------------------
Tintermedium-Thint.18G0192300.1              ---EP-------LPINV----------------------
Tintermedium-Thint.17G0230800.1              ---EP-------LPINV----------------------
Tintermedium-Thint.17G0230900.1              ---EP-------LPINV----------------------
Ecoracana-ELECO.r07.2AG0138000.1             ---KP-------LPIN-----------------------
Ecoracana-ELECO.r07.2BG0193650.1             ---KP-------LPIN-----------------------
Pvaginatum-Pavag04G202700.1                  ---KP-------LPIN-----------------------
Sviridis-Sevir.1G245000.1                    ---KP-------LPIN-----------------------
Sviridis-Sevir.6G187100.1                    ---KP-------LPIN-----------------------
Pvirgatum-Pavir.1NG356200.1                  ---KP-------LPIN-----------------------
Sbicolor-Sobic.004G220300.1                  ---KP-------LPINVVN--------------------
Zmays-Zm00001d017274_T001                    ---KP-------LPINVK---------------------
Zmays-Zm00001d051161_T003                    ---KP-------LPININ---------------------
Sviridis-Sevir.7G178200.1                    ---KP-------LPIC-----------------------
Osativa-LOC_Os02g41630.2                     ---EP-------LPIN-----------------------
Ecoracana-ELECO.r07.4AG0310430.1             ---KP-------LPIC-----------------------
Ecoracana-ELECO.r07.4BG0341620.1             ---KP-------LPIS-----------------------
Pvaginatum-Pavag06G160200.1                  ---EP-------LPIC-----------------------
Pvirgatum-Pavir.7KG238255.1                  ---KP-------LPIC-----------------------
Sbicolor-Sobic.006G148800.1                  ---KP-------LPIC-----------------------
Zmays-Zm00001d003016_T001                    ---KP-------LPIC-----------------------
Osativa-LOC_Os04g43760.1                     ---EP-------LPIC-----------------------
Platifolius-Phala.02G210400.1                ---EP-------LPIC-----------------------
Platifolius-Phala.06G165100.1                ---EP-------LPIC-----------------------
Streptochaeta-strangu_019386-RA              ---EP-------LPIC-----------------------
Streptochaeta-strangu_020769-RA              ---EP-------LPIC-----------------------
Sviridis-Sevir.2G448300.1                    ---EP-------LPIC-----------------------
Sviridis-Sevir.7G177900.1                    ---EP-------LPIC-----------------------
```

**Fig. S8. Alignment of PTAL and PAL proteins from monocot.**

The amino acid sequences of PTAL and PAL enzymes from different monocot species were aligned using MAFFT to predict residues that are critical for the acquisition of TAL activity. PTAL sequences are found between two horizontal lines. The residues that are required for the general aromatic ammonia-lyase activity are shown in blue. Sixteen candidate residues determined by phylogeny-guided alignment analysis are highlighted in magenta and purple: 8 residues highly conserved uniquely in either PTALs and PALs are shown in magenta, and the additional 8 residues were highly conserved among PTALs but not among PALs are shown in purple.

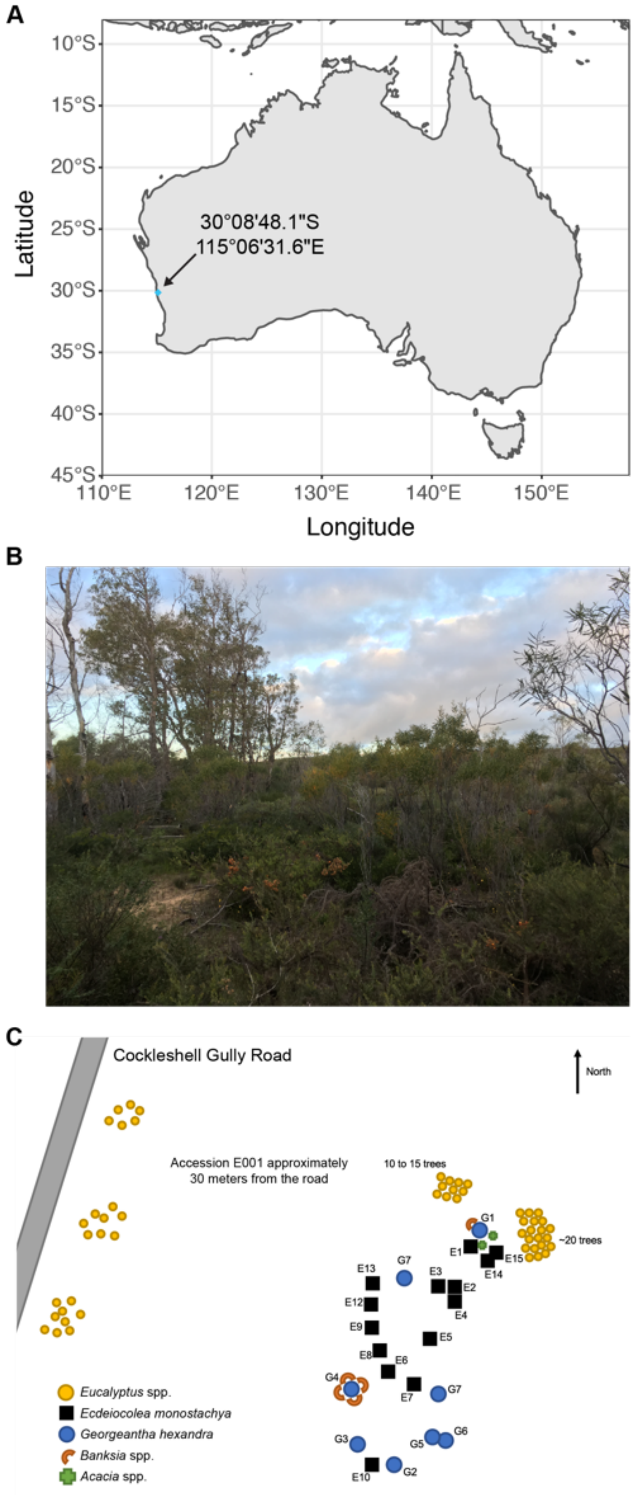


**Fig. S9. Identification and sampling site of *E. monostachya* accession E001.**
(**A**) Sampling sites in western Australia near Jurien Bay.
(**B**) Photograph of sampling site of *E. monostachya* accession E001.
(**C**) Sample site map of *E. monostachya* accession E001 with other plant species.

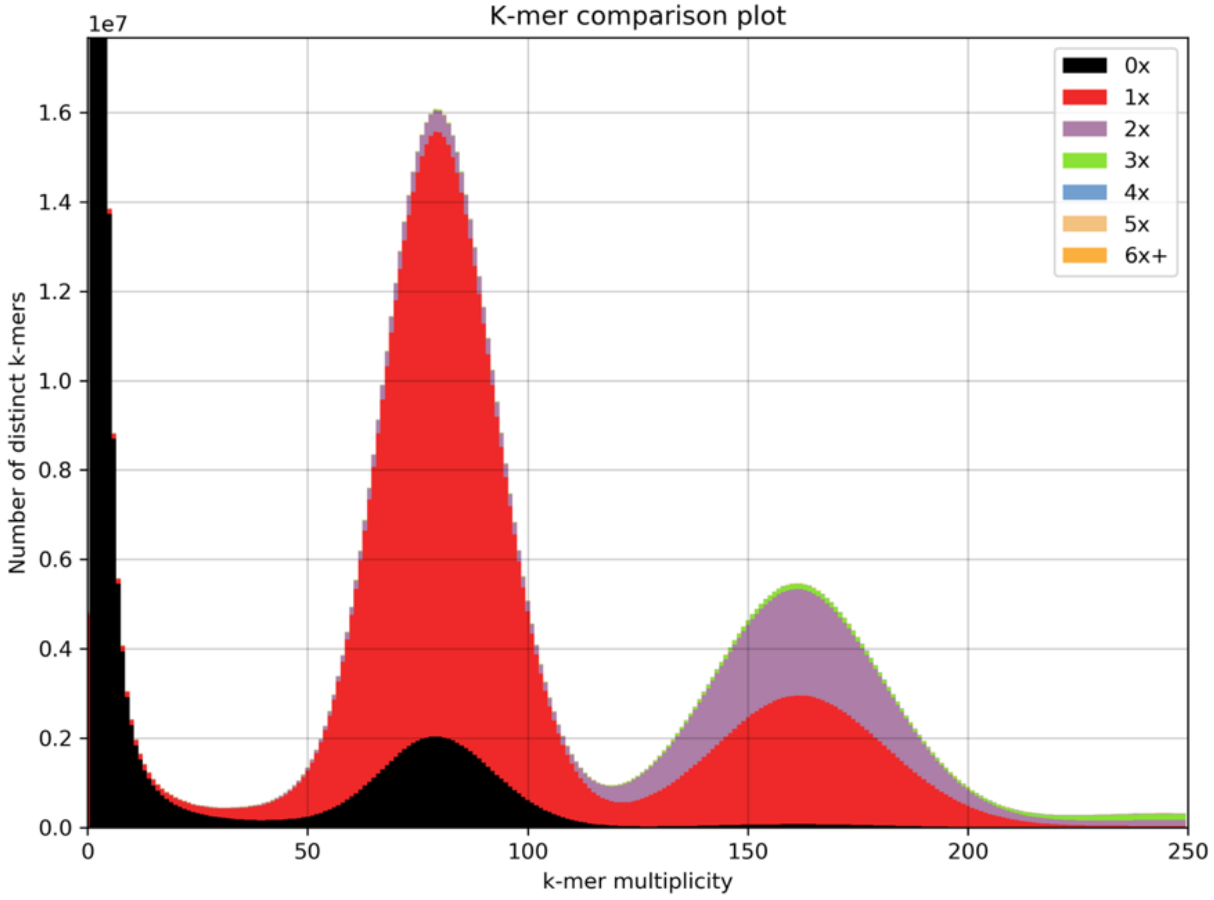


**Fig. S10. Distribution of *k*-mers from *E. monostachya* short read sequencing relative to genome.**

The stacked histogram was generated using the *k*-mer analysis toolkit (KAT; https://github.com/TGAC/KAT) with k=27 and shows the number of distinct *k*-mers present at varying copy numbers, including zero, in the finished genome assembly relative to their multiplicity in the short read data. *x*-axis, the occurrence of a *k*-mer in the short read data (Illumina). *y*-axis, the observed number of distinct *k*-mers. Color shading shows the number of times *k*-mers are observed in the assembled genome.

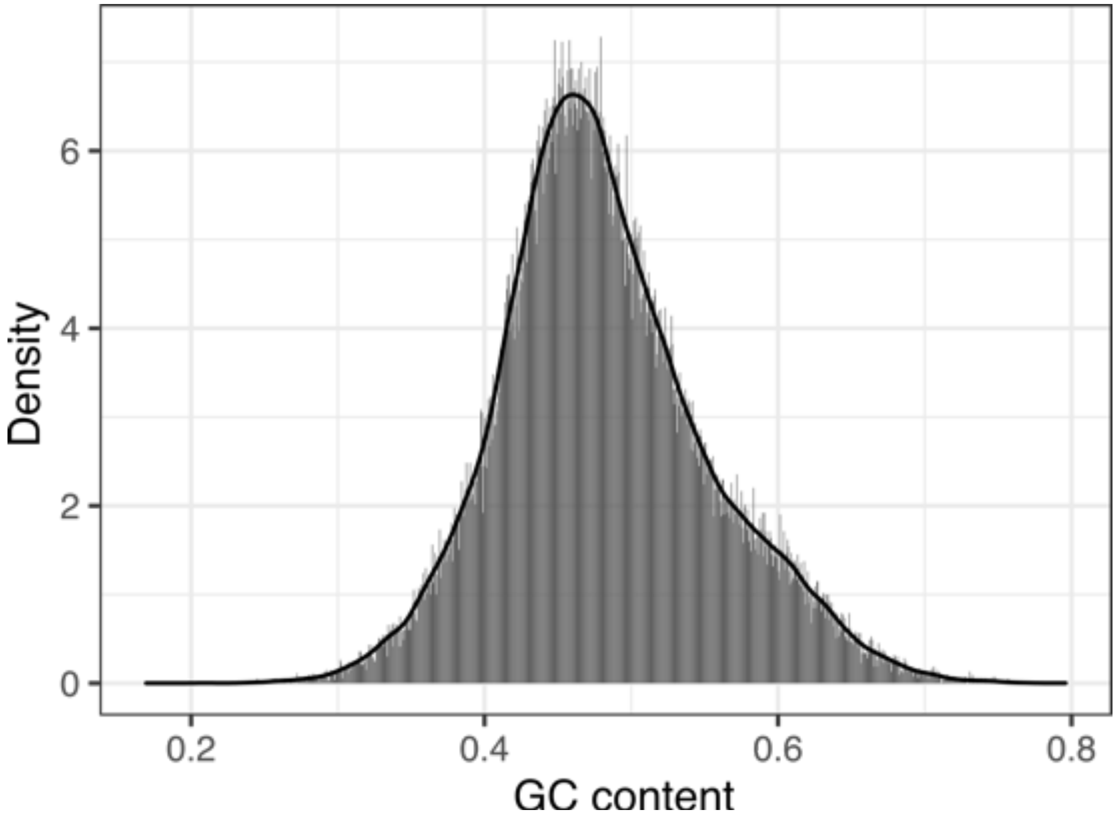


**Fig. S11. Distribution of *k*-mers from *E. monostachya* short read sequencing relative to genome.**
Histogram of GC content was estimated from the evidence-based gene annotations (84,700 gene models) fit with a Gaussian density distribution.

**Table S1. List of species used in the phylogenetic tree analysis and total putative paralogs per species estimated by PUG from gene trees**

| Order | Clade/Grade in phylogeny | Family | Species | Version | Source (reference) | Total putative paralogs |
|---|---|---|---|---|---|---|
| Amborellales | Amborellales | Amborellaceae | *Amborella trichopoda* Baill. | v.1.0 | Phytozome (*110*) | 19874 |
| Nymphaeales | Amborellales | Nymphaeaceae | *Nymphaea colorata* | v.1.2 | Phytozome (*111*) | 38172 |
| Laurales | Amborellales | Lauraceae | *Cinnamomum kanehirae* Hayata | v.3 | Phytozome (*112*) | 50536 |
| Ranunculales | Ranunculales | Ranunculaceae | *Aquilegia coerulea* E.James | v.3.1 | Phytozome (*113*) | 43154 |
| Apiales | Asterid | Apiaceae | *Daucus carota* L. | v.2.0 | Phytozome (*114*) | 40571 |
| Asterales | Asterid | Asteraceae | *Helianthus annuus* L. | v.1.2 | Phytozome (*115*) | 116488 |
| Solanales | Asterid | Solanaceae | *Solanum lycopersicum* L. | ITAG5.0 | Phytozome (*116*) | 49105 |
| Vitales | Rosid | Vitaceae | *Vitis vinifera* L. | v.2.1 | Phytozome (*41*) | 55313 |
| Caryophyllales | Caryophyllales | Chenopodiaceae | *Spinacia oleracea* L. | v.3 | Phytozome (*117*) | 49296 |
| Caryophyllales | Caryophyllales | Chenopodiaceae | *Beta vulgaris* L. | EL10.2_2 | Phytozome (*118*) | 27268 |
| Fabales | Rosid | Fabaceae | *Glycine max* (L.) Merr. | v.1.1 | Phytozome (*119*) | 142145 |
| Brassicales | Rosid | Brassicaceae | *Arabidopsis thaliana* L. | TAIR10 | Phytozome (*120*) | - |
| Malpighiales | Malpighiales | Salicaceae | *Populus trichocarpa* Torr. & A.*Gray ex. Hook.* | v.4.1 | Phytozome (*121*) | 64690 |
| Acorales | Alismatales and Acorales | Acoraceae | *Acorus americanus* Raf. | v.1.1 | Phytozome (new assembly) | 38115 |
| Alismatales | Alismatales and Acorales | Zosteraceae | *Zostera marina* L. | v.3.1 | Phytozome (*122*) | 15952 |
| Alismatales | Alismatales and Acorales | Araceae | *Spirodela polyrhiza* (L.) Schleid. | v.2 | Phytozome (*123*) | 16107 |
| Dioscoreales | Dioscoreales and Pandanales | Dioscoreaceae | *Dioscorea alata* L. | v.2.1 | Phytozome (*124*) | 24645 |
| Asparagales | Asparagales | Asparagaceae | *Asparagus officinalis* L. | v.1.1 | Phytozome (*125*) | 17273 |
| Arecales | Arecaceae | Arecaceae | *Phoenix dactylifera* L. | GCF_009389715.1 | NCBI (*126*) | 67833 |
| Zingiberales | Musaceae | Musacaceae | *Musa acuminata* Colla | v.1 | Phytozome (*127*) | 64579 |
| Poales | Other Poales | Bromeliaceae | *Ananas comosus* (L.) Merr. | v.3 | Phytozome (*128*) | 19100 |
| Poales | Other Poales | Typhaceae | *Typha latifolia* L. | v.1.1 | Phytozome (this study) | 24155 |
| Poales | Other Poales | Cyperaceae | *Carex scoparia* Schkuhr ex Willd. | GCA_024447655.1 | NCBI (*129*) | 29886 |
| Poales | Other Poales | Juncaceae | *Juncus effusus* L. | GCA_024447645.1 | NCBI (*129*) | 29969 |
| Poales | Poaceae | Joinvilleaceae | *Joinvillea ascendens* Gaudich. ex Brongn. & Gris | v.1.1 | Phytozome (this study) | 51896 |
| Poales | Poaceae | Ecdeiocoleaceae | *Ecdeiocolea monostachya* F.Muell | v.1.0 | NCBI (this study) | 102984 |
| Poales | Poaceae | Poaceae | *Streptochaeta angustifolia* Soderstr. | v.1.0 | (*7*) | 88410 |
| Poales | Poaceae | Poaceae | *Pharus latifolius* L. | v.1.1 | Phytozome (this study) | 53897 |
| Poales | Poaceae | Poaceae | *Oryza sativa* L. | v.7.0 | Phytozome (*130*) | 49391 |
| Poales | Poaceae | Poaceae | *Brachypodium distachyon* (L.) P.Beauv. | v.2.1 | Phytozome (*131*) | 36958 |
| Poales | Poaceae | Poaceae | *Hordeum vulgare* L. | r1 | Phytozome (*132*) | 69233 |
| Poales | Poaceae | Poaceae | *Triticum aestivum* L. | v.2.1 | Phytozome (*133*) | 451770 |
| Poales | Poaceae | Poaceae | *Oropetium thomaeum* (L.f.) Trin. | v.1.0 | Phytozome (*134*) | 20205 |
| Poales | Poaceae | Poaceae | *Eleusine coracana* Gaertn. | v.1.1 | Phytozome (*135*) | 144538 |
| Poales | Poaceae | Poaceae | *Setaria viridis* (L.) P.Beauv. | v.4.1 | Phytozome (*136*) | 40540 |
| Poales | Poaceae | Poaceae | *Panicum virgatum* L. | v.5.1 | Phytozome (*137*) | 212508 |
| Poales | Poaceae | Poeaceae | *Paspalum vaginatum* Sw. | v.3.1 | Phytozome (*138*) | 56010 |
| Poales | Poaceae | Poaceae | *Zea mays* L. | RefGen_V4 | Phytozome | 64387 |
| Poales | Poaceae | Poaceae | *Sorghum bicolor* (L.) Moench | v.3.1 | Phytozome (*139*) | 44420 |

**Table S2. Total counts of unique gene duplications mapped to placement of known WGM events on the species tree (tau, sigma, rho, maize, wheat, and gamma) and to portions of the tree without expected WGM events (BOP-PACMAD divergence, prior to PACMAD diversification, and prior to eudicot diversification).**

| **WGM Event** | **Total Unique Pairs** | |
|---|---|---|
| | **80 BSV** | **50 BSV** |
| *tau* | 972 | 1,258 |
| *sigma* | 1,122 | 1,325 |
| *rho* | 3,004 | 3,284 |
| Maize | 2,196 | 2,232 |
| Wheat | 3,200 | 3,290 |
| BOP-PACMAD | 868 | 995 |
| PACMAD | 936 | 1,020 |
| *gamma* | 1212 | 1502 |
| Eudicots | 803 | 920 |

**Table S3. Kinetics parameters of recombinant PTAL ortholog proteins with or without mutations.**

| | TAL assay | | | PAL assay | | |
|---|---|---|---|---|---|---|
| **protein name** | **$K_m$ (μM)** | **$k_{cat}$ ($s^{-1}$)** | **$k_{cat}/K_m$ ($s^{-1}$ $mM^{-1}$)** | **$K_m$ (μM)** | **$k_{cat}$ ($s^{-1}$)** | **$k_{cat}/K_m$ ($s^{-1}$ $mM^{-1}$)** |
| **SbPTAL** | 10.8 ± 2.2 | 0.09 ± 0.00 | 7.96 ± 1.18 | 150.1 ± 14.4 | 0.69 ± 0.01 | 4.63 ± 0.49 |
| **BdPTAL** | 19.1 ± 2.4 | 0.09 ± 0.00 | 4.78 ± 0.36 | 216.6 ± 10.3 | 1.05 ± 0.05 | 4.84 ± 0.11 |
| **SaPTAL-a** | 13.3 ± 1.0 | 0.04 ± 0.00 | 3.32 ± 0.13 | 154.5 ± 3.7 | 0.39 ± 0.01 | 2.51 ± 0.03 |
| **SaPTAL-b** | 16.2 ± 0.5 | 0.06 ± 0.00 | 3.57 ± 0.07 | 227.4 ± 1.5 | 0.56 ± 0.02 | 2.46 ± 0.10 |
| **EmoPTAL** | 16.3 ± 1.2 | 0.04 ± 0.0 | 2.55 ± 0.16 | 64.1 ± 3.1 | 0.48 ± 0.00 | 7.54 ± 0.39 |
| **JaPTAL** | 11.0 ± 0.4 | 0.04 ± 0.00 | 3.68 ± 0.22 | 65.6 ± 1.4 | 0.45 ± 0.02 | 6.80 ± 0.18 |
| **JaPAL** | 3532.5 ± 446.5 | 0.03 ± 0.00 | 0.01 ± 0.00 | 24.4 ± 0.1 | 1.92 ± 0.01 | 78.60 ± 0.00 |
| **EmoPAL** | 4226.2 ± 150.6 | 0.03 ± 0.00 | 0.01 ± 0.00 | 27.7 ± 1.9 | 1.23 ± 0.02 | 44.70 ± 2.39 |
| **BdPAL** | 3448.6 ± 1045.4 | 0.02 ± 0.01 | 0.01 ± 0.00 | 21.3 ± 2.1 | 0.86 ± 0.03 | 40.74 ± 2.68 |
| **SbPAL** | 5347.8 ± 1284.4 | 0.04 ± 0.01 | 0.01 ± 0.00 | 43.9 ± 2.7 | 0.97 ± 0.02 | 22.05 ± 1.08 |
| **SbPTAL$^{H123F}$** | 750.7 ± 36.8 | 0.03 ± 0.00 | 0.04 ± 0.00 | 3.8 ± 2.0 | 0.10 ± 0.00 | 30.97 ± 13.70 |
| **BdPTAL$^{H123F}$** | 765.2 ± 65.8 | 0.03 ± 0.00 | 0.04 ± 0.00 | 6.0 ± 1.4 | 0.17 ± 0.00 | 29.92 ± 6.75 |
| **SaPTAL-a$^{H118F}$** | 531.0 ± 13.2 | 0.03 ± 0.00 | 0.06 ± 0.00 | 6.0 ± 0.8 | 0.20 ± 0.00 | 33.60 ± 5.10 |
| **SaPTAL-b$^{H126F}$** | 723.6 ± 54.8 | 0.05 ± 0.00 | 0.06 ± 0.01 | 6.3 ± 0.5 | 0.23 ± 0.00 | 37.05 ±2.12 |
| **EmoPTAL$^{H127F}$** | 613.5 ±18.1 | 0.04 ± 0.0 | 0.07 ± 0.01 | 3.6 ± 0.5 | 0.12 ± 0.00 | 32.62 ± 2.12 |
| **JaPTAL$^{H125F}$** | 535.4 ± 80.5 | 0.02 ± 0.00 | 0.04 ± 0.00 | 6.8 ± 0.7 | 0.08 ± 0.00 | 12.28 ± 0.71 |
| **JaPAL$^{F140H}$** | 222.7 ± 13.5 | 0.03 ± 0.00 | 0.13 ± 0.01 | 697.0 ± 169.2 | 0.60 ± 0.04 | 0.89 ± 0.18 |
| **EmoPAL$^{F134H}$** | 450.1 ± 14.2 | 0.02 ± 0.00 | 0.05 ± 0.00 | 1305.3 ± 25.6 | 0.42 ± 0.01 | 0.32 ± 0.01 |
| **BdPAL$^{F137H}$** | 371.3 ± 15.4 | 0.04 ± 0.00 | 0.11 ± 0.01 | 1082.0 ± 58.1 | 0.75 ± 0.01 | 0.70 ± 0.03 |
| **SbPAL$^{F135H}$** | 412.8 ± 6.5 | 0.04 ± 0.00 | 0.10 ± 0.00 | 1051.6 ± 58.5 | 0.88 ± 0.05 | 0.84 ± 0.02 |
| **JaPAL$^{F140H_MUT8}$** | 17.9 ± 2.0 | 0.03 ± 0.00 | 1.81 ± 0.18 | 141.0 ± 6.3 | 0.60 ± 0.01 | 4.25 ± 0.22 |
| **JaPAL$^{F140H_MUT16}$** | 42.2 ± 2.1 | 0.02 ± 0.00 | 0.56 ± 0.02 | 454.9 ± 4.6 | 0.46 ± 0.01 | 1.01 ± 0.0 |
| **JaPAL$^{F140H_MUT8_I102V}$** | 24.5 ± 1.4 | 0.04 ± 0.00 | 1.67 ± 0.08 | 231.7 ± 3.5 | 0.74 ± 0.01 | 3.19 ± 0.08 |
| **JaPAL$^{F140H_MUT8_I122S}$** | 282.3 ± 29.0 | 0.02 ± 0.00 | 0.07 ± 0.00 | 2290.9 ± 344.4 | 0.29 ± 0.04 | 0.31 ± 0.00 |
| **JaPAL$^{F140H_MUT8_G121A}$** | 12.6 ± 0.6 | 0.03 ± 0.00 | 2.12 ± 0.07 | 58.1 ± 3.4 | 0.57 ± 0.02 | 9.88 ± 0.31 |
| **JaPAL$^{F140H_MUT8_L138I}$** | 18.3 ± 0.6 | 0.05 ± 0.00 | 2.50 ± 0.02 | 74.3 ± 1.1 | 0.66 ± 0.01 | 8.86 ± 0.01 |
| **JaPAL$^{F140H_MUT8_S267A}$** | 43.1 ± 1.5 | 0.04 ± 0.00 | 0.96 ± 0.03 | 316.9 ± 6.6 | 0.85 ± 0.04 | 2.67 ± 0.06 |
| **JaPAL$^{F140H_MUT8_T444P}$** | 25.3 ± 2.4 | 0.04 ± 0.00 | 1.61 ± 0.09 | 191.9 ± 3.6 | 0.90 ± 0.05 | 4.70 ± 0.22 |
| **JaPAL$^{F140H_MUT8_A448S}$** | 23.3 ± 1.0 | 0.04 ± 0.00 | 1.62 ± 0.04 | 167.7 ± 11.3 | 0.80 ± 0.03 | 4.79 ± 0.43 |
| **JaPAL$^{F140H_MUT8_V500I}$** | 25.1 ± 3.2 | 0.05 ± 0.00 | 1.82 ± 0.14 | 150.7 ± 3.7 | 0.82 ± 0.02 | 5.45 ± 0.14 |
| **JaPAL$^{S112I}$** | 353.5 ± 45.0 | 0.05 ± 0.00 | 0.13 ± 0.01 | 2.6 ± 0.5 | 0.21 ± 0.04 | 79.80 ± 0.00 |
| **JaPAL$^{F140H_S112I}$** | 17.2 ± 0.7 | 0.03 ± 0.00 | 1.79 ± 0.10 | 67.3 ± 2.5 | 0.77 ± 0.01 | 11.46 ± 0.21 |
| **JaPTAL$^{H125F_I97S}$** | Only trace activity detected | | | 9.42 ± 0.8 | 0.02 ± 0.00 | 1.66 ± 0.12 |
| **AtPAL1** | 3069.8 ± 433.4 | 0.05 ± 0.00 | 0.02 ± 0.00 | 52.2 ± 3.1 | 1.42 ± 0.07 | 27.31 ± 0.85 |
| **AtPAL1$^{S114I}$** | 515.4 ± 54.3 | 0.02 ± 0.00 | 0.04 ± 0.00 | 10.1 ± 1.9 | 0.23 ± 0.00 | 23.71 ± 4.63 |
| **AtPAL1$^{F144H}$** | 313.9 ± 13.9 | 0.01 ± 0.00 | 0.03 ± 0.00 | 1198.9 ± 21.1 | 1.58 ± 0.04 | 1.32 ± 0.02 |
| **AtPAL1$^{F144H_S114I}$** | 20.2 ± 0.2 | 0.02 ± 0.00 | 1.05 ± 0.02 | 87.3 ± 2.9 | 0.88 ± 0.01 | 10.07 ± 0.44 |

**Table S4. Residues potentially involved in the transition from PAL to PTAL in graminids.**

| Residue No. | PAL | PTAL | probability of positive selection |
|---|---|---|---|
| 70 | A/S | G | 0.945 |
| 102 | V | I | 0.929 |
| 103 | M/L | L/V | 0.934 |
| 106 | M | L/V/I | 0.935 |
| 107 | N/M/A/S | A | 0.792 |
| 110 | T/V/G | G | 0.986 |
| 112 | S | I | 0.938 |
| 121 | A | G | 0.940 |
| 128 | K | R/K | 0.817 |
| 129 | E/Q/K | D | 0.890 |
| 131 | G/A | P/H | 0.849 |
| 135 | R/K/Q/A | V | 0.957 |
| 138 | I | L | 0.952 |
| 140 | F | H | 0.999 |
| 145 | V/I/A | I | 0.924 |
| 226 | S/A | A | 0.942 |
| 253 | E | K/T | 0.932 |
| 267 | A | S | 0.940 |
| 270 | S | A/S | 0.953 |
| 271 | G/A | A | 0.940 |
| 279 | E/D | D | 0.926 |
| 334 | Y/F | F | 0.939 |
| 348 | Q/M | L/M | 0.735 |
| 390 | L | I/V | 0.916 |
| 422 | A | N/S | 0.999 |
| 444 | P | T | 0.953 |
| 448 | S | A | 0.940 |
| 462 | A | T/A | 0.725 |
| 500 | I | V | 0.929 |
| 502 | S/A | A | 0.713 |

The numbering of residues is in reference to JaPAL. The histidine H140 in PTAL, shown in blue, have been predicted to play a critical role in the acquisition of TAL activity. The sixteen residues highlighted with magenta and purple were mutated in the JaPAL$^{F140H_MUT16}$ mutant, while the eight out of these sixteen residues mutated in the JaPAL$^{F140H_MUT8}$ mutant are marked with magenta. Their positions in the protein sequence alignment are shown in **fig. S8**. Other residues under positive selection with a probability greater than 0.7 but were not biochemically tested are shown in black.

**Table S5. Genomic libraries included in the *Pharus latifolius* (var. MRM_UAlabama_McKain320) genome assembly and their respective assembled sequence coverage levels in the final release.**

| Library | Sequencing Platform | Average Read/Insert Size | Read Number | Assembled Sequence Coverage (x) |
|---|---|---|---|---|
| JAIJ | Illumina (2x150) | 400 | 1,313,004,860 | 170.65 |
| JBMT | Illumina-HiC (2x150) | N/A | 588,146,836 | 76.53 |
| PBZA/PBZB | PACBIO | 19,192* | 2,058,145 | 35.08 |
| **Total** | | N/A | 1,903,209,841 | 282.26 |

A total of 2,058,145 PACBIO reads (35.08x) were assembled using HiFiAsm assembler (*140*), and formed the starting point of the version 1.0 release. The 1,313,004,860 Illumina fragment 2x150 reads (170.65x) was used for fixing homozygous snp/indel errors in the consensus. Chromosomes were scaffolded using the 588,146,836 2x150 (76.53x) HiC reads.

*Average read length of PACBIO reads.

**Table S6. PACBIO CCS library statistics for the libraries included in the *Pharus latifolius* (var. MRM_UAlabama_McKain320) genome assembly and their respective assembled sequence coverage levels.**

| Cutoff | Number of Reads | Basepairs | Average Read Length | Coverage |
|---|---|---|---|---|
| 0 | 2,058,145 | 39,217,569,760 | 19,192 | 35.08x |
| 1,000 | 2,057,958 | 39,217,524,178 | 19,192 | 35.08x |
| 2,000 | 2,056,519 | 39,215,080,637 | 19,195 | 35.07x |
| 3,000 | 2,048,241 | 39,193,485,304 | 19,207 | 35.05x |
| 4,000 | 2,027,917 | 39,121,459,634 | 19,239 | 34.99x |
| 5,000 | 2,006,311 | 39,024,900,196 | 19,272 | 34.90x |
| 6,000 | 1,991,711 | 38,944,815,040 | 19,295 | 34.83x |
| 7,000 | 1,977,958 | 38,855,464,457 | 19,316 | 34.75x |
| 8,000 | 1,967,028 | 38,773,911,238 | 19,333 | 34.67x |
| 9,000 | 1,957,555 | 38,693,375,134 | 19,348 | 34.60x |
| 10,000 | 1,947,335 | 38,596,180,530 | 19,365 | 34.51x |
| 11,000 | 1,935,684 | 38,473,730,818 | 19,383 | 34.41x |
| 12,000 | 1,922,073 | 38,317,032,622 | 19,405 | 34.26x |
| 13,000 | 1,905,463 | 38,109,121,466 | 19,431 | 34.08x |
| 14,000 | 1,884,401 | 37,824,381,317 | 19,466 | 33.83x |
| 15,000 | 1,856,914 | 37,425,170,159 | 19,510 | 33.47x |
| 16,000 | 1,815,982 | 36,789,040,194 | 19,578 | 32.90x |
| 17,000 | 1,718,889 | 35,177,726,295 | 19,742 | 31.46x |
| 18,000 | 1,449,623 | 30,450,142,349 | 20,239 | 27.23x |
| 19,000 | 1,093,009 | 23,854,196,033 | 21,032 | 21.33x |

**Table S7. Summary statistics of the initial output of the RACON polished HiFiAsm+HIC assembly of the *Pharus latifolius* (var. MRM_UAlabama_McKain320) genome.**

| Minimum Scaffold Length | Number of Scaffolds | Number of Contigs | Scaffold Size | Basepairs | % Non-gap Basepairs |
|---|---|---|---|---|---|
| 5 Mb | 42 | 42 | 1,067,417,875 | 1,067,417,875 | 100.00% |
| 2.5 Mb | 51 | 51 | 1,097,864,293 | 1,097,864,293 | 100.00% |
| 1 Mb | 55 | 55 | 1,104,624,527 | 1,104,624,527 | 100.00% |
| 500 Kb | 59 | 59 | 1,107,616,505 | 1,107,616,505 | 100.00% |
| 250 Kb | 60 | 60 | 1,107,916,669 | 1,107,916,669 | 100.00% |
| 100 Kb | 67 | 67 | 1,108,868,307 | 1,108,868,307 | 100.00% |
| 50 Kb | 285 | 285 | 1,121,497,893 | 1,121,497,893 | 100.00% |
| 25 Kb | 3,205 | 3,205 | 1,233,074,669 | 1,233,074,669 | 100.00% |
| 10 Kb | 3,267 | 3,267 | 1,234,523,116 | 1,234,523,116 | 100.00% |
| 5 Kb | 3,267 | 3,267 | 1,234,523,116 | 1,234,523,116 | 100.00% |
| 2.5 Kb | 3,267 | 3,267 | 1,234,523,116 | 1,234,523,116 | 100.00% |
| 1 Kb | 3,267 | 3,267 | 1,234,523,116 | 1,234,523,116 | 100.00% |
| 0 bp | 3,267 | 3,267 | 1,234,523,116 | 1,234,523,116 | 100.00% |

The table shows total contigs and total assembled base pairs for each set of scaffolds greater than the size listed in the left hand column.

**Table S8. Final summary assembly statistics for the version 1 chromosome scale assembly of *Pharus latifolius* (var. MRM_UAlabama_McKain320).**

| | |
|---|---|
| **Scaffold total** | 198 |
| **Contig total** | 222 |
| **Scaffold sequence total** | 1,118.2 Mb |
| **Chromosome Sequence** | 1,106.8 Mb |
| **Contig sequence total** | 1,117.9 Mb (0.03% gap) |
| **Scaffold N/L50** | 6 / 97.0 Mb |
| **Contig N/L50** | 8 / 57.1 Mb |

**Table S9. Genomic libraries included in the *Typha latifolia* (var. 2019.6) genome assembly and their respective assembled sequence coverage levels in the final release.**

| Library | Sequencing Platform | Average Read/Insert Size | Read Number | Assembled Sequence Coverage (x) |
|---|---|---|---|---|
| KCWP | Illumina-OmniC (2x150) | N/A | 395,712,768 | 230.01 |
| ISAW | Illumina (2x150) | 400 | 359,449,236 | 261.16 |
| | PACBIO | 21,123* | 1,066,852 | 104.23 |
| Total | | N/A | 756,228,856 | 595.4 |

A total of 1,066,852 PACBIO reads (104.23x per haplotype) were assembled using HiFiAsm+HIC assembler (*140*), and formed the starting point of the version 1.0 release. The 359,449,236 Illumina fragment 2x150 reads (261.16x per haplotype) was used for fixing homozygous snp/indel errors in the consensus. Chromosomes were scaffolded using the 395,712,768 2x150 (230.01x) OmniC reads.

*Average read length of PACBIO reads.

**Table S10. PACBIO CCS library statistics for the libraries included in the *Typha latifolia* (var. 2019.6) genome assembly and their respective assembled sequence coverage levels.**

| Cutoff | Number of Reads | Basepairs | Average Read Length | Coverage |
|---|---|---|---|---|
| 0 | 1,066,852 | 22,411,463,488 | 21,123 | 104.23x |
| 1,000 | 1,066,770 | 22,411,444,606 | 21,123 | 104.23x |
| 2,000 | 1,065,878 | 22,409,980,209 | 21,126 | 104.23x |
| 3,000 | 1,064,137 | 22,405,562,694 | 21,132 | 104.22x |
| 4,000 | 1,060,686 | 22,393,309,138 | 21,144 | 104.15x |
| 5,000 | 1,055,931 | 22,371,877,766 | 21,161 | 104.05x |
| 6,000 | 1,050,223 | 22,340,333,463 | 21,182 | 103.91x |
| 7,000 | 1,042,696 | 22,291,295,367 | 21,209 | 103.67x |
| 8,000 | 1,033,792 | 22,224,362,976 | 21,241 | 103.37x |
| 9,000 | 1,023,756 | 22,139,118,284 | 21,277 | 102.98x |
| 10,000 | 1,016,020 | 22,065,885,615 | 21,305 | 102.63x |
| 11,000 | 1,011,577 | 22,019,480,734 | 21,321 | 102.42x |
| 12,000 | 1,009,671 | 21,997,671,638 | 21,328 | 102.32x |
| 13,000 | 1,008,094 | 21,977,925,307 | 21,334 | 102.22x |
| 14,000 | 1,005,431 | 21,941,866,652 | 21,344 | 102.06x |
| 15,000 | 1,000,579 | 21,871,231,494 | 21,362 | 101.72x |
| 16,000 | 990,031 | 21,707,077,668 | 21,401 | 100.97x |
| 17,000 | 968,648 | 21,353,179,592 | 21,482 | 99.32x |
| 18,000 | 925,436 | 20,594,173,169 | 21,648 | 95.79x |
| 19,000 | 834,937 | 18,914,964,217 | 22,013 | 87.98x |

**Table S11. Summary statistics of the initial output of the HAP1 RACON polished HiFiAsm+HIC assembly of the *Typha latifolia* (var. 2019.6) genome.**

| Minimum Scaffold Length | Number of Scaffolds | Number of Contigs | Scaffold Size | Basepairs | % Non-gap Basepairs |
|---|---|---|---|---|---|
| 5 Mb | 15 | 15 | 190,010,365 | 190,010,365 | 100.00% |
| 2.5 Mb | 20 | 20 | 207,810,681 | 207,810,681 | 100.00% |
| 1 Mb | 24 | 24 | 215,055,183 | 215,055,183 | 100.00% |
| 500 Kb | 25 | 25 | 215,708,286 | 215,708,286 | 100.00% |
| 250 Kb | 25 | 25 | 215,708,286 | 215,708,286 | 100.00% |
| 100 Kb | 49 | 49 | 218,941,164 | 218,941,164 | 100.00% |
| 50 Kb | 154 | 154 | 225,917,788 | 225,917,788 | 100.00% |
| 25 Kb | 395 | 395 | 235,196,222 | 235,196,222 | 100.00% |
| 10 Kb | 410 | 410 | 235,539,149 | 235,539,149 | 100.00% |
| 5 Kb | 410 | 410 | 235,539,149 | 235,539,149 | 100.00% |
| 2.5 Kb | 410 | 410 | 235,539,149 | 235,539,149 | 100.00% |
| 1 Kb | 410 | 410 | 235,539,149 | 235,539,149 | 100.00% |
| 0 bp | 410 | 410 | 235,539,149 | 235,539,149 | 100.00% |

The table shows total contigs and total assembled base pairs for each set of scaffolds greater than the size listed in the left hand column.

**Table S12. Summary statistics of the initial output of the HAP2 RACON polished HiFiAsm+HIC assembly of the *Typha latifolia* (var. 2019.6) genome.**

| Minimum Scaffold Length | Number of Scaffolds | Number of Contigs | Scaffold Size | Basepairs | % Non-gap Basepairs |
|---|---|---|---|---|---|
| 5 Mb | 16 | 16 | 178,012,269 | 178,012,269 | 100.00% |
| 2.5 Mb | 24 | 24 | 207,147,819 | 207,147,819 | 100.00% |
| 1 Mb | 28 | 28 | 214,409,968 | 214,409,968 | 100.00% |
| 500 Kb | 29 | 29 | 214,915,898 | 214,915,898 | 100.00% |
| 250 Kb | 30 | 30 | 215,308,267 | 215,308,267 | 100.00% |
| 100 Kb | 55 | 55 | 218,634,780 | 218,634,780 | 100.00% |
| 50 Kb | 155 | 155 | 225,215,235 | 225,215,235 | 100.00% |
| 25 Kb | 377 | 377 | 232,841,129 | 232,841,129 | 100.00% |
| 10 Kb | 421 | 421 | 233,857,847 | 233,857,847 | 100.00% |
| 5 Kb | 421 | 421 | 233,857,847 | 233,857,847 | 100.00% |
| 2.5 Kb | 421 | 421 | 233,857,847 | 233,857,847 | 100.00% |
| 1 Kb | 421 | 421 | 233,857,847 | 233,857,847 | 100.00% |
| 0 bp | 421 | 421 | 233,857,847 | 233,857,847 | 100.00% |

The table shows total contigs and total assembled base pairs for each set of scaffolds greater than the size listed in the left hand column.

**Table S13. Final summary assembly statistics for the version 1 HAP1 chromosome scale assembly of *Typha latifolia* (var. 2019.6).**

| | |
|---|---|
| **Scaffold total** | 16 |
| **Contig total** | 30 |
| **Scaffold sequence total** | 215.3 Mb |
| **Chromosome Sequence** | 215.1 Mb |
| **Contig sequence total** | 215.2 Mb (0.1% gap) |
| **Scaffold N/L50** | 6 / 14.6 Mb |
| **Contig N/L50** | 8 / 9.9 Mb |

**Table S14. Final summary assembly statistics for the version 1 HAP2 chromosome scale assembly of *Typha latifolia* (var. 2019.6).**

| | |
|---|---|
| **Scaffold total** | 15 |
| **Contig total** | 20 |
| **Scaffold sequence total** | 214.7 Mb |
| **Chromosome Sequence** | 214.7 Mb |
| **Contig sequence total** | 214.7 Mb (0.0% gap) |
| **Scaffold N/L50** | 6 / 14.6 Mb |
| **Contig N/L50** | 6 / 13.5 Mb |

### Table S15. Primers used in this study

| sequence (5' to 3') | purpose | template | Laboratory ID |
|---|---|---|---|
| **nested PCR and in-fusion cloning** | | | |
| CGCGCGGCAGCCATATGATGGCGTTCCAGAACGAC | in-fusion cloning of JaPTAL into pET28a | *Joinvillea ascendens* cDNA | pHM1810 |
| GCTCGAATTCGGATCCTCAGCAGATTGGCAGGGG | in-fusion cloning of JaPTAL into pET28a | *Joinvillea ascendens* cDNA | pHM1811 |
| CAATTGCAGGGAGATCGAGC | nested PCR for JaPAL | *Joinvillea ascendens* cDNA | pHM1869 |
| TGCTGTTGTAAGGTGGGGAT | nested PCR for JaPAL | *Joinvillea ascendens* cDNA | pHM1870 |
| CGCGCGGCAGCCATATGATGGAGTGCGAGAACGGC | in-fusion cloning of JaPAL into pET28a | *Joinvillea ascendens* cDNA | pHM1812 |
| GCTCGAATTCGGATCCTCAGCAGATTGGCAGGGG | in-fusion cloning of JaPAL into pET28a | *Joinvillea ascendens* cDNA | pHM1813 |
| TCTTCTTCCACACCAAACG | nested PCR for SaPTAL-a | *Streptochaeta angustifolia* cDNA | pHM1851 |
| GCACAAGAAGGATGCTAGAAAC | nested PCR for SaPTAL-a | *Streptochaeta angustifolia* cDNA | pHM1852 |
| CGCGCGGCAGCCATATGATGGCGAGCCAGAGGGAC | in-fusion cloning of SaPTAL-a into pET28a | *Streptochaeta angustifolia* cDNA | pHM1814 |
| GCTCGAATTCGGATCCTTAGCAGATGGGCAGGGG | in-fusion cloning of SaPTAL-a into pET28a | *Streptochaeta angustifolia* cDNA | pHM1815 |
| ATGGTGGCCCAGAGCGAC | nested PCR for SaPTAL-b | *Streptochaeta angustifolia* cDNA | pHM1841 |
| TTAGCAGATTGGAAGGGGC | nested PCR for SaPTAL-b | *Streptochaeta angustifolia* cDNA | pHM1842 |
| CGCGCGGCAGCCATATGATGGTGGCCCAGAGCGAC | in-fusion cloning of SaPTAL-b into pET28a | *Streptochaeta angustifolia* cDNA | pHM1816 |
| GCTCGAATTCGGATCCTTAGCAGATTGGAAGGGGC | in-fusion cloning of SaPTAL-b into pET28a | *Streptochaeta angustifolia* cDNA | pHM1817 |
| CAAGAAGAGCACGCCAACTC | nested PCR for SbPTAL | *Sorghum bicolor* RTx430 cDNA | pHM2009 |
| GCCACACACACATACGGATC | nested PCR for SbPTAL | *Sorghum bicolor* RTx430 cDNA | pHM2010 |
| GCGCGGCAGCCATATGATGGCGGGCAACGGCGCC | in-fusion cloning of SbPTAL into pET28a | *Sorghum bicolor* RTx430 cDNA | pHM2011 |
| GCTCGAATTCGGATCCTTAGTTGACGACGTTGAT | in-fusion cloning of SbPTAL into pET28a | *Sorghum bicolor* RTx430 cDNA | pHM2012 |
| CCACTGTCAGTCACGCAATT | nested PCR for SbPAL | *Sorghum bicolor* RTx430 cDNA | pHM2066 |
| TGCAACAGCCAAGAACATGC | nested PCR for SbPAL | *Sorghum bicolor* RTx430 cDNA | pHM2067 |
| GCGCGGCAGCCATATGATGGAGTGCGAGACGGGT | in-fusion cloning of SbPAL into pET28a | *Sorghum bicolor* RTx430 cDNA | pHM2068 |
| GCTCGAATTCGGATCCTCAGCAGAGCGGCAGTGG | in-fusion cloning of SbPAL into pET28a | *Sorghum bicolor* RTx430 cDNA | pHM2069 |
| CTCTGCAATTCGACGAGCTC | nested PCR for BdPAL | *Brachypodium distachyon* BL31 cDNA | pHM2072 |
| AGTTCTACTGGCTGCCTACC | nested PCR for BdPAL | *Brachypodium distachyon* BL31 cDNA | pHM2073 |
| GCGCGGCAGCCATATGATGGAGTACGAGAACGGG | in-fusion cloning of BdPAL into pET28a | *Brachypodium distachyon* BL31 cDNA | pHM2074 |
| GCTCGAATTCGGATCCTCAGCAGAGAGGCAGGGG | in-fusion cloning of BdPAL into pET28a | *Brachypodium distachyon* BL31 cDNA | pHM2075 |
| AGCTCCTATCTTCTTTCTTTCT | nested PCR for AtPAL1 | *Arabidopsis thaliana* cDNA | pHM2536 |
| AACCACTTCACAGACAATCA | nested PCR for AtPAL1 | *Arabidopsis thaliana* cDNA | pHM2537 |
| CGCGCGGCAGCCATATGATGGAGATTAACGGGGCACAC | in-fusion cloning of AtPAL1 into pET28a | *Arabidopsis thaliana* cDNA | pHM2522 |
| GCTCGAATTCGGATCCTTAACATATTGGAATGGGAGCTCCG | in-fusion cloning of AtPAL1 into pET28a | *Arabidopsis thaliana* cDNA | pHM2523 |
| **Sequencing analysis** | | | |
| CGACTCACTATAGGGGAATTGTG | sequencing of pET28a vectors | All of the pET28a construct generated | pHM1826 |
| GCTAGTTATTGCTCAGCGGTG | sequencing of pET28a vectors | All of the pET28a construct generated | pHM1827 |
| CATTCAAGATCGCCGGCATC | sequencing of JaPTAL-pET28a | JaPTAL-pET28a | pHM1828 |
| CTAACATCGAACTTGGCCGG | sequencing of JaPTAL-pET28a | JaPTAL-pET28a | pHM1829 |
| TCTTCCTGGCAGAGACAAGG | sequencing of JaPTAL-pET28a | JaPTAL-pET28a | pHM1863 |
| TTCCTCAATGCCGGAGTCTT | sequencing of JaPAL-pET28a | JaPAL-pET28a | pHM1830 |
| CTTCTGCGAAGTCATGACCG | sequencing of JaPAL-pET28a | JaPAL-pET28a | pHM1831 |
| CAACCCAGTGACCAACCATG | sequencing of JaPAL-pET28a | JaPAL-pET28a | pHM1832 |

| | | | |
|---|---|---|---|
| CTACGACGCCAACATTCTCG | sequencing of SaPTAL-a-pET28a | SaPTAL-a-pET28a | pHM1833 |
| ACATCGGCAAGCTCATGTTC | sequencing of SaPTAL-a-pET28a | SaPTAL-a-pET28a | pHM1834 |
| TTGATGGCAGGAAGGTGGAT | sequencing of SaPTAL-b-pET28a | SaPTAL-b-pET28a | pHM1835 |
| ATCGGAAAGCTCATGTTCGC | sequencing of SaPTAL-b-pET28a | SaPTAL-b-pET28a | pHM1836 |
| CCCCAAGGAAGGTCTGGC | sequencing of SbPTAL-pET28a | SbPTAL-pET28a | pHM2015 |
| ACATCGGCAAGCTCATGTTC | sequencing of SbPTAL-pET28a | SbPTAL-pET28a | pHM2016 |
| CATCGTCAATGGCACCTCC | sequencing of BdPTAL-pET28a | BdPTAL$^{H123F}$-pET28a | pHM2026 |
| CTCATGTTCGCGCAGTTCTC | sequencing of BdPTAL-pET28a | BdPTAL$^{H123F}$-pET28a | pHM2027 |
| GTCTCGCCATGGTCAACG | sequencing of SbPAL-pET28a | SbPAL-pET28a | pHM2070 |
| CCATCGGCAAGCTCATGTTC | sequencing of SbPAL-pET28a | SbPAL-pET28a | pHM2071 |
| CCTTGCCATGGTGAACGG | sequencing of BdPAL-pET28a | BdPAL-pET28a | pHM2076 |
| CAAGCTCATGTTTGCCCAGT | sequencing of BdPAL-pET28a | BdPAL-pET28a | pHM2077 |
| **Site-directed mutagenesis (1)** | | | |
| CTCAGGTTTCTGAACGCCGGGATCTTC | site-directed mutagenesis (H123F) | BdPTAL-pET28a | pHM1894 |
| GTTCAGAAACCTGAGGAGCTCGACCTG | site-directed mutagenesis (H123F) | BdPTAL-pET28a | pHM1895 |
| CTTAGATTCCTCAATGCCGGAATCTT | site-directed mutagenesis (F140H) | JaPTAL-pET28a | pHM1896 |
| ATTGAGGAATCTAAGGAGCTCTATTTG | site-directed mutagenesis (F140H) | JaPTAL-pET28a | pHM1897 |
| AATTAGACACCTCAATGCCGGAGTCTT | site-directed mutagenesis (H128F) | JaPAL-pET28a | pHM1904 |
| TTGAGGTGTCTAATTAGCTCTCTTTGG | site-directed mutagenesis (H128F) | JaPAL-pET28a | pHM1905 |
| CTCCGGTTTCTGAATGCTGGAATCTT | site-directed mutagenesis (H118F) | SaPTAL-a-pET28a | pHM1900 |
| ATTCAGAAACCGGAGGAGCTCCACCTG | site-directed mutagenesis (H118F) | SaPTAL-a-pET28a | pHM1901 |
| CTTCGGTTTCTCAATGCCGGAATCTT | site-directed mutagenesis (H127F) | SaPTAL-b-pET28a | pHM1902 |
| ATTGAGAAACCGAAGGAGCTCCACCTG | site-directed mutagenesis (H127F) | SaPTAL-b-pET28a | pHM1903 |
| CTCAGGTTTCTCAACGCCGGGATCTTCGGCACC | site-directed mutagenesis (H125F) | SbPTAL-pET28a | pHM2013 |
| GTTGAGAAACCTGAGCAGCTCGACCTGGAGCGC | site-directed mutagenesis (H125F) | SbPTAL-pET28a | pHM2014 |
| ATCAGACACCTCAATGCCGGCGCCTTCGGCACC | site-directed mutagenesis (F135H) | SbPAL-pET28a | pHM2083 |
| ATTGAGGTGTCTGATGAGCTCCCTCTGGAGCGCG | site-directed mutagenesis (F135H) | SbPAL-pET28a | pHM2084 |
| ATCCGACACCTTAATGCGGGAGCCTTCGGCACC | site-directed mutagenesis (F138H) | BdPAL-pET28a | pHM2085 |
| ATTAAGGTGTCGGATGAGCTCTCTCTGCAGAGCGC | site-directed mutagenesis (F138H) | BdPAL-pET28a | pHM2086 |
| CTTAGATTCCTCAATGCCGGAGTCTTCGGCACC | site-directed mutagenesis (H140F) | JaPAL$^{F140H_MUT8}$-pET28a, JaPAL$^{F140H_MUT8}$ -pET28a | pHM2232 |
| ATTGAGGAATCTAAGTAGCTCTCTTTGGAGAGC | site-directed mutagenesis (H140F) | JaPAL$^{F140H_MUT8}$-pET28a | pHM2233 |
| ATTGAGGAATCTAAGTAGCTCTACTTGGAGAGC | site-directed mutagenesis (H140F) | JaPAL$^{F140H_MUT16}$-pET28a | pHM2234 |
| **Site-directed mutagenesis (2)** | | | |
| GCGACTGGGTCATGAGCAGCATGATGAACGGC | site-directed mutagenesis (I102V) | JaPAL$^{F140H_MUT8}$-pET28a | pHM2354 |
| TCATGACCCAGTCGCTGCTGGCCTTGACG | site-directed mutagenesis (I102V) | JaPAL$^{F140H_MUT8}$-pET28a | pHM2355 |
| ACCGACAGCTACGGTGTCACCACTGG | site-directed mutagenesis (I112S) | JaPAL$^{F140H_MUT8}$-pET28a | pHM2328 |
| ACCGTAGCTGTCGGTGCCGTTCATCA | site-directed mutagenesis (I112S) | JaPAL$^{F140H_MUT8}$-pET28a | pHM2329 |
| CTTTGGAGCCACCTCCCACAGGAGGACC | site-directed mutagenesis (G121A) | JaPAL$^{F140H_MUT8}$-pET28a | pHM2356 |
| GAGGTGGCTCCAAAGCCAGTGGTGACACC | site-directed mutagenesis (G121A) | JaPAL$^{F140H_MUT8}$-pET28a | pHM2357 |
| GAGAGCTAATTAGACACCTCAATGCCGGAGTC | site-directed mutagenesis (L138I) | JaPAL$^{F140H_MUT8}$-pET28a | pHM2385 |
| GTCTAATTAGCTCTCTTTGGAGAGCACCAC | site-directed mutagenesis (L138I) | JaPAL$^{F140H_MUT8}$-pET28a | pHM2386 |
| CGGCACGGCCGTGGGTTCTGGTCTTG | site-directed mutagenesis (S267A) | JaPAL$^{F140H_MUT8}$-pET28a | pHM2334 |
| CCCACGGCCGTGCCGTTCACCATGGC | site-directed mutagenesis (S267A) | JaPAL$^{F140H_MUT8}$-pET28a | pHM2335 |
| TGGCCTGCCTTCCAACCTGGCCGGTG | site-directed mutagenesis (T444P) | JaPAL$^{F140H_MUT8}$-pET28a | pHM2336 |
| TTGGAAGGCAGGCCATTGTTGTAGAAG | site-directed mutagenesis (T444P) | JaPAL$^{F140H_MUT8}$-pET28a | pHM2337 |
| CAACCTGTCCGGTGGGCGCAACCCGA | site-directed mutagenesis (A448S) | JaPAL$^{F140H_MUT8}$-pET28a | pHM2338 |
| CCACCGGACAGGTTGGAAGTCAGGCC | site-directed mutagenesis (A448S) | JaPAL$^{F140H_MUT8}$-pET28a | pHM2339 |

| | | | |
|---|---|---|---|
| TGGCCTTATCTCATCCAGGAAGACCG | site-directed mutagenesis (V500I) | JaPAL$^{F140H_MUT8}$-pET28a | pHM2340 |
| GATGAGATAAGGCCAAGCGAGTTGAC | site-directed mutagenesis (V500I) | JaPAL$^{F140H_MUT8}$-pET28a | pHM2341 |
| **Site-directed mutagenesis (3)** | | | |
| GGAGATAGCTATGGTGTCACCACTGGCTTCG | site-directed mutagenesis (I97S) | JaPTAL$^{H128F}$-pET28a | pHM2456 |
| ACCATAGCTATCTCCACCGTTCGCCACG | site-directed mutagenesis (I97S) | JaPTAL$^{H128F}$-pET28a | pHM2457 |
| ACCGACATATACGGTGTCACCACTGGCT | site-directed mutagenesis (S112I) | JaPAL$^{F140H}$-pET28a | pHM2458 |
| ACCGTATATGTCGGTGCCGTTCATCA | site-directed mutagenesis (S112I) | JaPAL$^{F140H}$-pET28a | pHM2459 |
| ACTGATATATATGGTGTTACTACTGGTTTTGGTG | site-directed mutagenesis (S116I) | AtPAL1-pET28a | pHM2524 |
| ACCATATATATCAGTGCCTTTGTTCATACTCTC | site-directed mutagenesis (S116I) | AtPAL1-pET28a | pHM2525 |
| TATTAGACACCTTAACGCCGGAATATTCG | site-directed mutagenesis (F144H) | AtPAL1-pET28a | pHM2526 |
| TTAAGGTGTCTAATAAGTTCCTTCTGAAGTGCG | site-directed mutagenesis (F144H) | AtPAL1-pET28a | pHM2527 |

**Table S16. Key resource information in this study**

| REAGENT or RESOURCE | SOURCE | IDENTIFIER |
|---|---|---|
| **Chemicals, peptides, and recombinant proteins** | | |
| JaPTAL | Phytozome v13 | Joasc.05G060400 |
| JaPAL | Phytozome v13 | Joasc.05G060500 |
| SbPTAL | Phytozome v13 | SbiRTX430.04G229900.1 |
| SbPAL | Phytozome v13 | SbiRTX430.04G230000.1 |
| BdPTAL | Phytozome v13 | Bradi3g49250.2 |
| BdPAL | Phytozome v13 | Bradi5g15830.1 |
| AtPAL1 | Phytozome v13 | AT2G37040.1 |
| **Deposited data** | | |
| Genome of *Joinvillea ascendens* | Joint Genome Institute (JGI) Open Green Genome Initiative (OGG) https://phytozome-next.jgi.doe.gov/ogg | https://phytozome-next.jgi.doe.gov/ogg/info/Jascendens_v1_1 |
| Genome of *Ecdeiocolea monostachya* accession E001 | DDBJ/ENA/GenBank | BioProject codes PRJNA894727 (Illumina/Oxford Nanopore Technology hybrid assembly) and PRJNA1179411 (PacBio HiFi assembly) |
| Genome of *Pharus latifolius* | Joint Genome Institute (JGI) Open Green Genome Initiative (OGG) https://phytozome-next.jgi.doe.gov/ogg | https://phytozome-next.jgi.doe.gov/info/Platifolius_v1_1 |
| Genome of *Typha latifolia* | Joint Genome Institute (JGI) Open Green Genome Initiative (OGG) https://phytozome-next.jgi.doe.gov/ogg | https://phytozome-next.jgi.doe.gov/info/Tlatifoliavar_2019_6_v1_1 |
| **Experimental models:** | | |
| *Joinvillea ascendens* subsp. glabra | The National Tropical Garden (Kalaheo, HI), from seed collected in New Caledonia. | Accession No. 800379 |
| *Ecdeiocolea monostachya* | Western Australia near Jurien Bay | Accession No. E001 |
| *Pharus latifolius* | Missouri Botanical Garden (St. | Accession No. 1993-0885-2 |

| | | |
|---|---|---|
| | Louis, MO) | |
| *Typha latifolia* | Penn State Arboretum (University Park, PA) | Accession No. CWD-2019.6 |
| **Software and algorithms** | | |
| ape 5.0 | Paradis and Schliep (*107*) | http://cran.r-project.org/package=ape |
| Augustus | Keller et al. (*82*) | https://github.com/Gaius-Augustus/Augustus |
| BLAST | Camacho et al. | https://www.ncbi.nlm.nih.gov |
| BLAT | Kent (*98*) | http://genome.ucsc.edu/cgi-bin/hgBlat |
| BUSCO | Manni et al. (*92*) | https://busco.ezlab.org/ |
| bwa-mem | Li (*99*) | https://github.com/lh3/bwa |
| bwa-mem2 | Li (*99*) | https://github.com/bwa-mem2/bwa-mem2 |
| COGE | Lyons and Freeling (*108*) | https://genomevolution.org/coge/ |
| Cufflinks/Cuffmerge | Trapnell et al. | http://cole-trapnell-lab.github.io/cufflinks/ |
| EVM | Haas et al. (*91*) | https://github.com/EVidenceModeler/EVidenceModeler/wiki |
| findGSE | Sun et al. (*77*) | https://github.com/schneebergerlab/findGSE |
| GATK | Van der Auwera and O'Connor (*100*) | https://gatk.broadinstitute.org |
| GeMoMa | Keilwagen et al. (*84*) | https://github.com/Jstacs/Jstacs |
| GeneMarkS-T | Tang et al. (*88*) | https://genemark.bme.gatech.edu/ |
| GENESPACE | Lovell et al. (*35*) | https://github.com/jtlovell/GENESPACE |
| GenomeScope | Vurture et al. (*76*) | http://genomescope.org/ |
| GSNAP | Wu et al. (*74*) | http://research-pub.gene.com/gmap/ |
| HiFiAsm | Cheng et al. (*79,80*) | https://github.com/chhylp123/hifiasm |
| hisat2 | Kim et al. (*85*) | http://daehwankimlab.github.io/hisa |

| | | |
|---|---|---|
| | | t2/ |
| IQ-TREE 2 | Minh et al. (*103*) | http://www.iqtree.org |
| jellyfish | Marcais and Kingsford | https://github.com/gmarcais/Jellyfish |
| Juicer | Durand et al. (*96*) | https://github.com/aidenlab/juicer |
| KmerGenie | Chikhi and Medvedev (*78*) | http://kmergenie.bx.psu.edu/ |
| *k*-mer analysis toolkit | Mapleson et al. | https://github.com/TGAC/KAT |
| kraken2 | Wood et al. (*86*) | https://github.com/DerrickWood/kraken2 |
| MAFFT | Katoh and Standley (*102*) | https://mafft.cbrc.jp/ |
| MAKER | Campbell et al. (*72*) | https://github.com/Yandell-Lab/maker |
| MaSuRCA | Zimin et al. | https://github.com/alekseyzimin/masurca |
| MECAT | Xiao et al. (*71*) | https://thub.com/xiaochuanle/MECAT |
| ModelTest-NG | Darriba et al. (*106*) | https://github.com/ddarriba/mgiodeltest |
| OrthoFinder | Emms and Kelly (*101*) | https://github.com/davidemms/OrthoFinder |
| PAL2NAL | Suyama et al. (*104*) | https://www.bork.embl.de/pal2nal/ |
| PAML | Yang (*58*) | https://github.com/abacus-gene/paml |
| PASA | Haas et al. (*90*) | https://github.com/PASApipeline/PASApipeline/ |
| PERTRAN | Shu et al. (*73*) | N/A |
| PUG | McKain et al. (*3*) | https://github.com/mrmckain/PUG |
| purge_dups | Guan et al. (*81*) | https://github.com/dfguan/purge_dups |
| R | R Development Core Team, 2020 | www.R-project.org/ |
| RACON | Vaser et al. (*34*) | https://github.com/isovic/racon |
| RAxML-NG | Kozlov et al. (*105*) | https://github.com/amkozlov/raxml-ng |

| | | |
|---|---|---|
| RepeatModeler | Smit and Hubley | https://github.com/Dfam-consortium/RepeatModeler |
| samtools | Danece et al. | https://github.com/samtools/samtools |
| Solver | Microsoft Office Excel | N/A |
| SNAP | Korf (*83*) | https://github.com/KorfLab/SNAP |
| StringTie | Pertea et al. (*78*) | https://github.com/gpertea/stringtie |
| SWISS-MODEL | Waterhouse et al. (*109*) | https://swissmodel.expasy.org/ |
| TransDecoder | N/A | https://github.com/TransDecoder/TransDecoder |
| TreeSolve | Kordi et al. (*54*) | https://compbio.engr.uconn.edu/software/TreeSolve/ |
| Trimmomatic | Bolger et al. (*94*) | http://www.usadellab.org/cms/?page=trimmomatic |
| Trinity | Grabherr et al. (*89*) | https://github.com/trinityrnaseq/trinityrnaseq/wiki |
| 3D-DNA | Dudchenko et al. (*97*) | https://github.com/aidenlab/3d-dna |

### References for Supplementary Information:


70. J. Sebastian, M. K. Wong, E. Tang, J. R. Dinneny, Methods to Promote Germination of Dormant Setaria viridis Seeds. *PLOS ONE* **9**, e95109 (2014).

71. C.-L. Xiao, Y. Chen, S.-Q. Xie, K.-N. Chen, Y. Wang, Y. Han, F. Luo, Z. Xie, MECAT: fast mapping, error correction, and de novo assembly for single-molecule sequencing reads. *Nat. Methods* **14**, 1072–1074 (2017).

72. M. S. Campbell, C. Holt, B. Moore, M. Yandell, Genome Annotation and Curation Using MAKER and MAKER-P. *Current Protocols in Bioinformatics* **48**, 4.11.1-4.11.39 (2014).

73. S. Shu, D. Goodstein, D. Rokhsar, "PERTRAN: Genome-guided RNA-seq Read Assembler" (2013; https://www.semanticscholar.org/paper/PERTRAN%3A-Genome-guided-RNA-seq-Read-Assembler-Shu-Goodstein/d50df620c234841872bd568d92f555f3d45326d1#citing-papers).

74. T. D. Wu, S. Nacu, Fast and SNP-tolerant detection of complex variants and splicing in short reads. *Bioinformatics* **26**, 873–881 (2010).

75. C. N. Stewart, L. E. Via, A rapid CTAB DNA isolation technique useful for RAPD fingerprinting and other PCR applications. *Biotechniques* **14**, 748–750 (1993).

76. G. W. Vurture, F. J. Sedlazeck, M. Nattestad, C. J. Underwood, H. Fang, J. Gurtowski, M. C. Schatz, GenomeScope: fast reference-free genome profiling from short reads. *Bioinformatics* **33**, 2202–2204 (2017).

77. H. Sun, J. Ding, M. Piednoël, K. Schneeberger, findGSE: estimating genome size variation within human and Arabidopsis using k-mer frequencies. *Bioinformatics* **34**, 550–557 (2018).

78. R. Chikhi, P. Medvedev, Informed and Automated k-Mer Size Selection for Genome Assembly, *arXiv.org* (2013). https://arxiv.org/abs/1304.5665v1.

79. H. Cheng, G. T. Concepcion, X. Feng, H. Zhang, H. Li, Haplotype-resolved de novo assembly using phased assembly graphs with hifiasm. *Nat. Methods* **18**, 170–175 (2021).

80. H. Cheng, E. D. Jarvis, O. Fedrigo, K.-P. Koepfli, L. Urban, N. J. Gemmell, H. Li, Haplotype-resolved assembly of diploid genomes without parental data. *Nat. Biotechnol.* **40**, 1332–1335 (2022).

81. D. Guan, S. A. McCarthy, J. Wood, K. Howe, Y. Wang, R. Durbin, Identifying and removing haplotypic duplication in primary genome assemblies. *Bioinformatics* **36**, 2896–2898 (2020).

82. O. Keller, M. Kollmar, M. Stanke, S. Waack, A novel hybrid gene prediction method employing protein multiple sequence alignments. *Bioinformatics* **27**, 757–763 (2011).

83. I. Korf, Gene finding in novel genomes. *BMC Bioinformatics* **5**, 59 (2004).

84. J. Keilwagen, F. Hartung, J. Grau, "GeMoMa: Homology-Based Gene Prediction Utilizing Intron Position Conservation and RNA-seq Data" in *Gene Prediction: Methods and Protocols*, M. Kollmar, Ed. (Springer, New York, NY, 2019; https://doi.org/10.1007/978-1-4939-9173-0_9), pp. 161–177.

85. D. Kim, J. M. Paggi, C. Park, C. Bennett, S. L. Salzberg, Graph-based genome alignment and genotyping with HISAT2 and HISAT-genotype. *Nat. Biotechnol* **37**, 907–915 (2019).

86. D. E. Wood, J. Lu, B. Langmead, Improved metagenomic analysis with Kraken 2. *Genome Biol.* **20**, 257 (2019).

87. M. Pertea, G. M. Pertea, C. M. Antonescu, T.-C. Chang, J. T. Mendell, S. L. Salzberg, StringTie enables improved reconstruction of a transcriptome from RNA-seq reads. *Nat. Biotechnol.* **33**, 290–295 (2015).

88. S. Tang, A. Lomsadze, M. Borodovsky, Identification of protein coding regions in RNA transcripts. *Nucl, Acids Res.* **43**, e78 (2015).

89. M. G. Grabherr, B. J. Haas, M. Yassour, J. Z. Levin, D. A. Thompson, I. Amit, X. Adiconis, L. Fan, R. Raychowdhury, Q. Zeng, Z. Chen, E. Mauceli, N. Hacohen, A. Gnirke, N. Rhind, F. di Palma, B. W. Birren, C. Nusbaum, K. Lindblad-Toh, N. Friedman, A. Regev, Full-length transcriptome assembly from RNA-Seq data without a reference genome. *Nat. Biotechnol.* **29**, 644–652 (2011).

90. B. J. Haas, A. L. Delcher, S. M. Mount, J. R. Wortman, R. K. Smith Jr, L. I. Hannick, R. Maiti, C. M. Ronning, D. B. Rusch, C. D. Town, S. L. Salzberg, O. White, Improving the Arabidopsis genome annotation using maximal transcript alignment assemblies. *Nucl. Acids Res.* **31**, 5654–5666 (2003).

91. B. J. Haas, S. L. Salzberg, W. Zhu, M. Pertea, J. E. Allen, J. Orvis, O. White, C. R. Buell, J. R. Wortman, Automated eukaryotic gene structure annotation using EVidenceModeler and the Program to Assemble Spliced Alignments. *Genome Biol.* **9**, R7 (2008).

92. M. Manni, M. R. Berkeley, M. Seppey, F. A. Simão, E. M. Zdobnov, BUSCO Update: Novel and Streamlined Workflows along with Broader and Deeper Phylogenetic Coverage for Scoring of Eukaryotic, Prokaryotic, and Viral Genomes. *Molecular Biology and Evolution* **38**, 4647–4654 (2021).

93. J. Salinas, G. Matassi, L. M. Montero, G. Bernardi, Compositional compartmentalization and compositional patterns in the nuclear genomes of plants. *Nucl. Acids Res.* **16**, 4269–4285 (1988).

94. A. M. Bolger, M. Lohse, B. Usadel, Trimmomatic: a flexible trimmer for Illumina sequence data. *Bioinformatics* **30**, 2114–2120 (2014).

95. R. Chikhi, P. Medvedev, Informed and Automated k-Mer Size Selection for Genome Assembly, *arXiv.org* (2013). https://arxiv.org/abs/1304.5665v1.

96. N. C. Durand, M. S. Shamim, I. Machol, S. S. P. Rao, M. H. Huntley, E. S. Lander, E. L. Aiden, Juicer Provides a One-Click System for Analyzing Loop-Resolution Hi-C Experiments. *Cells* **3**, 95–98 (2016).

97. O. Dudchenko, S. S. Batra, A. D. Omer, S. K. Nyquist, M. Hoeger, N. C. Durand, M. S. Shamim, I. Machol, E. S. Lander, A. P. Aiden, E. L. Aiden, De novo assembly of the Aedes aegypti genome using Hi-C yields chromosome-length scaffolds. *Science* **356**, 92–95 (2017).

98. W. J. Kent, BLAT—The BLAST-Like Alignment Tool. *Genome Res.* **12**, 656–664 (2002).

99. H. Li, Aligning sequence reads, clone sequences and assembly contigs with BWA-MEM. arXiv arXiv:1303.3997 [Preprint] (2013). https://doi.org/10.48550/arXiv.1303.3997.

100. G. van der Auwera, B. D. O'Connor, *Genomics in the Cloud: Using Docker, GATK, and WDL in Terra* (O'Reilly Media, Incorporated, 2020).

101. D. M. Emms, S. Kelly, OrthoFinder: solving fundamental biases in whole genome comparisons dramatically improves orthogroup inference accuracy. *Genome Biol.* **16**, 157 (2015).

102. K. Katoh, D. M. Standley, MAFFT multiple sequence alignment software version 7: improvements in performance and usability. *Mol. Biol. Evol.* **30**, 772–780 (2013).

103. B. Q. Minh, H. A. Schmidt, O. Chernomor, D. Schrempf, M. D. Woodhams, A. von Haeseler, R. Lanfear, IQ-TREE 2: New Models and Efficient Methods for Phylogenetic Inference in the Genomic Era. *Mol. Biol. Evol.* **37**, 1530–1534 (2020).

104. M. Suyama, D. Torrents, P. Bork, PAL2NAL: robust conversion of protein sequence alignments into the corresponding codon alignments. *Nucl. Acids Res.* **34**, W609–W612 (2006).

105. A. M. Kozlov, D. Darriba, T. Flouri, B. Morel, A. Stamatakis, RAxML-NG: a fast, scalable and user-friendly tool for maximum likelihood phylogenetic inference. *Bioinformatics* **35**, 4453–4455 (2019).

106. D. Darriba, D. Posada, A. M. Kozlov, A. Stamatakis, B. Morel, T. Flouri, ModelTest-NG: A New and Scalable Tool for the Selection of DNA and Protein Evolutionary Models. *Mol. Biol. Evol.* **37**, 291–294 (2020).

107. E. Paradis, K. Schliep, ape 5.0: an environment for modern phylogenetics and evolutionary analyses in R. *Bioinformatics* **35**, 526–528 (2019).

108. E. Lyons, M. Freeling, How to usefully compare homologous plant genes and chromosomes as DNA sequences. *Plant J.* **53**, 661–673 (2008).

109. A. Waterhouse, M. Bertoni, S. Bienert, G. Studer, G. Tauriello, R. Gumienny, F. T. Heer, T. A. P. de Beer, C. Rempfer, L. Bordoli, R. Lepore, T. Schwede, SWISS-MODEL: homology modelling of protein structures and complexes. *Mol. Biol. Evol.* **46**, W296–W303 (2018).

110. AMBORELLA GENOME PROJECT, V. A. Albert, W. B. Barbazuk, C. W. dePamphilis, J. P. Der, J. Leebens-Mack, H. Ma, J. D. Palmer, S. Rounsley, D. Sankoff, S. C. Schuster, D. E. Soltis, P. S. Soltis, S. R. Wessler, R. A. Wing, V. A. Albert, J. S. S. Ammiraju, W. B. Barbazuk, S. Chamala, A. S. Chanderbali, C. W. dePamphilis, J. P. Der, R. Determann, J. Leebens-Mack, H. Ma, P. Ralph, S. Rounsley, S. C. Schuster, D. E. Soltis, P. S. Soltis, J. Talag, L. Tomsho, B. Walts, S. Wanke, R. A. Wing, V. A. Albert, W. B. Barbazuk, S. Chamala, A. S. Chanderbali, T.-H. Chang, R. Determann, T. Lan, D. E. Soltis, P. S. Soltis, S. Arikit, M. J. Axtell, S. Ayyampalayam, W. B. Barbazuk, J. M. Burnette, S. Chamala, E. De Paoli, C. W. dePamphilis, J. P. Der, J. C. Estill, N. P. Farrell, A. Harkess, Y. Jiao, J. Leebens-Mack, K. Liu, W. Mei, B. C. Meyers, S. Shahid, E. Wafula, B. Walts, S. R. Wessler, J. Zhai, X. Zhang, V. A. Albert, L. Carretero-Paulet, C. W. dePamphilis, J. P. Der, Y. Jiao, J. Leebens-Mack, E. Lyons, D. Sankoff, H. Tang, E. Wafula, C. Zheng, V. A. Albert, N. S. Altman, W. B. Barbazuk, L. Carretero-Paulet, C. W. dePamphilis, J. P. Der, J. C. Estill, Y. Jiao, J. Leebens-Mack, K. Liu, W. Mei, E. Wafula, N. S. Altman, S. Arikit, M. J. Axtell, S. Chamala, A. S. Chanderbali, F. Chen, J.-Q. Chen, V. Chiang, E. De Paoli, C. W. dePamphilis, J. P. Der, R. Determann, B. Fogliani, C. Guo, J. Harholt, A. Harkess, C. Job, D. Job, S. Kim, H. Kong, J. Leebens-Mack, G. Li, L. Li, J. Liu, H. Ma, B. C. Meyers, J. Park, X. Qi, L. Rajjou, V. Burtet-Sarramegna, R. Sederoff, S. Shahid, D. E. Soltis, P. S. Soltis, Y.-H. Sun, P. Ulvskov, M. Villegente, J.-Y. Xue, T.-F. Yeh, X. Yu, J. Zhai, J. J. Acosta, V. A. Albert, W. B. Barbazuk, R. A. Bruenn, S. Chamala, A. de Kochko, C. W. dePamphilis, J. P. Der, L. R. Herrera-Estrella, E. Ibarra-Laclette, M. Kirst, J. Leebens-Mack, S. P. Pissis, V. Poncet, S. C. Schuster, D. E. Soltis, P. S. Soltis, L. Tomsho, The Amborella Genome and the Evolution of Flowering Plants. *Science* **342**, 1241089 (2013).

111. L. Zhang, F. Chen, X. Zhang, Z. Li, Y. Zhao, R. Lohaus, X. Chang, W. Dong, S. Y. W. Ho, X. Liu, A. Song, J. Chen, W. Guo, Z. Wang, Y. Zhuang, H. Wang, X. Chen, J. Hu, Y. Liu, Y. Qin, K. Wang, S. Dong, Y. Liu, S. Zhang, X. Yu, Q. Wu, L. Wang, X. Yan, Y. Jiao, H. Kong, X. Zhou, C. Yu, Y. Chen, F. Li, J. Wang, W. Chen, X. Chen, Q. Jia, C. Zhang, Y. Jiang, W. Zhang, G. Liu, J. Fu, F. Chen, H. Ma, Y. Van de Peer, H. Tang, The water lily genome and the early evolution of flowering plants. *Nature* **577**, 79–84 (2020).

112. S.-M. Chaw, Y.-C. Liu, Y.-W. Wu, H.-Y. Wang, C.-Y. I. Lin, C.-S. Wu, H.-M. Ke, L.-Y. Chang, C.-Y. Hsu, H.-T. Yang, E. Sudianto, M.-H. Hsu, K.-P. Wu, L.-N. Wang, J. H. Leebens-Mack, I. J. Tsai, Stout camphor tree genome fills gaps in understanding of flowering plant genome evolution. *Nature Plants* **5**, 63–73 (2019).

113. D. L. Filiault, E. S. Ballerini, T. Mandáková, G. Aköz, N. J. Derieg, J. Schmutz, J. Jenkins, J. Grimwood, S. Shu, R. D. Hayes, U. Hellsten, K. Barry, J. Yan, S. Mihaltcheva, M. Karafiátová, V. Nizhynska, E. M. Kramer, M. A. Lysak, S. A. Hodges, M. Nordborg, The Aquilegia genome provides insight into adaptive radiation and reveals an extraordinarily polymorphic chromosome with a unique history. *Elife* **7**, e36426 (2018).

114. M. Iorizzo, S. Ellison, D. Senalik, P. Zeng, P. Satapoomin, J. Huang, M. Bowman, M. Iovene, W. Sanseverino, P. Cavagnaro, M. Yildiz, A. Macko-Podgórni, E. Moranska, E. Grzebelus, D. Grzebelus, H. Ashrafi, Z. Zheng, S. Cheng, D. Spooner, A. Van Deynze, P. Simon, A high-quality carrot genome assembly provides new insights into carotenoid accumulation and asterid genome evolution. *Nat Genet* **48**, 657–666 (2016).

115. H. Badouin, J. Gouzy, C. J. Grassa, F. Murat, S. E. Staton, L. Cottret, C. Lelandais-Brière, G. L. Owens, S. Carrère, B. Mayjonade, L. Legrand, N. Gill, N. C. Kane, J. E. Bowers, S. Hubner, A. Bellec, A. Bérard, H. Bergès, N. Blanchet, M.-C. Boniface, D. Brunel, O. Catrice, N. Chaidir, C. Claudel, C. Donnadieu, T. Faraut, G. Fievet, N. Helmstetter, M. King, S. J. Knapp, Z. Lai, M.-C. Le Paslier, Y. Lippi, L. Lorenzon, J. R. Mandel, G. Marage, G. Marchand, E. Marquand, E. Bret-Mestries, E. Morien, S. Nambeesan, T. Nguyen, P. Pegot-Espagnet, N. Pouilly, F. Raftis, E. Sallet, T. Schiex, J. Thomas, C. Vandecasteele, D. Varès, F. Vear, S. Vautrin, M. Crespi, B. Mangin, J. M. Burke, J. Salse, S. Muños, P. Vincourt, L. H. Rieseberg, N. B. Langlade, The sunflower genome provides insights into oil metabolism, flowering and Asterid evolution. *Nature* **546**, 148–152 (2017).

116. Y. Zhou, Z. Zhang, Z. Bao, H. Li, Y. Lyu, Y. Zan, Y. Wu, L. Cheng, Y. Fang, K. Wu, J. Zhang, H. Lyu, T. Lin, Q. Gao, S. Saha, L. Mueller, Z. Fei, T. Städler, S. Xu, Z. Zhang, D. Speed, S. Huang, Graph pangenome captures missing heritability and empowers tomato breeding. *Nature* **606**, 527–534 (2022).

117. A. M. Hulse-Kemp, H. Bostan, S. Chen, H. Ashrafi, K. Stoffel, W. Sanseverino, L. Li, S. Cheng, M. C. Schatz, T. Garvin, L. J. du Toit, E. Tseng, J. Chin, M. Iorizzo, A. Van Deynze, An anchored chromosome-scale genome assembly of spinach improves annotation and reveals extensive gene rearrangements in euasterids. *The Plant Genome* **14**, e20101 (2021).

118. J. M. McGrath, A. Funk, P. Galewski, S. Ou, B. Townsend, K. Davenport, H. Daligault, S. Johnson, J. Lee, A. Hastie, A. Darracq, G. Willems, S. Barnes, I. Liachko, S. Sullivan, S. Koren, A. Phillippy, J. Wang, T. Liu, J. Pulman, K. Childs, S. Shu, A. Yocum, D. Fermin, E. Mutasa-Göttgens, P. Stevanato, K. Taguchi, R. Naegele, K. M. Dorn, A contiguous de novo genome assembly of sugar beet EL10 (Beta vulgaris L.). *DNA Research* **30**, dsac033 (2023).

119. J. Schmutz, S. B. Cannon, J. Schlueter, J. Ma, T. Mitros, W. Nelson, D. L. Hyten, Q. Song, J. J. Thelen, J. Cheng, D. Xu, U. Hellsten, G. D. May, Y. Yu, T. Sakurai, T. Umezawa, M. K. Bhattacharyya, D. Sandhu, B. Valliyodan, E. Lindquist, M. Peto, D. Grant, S. Shu, D. Goodstein, K. Barry, M. Futrell-Griggs, B. Abernathy, J. Du, Z. Tian, L. Zhu, N. Gill, T. Joshi, M. Libault, A. Sethuraman, X.-C. Zhang, K. Shinozaki, H. T. Nguyen, R. A. Wing, P. Cregan, J. Specht, J. Grimwood, D. Rokhsar, G. Stacey, R. C. Shoemaker, S. A. Jackson, Genome sequence of the palaeopolyploid soybean. *Nature* **463**, 178–183 (2010).

120. P. Lamesch, T. Z. Berardini, D. Li, D. Swarbreck, C. Wilks, R. Sasidharan, R. Muller, K. Dreher, D. L. Alexander, M. Garcia-Hernandez, A. S. Karthikeyan, C. H. Lee, W. D. Nelson, L. Ploetz, S. Singh, A. Wensel, E. Huala, The Arabidopsis Information Resource (TAIR): improved gene annotation and new tools. *Nucleic Acids Research* **40**, D1202–D1210 (2012).

121. G. A. Tuskan, S. DiFazio, S. Jansson, J. Bohlmann, I. Grigoriev, U. Hellsten, N. Putnam, S. Ralph, S. Rombauts, A. Salamov, J. Schein, L. Sterck, A. Aerts, R. R. Bhalerao, R. P. Bhalerao, D.

Blaudez, W. Boerjan, A. Brun, A. Brunner, V. Busov, M. Campbell, J. Carlson, M. Chalot, J. Chapman, G.-L. Chen, D. Cooper, P. M. Coutinho, J. Couturier, S. Covert, Q. Cronk, R. Cunningham, J. Davis, S. Degroeve, A. Dejardin, C. dePamphilis, J. Detter, B. Dirks, I. Dubchak, S. Duplessis, J. Ehlting, B. Ellis, K. Gendler, D. Goodstein, M. Gribskov, J. Grimwood, A. Groover, L. Gunter, B. Hamberger, B. Heinze, Y. Helariutta, B. Henrissat, D. Holligan, R. Holt, W. Huang, N. Islam-Faridi, S. Jones, M. Jones-Rhoades, R. Jorgensen, C. Joshi, J. Kangasjarvi, J. Karlsson, C. Kelleher, R. Kirkpatrick, M. Kirst, A. Kohler, U. Kalluri, F. Larimer, J. Leebens-Mack, J.-C. Leple, P. Locascio, Y. Lou, S. Lucas, F. Martin, B. Montanini, C. Napoli, D. R. Nelson, C. Nelson, K. Nieminen, O. Nilsson, V. Pereda, G. Peter, R. Philippe, G. Pilate, A. Poliakov, J. Razumovskaya, P. Richardson, C. Rinaldi, K. Ritland, P. Rouze, D. Ryaboy, J. Schmutz, J. Schrader, B. Segerman, H. Shin, A. Siddiqui, F. Sterky, A. Terry, C.-J. Tsai, E. Uberbacher, P. Unneberg, J. Vahala, K. Wall, S. Wessler, G. Yang, T. Yin, C. Douglas, M. Marra, G. Sandberg, Y. Van de Peer, D. Rokhsar, The Genome of Black Cottonwood, Populus trichocarpa (Torr. & Gray). *Science* **313**, 1596–1604 (2006).

122. X. Ma, J. L. Olsen, T. B. H. Reusch, G. Procaccini, D. Kudrna, M. Williams, J. Grimwood, S. Rajasekar, J. Jenkins, J. Schmutz, Y. V. de Peer, Improved chromosome-level genome assembly and annotation of the seagrass, *Zostera marina* (eelgrass). F1000Research 10:289 [Preprint] (2021). https://doi.org/10.12688/f1000research.38156.1.

123. W. Wang, G. Haberer, H. Gundlach, C. Gläßer, T. Nussbaumer, M. C. Luo, A. Lomsadze, M. Borodovsky, R. A. Kerstetter, J. Shanklin, D. W. Byrant, T. C. Mockler, K. J. Appenroth, J. Grimwood, J. Jenkins, J. Chow, C. Choi, C. Adam, X.-H. Cao, J. Fuchs, I. Schubert, D. Rokhsar, J. Schmutz, T. P. Michael, K. F. X. Mayer, J. Messing, The Spirodela polyrhiza genome reveals insights into its neotenous reduction fast growth and aquatic lifestyle. *Nat Commun* **5**, 3311 (2014).

124. J. V. Bredeson, J. B. Lyons, I. O. Oniyinde, N. R. Okereke, O. Kolade, I. Nnabue, C. O. Nwadili, E. Hřibová, M. Parker, J. Nwogha, S. Shu, J. Carlson, R. Kariba, S. Muthemba, K. Knop, G. J. Barton, A. V. Sherwood, A. Lopez-Montes, R. Asiedu, R. Jamnadass, A. Muchugi, D. Goodstein, C. N. Egesi, J. Featherston, A. Asfaw, G. G. Simpson, J. Doležel, P. S. Hendre, A. Van Deynze, P. L. Kumar, J. E. Obidiegwu, R. Bhattacharjee, D. S. Rokhsar, Chromosome evolution and the genetic basis of agronomically important traits in greater yam. *Nat Commun* **13**, 2001 (2022).

125. A. Harkess, J. Zhou, C. Xu, J. E. Bowers, R. Van der Hulst, S. Ayyampalayam, F. Mercati, P. Riccardi, M. R. McKain, A. Kakrana, H. Tang, J. Ray, J. Groenendijk, S. Arikit, S. M. Mathioni, M. Nakano, H. Shan, A. Telgmann-Rauber, A. Kanno, Z. Yue, H. Chen, W. Li, Y. Chen, X. Xu, Y. Zhang, S. Luo, H. Chen, J. Gao, Z. Mao, J. C. Pires, M. Luo, D. Kudrna, R. A. Wing, B. C. Meyers, K. Yi, H. Kong, P. Lavrijsen, F. Sunseri, A. Falavigna, Y. Ye, J. H. Leebens-Mack, G. Chen, The asparagus genome sheds light on the origin and evolution of a young Y chromosome. *Nat Commun* **8**, 1279 (2017).

126. K. M. Hazzouri, M. Gros-Balthazard, J. M. Flowers, D. Copetti, A. Lemansour, M. Lebrun, K. Masmoudi, S. Ferrand, M. I. Dhar, Z. A. Fresquez, U. Rosas, J. Zhang, J. Talag, S. Lee, D. Kudrna, R. F. Powell, I. J. Leitch, R. R. Krueger, R. A. Wing, K. M. A. Amiri, M. D. Purugganan, Genome-wide association mapping of date palm fruit traits. *Nat Commun* **10**, 4680 (2019).

127. A. D'Hont, F. Denoeud, J.-M. Aury, F.-C. Baurens, F. Carreel, O. Garsmeur, B. Noel, S. Bocs, G. Droc, M. Rouard, C. Da Silva, K. Jabbari, C. Cardi, J. Poulain, M. Souquet, K. Labadie, C. Jourda, J. Lengellé, M. Rodier-Goud, A. Alberti, M. Bernard, M. Correa, S. Ayyampalayam, M. R. Mckain, J. Leebens-Mack, D. Burgess, M. Freeling, D. Mbéguié-A-Mbéguié, M. Chabannes, T. Wicker, O. Panaud, J. Barbosa, E. Hribova, P. Heslop-Harrison, R. Habas, R. Rivallan, P. Francois, C. Poiron, A. Kilian, D. Burthia, C. Jenny, F. Bakry, S. Brown, V. Guignon, G. Kema, M. Dita, C. Waalwijk, S. Joseph, A. Dievart, O. Jaillon, J. Leclercq, X. Argout, E. Lyons, A. Almeida, M. Jeridi, J.

Dolezel, N. Roux, A.-M. Risterucci, J. Weissenbach, M. Ruiz, J.-C. Glaszmann, F. Quétier, N. Yahiaoui, P. Wincker, The banana (Musa acuminata) genome and the evolution of monocotyledonous plants. *Nature* **488**, 213–217 (2012).

128. R. Ming, R. VanBuren, C. M. Wai, H. Tang, M. C. Schatz, J. E. Bowers, E. Lyons, M.-L. Wang, J. Chen, E. Biggers, J. Zhang, L. Huang, L. Zhang, W. Miao, J. Zhang, Z. Ye, C. Miao, Z. Lin, H. Wang, H. Zhou, W. C. Yim, H. D. Priest, C. Zheng, M. Woodhouse, P. P. Edger, R. Guyot, H.-B. Guo, H. Guo, G. Zheng, R. Singh, A. Sharma, X. Min, Y. Zheng, H. Lee, J. Gurtowski, F. J. Sedlazeck, A. Harkess, M. R. McKain, Z. Liao, J. Fang, J. Liu, X. Zhang, Q. Zhang, W. Hu, Y. Qin, K. Wang, L.-Y. Chen, N. Shirley, Y.-R. Lin, L.-Y. Liu, A. G. Hernandez, C. L. Wright, V. Bulone, G. A. Tuskan, K. Heath, F. Zee, P. H. Moore, R. Sunkar, J. H. Leebens-Mack, T. Mockler, J. L. Bennetzen, M. Freeling, D. Sankoff, A. H. Paterson, X. Zhu, X. Yang, J. A. C. Smith, J. C. Cushman, R. E. Paull, Q. Yu, The pineapple genome and the evolution of CAM photosynthesis. *Nat Genet* **47**, 1435–1442 (2015).

129. J. Planta, Y.-Y. Liang, H. Xin, M. T. Chansler, L. A. Prather, N. Jiang, J. Jiang, K. L. Childs, Chromosome-scale genome assemblies and annotations for Poales species Carex cristatella, Carex scoparia, Juncus effusus, and Juncus inflexus. *G3 Genes|Genomes|Genetics* **12**, jkac211 (2022).

130. S. Ouyang, W. Zhu, J. Hamilton, H. Lin, M. Campbell, K. Childs, F. Thibaud-Nissen, R. L. Malek, Y. Lee, L. Zheng, J. Orvis, B. Haas, J. Wortman, C. R. Buell, The TIGR Rice Genome Annotation Resource: improvements and new features. *Nucleic Acids Research* **35**, D883–D887 (2007).

131. J. P. Vogel, D. F. Garvin, T. C. Mockler, J. Schmutz, D. Rokhsar, M. W. Bevan, K. Barry, S. Lucas, M. Harmon-Smith, K. Lail, H. Tice, J. Schmutz (Leader), J. Grimwood, N. McKenzie, M. W. Bevan, N. Huo, Y. Q. Gu, G. R. Lazo, O. D. Anderson, J. P. Vogel (Leader), F. M. You, M.-C. Luo, J. Dvorak, J. Wright, M. Febrer, M. W. Bevan, D. Idziak, R. Hasterok, D. F. Garvin, E. Lindquist, M. Wang, S. E. Fox, H. D. Priest, S. A. Filichkin, S. A. Givan, D. W. Bryant, J. H. Chang, T. C. Mockler (Leader), H. Wu, W. Wu, A.-P. Hsia, P. S. Schnable, A. Kalyanaraman, B. Barbazuk, T. P. Michael, S. P. Hazen, J. N. Bragg, D. Laudencia-Chingcuanco, J. P. Vogel, D. F. Garvin, Y. Weng, N. McKenzie, M. W. Bevan, G. Haberer, M. Spannagl, K. Mayer (Leader), T. Rattei, T. Mitros, D. Rokhsar, S.-J. Lee, J. K. C. Rose, L. A. Mueller, T. L. York, T. Wicker (Leader), J. P. Buchmann, J. Tanskanen, A. H. Schulman (Leader), H. Gundlach, J. Wright, M. Bevan, A. Costa de Oliveira, L. da C. Maia, W. Belknap, Y. Q. Gu, N. Jiang, J. Lai, L. Zhu, J. Ma, C. Sun, E. Pritham, J. Salse (Leader), F. Murat, M. Abrouk, G. Haberer, M. Spannagl, K. Mayer, R. Bruggmann, J. Messing, F. M. You, M.-C. Luo, J. Dvorak, N. Fahlgren, S. E. Fox, C. M. Sullivan, T. C. Mockler, J. C. Carrington, E. J. Chapman, G. D. May, J. Zhai, M. Ganssmann, S. Guna Ranjan Gurazada, M. German, B. C. Meyers, P. J. Green (Leader), J. N. Bragg, L. Tyler, J. Wu, Y. Q. Gu, G. R. Lazo, D. Laudencia-Chingcuanco, J. Thomson, J. P. Vogel (Leader), S. P. Hazen, S. Chen, H. V. Scheller, J. Harholt, P. Ulvskov, S. E. Fox, S. A. Filichkin, N. Fahlgren, J. A. Kimbrel, J. H. Chang, C. M. Sullivan, E. J. Chapman, J. C. Carrington, T. C. Mockler, L. E. Bartley, P. Cao, K.-H. Jung, M. K. Sharma, M. Vega-Sanchez, P. Ronald, C. D. Dardick, S. De Bodt, W. Verelst, D. Inzé, M. Heese, A. Schnittger, X. Yang, U. C. Kalluri, G. A. Tuskan, Z. Hua, R. D. Vierstra, D. F. Garvin, Y. Cui, S. Ouyang, Q. Sun, Z. Liu, A. Yilmaz, E. Grotewold, R. Sibout, K. Hematy, G. Mouille, H. Höfte, T. Michael, J. Pelloux, D. O'Connor, J. Schnable, S. Rowe, F. Harmon, C. L. Cass, J. C. Sedbrook, M. E. Byrne, S. Walsh, J. Higgins, M. Bevan, P. Li, T. Brutnell, T. Unver, H. Budak, H. Belcram, M. Charles, B. Chalhoub, I. Baxter, Genome sequencing and analysis of the model grass Brachypodium distachyon. *Nature* **463**, 763–768 (2010).

132. S. Beier, A. Himmelbach, C. Colmsee, X.-Q. Zhang, R. A. Barrero, Q. Zhang, L. Li, M. Bayer, D. Bolser, S. Taudien, M. Groth, M. Felder, A. Hastie, H. Šimková, H. Staňková, J. Vrána, S. Chan, M. Muñoz-Amatriaín, R. Ounit, S. Wanamaker, T. Schmutzer, L. Aliyeva-Schnorr, S. Grasso, J. Tanskanen, D. Sampath, D. Heavens, S. Cao, B. Chapman, F. Dai, Y. Han, H. Li, X. Li, C. Lin, J.

K. McCooke, C. Tan, S. Wang, S. Yin, G. Zhou, J. A. Poland, M. I. Bellgard, A. Houben, J. Doležel, S. Ayling, S. Lonardi, P. Langridge, G. J. Muehlbauer, P. Kersey, M. D. Clark, M. Caccamo, A. H. Schulman, M. Platzer, T. J. Close, M. Hansson, G. Zhang, I. Braumann, C. Li, R. Waugh, U. Scholz, N. Stein, M. Mascher, Construction of a map-based reference genome sequence for barley, Hordeum vulgare L. *Sci Data* **4**, 170044 (2017).

133. T. Zhu, L. Wang, H. Rimbert, J. C. Rodriguez, K. R. Deal, R. De Oliveira, F. Choulet, G. Keeble-Gagnère, J. Tibbits, J. Rogers, K. Eversole, R. Appels, Y. Q. Gu, M. Mascher, J. Dvorak, M.-C. Luo, Optical maps refine the bread wheat Triticum aestivum cv. Chinese Spring genome assembly. *The Plant Journal* **107**, 303–314 (2021).

134. R. VanBuren, D. Bryant, P. P. Edger, H. Tang, D. Burgess, D. Challabathula, K. Spittle, R. Hall, J. Gu, E. Lyons, M. Freeling, D. Bartels, B. Ten Hallers, A. Hastie, T. P. Michael, T. C. Mockler, Single-molecule sequencing of the desiccation-tolerant grass Oropetium thomaeum. *Nature* **527**, 508–511 (2015).

135. K. M. Devos, P. Qi, B. A. Bahri, D. M. Gimode, K. Jenike, S. J. Manthi, D. Lule, T. Lux, L. Martinez-Bello, T. H. Pendergast, C. Plott, D. Saha, G. S. Sidhu, A. Sreedasyam, X. Wang, H. Wang, H. Wright, J. Zhao, S. Deshpande, S. de Villiers, M. M. Dida, J. Grimwood, J. Jenkins, J. Lovell, K. F. X. Mayer, E. E. Mneney, H. F. Ojulong, M. C. Schatz, J. Schmutz, B. Song, K. Tesfaye, D. A. Odeny, Genome analyses reveal population structure and a purple stigma color gene candidate in finger millet. *Nat Commun* **14**, 3694 (2023).

136. S. Mamidi, A. Healey, P. Huang, J. Grimwood, J. Jenkins, K. Barry, A. Sreedasyam, S. Shu, J. T. Lovell, M. Feldman, J. Wu, Y. Yu, C. Chen, J. Johnson, H. Sakakibara, T. Kiba, T. Sakurai, R. Tavares, D. A. Nusinow, I. Baxter, J. Schmutz, T. P. Brutnell, E. A. Kellogg, A genome resource for green millet Setaria viridis enables discovery of agronomically valuable loci. *Nat Biotechnol* **38**, 1203–1210 (2020).

137. J. T. Lovell, A. H. MacQueen, S. Mamidi, J. Bonnette, J. Jenkins, J. D. Napier, A. Sreedasyam, A. Healey, A. Session, S. Shu, K. Barry, S. Bonos, L. Boston, C. Daum, S. Deshpande, A. Ewing, P. P. Grabowski, T. Haque, M. Harrison, J. Jiang, D. Kudrna, A. Lipzen, T. H. Pendergast, C. Plott, P. Qi, C. A. Saski, E. V. Shakirov, D. Sims, M. Sharma, R. Sharma, A. Stewart, V. R. Singan, Y. Tang, S. Thibivillier, J. Webber, X. Weng, M. Williams, G. A. Wu, Y. Yoshinaga, M. Zane, L. Zhang, J. Zhang, K. D. Behrman, A. R. Boe, P. A. Fay, F. B. Fritschi, J. D. Jastrow, J. Lloyd-Reilley, J. M. Martínez-Reyna, R. Matamala, R. B. Mitchell, F. M. Rouquette, P. Ronald, M. Saha, C. M. Tobias, M. Udvardi, R. A. Wing, Y. Wu, L. E. Bartley, M. Casler, K. M. Devos, D. B. Lowry, D. S. Rokhsar, J. Grimwood, T. E. Juenger, J. Schmutz, Genomic mechanisms of climate adaptation in polyploid bioenergy switchgrass. *Nature* **590**, 438–444 (2021).

138. G. Sun, N. Wase, S. Shu, J. Jenkins, B. Zhou, J. V. Torres-Rodríguez, C. Chen, L. Sandor, C. Plott, Y. Yoshinga, C. Daum, P. Qi, K. Barry, A. Lipzen, L. Berry, C. Pedersen, T. Gottilla, A. Foltz, H. Yu, R. O'Malley, C. Zhang, K. M. Devos, B. Sigmon, B. Yu, T. Obata, J. Schmutz, J. C. Schnable, Genome of Paspalum vaginatum and the role of trehalose mediated autophagy in increasing maize biomass. *Nat Commun* **13**, 7731 (2022).

139. R. F. McCormick, S. K. Truong, A. Sreedasyam, J. Jenkins, S. Shu, D. Sims, M. Kennedy, M. Amirebrahimi, B. D. Weers, B. McKinley, A. Mattison, D. T. Morishige, J. Grimwood, J. Schmutz, J. E. Mullet, The Sorghum bicolor reference genome: improved assembly, gene annotations, a transcriptome atlas, and signatures of genome organization. *The Plant Journal* **93**, 338–354 (2018).

140. H. Cheng, G. T. Concepcion, X. Feng, H. Zhang, H. Li, Haplotype-resolved de novo assembly using phased assembly graphs with hifiasm. *Nat Methods* **18**, 170–175 (2021).
